# Normal aging impacts the extent and diversity of neural plasticity induced in the mouse brain with repetitive transcranial magnetic stimulation

**DOI:** 10.1101/2025.07.24.665914

**Authors:** Rebecca C S Ong, Alexander D Tang

## Abstract

Repetitive transcranial magnetic stimulation (rTMS) is an attractive tool to promote healthy brain ageing in older adults and treat age-related neurological conditions. Despite its popularity, the neurological processes and plasticity mechanisms altered by rTMS in the aged brain, and where these changes occur in the brain are unknown. Furthermore, it is not known why different rTMS protocols induce different changes in the aged brain, or why rTMS is less effective in older adults compared to younger adults. Using spatial transcriptomics, we uncovered that rTMS primarily acts on genes related to synaptic plasticity in both cortical and subcortical circuits in aged mice, but the specific changes were dependent on the brain region and even down to individual cortical layers in the motor and somatosensory cortices. Comparing our results from aged mice to young adult mice revealed that rTMS acts on a larger variety of neural plasticity mechanisms in the young adult brain, and that rTMS was less effective at altering gene expression related to neural plasticity in the aged brain, but this varied between brain regions and the protocol of rTMS applied. These findings provide a comprehensive map of the mechanisms altered by rTMS across the aged brain and highlight the need to consider the effect of ageing when optimising rTMS protocols for older populations.

## INTRODUCTION

Cognitive processes such as learning and memory are fundamental to how we perceive, interpret, and interact with the world around us. To facilitate these cognitive processes, the brain requires constant adaptation in the form of neural plasticity, whereby single cells to entire neural circuits alter their structure and function. Unfortunately, normal aging is associated with a decline and neural plasticity (Burke & Barnes, 2006) and is a risk factor for several neurological conditions (Roca, Lang, & Chassagne, 2019). As a result, there has been great effort to develop tools that can enhance cognitive processing and neural plasticity in older adults. One such tool is repetitive transcranial magnetic stimulation (rTMS) which applies rapid pulses of magnetic fields through the cranium to alter neural activity and plasticity in the brain (Tang, Thickbroom, & Rodger, 2017). Due to the non-invasive nature and excellent safety profile of rTMS, it has become an extremely popular tool to induce neural plasticity in older adults to promote healthy brain ageing (Hsu, Ku, Zanto, & Gazzaley, 2015) or as a treatment for age-related neurological conditions (stroke, Alzheimer’s disease etc.). When delivering rTMS, the same protocols are used are used irrespective of age, however, multiple reports have shown that rTMS is less effective in inducing neural plasticity in older adults relative to young adults (Opie, Cirillo, & Semmler, 2018; Opie, Vosnakis, Ridding, Ziemann, & Semmler, 2017; Todd, Kimber, Ridding, & Semmler, 2010), blunting its use as a basic tool to modulate the aged brain.

Characterising the neural plasticity mechanisms affected by rTMS in the aged brain, and how this varies between different brain regions and stimulation protocols is needed to better understand and optimise the use of rTMS in older adults. Spatial transcriptomics is a powerful method to study the molecular mechanisms of rTMS, and in the mouse brain, can map the changes to cortical and subcortical circuits in the same brain section. Spatial transcriptomics has recently been used to characterise the effect of two common rTMS protocols, intermittent and continuous theta burst stimulation (iTBS and cTBS respectively), across the young adult mouse brain(R. C. S. Ong & Tang, 2024). In that study, rTMS was found to act on a diverse range of neural plasticity mechanisms (synaptic and non-synaptic) and general cellular functions, with the specific changes varying between brain regions, cortical layers, and the protocol of theta burst stimulation applied. Here, we use spatial transcriptomics to uncover and map the neural plasticity mechanisms induced across the aged mouse brain following iTBS and cTBS to the sensorimotor cortex. In addition, we compare our findings to the transcriptomic changes that occur in young adult mice to uncover why current protocols of rTMS are less effective in older adults.

## METHODS

### Animals

Twenty-seven aged (13-16 months) male C57BL/6J mice, supplied by the Animal Resource Center (Murdoch, Australia), were individually housed upon arrival, and maintained under a standard 12 h light/dark cycle with *ad libitum* access to food and water. All animal procedures were approved by the University of Western Australia Animal Ethics Committee (AEC No. 2022/ET000022) and performed in accordance with the National Health and Medical Research Council (NHMRC) of Australia Code of Practice for the Care and Use of Animals for Scientific Purposes.

### rTMS parameters and administration

A single session of rTMS or sham was delivered to awake and non-restrained mice. All animals were handled for a minimum of 5 mins per day for 7 consecutive days prior to active rTMS or sham delivery to acclimatise the animals to the coil and noise from the stimulator and prevent any stress response.

To deliver focal stimulation appropriate for the mouse brain, custom built rodent-specific circular coils were used (8mm in height and width) ( Tang et al., 2016) to deliver bilateral stimulation to the sensorimotor cortices. Stimulation was delivered as monophasic pulses (300 μs rise time, 100 μs fall time) produced by a waveform generator (33500B; Agilent Technologies, USA) connected to a bipolar programmable power supply (BOP 100-4 M; Kepco Inc, USA). For all experiments, mice received a single session of active rTMS or sham stimulation. Active rTMS was delivered in the form of theta burst stimulation (TBS), a pattern consisting of 3 pulses delivered at 50 Hz repeated in 5 Hz intervals. Two paradigms of TBS were tested: intermittent TBS (iTBS; 2s train of TBS repeated once every 10s for a total of 600 pulses in 192s) and continuous TBS (cTBS; a continual train of TBS for a total of 600 pulses in 40s). An input voltage of 60.2V peak-to-peak was used to ensure the coil temperature was < 37°C throughout the duration of stimulation, preventing any burning of the animal and coil. Using a Gauss meter (GM08; Hirst Magnetic Instruments, UK), a peak electromagnetic field intensity of 210mT was measured at the base of the coil, estimated to induce an electric field of ∼ 30V/m at the cortical surface based off our previous electric field modelling (R. C. S. Ong & Tang, 2024). Sham stimulation lasted 192s and was delivered under the same conditions as rTMS, but the coil was disconnected from the stimulator.

### 10X Visium spatial transcriptomics

#### Sample preparation & sequencing

For spatial transcriptomic experiments, 4 mice were used for each iTBS, cTBS, and sham stimulation group. All animals were euthanised 3 hours following rTMS stimulation using a two-step procedure of methoxyflurane inhalation (Clifford Hallam Healthcare) and an overdose intraperitoneal injection of sodium pentobarbitone (Lethabarb, Virbac). Cardiac perfusion of ice-cold saline (20 mL, 0.9%) was performed immediately prior to the removal of the brains. Following cardiac perfusion with ice-cold saline, whole brains were immediately removed and dissected along the midline. Right hemispheres from each brain were then placed in an embedding mould (1 cm x 1 cm) prepared with a thin layer of optimal cutting temperature (OCT) embedding medium, with further OCT added to fully cover the tissue. The embedding mould was then placed in liquid nitrogen-cooled isopentane for approximately 1-2 mins to snap-freeze the tissue.

Fresh-frozen tissues were cryosectioned coronally (approx. 0.26 – 0.74 mm anterior from bregma; approx. Allen brain reference atlas coronal section 54) at 10μm thickness onto designated capture regions on Visium Spatial Gene Expression (GEX) Slides and were stored at −80°C prior to further processing. Mounted tissues were fixed with methanol and stained with hematoxylin and eosin (H&E) to visualise tissue morphology as reference for the sequencing data. Stained sections were imaged using a Nikon Ti2 inverted microscopy at 10x magnification (numerical aperture = 0.45, 2424 x 2424-pixel resolution). Tissue optimisation slides were then incubated with a permeabilisation enzyme for 12 minutes to release poly-A messenger RNA (mRNA) from the tissue onto the slide. This optimal tissue permeabilisation timepoint was selected based on our previous spatial transcriptomics work using mice of the same species and sex at 12-weeks old (i.e., male C57BL/6J mice)(R. C. S. Ong & Tang, 2024). mRNA captured on spatially barcoded probes on the slide then underwent reverse transcription and second-strand synthesis, generating cDNA strands used for library construction. In brief, cDNA was released from slides into individual tubes and amplified for 16 cycles using a Thermal Cycler (Biorad T-100). Sequencing-ready cDNA libraries were constructed according to the manufacturer’s protocol whereby cDNA from each sample underwent fragmentation, end repair and A-tailing, adaptor ligation, and sample indexing (Dual Index Kit TT Set A, 10X Genomics) and amplification. Quality of all dual-indexed, paired-end cDNA libraries were checked on an Agilent TapeStation prior to sequencing. Libraries were submitted to the Australian Genome Research Facility (Melbourne, Australia) and were sequenced on the 10B flow cell (NovaSeq X Series 10B Reagent Kit, 300 cycles) using a NovaSeq X instrument (Illumina). Samples were sequenced at a depth of approximately 212 million read-pairs per sample (Data S1). Sequencing read configurations was performed according to the Visium protocol: read 1, 28bp; i7 index, 10bp; i5 index, 10bp; read 2, 90bp.

#### Spatial transcriptomics data processing

Raw sequencing BCL data files were demultiplexed and converted into FASTQ files using bcl2fastq (v2.20). The 10X Genomics SpaceRanger (v.2.1.1) software was used to align FASTQ files to the *Mus musculus* mm10 reference genome and match the sequenced data with the corresponding H&E tissue images. Unique molecular identifier (UMI) counts were obtained, reflecting the number of genes identified in each spot within the spatial fiducial frame. Only spots underlying the tissue section were used for subsequent analysis. Further analysis was performed using R (v4.1.0) where raw counts and images from all samples were integrated using the spatial transcriptomics toolkit *STUtility* (v.1.1.1) (Bergenstråhle, Larsson, & Lundeberg, 2020). A combination of functions in the *STUtility* package and *Seurat* (v4.0.3)(Hao et al., 2021) were used to analyse the data. As a quality control measure, spots expressing more than 30% of mitochondrial and ribosomal genes were removed as they are commonly used as an indicator of sample quality and a reflection of high levels of cell death/lysis (Márquez-Jurado et al., 2018). Remaining reads from samples were then batch corrected and normalised using the Seurat built-in *SCTransform* function, to account for differences in sequencing depth between samples across and within spatial transcriptomics slides. To visualise the spatial gene expression data based on spots that contained similar transcriptional profiles, the Seurat built-in *FindNeighbours* function was used to construct a shared-nearest-neighbours graph using the first 16 principal component (PC) dimensions before spots were clustered using the *FindClusters* function with a 0.4 clustering resolution. For the re-clustering analysis, clusters corresponding to the cortex were subsetted and spots were re-clustered with a resolution of 0.4, whilst clusters corresponding to the striatum were subsetted and re-clustered with a resolution of 0.2. Visual comparisons to the Allen Mouse Brain Atlas were performed to annotate each cluster guided by their anatomical location.

To identify genes differentially expressed following stimulation in each of the brain regions, compared to sham stimulation control, Visium clusters that corresponded to each of the different regions were first subsetted prior to differential expression analysis. The Seurat *FindMarkers* function was then implemented, and from the list of differentially expressed genes with a p.adj value (Bonferroni correction) less than/equal to 0.05 (Data S2 and S3), significant genes of interest were determined by further filtering for genes that has an absolute log2FC greater than 0.25. Of the significant differentially expressed genes, each was manually assigned into one/or more of seven categories relating to different neural processes of interest (synapse and/or synaptic plasticity, neurotransmitter, intrinsic plasticity and/or membrane excitability, oligodendrocyte and/or myelin, neurogenesis, neurotrophin, and inflammation/apoptosis) using existing literature to known functions. Gene ontology (GO) enrichment analysis was performed using the *clusterProfiler* (v4.2.2) (Yu, Wang, Han, & He, 2012) and the GO database (Ashburner et al., 2000). Data plots were generated with ggplot2 (v3.3.3) (Wickham, 2016) or visualisation tools in STUtility (e.g., *STFeaturePlot*).

### Bulk RNA-sequencing

#### Sample preparation & sequencing

For bulk RNA-sequencing, 5 mice were used for each iTBS, cTBS, and sham stimulation group. All animals were euthanised using the same procedures as performed for spatial transcriptomics, stated above. Right sensorimotor cortices were dissected for all animals across each stimulation group and RNA for each individual tissue sample was extracted using the RNeasy Mini Kit (Qiagen) and RNase-Free DNase Set (Qiagen) following standard manufacturer’s instructions. RNA samples were quantified using a LabChip GX Touch Nucleic Acid Analyser and were all verified to have a minimum RNA integrity number (RIN) of > 7 required for sequencing.

Library preparation and whole transcriptome sequencing of all RNA samples was performed by the Australian Genome Research Facility (Melbourne, Australia). In brief, ribosomal RNAs (rRNA) were depleted using the Illumina Ribo-Zero Plus rRNA Depletion Kit (Illumina) prior to the generation of cDNA libraries using the TruSeq Stranded mRNA Total Library Prep Kit (Illumina). Libraries were pooled prior to sequencing. Samples were sequencing on the Illumina NovaSeq 6000, generating 50 million, 150 base-pair (bp) paired-end reads using the Illumina DRAGEN BCL Convert 07.021.645.4.0.3 pipeline. Libraries for each sample were run in duplicate (i.e., over two sequencing lanes).

#### Bulk RNA-sequencing data analysis

Raw RNA-sequencing FASTQ files from each sample underwent quality checks to identify any variability amongst sequencing replicates using FastQC (v.0.11.9) (Chen, Zhou, Chen, & Gu, 2018). Replicates were concatenated and the first 15bp of reads were trimmed using the *fastq* package. Trimmed reads were aligned to the ENSEMBL *Mus musculus* GRCm39 reference genome using STAR aligner (v.2.7.10a) (Dobin et al., 2013). In R (v.4.1.0), reads were quantified, and a raw counts table was generated using GenomicAlignments (v.1.26.0)(Lawrence et al., 2013). All sample read counts were normalised using the variance stabilising transformation (VST) in DESeq2 (v.1.30.1) (Love, Huber, & Anders, 2014). Differential gene expression analysis between rTMS treated (iTBS and cTBS) animals and sham controls was performed using the standard DESeq2 analysis pipeline. In brief, the Wald test was used to generate *P*-values that were adjusted for multiple comparisons through the Benjamini-Hochberg false discovery rate correction (Benjamini & Hochberg, 1995). Genes with an adjusted *P*-value (p.adj) of less than 0.05 and had an absolute log2 fold difference (log2FC) that was greater than/equal to one were considered to be statistically significant.

## RESULTS

### Spatial transcriptomics uncovers cortical region and layer dependent neural plasticity mechanisms of subthreshold theta burst rTMS in the aged brain

To determine whether rTMS has a large gross effect on the cortex, we first used whole transcriptome bulk RNA-sequencing from dissected sensorimotor tissue (i.e. motor and somatosensory cortex dissected together) (Fig. 1a). Principal component analysis (PCA) revealed divergent expression patterns within treatment groups with no distinct clustering of samples (Fig. S1a). Given that all animals were habituated to handling, TMS coil, and machine noise prior to the experiment, we believe this variability to be due to inherent biological variability despite using inbred mice, and littermates where possible. Reflecting this within-group variability, there were no significant changes in gene expression following cTBS (Fig. S1b) and only 1 differentially expressed gene follow iTBS, with a downregulation of lipocalin 2 *lcn2*, encoding for a neurotoxin highly expressed in activated glial cells (Jung & Ryu, 2023; Kim et al., 2023) (Fig. S1c).

**Fig. 1:**
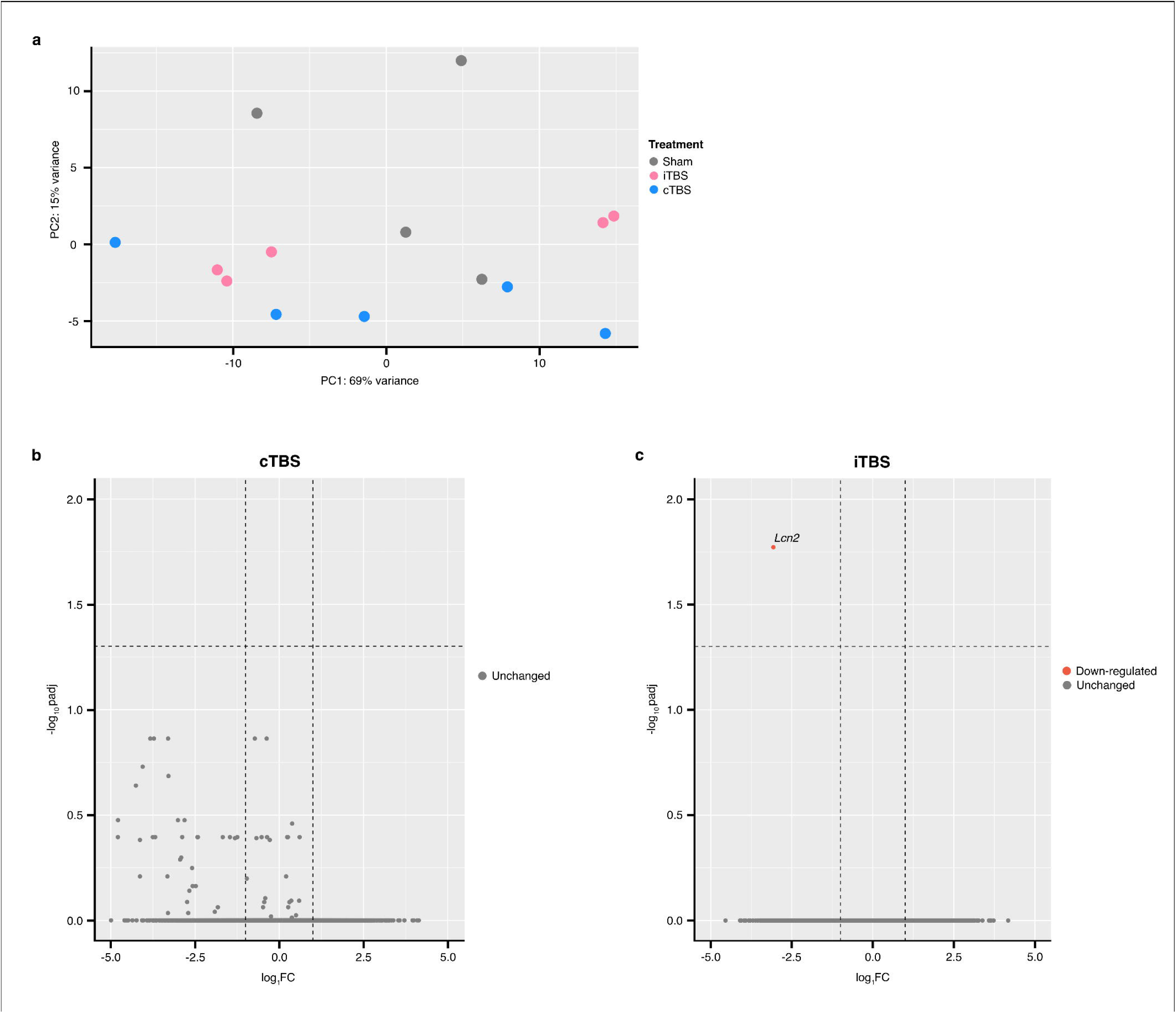
Primary motor (M1) cortex layer specific gene expression changes induced 3 h following subthreshold rTMS protocols in the aged mouse brain. **a,** Schematic of the experimental design to assess changes in transcriptional profiles of brain regions in aged mice following rTMS neuromodulation. **b,** Subclustering of Visium spots corresponding to the M1 cortex. Visium samples were aligned to the Allen Reference Mouse Brain Atlas to identify the M1 region within each Visium sample. Four distinct clusters corresponding to the different cortical layers (Ctx L1-L6) were resolved from unsupervised clustering of Visium spots in the M1 cortex. **c,** Barplot of the number of up- and down-regulated significant differentially expressed genes (DEGs) (p.adjusted ≤ 0.05, absolute log2 fold-change > 0.25) in each M1 cortical layer. Due to the large variability in the number of Ctx L1 spots between samples, no differential expression analysis was performed on this region. The overlap of DEGs between each cortical layer following **d,** iTBS and **e,** cTBS are displayed as venn diagrams. Each gene was assessed and where applicable, categorised to various neural plasticity processes based on their known function in the literature. **f,** Spatial gene expression maps from representative sham, iTBS, and cTBS samples for various genes are displayed. Coloured columns used to visualise the extent of regulation for each gene within the different cortical layers following each stimulation protocol, with the intensity of blue and red indicating an increase and decrease in expression (log2FC) respectively. **f,** Table denoting the number of DEGs that were identified to be involved in one of the different categories of neural plasticity processes of interest. Genes in **d,** and **e,** are colour coded as indicated in the table.

It was recently shown that the higher resolution of spatial transcriptomics can uncover changes to gene expression in different brain regions, following rTMS down to the changes in each cortical layer which cannot be detected with low resolution bulk RNA-sequencing (R. C. S. Ong & Tang, 2024). Therefore, to determine the changes specific to different brain regions, we aligned our H&E images with the Allen Brain Atlas which allowed us to subset the spots in the motor and somatosensory cortices (R. C. S. Ong & Tang, 2024). Spots aligning to either the motor or somatosensory cortex were then sub-clustered to distinguish between the different cortical layers specific to each cortical region (Fig. 1b and 2a).

**Fig. 2:**
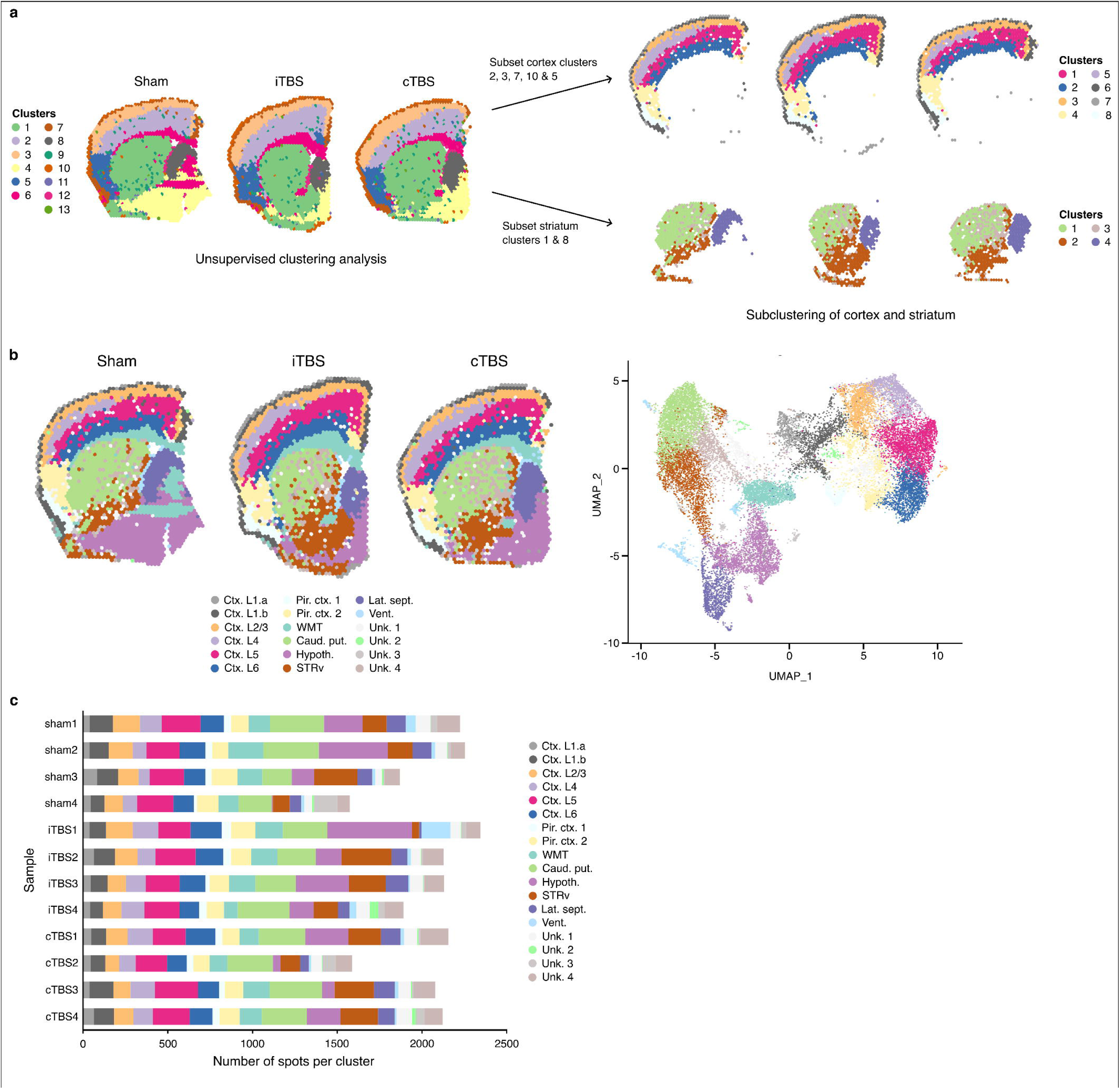
Somatosensory (SS) cortex layer specific gene expression changes induced 3 h following subthreshold rTMS protocols in the aged mouse brain. **a**, Subclustering of Visium spots corresponding to the SS cortex, as defined using alignment of Visium samples to the Allen Reference Mouse Brain Atlas. Seven different clusters were resolved, of which, six were annotated as the different cortical layers of the SS cortex (Ctx L1-L6). Ctx L1 spots were resolved as two clusters and were variable between samples, thus no further analysis was performed on this region. **b,** Barplot displaying the number of up- and down-regulated significant differentially expressed genes (DEGs) (p.adjusted ≤ 0.05, absolute log2 fold-change > 0.25) in each SS cortical layer. Overlapping DEGs between each cortical layer following **c,** iTBS and **d,** cTBS are displayed as venn diagrams. Genes were categorised into different neural plasticity processes, where applicable, based on their known function in the literature. Categorised genes are colour coded as displayed in **e**, the table, denoting the number of DEGs found for each neural plasticity process. **f,** Spatial gene expression maps taken from representative sham, iTBS, and cTBS samples for various genes with accompanying coloured columns used to visualise the extent of regulation for each genes within the different cortical layers.

Sub-clustering of the motor cortex revealed four distinct clusters that aligned with the different cortical layers typically observed within this brain region (Fig. 1b). Using the Allen Mouse Brain Atlas as reference, these clusters were annotated as cortical layer 1 (Ctx. L1), cortical layer 2/3 (Ctx. L2/3), cortical layer 5 (Ctx. L5), and cortical layer 6 (Ctx. L6). We observed a large variability in the number of spots within the Ctx. L1 cluster, presumably due to mechanical damage from sectioning, and as such, we chose to exclude Ctx. L1 from any further analysis. Across the remaining cortical layers, the number of gene expression changes decreased with depth (Ctx. L2/3; 35 DEGs following iTBS, and 16 DEGs following cTBS) to deepest (Ctx. L6; 9 DEGs following iTBS, and 10 DEGs following cTBS) that was seen in both TBS protocols (Fig. 1c). Comparing the DEGs identified in each cortical layer, there were gene expression changes induced by stimulation that were cortical-layer specific, as well as genes that were altered in expression across multiple cortical layers (Fig. 1d and 1e). We identified genes involved in various different cellular processes, some of which were specific to certain cortical layers and stimulation protocol (Fig. 1f, and Data S4 and S5). For example, modulation of transcription factor genes (e.g., *Nr4a1*, and *Junb*) were only observed following iTBS and was confined to layer 2/3 of the motor cortex (Fig. 1f). Notably, *Junb* is an immediate early gene that has been shown to have increased expression as early as 5 minutes following rTMS (Hwang, Choi, Bang, Park, & Oh, 2022). Across both TBS protocols, changes in the expression of genes known to play a role in inflammation (e.g., *Ikbkb*), mitochondrial function (e.g., *Cox7c*, and *Dusp26*), and cytoskeletal organisation (e.g., *Actb*, and *Tuba1b*) were observed, all of which had varying levels of expression change and regulation specific to particular motor cortex layers (Fig. 1f). To further assist in the interpretation of what different cellular processes altered, we manually assigned genes into several neural plasticity categories known to be affected by rTMS^8^. In the motor cortex, we found that both stimulation protocols predominantly modulated the expression of genes related to synaptic plasticity across all cortical layers (Fig. 1g). Genes encoding for proteins associated with synaptic function (e.g., *Syngap1*, and *Cdk5r2*) and synaptic vesicles (e.g., *Syngr1*, *Dnm1*, and *Cplx2*) were downregulated within specific layers of the motor cortex but these specific gene changes were found only following iTBS.

Sub-clustering of the somatosensory cortex revealed five distinct clusters that aligned with cortical layers 1, 2/3, 4, 5 and 6, however cortical layer 1 was excluded from the analysis for the same reasons it was excluded for the motor cortex analyses (Fig. 2a). We found that across layers 2/3 to 6 in the somatosensory cortex layers, iTBS had an equal or greater influence on gene expression compared to cTBS, with a large proportion of genes found to be upregulated within this cortical region following stimulation (Fig. 2b). Of the cortical layers, both stimulation protocols induced the least number of gene expression changes within cortical layer 5 (5 DEGs following iTBS and 4 DEGs following cTBS; Fig. 2c and 2d), though across the board, there was a lower number of DEGs following both protocols in the somatosensory cortex compared to the motor cortex. Assessing the known function of each DEG, we found that iTBS induced changes across numerous cellular functions including genes associated with synaptic plasticity (e.g., *Adcy1*, and *Camk2n1*), cellular structure (e.g., *Tuba1b*), and mitochondria (e.g., *Ndufa4*) (Fig. 2e and 2f, and Data S6 and S7). We also found changes in several ribosomal genes and non-coding RNAs including *Malat1*, a long non-coding RNA (lncRNA) known to regulate gene transcription particularly relating to synapse formation (Bernard et al., 2010; Wang et al., 2022) (Fig. 2f). In contrast, cTBS largely only affected genes relating to synaptic plasticity-related processes across the somatosensory cortex, some of which were similarly regulated following iTBS (e.g., *Camk2n1*) whilst others were unique to cTBS (e.g., *Ncdn*) (Fig. 2e and 2f).

### Subthreshold theta burst rTMS induces neural plasticity in subcortical brain regions

Neuroimaging studies in human participants show rTMS to the cortex induces changes in both cortical and sub-cortical brain structures, however, these methods cannot determine the specific neural plasticity changes within each region and how they differ between regions. To that end, we looked at the gene expression profiles in subcortical brain regions and white matter tracts that were identified through an unsupervised clustering-based analysis of the entire coronal section and further subclustering of certain regions (e.g., striatum) (Fig. S2a and S2b). In addition to a cluster for white matter tracts (WMT), clusters for the caudate putamen (Caud. put.), ventral striatum (STRv), and lateral septum (Lat. sept.) were included for differential gene expression analysis given that the number of spots per cluster were comparable across samples (Fig. S2c).

Comparing the gene expression profiles of the various subcortical structures, we found comparable numbers of genes significantly differentially expressed following iTBS and cTBS across all regions (Fig. 3a). To assess how the effect of stimulation differs between cortical and subcortical structures, we also looked at the number of DEGs in the different layers of the sensorimotor cortex as a whole. Surprisingly, despite the stimulation being targeted towards the sensorimotor cortex and having the highest induced electric field intensity, the number of DEGs in the subcortical brain regions was similar to the cortical regions, and in some cases, were greater than the cortex (Fig. 3a). Specifically, following iTBS, the number of DEGs amongst cortical layers ranged from 10 (Ctx. L4) to 15 (Ctx. L5) compared to sub-cortical structures ranging from 8 (STRv) to 23 (WMT). Following cTBS, the number of DEGs amongst cortical layers ranged from 7 (Ctx. L6) to 11 (Ctx. L2/3) compared to sub-cortical structures ranging from 10 (STRv) to 23 (Lat. sept.). Looking at the directionality of gene expression changes, there was an approximately equal proportion of genes that were up-, and down-regulated within the subcortical structures, except for the white matter tracts whereby both stimulation protocols largely induced an increase in gene expression (Fig. 3a). Following iTBS and cTBS, we found changes in the expression of genes that were stimulation protocol-specific within the various subcortical structures (Fig. 3b-e). However, we also found some overlap in the genes altered following both protocols, with some gene expression changes that were consistently observed regardless of stimulation protocol and brain region (Fig. 3b-e). Notably, *Bc1*, a non-protein-coding gene well characterised to be involved in experience-dependent synaptic plasticity (Briz et al., 2017; Chung, Dahan, Alarcon, & Fenton, 2017; Mus, Hof, & Tiedge, 2007; Wu et al., 2024) was found to be downregulated in all subcortical brain regions following both iTBS and cTBS, with the exception of the lateral septum where only cTBS modulated its expression. We also observed the downregulation of several other synaptic plasticity-related genes (e.g., *Slc17a7, Malat1*, and *Kalrn*; Fig. 3b-f and Data S8) that were uniquely modulated depending on the stimulation protocol and subcortical structure. Interestingly, *Ttr*, which codes for the transthyretin protein previously found to have a decreased expression in the hippocampus of memory-impaired aged mice (Brouillette & Quirion, 2008), was found to have differing changes in expression between brain regions following the same stimulation protocol, with increases in its expression found in the white matter tracts and caudate putamen, whilst it was downregulated in the lateral septum following iTBS (Fig. 3b-f and Data S8). Through assessing the functions of each significant DEG in the literature, we also identified changes in genes relating to several other neural processes alongside changes to numerous ribosomal genes (Fig. 3f and Data S8). Whilst the number of genes that could be categorised into one of our seven neural plasticity process categories were limited, we found a downregulation of genes relating to intrinsic plasticity (e.g., *Atp1a3*, and *Kcnq2*) in the ventral striatum and lateral septum, and an upregulation of inflammatory-related genes (e.g., *Snhg8*, and *Banf1*) within the white matter tracts, but only following iTBS (Fig. 3f). In contrast, while there was no evidence that cTBS induced changes in the expression of genes relating to intrinsic plasticity and inflammation, there was an upregulation of myelin-related genes (e.g., *Bcas1*, *Mbp*, and *Mobp*) within the within the white matter tracts that was unique to cTBS (Fig. 3f).

**Fig. 3:**
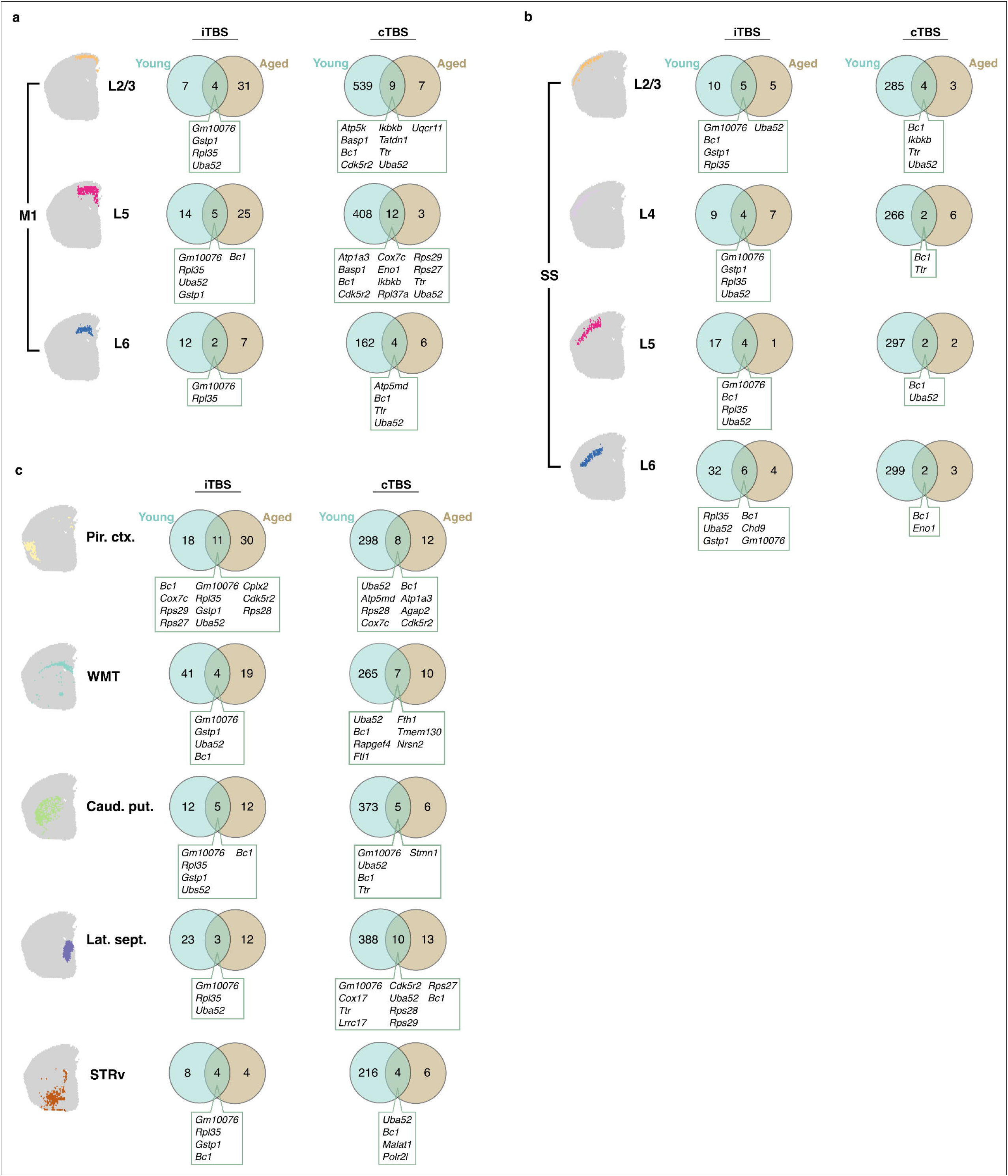
Sub-cortical gene expression changes induced 3 h following a single session of subthreshold rTMS in the aged mouse brain. **a**, Number of genes that had a significant increase or decrease in expression (p.adjusted ≤ 0.05, absolute log2 fold-change > 0.25) in each annotated brain region. Venn diagrams of the overlap of significant differentially expressed genes (DEGs) following iTBS and cTBS and the corresponding spatial plots of the sub-cortical regions are displayed: **b**, white matter tracts, **c**, caudoputamen, **d**, striatum ventral region, and the **e**, lateral septum. Genes were categories into one of seven different neural plasticity processes of interest, as summarised in f. **f,** Table summarising the number of DEGs found in each neural plasticity process.

### rTMS induced neural plasticity is less diverse in the aged mouse brain compared to young adults

To determine if the neural plasticity induced across the brain following iTBS and cTBS are affected by normal ageing, we compared our spatial transcriptomic data from the aged mouse brain to a published dataset from young adult mice (12 weeks old) that received iTBS or cTBS to the sensorimotor cortices (Ong & Tang, 2025). In total, there were 12 brain regions across both age groups for comparison (Fig. 4a-c).

**Fig. 4:**
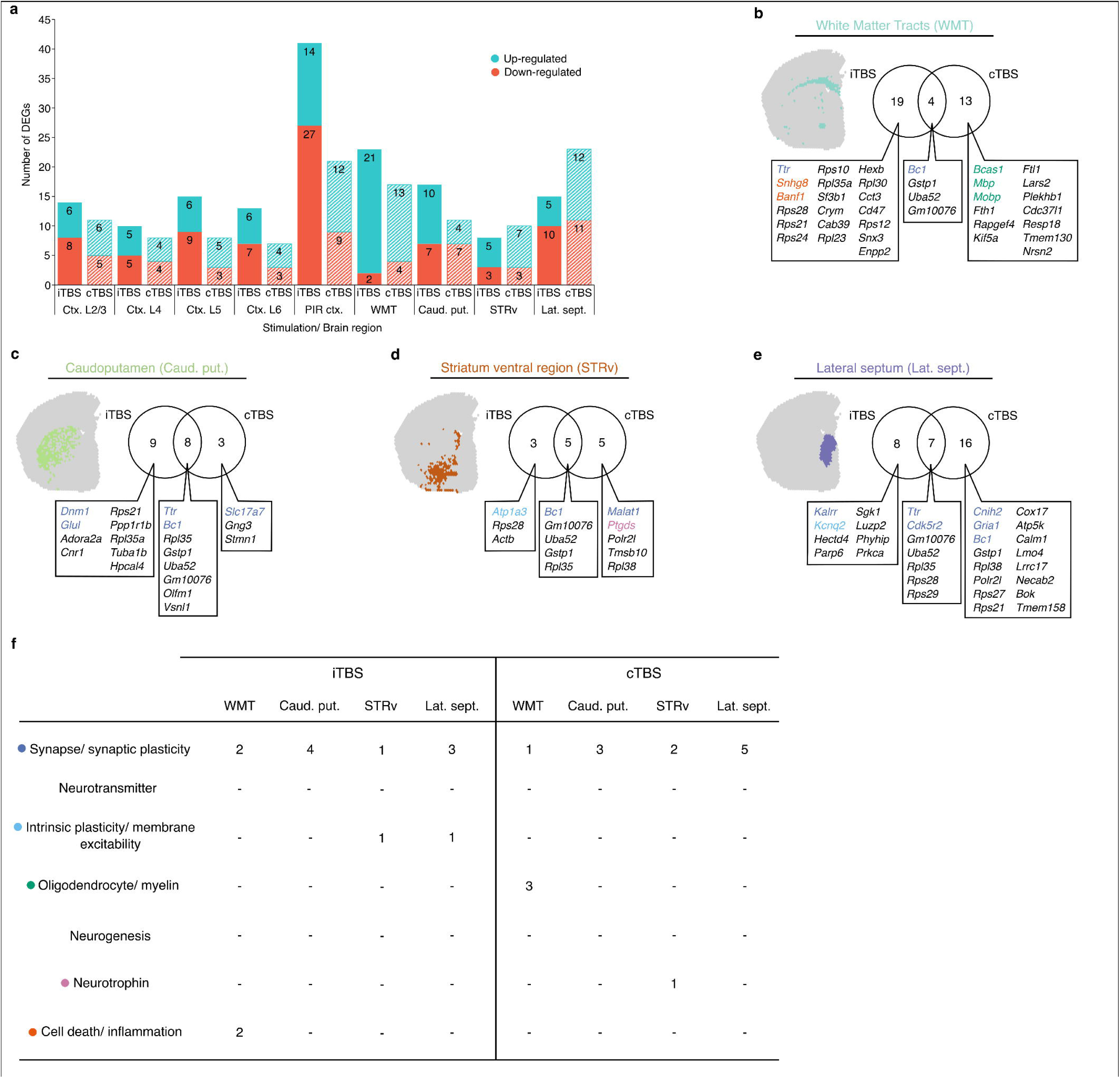
Comparison of subthreshold rTMS-induced gene expression changes in young adult and aged mice assessed 3 h following stimulation. Venn diagrams displaying the overlap between genes in young adult and aged that had a significant differential expression following iTBS, and cTBS neuromodulation within the **a**, layers of the primary motor cortex (M1), **b**, layers of the somatosensory cortex (SS), **c**, and sub-cortical structures.

In layers 2/3 and 5 of the motor cortex, iTBS induced a greater number of significant gene expression changes in the aged brain compared to the young adult brain but was slightly decreased in layer 6 (Fig. 4a). Interestingly, in the cTBS groups there was a sizeable reduction in the number DEGS in the aged brain compared to young adults across all motor cortex layers (Fig. 4a). Comparing the list of significant DEGs within each layer of the motor cortex, there was little overlap between the age groups for each stimulation protocol, suggesting age-dependent mechanisms-of-action of iTBS and cTBS. Of the overlapping genes within the motor cortex between age groups, we identified genes related to synaptic plasticity (e.g., *Basp1*, *Bc1*), intrinsic plasticity (e.g, *Atp1a3*), ribosomal function (e.g., *Uba52,* and genes beginning with ‘*Rps*’ or ‘*Rpl*’), and mitochondria function (e.g., *Atp5k*, and *Uqcr11*).

In the somatosensory cortex, both iTBS and cTBS induced a smaller number of DEGs in the aged brain compared to the young adult brain (Fig. 4b). Similar to the motor cortex, the reduction in DEGs was larger in the cTBS group suggesting that the effects of cTBS is impacted by normal aging to a greater extent than iTBS (Fig. 4b). Looking at the overlap in DEGs irrespective of age and stimulation protocol, we identified a small number of genes that were altered in expression independent of age. These includes genes involved in synaptic plasticity (e.g., *Bc1*, and *Ttr*), ribosomal function (e.g., *Uba52*, and *Rpl35*), DNA binding (e.g., *Eno1*, and *Chd9*) and inflammation (e.g., *Ikbkb*). In addition, there were unique DEGs between the age groups, suggesting age-dependent effects of TBS on the somatosensory cortex.

Across deeper brain regions (i.e., piriform cortex and subcortical structures), iTBS and cTBS had a greater effect on gene expression in the young adult brain compared to the aged brain, except for iTBS in the piriform cortex and caudate putamen suggesting that there are some brain regions where normal brain aging does not impact the efficacy of rTMS. The difference in the number of genes regulated in the young adult vs. aged brain was substantially higher for cTBS (Fig. 4c) suggesting that just like the cortex, normal aging has a greater impact on cTBS neuromodulation. Interestingly, for iTBS there was one DEG (*Gm10076*) that had altered expression across all brain regions, cortical and sub-cortical, regardless of age, but the function of this transcript is currently unknown (McKellar et al., 2023).

Comparing the categories of neural plasticity processes that were identified from the significant DEGs in various brain regions across age groups, both TBS protocols primarily induced changes in synaptic plasticity-related genes with limited influence seen on other plasticity mechanisms (e.g., intrinsic and myelin plasticity) in the aged brain. In young adults, iTBS primarily affected the expression of synaptic and myelin plasticity genes in both cortical and sub-cortical regions. In contrast, cTBS in young adults altered a greater number of DEGs and more neuronal processes/plasticity mechanisms across the brain relative to iTBS, which was greatly reduced in our aged mice cohort.

To determine whether the differences in the number of DEGs altered by rTMS in the young adult and aged brain was due to an impact of ageing on baseline gene expression, we computed differential gene expression between sham adult and sham groups (i.e. DEGs altered by ageing irrespective of rTMS) for each brain region. We then compared these lists to the genes that were differentially expressed following rTMS in the young adult brain, but not in the aged brain (Data S9). The analysis showed that genes altered by ageing alone had some overlap to the genes exclusively altered by rTMS in the young brain, ranging from no overlap at all following iTBS in L2/3 of M1 and SS, to 176/374 genes in the caudoputamen following cTBS. Generally, the amount of overlap was greater in subcortical brain regions but the exact changes was also dependent on the rTMS protocol, further suggesting protocol dependent effects of rTMS. Given that there was only a small amount of overlap in the genes altered in expression by age alone and those exclusively altered in the young adult brain, our findings suggest that age-related baseline gene expression cannot fully explain the differences in the number of DEGs altered by rTMS in the aged and young adult brain.

## Discussion

Using spatial transcriptomics, we mapped the neural plasticity and biological processes altered in cortical and subcortical brain regions following two common protocols of rTMS targeted to the aged sensorimotor cortex. Given that the maximum induced electric field from our rTMS delivery occurs in both the motor and somatosensory cortices, it could be expected that for the same protocol of rTMS, similar changes in gene expression between the motor and somatosensory would be seen. However, while we found further evidence that iTBS and cTBS act on synaptic plasticity in the motor and somatosensory cortices (Tang et al., 2021), as well as all subcortical brain regions investigated, changes to other plasticity mechanisms and biological processes were cortical region and protocol dependent. For example, both protocols of rTMS led to a greater number of DEGs in the aged motor cortex compared to somatosensory cortex and a greater diversity in the plasticity and biological processes altered. Similarly, there were only a few genes that were altered in expression in the motor and somatosensory cortices by both protocols of rTMS (e.g., *BC1* related to synaptic plasticity) and all other DEGs showed a complex pattern of altered expression that differed between cortical region, cortical layer, and rTMS protocol. These results show that while rTMS may be acting on common neural plasticity mechanisms at a high level (e.g., synaptic plasticity), the genes and biological processes leading to these common mechanisms vary substantially between neural circuits.

By comparing our spatial transcriptomic data in the aged brain to an identical dataset in young adult mice, we were able to provide the first characterisation of the effect of normal aging on rTMS neuromodulation at the molecular level. Unexpectedly, we did not find a consistent effect of aging on the extent of gene expression changes and instead found the effect to be protocol and brain region dependent. Specifically, there was a large and consistent effect of aging on cTBS neuromodulation with less changes to gene expression across all brain regions and cortical layers in the aged brain, but the effect following iTBS was inconsistent with some brain regions and cortical layers having a greater number of DEGs in the aged brain. In some brain regions, the difference in the number of DEGs following rTMS between young adult and the aged differed by two orders of magnitude (e.g., Layer 2/3 with cTBS). The stark differences in the number of DEGs altered by rTMS in the young adult and aged brain is likely to impact rTMS efficacy as our subsequent analysis of the types of neural plasticity mechanisms and biological processes affected by rTMS showed both protocols of rTMS led to less diversity compared to young adults. For example, there was a lack of changes to intrinsic plasticity mechanisms in the aged brain that would impact the efficacy of rTMS to alter neuronal excitability and promote activity dependent plasticity. Similarly, iTBS altered the expression of synaptic and oligodendrocyte/myelin plasticity related genes in the cortex and white matter tracts in young adults but not oligodendrocyte/myelin plasticity in any region of the aged brain. Given that rTMS is not always used for the purpose of inducing synaptic plasticity, such as oligodendrocyte plasticity in multiple sclerosis (Nguyen, Makowiecki, et al., 2024; Nguyen, Zarghami, et al., 2024), our results show the need to carefully consider both age, protocol, and desired neural circuit targets before applying rTMS.

Consistent with our previous work in adult mice (Ong & Tang, 2024), we found that rTMS targeted to the cortex results in robust changes to gene expression in underlying white matter and subcortical regions in aged mice. We have speculated that this neuromodulation is indirect (i.e. not due to the E-field at the distant regions) and most likely due to rTMS-induced changes to neuronal activity at the site of stimulation leading to activity-dependent plasticity in synaptically connected brain regions. Whether the downstream changes are incidental or functional remains to be confirmed but we have previously shown that iTBS of similar intensity to the adult sensorimotor improves motor learning and skill accuracy (Tang et al., 2018), processes that involve plasticity in several brain regions including the white matter tracts (Xin & Chan, 2020), striatum (Cataldi, Stanley, Miniaci, & Sulzer, 2022), and caudate putamen (Badea et al., 2019) regions where we found changes to gene expression. However, it is worth noting that stimulation with suprathreshold intensities may engage additional mechanisms to what we propose for subthreshold stimulation, like the coincidence of backward propagating action potentials in the post-synaptic neuron arriving at the synapse in short latency of the forward propagating action potential in the pre-synaptic neuron to induce synaptic plasticity (Lenz et al., 2015).

Our transcriptomic data in mice supports previous rTMS studies using older adults which show that current protocols of rTMS are less effective in inducing neural plasticity in the aged brain (Opie et al., 2018, 2017; Todd et al., 2010). This raises the question of whether the difference in efficacy is due to a decreased capacity of the aged brain to undergo plasticity, or whether the current protocols of rTMS are just not optimised for the aged brain. For plasticity mechanisms such as synaptic plasticity (Ryan, Guévremont, Luxmanan, Abraham, & Williams, 2015) and oligodendrocyte/myelin plasticity (Hill, Li, & Grutzendler, 2018) that are known to have an age-related decline, age is likely to impact rTMS efficacy. However, since the effect of ageing was not the same for iTBS and cTBS, and in some areas iTBS induced greater plasticity in the aged brain, our data suggest that manipulating the rTMS parameters can also lead to improved efficacy. Therefore, we argue that while intrinsic biological factors are likely to contribute to the reduced efficacy of current rTMS protocols, improved efficacy could be achieved by altering the rTMS parameters specifically for the aged brain (e.g., stimulation frequency, number of pulses etc.). In addition,

While our study provides novel insight into the role of normal aging on the molecular mechanisms of rTMS, there are several points to consider when applying it in a translational context. The first is that our study characterised the changes to gene expression after a single session of iTBS or cTBS, whereas clinical rTMS is often applied over several days to weeks to achieve its therapeutic effect. It has been suggested that increasing the number of stimulation sessions delivered enhances the treatment efficacy of rTMS in older adults, producing comparable outcomes to that seen in younger adult cohorts (Cotovio, Boes, Press, Oliveira-Maia, & Pascual-Leone, 2022). However, how repeated sessions of rTMS would affect gene expression in the aged brain is unclear as a previous study in adult rats has shown that multiple sessions of rTMS leads to a complex and non-linear change to protein levels (Volz, Benali, Mix, Neubacher, & Funke, 2013), and repeated sessions of iTBS to the aged sensorimotor cortex in mice does not result in greater structural synaptic plasticity compared to a single session (Tang et al., 2021). Nevertheless, a spatial transcriptomic characterisation of the aged brain after multiple sessions of rTMS would complement our results to further the development of optimised protocols for clinical use. Similarly, future studies employing additional approaches spanning proteomic, electrophysiological and behavioural approaches across species are needed to determine the consequence of rTMS-induced transcriptomic changes. For example, studies beginning to employ single cell electrophysiology in human cortical tissue of different ages stimulated with rTMS (Vlachos, 2025), combined with spatial transcriptomics (Ong & Tang, 2025) will help uncover the relationship between the functional and transcriptomic changes induced by rTMS, as well as the impact of aging on rTMS plasticity in human neurons.

In conclusion, our study has shown that rTMS to the cortex of aged mice results in neural plasticity across cortical and subcortical brain regions. Similar to young adult mice, the neural plasticity and biological process affected by rTMS in the aged brain were largely dependent on the brain region, cortical layer and protocol of rTMS used. However, for the current TBS protocols of rTMS, the aged brain generally induced less neuromodulation and diversity in neural plasticity. These findings highlight the need to account for age when applying and interpreting the effect of rTMS in older adults. Furthermore, we show the critical need to develop rTMS protocols specifically optimised for the aged brain.

## Acknowledgments

The authors thank N. Matigian for bioinformatics services (Queensland Cyber Infrastructure Foundation).

## Author contributions

Conceptualization: ADT, RCSO

Methodology: ADT, RCSO

Investigation: ADT, RCSO

Visualization: RCSO

Funding acquisition: ADT, RCSO

Project administration: ADT

Supervision: ADT

Writing – original draft: ADT, RCSO

Writing – review & editing: ADT, RCSO

## Data and materials availability

The aged bulk RNA-seq and spatial transcriptomics data have been deposited in the Gene Omnibus Expression database (GSE274580 and GSE274782 respectively). The young adult spatial transcriptomic dataset used for comparison was downloaded from the Gene Omnibus Expression database (GSE259405).

---

**Fig. S1:**
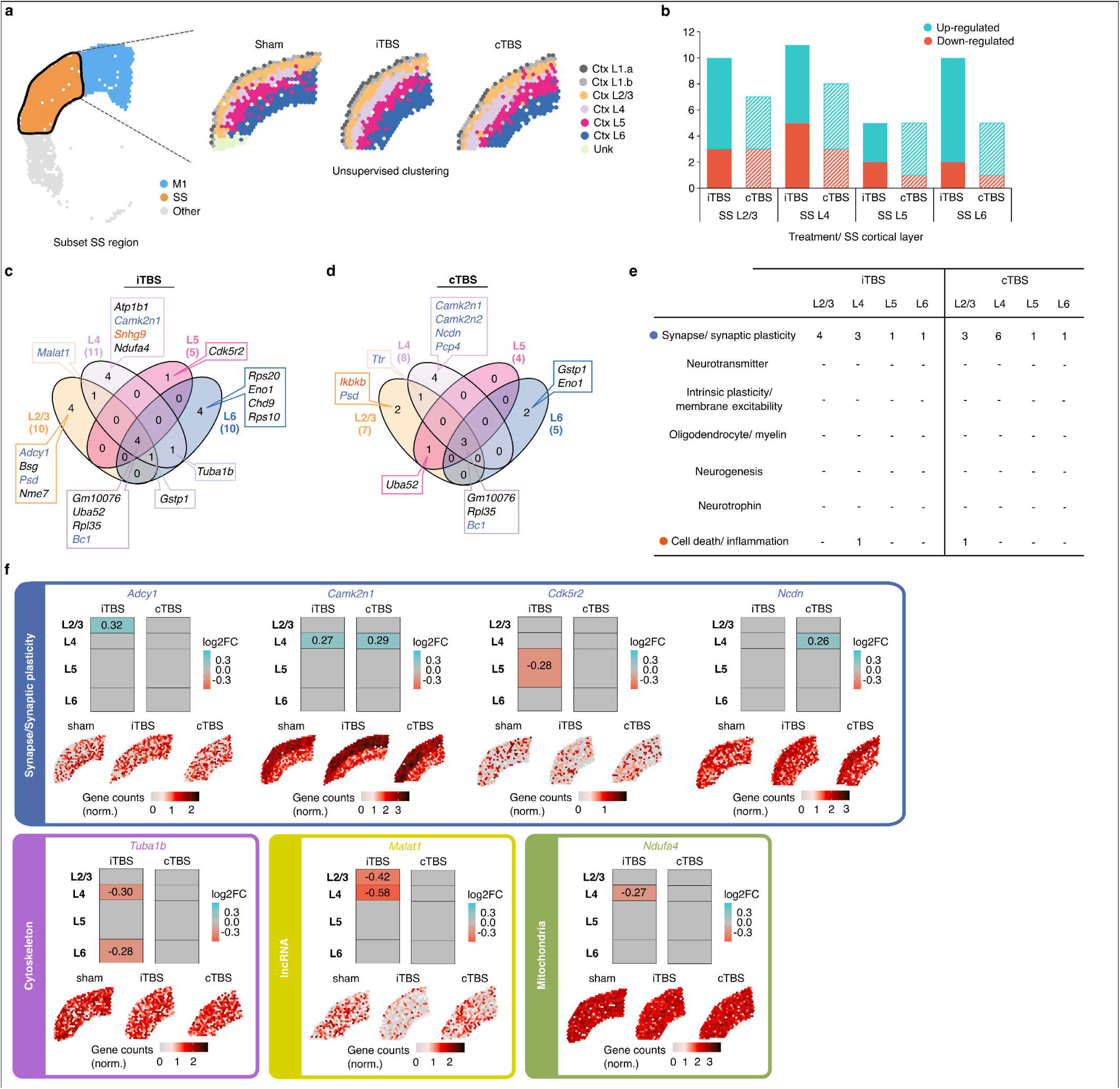
Bulk RNA-sequencing of the whole sensorimotor cortex of aged mice indicated little-to-no significant effect of stimulation on gene expression. **a**, Two-dimensional PCA plot of iTBS, cTBS and sham stimulation treated samples. **b**, Volcano plot of genes differentially expressed following cTBS treatment indicates no significant (p.adjusted < 0.05, absolute log2 fold-change >1) effect following stimulation on the aged sensorimotor cortex. **c**, Volcano plot of genes differentially expressed following iTBS treatment identified a single gene that was significantly downregulated following stimulation.

**Fig. S2:**
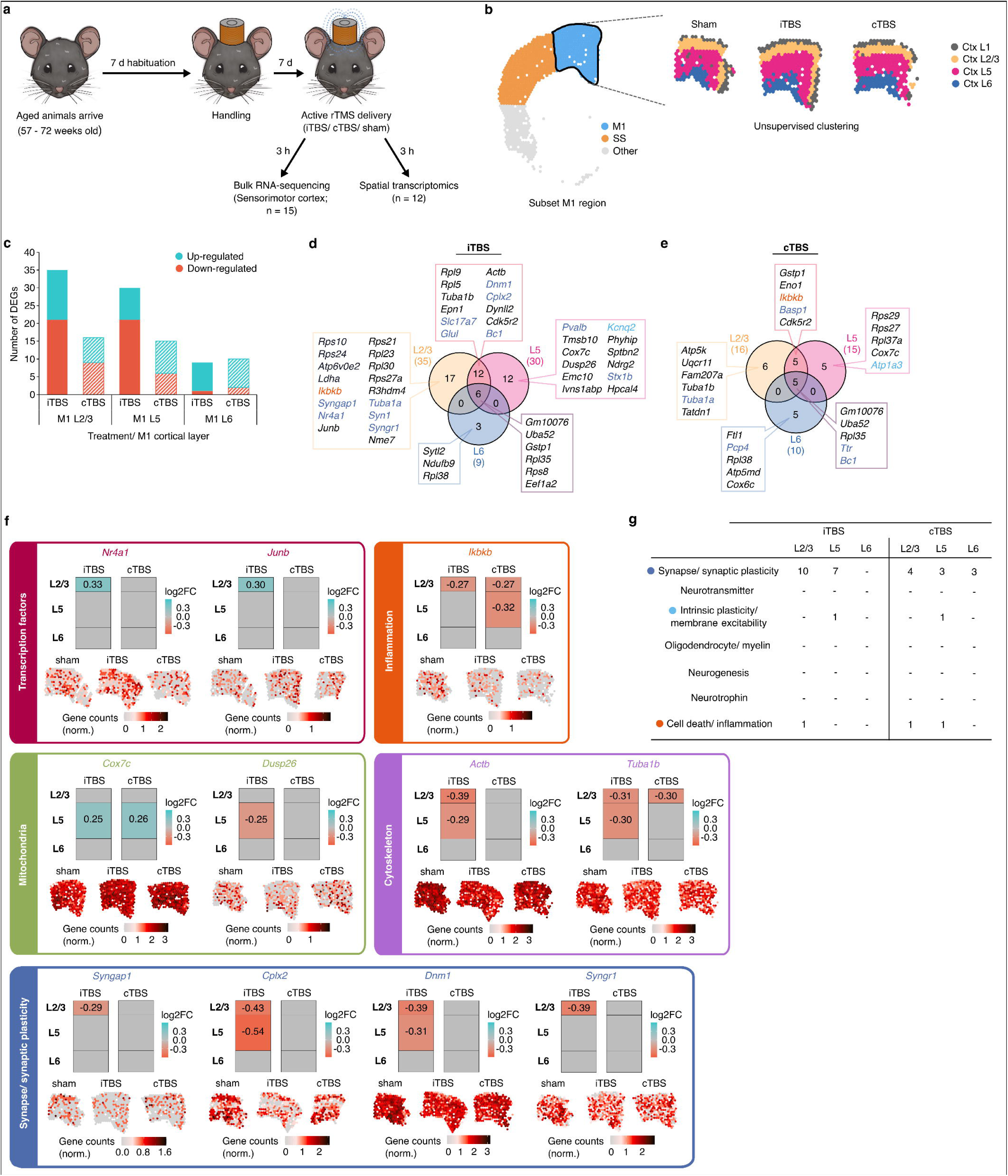
Clustering and quality checks of spatial transcriptomics samples. **a**, Initial unsupervised clustering analysis identified 13 clusters. Of these, clusters that overlaid the cortex and striatum were subsetted and an additional unsupervised clustering analysis was performed to further delineate additional cortical layers and sub-cortical structures. **b**, Final spatial transcriptome maps representative of each treatment group, coloured based on the different clusters identified. Clusters were also manually annotated based on their anatomical location based on the Allen Mouse Brain Atlas. A uniform manifold approximation and projection (UMAP) plot displays all the identified clusters. **c**, Bar graph displaying the number of spots within each cluster across all samples.

