## Supplementary material for "Normal aging impacts the extent and diversity of neural plasticity induced in the mouse brain with repetitive transcranial magnetic stimulation"

**Data S1.** Summary of spatial transcriptomics samples.

| | | | | | | | | | | | | |
|---|---|---|---|---|---|---|---|---|---|---|---|---|
| Sample ID | ET22A01 | ET22A02 | ET22A03 | ET22A04 | ET22A05 | ET22A06 | ET22A07 | ET22A09 | ET22A11 | ET22A12 | ET22A13 | ET22A14 |
| Stimulation condition | iTBS | cTBS | sham | sham | cTBS | iTBS | sham | cTBS | iTBS | sham | iTBS | cTBS |
| Visium slide number | 1 | 1 | 4 | 4 | 3 | 2 | 2 | 4 | 4 | 3 | 3 | 3 |
| Visium area number | A1 | B1 | A1 | C1 | A1 | A1 | B1 | B1 | D1 | B1 | C1 | D1 |
| Number of Reads | 213720183 | 203973068 | 218242715 | 211621221 | 231707408 | 200887856 | 231879150 | 208106723 | 203712276 | 219756357 | 197475094 | 205859379 |

**Data S2.** Differentially expressed genes (DEGs) identified in sub-cortical regions following iTBS, and cTBS. DEGs were identified using a cutoff of p.adj ≤ 0.05.

| Brain region | Stimulation | Gene Symbol | Gene Name | EntrezID | Avg log2FoldChange | % expressed in stimulated samples | % expressed in sham samples | p.adjust value (Bonferroni correction) | p.value |
|---|---|---|---|---|---|---|---|---|---|
| White Matter Tracts | iTBS | Bc1 | brain cytoplasmic RNA 1 | 100568459 | -0.433049208 | 0.816 | 0.885 | 0.017471008 | 1.18E-06 |
| White Matter Tracts | iTBS | Snx3 | sorting nexin 3 | 54198 | -0.256664141 | 0.791 | 0.877 | 0.002128645 | 1.44E-07 |
| White Matter Tracts | iTBS | Dync1h1 | dynein cytoplasmic 1 heavy chain 1 | 13424 | -0.235728648 | 0.827 | 0.913 | 0.002055333 | 1.39E-07 |
| White Matter Tracts | iTBS | Ikbkb | inhibitor of kappaB kinase beta | 16150 | -0.232327107 | 0.049 | 0.127 | 0.031666974 | 2.14E-06 |
| White Matter Tracts | iTBS | Atp6v0d1 | ATPase, H+ transporting, lysosomal V0 subunit D1 | 11972 | -0.221361557 | 0.773 | 0.868 | 0.020105921 | 1.36E-06 |
| White Matter Tracts | iTBS | Gprasp1 | G protein-coupled receptor associated sorting protein 1 | 67298 | -0.218646244 | 0.731 | 0.865 | 0.000181459 | 1.23E-08 |
| White Matter Tracts | iTBS | Cyfip2 | cytoplasmic FMR1 interacting protein 2 | 76884 | -0.206299765 | 0.747 | 0.862 | 0.000484227 | 3.28E-08 |
| White Matter Tracts | iTBS | Atp6v0a1 | ATPase, H+ transporting, lysosomal V0 subunit A1 | 11975 | -0.200278917 | 0.717 | 0.813 | 0.011909707 | 8.06E-07 |
| White Matter Tracts | iTBS | Itpr1 | inositol 1,4,5-trisphosphate receptor 1 | 16438 | -0.178168816 | 0.686 | 0.822 | 0.00638697 | 4.32E-07 |
| White Matter Tracts | iTBS | Syn2 | synapsin II | 20965 | -0.177677157 | 0.724 | 0.843 | 0.01599833 | 1.08E-06 |
| White Matter Tracts | iTBS | Sptbn2 | spectrin beta, non-erythrocytic 2 | 20743 | -0.174942322 | 0.686 | 0.818 | 0.003636036 | 2.46E-07 |
| White Matter Tracts | iTBS | Sv2a | synaptic vesicle glycoprotein 2 a | 64051 | -0.172060746 | 0.749 | 0.833 | 0.026762547 | 1.81E-06 |
| White Matter Tracts | iTBS | Kifap3 | kinesin-associated protein 3 | 16579 | -0.171420558 | 0.718 | 0.847 | 0.002705414 | 1.83E-07 |
| White Matter Tracts | iTBS | Fabp3 | fatty acid binding protein 3, muscle and heart | 14077 | -0.171345547 | 0.659 | 0.768 | 0.019439221 | 1.32E-06 |
| White Matter Tracts | iTBS | Map2 | microtubule-associated protein 2 | 17756 | -0.167642608 | 0.755 | 0.872 | 0.000599496 | 4.06E-08 |
| White Matter Tracts | iTBS | Epn1 | epsin 1 | 13854 | -0.167487046 | 0.771 | 0.88 | 0.027127363 | 1.84E-06 |
| White Matter Tracts | iTBS | Map1a | microtubule-associated protein 1 A | 17754 | -0.165190308 | 0.727 | 0.843 | 0.012618462 | 8.54E-07 |
| White Matter Tracts | iTBS | Atp6v1d | ATPase, H+ transporting, lysosomal V1 subunit D | 73834 | -0.164873352 | 0.76 | 0.858 | 0.03834351 | 2.59E-06 |
| White Matter Tracts | iTBS | Pdxp | pyridoxal (pyridoxine, vitamin B6) phosphatase | 57028 | -0.164469269 | 0.711 | 0.828 | 0.041748572 | 2.83E-06 |
| White Matter Tracts | iTBS | Nat8l | N-acetyltransferase 8-like | 269642 | -0.158046699 | 0.623 | 0.742 | 0.013943774 | 9.44E-07 |
| White Matter Tracts | iTBS | Enc1 | ectodermal-neural cortex 1 | 13803 | -0.156182649 | 0.652 | 0.765 | 0.043706695 | 2.96E-06 |
| White Matter Tracts | iTBS | Gnao1 | guanine nucleotide binding protein, alpha O | 14681 | -0.154016163 | 0.821 | 0.913 | 0.025468443 | 1.72E-06 |
| White Matter Tracts | iTBS | Atxn10 | ataxin 10 | 54138 | -0.142676998 | 0.764 | 0.88 | 0.045877523 | 3.10E-06 |
| White Matter Tracts | iTBS | Pde2a | phosphodiesterase 2A, cGMP-stimulated | 207728 | -0.135384779 | 0.662 | 0.79 | 0.011522815 | 7.80E-07 |
| White Matter Tracts | iTBS | Dync1i1 | dynein cytoplasmic 1 intermediate chain 1 | 13426 | 0.043993426 | 0.231 | 0.088 | 2.18E-05 | 1.48E-09 |
| White Matter Tracts | iTBS | Praf2 | PRA1 domain family 2 | 54637 | 0.068533939 | 0.236 | 0.113 | 0.012763434 | 8.64E-07 |
| White Matter Tracts | iTBS | Prmt1 | protein arginine N-methyltransferase 1 | 15469 | 0.071608496 | 0.439 | 0.275 | 0.025418944 | 1.72E-06 |
| White Matter Tracts | iTBS | C1d | C1D nuclear receptor co-repressor | 57316 | 0.071951824 | 0.291 | 0.128 | 3.46E-05 | 2.34E-09 |
| White Matter Tracts | iTBS | Cacybp | calcyclin binding protein | 12301 | 0.073311383 | 0.273 | 0.133 | 0.003565862 | 2.41E-07 |
| White Matter Tracts | iTBS | Actr3 | ARP3 actin-related protein 3 | 74117 | 0.074342733 | 0.498 | 0.338 | 0.037685894 | 2.55E-06 |
| White Matter Tracts | iTBS | Hspd1 | heat shock protein 1 (chaperonin) | 15510 | 0.074386776 | 0.417 | 0.255 | 0.020258934 | 1.37E-06 |
| White Matter Tracts | iTBS | Mmd2 | monocyte to macrophage differentiation-associated 2 | 75104 | 0.074645476 | 0.547 | 0.35 | 0.01576143 | 1.07E-06 |
| White Matter Tracts | iTBS | Aars | alanyl-tRNA synthetase | 234734 | 0.075200505 | 0.332 | 0.173 | 0.003284767 | 2.22E-07 |
| White Matter Tracts | iTBS | Ehmt2 | euchromatic histone lysine N-methyltransferase 2 | 110147 | 0.083212026 | 0.397 | 0.243 | 0.013814079 | 9.35E-07 |
| White Matter Tracts | iTBS | Atxn7l3 | ataxin 7-like 3 | 217218 | 0.083408609 | 0.363 | 0.205 | 0.002629173 | 1.78E-07 |
| White Matter Tracts | iTBS | Cct8 | chaperonin containing Tcp1, subunit 8 (theta) | 12469 | 0.084224245 | 0.487 | 0.31 | 0.001561987 | 1.06E-07 |
| White Matter Tracts | iTBS | Rnf126 | ring finger protein 126 | 70294 | 0.091064402 | 0.236 | 0.1 | 0.00016056 | 1.09E-08 |
| White Matter Tracts | iTBS | Ik | IK cytokine | 24010 | 0.091410334 | 0.321 | 0.152 | 5.97E-05 | 4.04E-09 |
| White Matter Tracts | iTBS | Commd4 | COMM domain containing 4 | 66199 | 0.092188233 | 0.531 | 0.357 | 0.017663457 | 1.20E-06 |
| White Matter Tracts | iTBS | Brinp1 | bone morphogenic protein/retinoic acid inducible neural specific 1 | 56710 | 0.093189454 | 0.392 | 0.243 | 0.014316736 | 9.69E-07 |
| White Matter Tracts | iTBS | Pdxk | pyridoxal (pyridoxine, vitamin B6) kinase | 216134 | 0.093963537 | 0.41 | 0.257 | 0.036084628 | 2.44E-06 |
| White Matter Tracts | iTBS | Gng2 | guanine nucleotide binding protein (G protein), gamma 2 | 14702 | 0.09478512 | 0.211 | 0.093 | 0.002334959 | 1.58E-07 |
| White Matter Tracts | iTBS | Cct2 | chaperonin containing Tcp1, subunit 2 (beta) | 12461 | 0.095425437 | 0.455 | 0.288 | 0.01317107 | 8.91E-07 |
| White Matter Tracts | iTBS | Mgll | monoglyceride lipase | 23945 | 0.09561621 | 0.341 | 0.187 | 0.003867957 | 2.62E-07 |
| White Matter Tracts | iTBS | Use1 | unconventional SNARE in the ER 1 homolog (S. cerevisiae) | 67023 | 0.097433564 | 0.448 | 0.26 | 0.001912664 | 1.29E-07 |
| White Matter Tracts | iTBS | Chd3 | chromodomain helicase DNA binding protein 3 | 216848 | 0.098713078 | 0.357 | 0.183 | 7.58E-05 | 5.13E-09 |
| White Matter Tracts | iTBS | Pip5k1c | phosphatidylinositol-4-phosphate 5-kinase, type 1 gamma | 18717 | 0.10321869 | 0.278 | 0.127 | 5.35E-05 | 3.62E-09 |
| White Matter Tracts | iTBS | Mrpl11 | mitochondrial ribosomal protein L11 | 66419 | 0.104494362 | 0.412 | 0.225 | 5.54E-05 | 3.75E-09 |
| White Matter Tracts | iTBS | Mrpl28 | mitochondrial ribosomal protein L28 | 68611 | 0.104999452 | 0.439 | 0.257 | 0.000673266 | 4.56E-08 |
| White Matter Tracts | iTBS | Calb1 | calbindin 1 | 12307 | 0.105974317 | 0.394 | 0.252 | 0.044200825 | 2.99E-06 |
| White Matter Tracts | iTBS | Nol7 | nucleolar protein 7 | 70078 | 0.106189258 | 0.551 | 0.36 | 0.003437455 | 2.33E-07 |
| White Matter Tracts | iTBS | Smpd1 | sphingomyelin phosphodiesterase 1, acid lysosomal | 20597 | 0.106524662 | 0.404 | 0.237 | 0.00226689 | 1.53E-07 |
| White Matter Tracts | iTBS | Hnrnpa0 | heterogeneous nuclear ribonucleoprotein A0 | 77134 | 0.106585064 | 0.397 | 0.232 | 0.005579856 | 3.78E-07 |
| White Matter Tracts | iTBS | Gm17018 | NA | NA | 0.106742041 | 0.431 | 0.263 | 0.010717744 | 7.25E-07 |
| White Matter Tracts | iTBS | Dgcr6 | DiGeorge syndrome critical region gene 6 | 13353 | 0.10762072 | 0.26 | 0.137 | 0.049144235 | 3.33E-06 |
| White Matter Tracts | iTBS | Cs | citrate synthase | 12974 | 0.109865098 | 0.379 | 0.203 | 0.000336764 | 2.28E-08 |
| White Matter Tracts | iTBS | Celsr2 | cadherin, EGF LAG seven-pass G-type receptor 2 | 53883 | 0.111046647 | 0.227 | 0.092 | 3.90E-05 | 2.64E-09 |
| White Matter Tracts | iTBS | Nudt19 | nudix (nucleoside diphosphate linked moiety X)-type motif 19 | 110959 | 0.111249745 | 0.282 | 0.127 | 3.01E-05 | 2.04E-09 |
| White Matter Tracts | iTBS | Smap2 | small ArfGAP 2 | 69780 | 0.111591752 | 0.403 | 0.245 | 0.0109304 | 7.40E-07 |
| White Matter Tracts | iTBS | Iqsec1 | IQ motif and Sec7 domain 1 | 232227 | 0.111709673 | 0.422 | 0.257 | 0.009541888 | 6.46E-07 |
| White Matter Tracts | iTBS | Grb2 | growth factor receptor bound protein 2 | 14784 | 0.111977286 | 0.518 | 0.352 | 0.030964313 | 2.10E-06 |
| White Matter Tracts | iTBS | Ube2v2 | ubiquitin-conjugating enzyme E2 variant 2 | 70620 | 0.113223351 | 0.31 | 0.157 | 0.000789442 | 5.34E-08 |
| White Matter Tracts | iTBS | Dkk3 | dickkopf WNT signaling pathway inhibitor 3 | 50781 | 0.115076524 | 0.412 | 0.242 | 0.000396191 | 2.68E-08 |
| White Matter Tracts | iTBS | Rufy3 | RUN and FYVE domain containing 3 | 52822 | 0.115076524 | 0.504 | 0.333 | 0.010396851 | 7.04E-07 |
| White Matter Tracts | iTBS | Kcnk1 | potassium channel, subfamily K, member 1 | 16525 | 0.115076524 | 0.403 | 0.24 | 0.001541284 | 1.04E-07 |
| White Matter Tracts | iTBS | Acly | ATP citrate lyase | 104112 | 0.116683984 | 0.493 | 0.305 | 0.000394228 | 2.67E-08 |
| White Matter Tracts | iTBS | Epdr1 | ependymin related protein 1 (zebrafish) | 105298 | 0.116726262 | 0.468 | 0.282 | 0.000311743 | 2.11E-08 |
| White Matter Tracts | iTBS | Bod1 | biorientation of chromosomes in cell division 1 | 69556 | 0.116735748 | 0.469 | 0.303 | 0.009964813 | 6.74E-07 |
| White Matter Tracts | iTBS | Prkce | protein kinase C, epsilon | 18754 | 0.116817858 | 0.437 | 0.273 | 0.003240253 | 2.19E-07 |
| White Matter Tracts | iTBS | Dmwd | dystrophia myotonica-containing WD repeat motif | 13401 | 0.118340542 | 0.43 | 0.248 | 0.000629985 | 4.26E-08 |
| White Matter Tracts | iTBS | Tspan5 | tetraspanin 5 | 56224 | 0.118475101 | 0.431 | 0.25 | 0.00028039 | 1.90E-08 |
| White Matter Tracts | iTBS | Rnf10 | ring finger protein 10 | 50849 | 0.118532083 | 0.394 | 0.22 | 0.000285255 | 1.93E-08 |
| White Matter Tracts | iTBS | Klhdc3 | kelch domain containing 3 | 71765 | 0.118578213 | 0.419 | 0.25 | 0.000771326 | 5.22E-08 |
| White Matter Tracts | iTBS | Gprasp2 | G protein-coupled receptor associated sorting protein 2 | 245607 | 0.118878095 | 0.323 | 0.147 | 1.75E-06 | 1.19E-10 |
| White Matter Tracts | iTBS | Hmgb1 | high mobility group box 1 | 15289 | 0.119146189 | 0.592 | 0.393 | 0.003647968 | 2.47E-07 |
| White Matter Tracts | iTBS | Sar1a | secretion associated Ras related GTPase 1A | 20224 | 0.120201557 | 0.399 | 0.222 | 0.000404872 | 2.74E-08 |
| White Matter Tracts | iTBS | Klhl2 | kelch-like 2, Mayven | 77113 | 0.120935193 | 0.551 | 0.35 | 0.002103163 | 1.42E-07 |
| White Matter Tracts | iTBS | Fam171b | family with sequence similarity 171, member B | 241520 | 0.121404144 | 0.466 | 0.285 | 0.001457007 | 9.86E-08 |
| White Matter Tracts | iTBS | Clpp | caseinolytic mitochondrial matrix peptidase proteolytic subunit | 53895 | 0.123269023 | 0.444 | 0.265 | 0.002141941 | 1.45E-07 |
| White Matter Tracts | iTBS | H13 | histocompatibility 13 | 14950 | 0.123689655 | 0.135 | 0.053 | 0.048384163 | 3.27E-06 |
| White Matter Tracts | iTBS | Reps2 | RALBP1 associated Eps domain containing protein 2 | 194590 | 0.124110336 | 0.394 | 0.228 | 0.00073276 | 4.96E-08 |
| White Matter Tracts | iTBS | Mbnl2 | muscleblind like splicing factor 2 | 105559 | 0.124671681 | 0.619 | 0.443 | 0.027265637 | 1.85E-06 |
| White Matter Tracts | iTBS | Arnt2 | aryl hydrocarbon receptor nuclear translocator 2 | 11864 | 0.125320556 | 0.363 | 0.177 | 1.17E-05 | 7.94E-10 |
| White Matter Tracts | iTBS | Rprml | reprimo-like | 104582 | 0.125829581 | 0.406 | 0.25 | 0.004303787 | 2.91E-07 |
| White Matter Tracts | iTBS | Hmox2 | heme oxygenase 2 | 15369 | 0.126366538 | 0.361 | 0.192 | 0.001109806 | 7.51E-08 |
| White Matter Tracts | iTBS | Ahsa1 | AHA1, activator of heat shock protein ATPase 1 | 217737 | 0.126421498 | 0.3 | 0.14 | 8.79E-05 | 5.95E-09 |
| White Matter Tracts | iTBS | Gsk3b | glycogen synthase kinase 3 beta | 56637 | 0.127645198 | 0.388 | 0.217 | 0.000151507 | 1.03E-08 |
| White Matter Tracts | iTBS | Asphd2 | aspartate beta-hydroxylase domain containing 2 | 72898 | 0.128932046 | 0.455 | 0.29 | 0.001743015 | 1.18E-07 |
| White Matter Tracts | iTBS | Rbfox3 | RNA binding protein, fox-1 homolog (C. elegans) 3 | 52897 | 0.129073676 | 0.276 | 0.132 | 0.000128826 | 8.72E-09 |
| White Matter Tracts | iTBS | Denr | density-regulated protein | 68184 | 0.12915171 | 0.381 | 0.193 | 1.98E-05 | 1.34E-09 |
| White Matter Tracts | iTBS | Atp6v1h | ATPase, H+ transporting, lysosomal V1 subunit H | 108664 | 0.129431817 | 0.209 | 0.087 | 0.000369757 | 2.50E-08 |
| White Matter Tracts | iTBS | Slc39a10 | solute carrier family 39 (zinc transporter), member 10 | 227059 | 0.12961785 | 0.231 | 0.073 | 1.85E-08 | 1.25E-12 |
| White Matter Tracts | iTBS | Rab18 | RAB18, member RAS oncogene family | 19330 | 0.129704736 | 0.291 | 0.147 | 0.00348324 | 2.36E-07 |
| White Matter Tracts | iTBS | Evl | Ena-vasodilator stimulated phosphoprotein | 14026 | 0.130104734 | 0.319 | 0.168 | 0.000501413 | 3.39E-08 |
| White Matter Tracts | iTBS | Pianp | PILR alpha associated neural protein | 319352 | 0.131635616 | 0.227 | 0.075 | 7.43E-08 | 5.03E-12 |
| White Matter Tracts | iTBS | Ahi1 | Abelson helper integration site 1 | 52906 | 0.131793919 | 0.455 | 0.298 | 0.015147866 | 1.03E-06 |
| White Matter Tracts | iTBS | Ndufa9 | NADH:ubiquinone oxidoreductase subunit A9 | 66108 | 0.132016387 | 0.303 | 0.127 | 1.12E-06 | 7.56E-11 |
| White Matter Tracts | iTBS | Sesn3 | sestrin 3 | 75747 | 0.132105254 | 0.327 | 0.153 | 9.80E-06 | 6.63E-10 |
| White Matter Tracts | iTBS | Fbxo44 | F-box protein 44 | 230903 | 0.132909818 | 0.394 | 0.2 | 3.17E-06 | 2.14E-10 |
| White Matter Tracts | iTBS | Gng10 | guanine nucleotide binding protein (G protein), gamma 10 | 14700 | 0.132998432 | 0.282 | 0.142 | 0.001857206 | 1.26E-07 |
| White Matter Tracts | iTBS | Mapre2 | microtubule-associated protein, RP/EB family, member 2 | 212307 | 0.133020724 | 0.285 | 0.15 | 0.018430322 | 1.25E-06 |
| White Matter Tracts | iTBS | Dnm3 | dynamin 3 | 103967 | 0.133223871 | 0.377 | 0.19 | 3.71E-06 | 2.51E-10 |
| White Matter Tracts | iTBS | Gda | guanine deaminase | 14544 | 0.133533429 | 0.251 | 0.08 | 1.60E-09 | 1.08E-13 |
| White Matter Tracts | iTBS | Rgs10 | regulator of G-protein signalling 10 | 67865 | 0.133540587 | 0.401 | 0.215 | 3.09E-05 | 2.09E-09 |
| White Matter Tracts | iTBS | Ybx1 | Y box protein 1 | 22608 | 0.134024011 | 0.619 | 0.413 | 0.000465951 | 3.15E-08 |
| White Matter Tracts | iTBS | Hectd4 | HECT domain E3 ubiquitin protein ligase 4 | 269700 | 0.134584683 | 0.431 | 0.253 | 7.95E-05 | 5.38E-09 |
| White Matter Tracts | iTBS | Eif3i | eukaryotic translation initiation factor 3, subunit I | 54709 | 0.134772326 | 0.48 | 0.275 | 2.24E-05 | 1.52E-09 |
| White Matter Tracts | iTBS | Herc1 | HECT and RLD domain containing E3 ubiquitin protein ligase family meml | 235439 | 0.134778439 | 0.361 | 0.202 | 0.000996738 | 6.74E-08 |
| White Matter Tracts | iTBS | Strbp | spermatid perinuclear RNA binding protein | 20744 | 0.135228783 | 0.338 | 0.177 | 0.000298761 | 2.02E-08 |
| White Matter Tracts | iTBS | Tspyl4 | TSPY-like 4 | 72480 | 0.135305849 | 0.339 | 0.17 | 4.72E-05 | 3.19E-09 |
| White Matter Tracts | iTBS | Ssr4 | signal sequence receptor, delta | 20832 | 0.135585383 | 0.496 | 0.3 | 0.000272262 | 1.84E-08 |
| White Matter Tracts | iTBS | Astn1 | astrotactin 1 | 11899 | 0.136112548 | 0.435 | 0.277 | 0.006271907 | 4.24E-07 |
| White Matter Tracts | iTBS | Disp2 | dispatched RND tramsporter family member 2 | 214240 | 0.136507905 | 0.3 | 0.137 | 6.01E-06 | 4.07E-10 |
| White Matter Tracts | iTBS | Ppp2r2a | protein phosphatase 2, regulatory subunit B, alpha | 71978 | 0.137054894 | 0.525 | 0.363 | 0.008980293 | 6.08E-07 |
| White Matter Tracts | iTBS | mt-Cytb | NA | NA | 0.137413366 | 1 | 1 | 0.013798673 | 9.34E-07 |
| White Matter Tracts | iTBS | Pet100 | PET100 homolog | 100503890 | 0.137630841 | 0.543 | 0.358 | 0.005545063 | 3.75E-07 |
| White Matter Tracts | iTBS | Adprh | ADP-ribosylarginine hydrolase | 11544 | 0.137907233 | 0.381 | 0.215 | 0.001627521 | 1.10E-07 |
| White Matter Tracts | iTBS | Setd3 | SET domain containing 3 | 52690 | 0.1382296 | 0.574 | 0.397 | 0.007795129 | 5.27E-07 |
| White Matter Tracts | iTBS | Lpgat1 | lysophosphatidylglycerol acyltransferase 1 | 226856 | 0.138535497 | 0.339 | 0.148 | 3.02E-07 | 2.05E-11 |
| White Matter Tracts | iTBS | Cope | coatomer protein complex, subunit epsilon | 59042 | 0.138535497 | 0.529 | 0.32 | 3.26E-05 | 2.21E-09 |
| White Matter Tracts | iTBS | Polr2l | polymerase (RNA) II (DNA directed) polypeptide L | 66491 | 0.139242836 | 0.513 | 0.35 | 0.005944743 | 4.02E-07 |
| White Matter Tracts | iTBS | Myh10 | myosin, heavy polypeptide 10, non-muscle | 77579 | 0.13935808 | 0.273 | 0.122 | 1.65E-05 | 1.12E-09 |
| White Matter Tracts | iTBS | Dst | dystonin | 13518 | 0.140479372 | 0.471 | 0.253 | 8.78E-06 | 5.94E-10 |
| White Matter Tracts | iTBS | Cuta | cutA divalent cation tolerance homolog | 67675 | 0.141855856 | 0.404 | 0.225 | 0.000576658 | 3.90E-08 |
| White Matter Tracts | iTBS | Phyhip | phytanoyl-CoA hydroxylase interacting protein | 105653 | 0.142043572 | 0.401 | 0.237 | 0.000504855 | 3.42E-08 |
| White Matter Tracts | iTBS | Nrcam | neuronal cell adhesion molecule | 319504 | 0.142963482 | 0.484 | 0.328 | 0.01958101 | 1.33E-06 |
| White Matter Tracts | iTBS | Eif5 | eukaryotic translation initiation factor 5 | 217869 | 0.143381894 | 0.439 | 0.237 | 2.96E-06 | 2.00E-10 |
| White Matter Tracts | iTBS | Ola1 | Obg-like ATPase 1 | 67059 | 0.144747391 | 0.372 | 0.182 | 2.93E-07 | 1.98E-11 |
| White Matter Tracts | iTBS | Tafa5 | TAFA chemokine like family member 5 | 106014 | 0.144862254 | 0.319 | 0.132 | 6.07E-08 | 4.11E-12 |
| White Matter Tracts | iTBS | Ablim2 | actin-binding LIM protein 2 | 231148 | 0.144988343 | 0.226 | 0.095 | 0.000154746 | 1.05E-08 |
| White Matter Tracts | iTBS | Rpl10 | ribosomal protein L10 | 110954 | 0.14536322 | 0.973 | 0.982 | 0.03599266 | 2.44E-06 |
| White Matter Tracts | iTBS | Hagh | hydroxyacyl glutathione hydrolase | 14651 | 0.148473734 | 0.449 | 0.262 | 8.43E-05 | 5.70E-09 |
| White Matter Tracts | iTBS | Trappc6b | trafficking protein particle complex 6B | 78232 | 0.148803399 | 0.332 | 0.167 | 5.32E-05 | 3.60E-09 |
| White Matter Tracts | iTBS | 1700037H04Rik | RIKEN cDNA 1700037H04 gene | 67326 | 0.149023856 | 0.253 | 0.092 | 9.18E-08 | 6.21E-12 |
| White Matter Tracts | iTBS | Lynx1 | Ly6/neurotoxin 1 | 23936 | 0.149202308 | 0.354 | 0.188 | 4.84E-05 | 3.28E-09 |
| White Matter Tracts | iTBS | Fam131a | family with sequence similarity 131, member A | 78408 | 0.149535995 | 0.421 | 0.248 | 0.000115909 | 7.84E-09 |
| White Matter Tracts | iTBS | Gars | glycyl-tRNA synthetase | 353172 | 0.149891295 | 0.231 | 0.083 | 8.64E-07 | 5.85E-11 |
| White Matter Tracts | iTBS | Hnrnpul2 | heterogeneous nuclear ribonucleoprotein U-like 2 | 68693 | 0.150288556 | 0.318 | 0.16 | 0.000264259 | 1.79E-08 |
| White Matter Tracts | iTBS | Rit2 | Ras-like without CAAX 2 | 19762 | 0.150946481 | 0.236 | 0.088 | 3.43E-06 | 2.32E-10 |
| White Matter Tracts | iTBS | Cct4 | chaperonin containing Tcp1, subunit 4 (delta) | 12464 | 0.151383528 | 0.394 | 0.2 | 2.47E-06 | 1.67E-10 |
| White Matter Tracts | iTBS | Chmp5 | charged multivesicular body protein 5 | 76959 | 0.1516024 | 0.401 | 0.22 | 0.000836798 | 5.66E-08 |
| White Matter Tracts | iTBS | Cdkn2d | cyclin dependent kinase inhibitor 2D | 12581 | 0.151835074 | 0.339 | 0.165 | 9.07E-06 | 6.14E-10 |
| White Matter Tracts | iTBS | Arrb1 | arrestin, beta 1 | 109689 | 0.152275142 | 0.388 | 0.228 | 0.001133648 | 7.67E-08 |
| White Matter Tracts | iTBS | Trim37 | tripartite motif-containing 37 | 68729 | 0.154506669 | 0.433 | 0.242 | 1.50E-06 | 1.01E-10 |
| White Matter Tracts | iTBS | Mycbp2 | MYC binding protein 2, E3 ubiquitin protein ligase | 105689 | 0.155614684 | 0.347 | 0.162 | 8.60E-07 | 5.82E-11 |
| White Matter Tracts | iTBS | Spryd3 | SPRY domain containing 3 | 223918 | 0.155992968 | 0.282 | 0.118 | 1.40E-06 | 9.47E-11 |
| White Matter Tracts | iTBS | Ptpra | protein tyrosine phosphatase, receptor type, A | 19262 | 0.156089907 | 0.551 | 0.362 | 0.002039282 | 1.38E-07 |
| White Matter Tracts | iTBS | Ube2e1 | ubiquitin-conjugating enzyme E2E 1 | 22194 | 0.156751723 | 0.478 | 0.317 | 0.004999805 | 3.38E-07 |
| White Matter Tracts | iTBS | Polr2g | polymerase (RNA) II (DNA directed) polypeptide G | 67710 | 0.156949477 | 0.33 | 0.162 | 8.73E-05 | 5.91E-09 |
| White Matter Tracts | iTBS | Smarcd3 | SWI/SNF related, matrix associated, actin dependent regulator of chroma | 66993 | 0.157245979 | 0.273 | 0.11 | 3.39E-07 | 2.29E-11 |
| White Matter Tracts | iTBS | Madd | MAP-kinase activating death domain | 228355 | 0.157773452 | 0.426 | 0.248 | 2.53E-05 | 1.71E-09 |
| White Matter Tracts | iTBS | Raly | hnRNP-associated with lethal yellow | 19383 | 0.1577882 | 0.352 | 0.183 | 0.000616583 | 4.17E-08 |
| White Matter Tracts | iTBS | Ppib | peptidylprolyl isomerase B | 19035 | 0.158244384 | 0.366 | 0.182 | 1.88E-05 | 1.28E-09 |
| White Matter Tracts | iTBS | Aatk | apoptosis-associated tyrosine kinase | 11302 | 0.159054222 | 0.704 | 0.492 | 0.004730756 | 3.20E-07 |
| White Matter Tracts | iTBS | Stip1 | stress-induced phosphoprotein 1 | 20867 | 0.159169109 | 0.482 | 0.297 | 0.000258512 | 1.75E-08 |
| White Matter Tracts | iTBS | Lin7b | lin-7 homolog B, crumbs cell polarity complex component | 22342 | 0.159416063 | 0.422 | 0.277 | 0.016234577 | 1.10E-06 |
| White Matter Tracts | iTBS | Sik3 | SIK family kinase 3 | 70661 | 0.159568444 | 0.408 | 0.23 | 0.000619079 | 4.19E-08 |
| White Matter Tracts | iTBS | Necap1 | NECAP endocytosis associated 1 | 67602 | 0.160824347 | 0.433 | 0.25 | 5.80E-05 | 3.92E-09 |
| White Matter Tracts | iTBS | Baalc | brain and acute leukemia, cytoplasmic | 118452 | 0.160880214 | 0.44 | 0.245 | 2.47E-06 | 1.67E-10 |
| White Matter Tracts | iTBS | Ddx3x | DEAD box helicase 3, X-linked | 13205 | 0.161042378 | 0.437 | 0.222 | 1.15E-07 | 7.79E-12 |
| White Matter Tracts | iTBS | Kcnab1 | potassium voltage-gated channel, shaker-related subfamily, beta membe | 16497 | 0.161123892 | 0.325 | 0.148 | 3.88E-07 | 2.63E-11 |
| White Matter Tracts | iTBS | Ctnnd2 | catenin (cadherin associated protein), delta 2 | 18163 | 0.161262773 | 0.245 | 0.097 | 3.22E-06 | 2.18E-10 |
| White Matter Tracts | iTBS | Mrpl54 | mitochondrial ribosomal protein L54 | 66047 | 0.161425265 | 0.399 | 0.213 | 2.21E-05 | 1.49E-09 |
| White Matter Tracts | iTBS | Rbbp7 | retinoblastoma binding protein 7, chromatin remodeling factor | 245688 | 0.161997572 | 0.271 | 0.107 | 2.79E-07 | 1.89E-11 |
| White Matter Tracts | iTBS | Prkcg | protein kinase C, gamma | 18752 | 0.16218912 | 0.301 | 0.142 | 4.89E-06 | 3.31E-10 |
| White Matter Tracts | iTBS | Sorl1 | sortilin-related receptor, LDLR class A repeats-containing | 20660 | 0.16232265 | 0.386 | 0.203 | 5.82E-06 | 3.94E-10 |
| White Matter Tracts | iTBS | Sf1 | splicing factor 1 | 22668 | 0.163022803 | 0.334 | 0.138 | 4.51E-08 | 3.05E-12 |
| White Matter Tracts | iTBS | Mat2b | methionine adenosyltransferase II, beta | 108645 | 0.163260058 | 0.421 | 0.23 | 2.09E-06 | 1.41E-10 |
| White Matter Tracts | iTBS | Ier5 | immediate early response 5 | 15939 | 0.163633345 | 0.455 | 0.298 | 0.002318663 | 1.57E-07 |
| White Matter Tracts | iTBS | Rpl17 | ribosomal protein L17 | 319195 | 0.16388795 | 0.998 | 0.997 | 0.001706124 | 1.15E-07 |
| White Matter Tracts | iTBS | Igfbp6 | insulin-like growth factor binding protein 6 | 16012 | 0.163986125 | 0.276 | 0.118 | 8.02E-07 | 5.43E-11 |
| White Matter Tracts | iTBS | Rps14 | ribosomal protein S14 | 20044 | 0.164294196 | 1 | 0.998 | 0.030014551 | 2.03E-06 |
| White Matter Tracts | iTBS | Emc6 | ER membrane protein complex subunit 6 | 66048 | 0.164320043 | 0.433 | 0.252 | 0.000109819 | 7.43E-09 |
| White Matter Tracts | iTBS | Nptx1 | neuronal pentraxin 1 | 18164 | 0.164474618 | 0.236 | 0.078 | 2.87E-08 | 1.94E-12 |
| White Matter Tracts | iTBS | Pnmal1 | NA | NA | 0.165140362 | 0.28 | 0.108 | 3.74E-08 | 2.53E-12 |
| White Matter Tracts | iTBS | Igip | IgA inducing protein | 109169 | 0.165702598 | 0.274 | 0.107 | 2.40E-07 | 1.62E-11 |
| White Matter Tracts | iTBS | Kif5b | kinesin family member 5B | 16573 | 0.16622801 | 0.5 | 0.293 | 0.002233998 | 1.51E-07 |
| White Matter Tracts | iTBS | Nat14 | N-acetyltransferase 14 | 269854 | 0.166395302 | 0.294 | 0.132 | 1.72E-06 | 1.16E-10 |
| White Matter Tracts | iTBS | Ppme1 | protein phosphatase methylesterase 1 | 72590 | 0.166748812 | 0.264 | 0.097 | 8.58E-08 | 5.81E-12 |
| White Matter Tracts | iTBS | Cirbp | cold inducible RNA binding protein | 12696 | 0.166859067 | 0.495 | 0.313 | 0.000620011 | 4.20E-08 |
| White Matter Tracts | iTBS | Cwc15 | CWC15 spliceosome-associated protein | 66070 | 0.16828389 | 0.439 | 0.218 | 1.63E-06 | 1.10E-10 |
| White Matter Tracts | iTBS | Ypel5 | yippee like 5 | 383295 | 0.168547733 | 0.448 | 0.278 | 0.000729973 | 4.94E-08 |
| White Matter Tracts | iTBS | AI593442 | expressed sequence AI593442 | 330941 | 0.168773648 | 0.285 | 0.137 | 4.60E-05 | 3.11E-09 |
| White Matter Tracts | iTBS | Magee1 | MAGE family member E1 | 107528 | 0.168981738 | 0.406 | 0.228 | 4.19E-05 | 2.83E-09 |
| White Matter Tracts | iTBS | Ap1b1 | adaptor protein complex AP-1, beta 1 subunit | 11764 | 0.169064171 | 0.455 | 0.287 | 0.000498821 | 3.38E-08 |
| White Matter Tracts | iTBS | Stx1a | syntaxin 1A (brain) | 20907 | 0.169242312 | 0.381 | 0.208 | 1.06E-05 | 7.15E-10 |
| White Matter Tracts | iTBS | Mrps12 | mitochondrial ribosomal protein S12 | 24030 | 0.169283599 | 0.511 | 0.35 | 0.00551951 | 3.73E-07 |
| White Matter Tracts | iTBS | Dnajc15 | DnaJ heat shock protein family (Hsp40) member C15 | 66148 | 0.169524308 | 0.487 | 0.262 | 7.55E-08 | 5.11E-12 |
| White Matter Tracts | iTBS | Celf5 | CUGBP, Elav-like family member 5 | 319586 | 0.169597805 | 0.318 | 0.155 | 8.96E-06 | 6.06E-10 |
| White Matter Tracts | iTBS | Taf10 | TATA-box binding protein associated factor 10 | 24075 | 0.1696751 | 0.662 | 0.525 | 0.041780746 | 2.83E-06 |
| White Matter Tracts | iTBS | Reep2 | receptor accessory protein 2 | 225362 | 0.169790783 | 0.437 | 0.282 | 0.001265554 | 8.56E-08 |
| White Matter Tracts | iTBS | Laptm4b | lysosomal-associated protein transmembrane 4B | 114128 | 0.171407524 | 0.431 | 0.232 | 1.09E-05 | 7.37E-10 |
| White Matter Tracts | iTBS | Ranbp1 | RAN binding protein 1 | 19385 | 0.17234073 | 0.397 | 0.193 | 1.28E-07 | 8.65E-12 |
| White Matter Tracts | iTBS | Pgp | phosphoglycolate phosphatase | 67078 | 0.172636414 | 0.486 | 0.297 | 0.000663347 | 4.49E-08 |
| White Matter Tracts | iTBS | Spin1 | spindlin 1 | 20729 | 0.17292818 | 0.491 | 0.332 | 0.000728306 | 4.93E-08 |
| White Matter Tracts | iTBS | Pfkl | phosphofructokinase, liver, B-type | 18641 | 0.173414461 | 0.282 | 0.115 | 4.02E-07 | 2.72E-11 |
| White Matter Tracts | iTBS | mt-Atp6 | NA | NA | 0.173670596 | 1 | 1 | 0.000276388 | 1.87E-08 |
| White Matter Tracts | iTBS | Nefm | neurofilament, medium polypeptide | 18040 | 0.173970214 | 0.384 | 0.213 | 2.64E-05 | 1.79E-09 |
| White Matter Tracts | iTBS | Mrpl53 | mitochondrial ribosomal protein L53 | 68499 | 0.175101296 | 0.542 | 0.367 | 0.001050317 | 7.11E-08 |
| White Matter Tracts | iTBS | Necab2 | N-terminal EF-hand calcium binding protein 2 | 117148 | 0.175233335 | 0.457 | 0.29 | 0.000306333 | 2.07E-08 |
| White Matter Tracts | iTBS | Snrpe | small nuclear ribonucleoprotein E | 20643 | 0.177286188 | 0.394 | 0.198 | 1.13E-06 | 7.62E-11 |
| White Matter Tracts | iTBS | Cmas | cytidine monophospho-N-acetylneuraminic acid synthetase | 12764 | 0.177334448 | 0.359 | 0.18 | 6.98E-06 | 4.73E-10 |
| White Matter Tracts | iTBS | Tmem223 | transmembrane protein 223 | 66836 | 0.177429837 | 0.426 | 0.22 | 1.06E-05 | 7.18E-10 |
| White Matter Tracts | iTBS | Mrpl48 | mitochondrial ribosomal protein L48 | 52443 | 0.177580221 | 0.471 | 0.287 | 0.000748181 | 5.06E-08 |
| White Matter Tracts | iTBS | Camk2g | calcium/calmodulin-dependent protein kinase II gamma | 12325 | 0.17781228 | 0.291 | 0.133 | 9.34E-06 | 6.32E-10 |
| White Matter Tracts | iTBS | Rusc1 | RUN and SH3 domain containing 1 | 72296 | 0.18011041 | 0.245 | 0.083 | 1.58E-08 | 1.07E-12 |
| White Matter Tracts | iTBS | Cdk17 | cyclin-dependent kinase 17 | 237459 | 0.180402772 | 0.309 | 0.132 | 1.31E-07 | 8.84E-12 |
| White Matter Tracts | iTBS | Srsf5 | serine and arginine-rich splicing factor 5 | 20384 | 0.182399549 | 0.572 | 0.383 | 0.000199255 | 1.35E-08 |
| White Matter Tracts | iTBS | Vps41 | VPS41 HOPS complex subunit | 218035 | 0.182444873 | 0.375 | 0.187 | 6.82E-07 | 4.62E-11 |
| White Matter Tracts | iTBS | Rpl13 | ribosomal protein L13 | 270106 | 0.182529273 | 1 | 1 | 0.041555533 | 2.81E-06 |
| White Matter Tracts | iTBS | Csnk1a1 | casein kinase 1, alpha 1 | 93687 | 0.18272649 | 0.619 | 0.44 | 0.001904821 | 1.29E-07 |
| White Matter Tracts | iTBS | Lmbrd1 | LMBR1 domain containing 1 | 68421 | 0.182776765 | 0.509 | 0.288 | 1.24E-07 | 8.41E-12 |
| White Matter Tracts | iTBS | Mapk9 | mitogen-activated protein kinase 9 | 26420 | 0.184366914 | 0.455 | 0.26 | 2.65E-06 | 1.79E-10 |
| White Matter Tracts | iTBS | Ssna1 | SS nuclear autoantigen 1 | 68475 | 0.184719824 | 0.412 | 0.207 | 3.08E-07 | 2.08E-11 |
| White Matter Tracts | iTBS | Gabbr2 | gamma-aminobutyric acid (GABA) B receptor, 2 | 242425 | 0.184881883 | 0.301 | 0.125 | 2.17E-08 | 1.47E-12 |
| White Matter Tracts | iTBS | Rps5 | ribosomal protein S5 | 20103 | 0.185417505 | 0.996 | 0.993 | 0.004613704 | 3.12E-07 |
| White Matter Tracts | iTBS | Dock3 | dedicator of cyto-kinesis 3 | 208869 | 0.186059676 | 0.271 | 0.095 | 2.19E-09 | 1.48E-13 |
| White Matter Tracts | iTBS | Gabrd | gamma-aminobutyric acid (GABA) A receptor, subunit delta | 14403 | 0.186095988 | 0.204 | 0.052 | 2.65E-10 | 1.80E-14 |
| White Matter Tracts | iTBS | Emc7 | ER membrane protein complex subunit 7 | 73024 | 0.186703811 | 0.556 | 0.357 | 0.000295127 | 2.00E-08 |
| White Matter Tracts | iTBS | Gabrg2 | gamma-aminobutyric acid (GABA) A receptor, subunit gamma 2 | 14406 | 0.186903874 | 0.404 | 0.213 | 1.86E-07 | 1.26E-11 |
| White Matter Tracts | iTBS | Dgkb | diacylglycerol kinase, beta | 217480 | 0.187934953 | 0.256 | 0.095 | 8.32E-08 | 5.63E-12 |
| White Matter Tracts | iTBS | Mmp17 | matrix metallopeptidase 17 | 23948 | 0.188657657 | 0.283 | 0.113 | 2.23E-08 | 1.51E-12 |
| White Matter Tracts | iTBS | Cadps | Ca2+-dependent secretion activator | 27062 | 0.188668931 | 0.305 | 0.13 | 5.09E-08 | 3.44E-12 |
| White Matter Tracts | iTBS | Rbis | ribosomal biogenesis factor | 69126 | 0.190074673 | 0.664 | 0.527 | 0.01145952 | 7.75E-07 |
| White Matter Tracts | iTBS | Icam5 | intercellular adhesion molecule 5, telencephalin | 15898 | 0.190396615 | 0.392 | 0.212 | 2.78E-06 | 1.88E-10 |
| White Matter Tracts | iTBS | Psmd14 | proteasome (prosome, macropain) 26S subunit, non-ATPase, 14 | 59029 | 0.190976639 | 0.329 | 0.135 | 1.13E-08 | 7.68E-13 |
| White Matter Tracts | iTBS | Ppa1 | pyrophosphatase (inorganic) 1 | 67895 | 0.191270014 | 0.329 | 0.172 | 0.000158302 | 1.07E-08 |
| White Matter Tracts | iTBS | Mrps18a | mitochondrial ribosomal protein S18A | 68565 | 0.19151733 | 0.464 | 0.267 | 3.30E-06 | 2.23E-10 |
| White Matter Tracts | iTBS | Eif3g | eukaryotic translation initiation factor 3, subunit G | 53356 | 0.191729372 | 0.294 | 0.142 | 0.000126723 | 8.58E-09 |
| White Matter Tracts | iTBS | Mettl9 | methyltransferase like 9 | 59052 | 0.191755584 | 0.404 | 0.213 | 1.45E-05 | 9.83E-10 |
| White Matter Tracts | iTBS | Nucks1 | nuclear casein kinase and cyclin-dependent kinase substrate 1 | 98415 | 0.191915506 | 0.525 | 0.338 | 0.000330606 | 2.24E-08 |
| White Matter Tracts | iTBS | Spcs2 | signal peptidase complex subunit 2 homolog (S. cerevisiae) | 66624 | 0.192481028 | 0.458 | 0.237 | 1.61E-07 | 1.09E-11 |
| White Matter Tracts | iTBS | Kcnj4 | potassium inwardly-rectifying channel, subfamily J, member 4 | 16520 | 0.19254194 | 0.289 | 0.122 | 1.45E-07 | 9.78E-12 |
| White Matter Tracts | iTBS | Vxn | vexin | 76982 | 0.192669645 | 0.314 | 0.15 | 8.18E-06 | 5.53E-10 |
| White Matter Tracts | iTBS | mt-Co3 | NA | NA | 0.193365685 | 1 | 1 | 1.53E-05 | 1.04E-09 |
| White Matter Tracts | iTBS | mt-Co1 | NA | NA | 0.1953381 | 1 | 1 | 2.66E-06 | 1.80E-10 |
| White Matter Tracts | iTBS | Dnm1l | dynamin 1-like | 74006 | 0.195449941 | 0.413 | 0.225 | 7.39E-07 | 5.00E-11 |
| White Matter Tracts | iTBS | Nbea | neurobeachin | 26422 | 0.196376627 | 0.245 | 0.082 | 7.09E-09 | 4.80E-13 |
| White Matter Tracts | iTBS | mt-Nd1 | NA | NA | 0.19656468 | 1 | 1 | 5.76E-05 | 3.90E-09 |
| White Matter Tracts | iTBS | Rpn1 | ribophorin I | 103963 | 0.197994527 | 0.466 | 0.283 | 0.000215313 | 1.46E-08 |
| White Matter Tracts | iTBS | Bola3 | bolA-like 3 (E. coli) | 78653 | 0.201455583 | 0.572 | 0.352 | 1.98E-06 | 1.34E-10 |
| White Matter Tracts | iTBS | Txndc15 | thioredoxin domain containing 15 | 69672 | 0.202001468 | 0.374 | 0.185 | 1.30E-06 | 8.78E-11 |

| | | | | | | | | | |
|---|---|---|---|---|---|---|---|---|---|
| White Matter Tracts | iTBS | Stard10 | START domain containing 10 | 56018 | 0.204303843 | 0.278 | 0.117 | 9.01E-07 | 6.10E-11 |
| White Matter Tracts | iTBS | Lysmd2 | LysM, putative peptidoglycan-binding, domain containing 2 | 70082 | 0.206707 | 0.325 | 0.128 | 1.06E-09 | 7.19E-14 |
| White Matter Tracts | iTBS | Rell2 | RELT-like 2 | 225392 | 0.207941218 | 0.336 | 0.152 | 1.09E-07 | 7.37E-12 |
| White Matter Tracts | iTBS | Dlgap1 | DLG associated protein 1 | 224997 | 0.207959641 | 0.451 | 0.263 | 1.44E-06 | 9.76E-11 |
| White Matter Tracts | iTBS | Rpl35 | ribosomal protein L35 | 66489 | 0.208400153 | 0.996 | 1 | 1.58E-05 | 1.07E-09 |
| White Matter Tracts | iTBS | Fam49a | NA | NA | 0.208689575 | 0.334 | 0.153 | 7.90E-08 | 5.35E-12 |
| White Matter Tracts | iTBS | Dpm3 | dolichyl-phosphate mannosyltransferase polypeptide 3 | 68563 | 0.212447956 | 0.323 | 0.17 | 0.000491385 | 3.33E-08 |
| White Matter Tracts | iTBS | Rpl37 | ribosomal protein L37 | 67281 | 0.212887212 | 1 | 1 | 0.014871922 | 1.01E-06 |
| White Matter Tracts | iTBS | Ogt | O-linked N-acetylglucosamine (GlcNAc) transferase (UDP-N-acetylglucosa | 108155 | 0.214371729 | 0.413 | 0.192 | 1.86E-09 | 1.26E-13 |
| White Matter Tracts | iTBS | H3f3b | H3.3 histone B | 15081 | 0.215215595 | 0.94 | 0.948 | 0.006800937 | 4.60E-07 |
| White Matter Tracts | iTBS | Ube2l3 | ubiquitin-conjugating enzyme E2L 3 | 22195 | 0.215415145 | 0.574 | 0.383 | 0.000229894 | 1.56E-08 |
| White Matter Tracts | iTBS | Ssbp4 | single stranded DNA binding protein 4 | 76900 | 0.217127796 | 0.415 | 0.22 | 1.18E-07 | 7.97E-12 |
| White Matter Tracts | iTBS | Polr2i | polymerase (RNA) II (DNA directed) polypeptide I | 69920 | 0.217591435 | 0.348 | 0.142 | 3.79E-09 | 2.56E-13 |
| White Matter Tracts | iTBS | Pin4 | peptidyl-prolyl cis/trans isomerase, NIMA-interacting, 4 (parvulin) | 69713 | 0.217629176 | 0.426 | 0.215 | 1.20E-07 | 8.13E-12 |
| White Matter Tracts | iTBS | Rps27a | ribosomal protein S27A | 78294 | 0.221362634 | 1 | 1 | 0.000273954 | 1.85E-08 |
| White Matter Tracts | iTBS | Ndufaf8 | NADH:ubiquinone oxidoreductase complex assembly factor 8 | 208501 | 0.221457001 | 0.345 | 0.142 | 6.48E-09 | 4.39E-13 |
| White Matter Tracts | iTBS | Rpl34 | ribosomal protein L34 | 68436 | 0.222874321 | 1 | 1 | 0.030868616 | 2.09E-06 |
| White Matter Tracts | iTBS | Snrpb | small nuclear ribonucleoprotein B | 20638 | 0.226359858 | 0.451 | 0.238 | 6.36E-07 | 4.31E-11 |
| White Matter Tracts | iTBS | Araf | Araf proto-oncogene, serine/threonine kinase | 11836 | 0.227405225 | 0.316 | 0.13 | 2.02E-08 | 1.36E-12 |
| White Matter Tracts | iTBS | Rpl39 | ribosomal protein L39 | 67248 | 0.2274344 | 0.998 | 0.998 | 0.01107004 | 7.49E-07 |
| White Matter Tracts | iTBS | Arhgef17 | Rho guanine nucleotide exchange factor (GEF) 17 | 207212 | 0.2294092 | 0.267 | 0.09 | 3.02E-10 | 2.04E-14 |
| White Matter Tracts | iTBS | Rps8 | ribosomal protein S8 | 20116 | 0.231711363 | 1 | 1 | 4.87E-05 | 3.30E-09 |
| White Matter Tracts | iTBS | Krtcap2 | keratinocyte associated protein 2 | 66059 | 0.231751695 | 0.545 | 0.362 | 0.000478247 | 3.24E-08 |
| White Matter Tracts | iTBS | Polr2j | polymerase (RNA) II (DNA directed) polypeptide J | 20022 | 0.232463332 | 0.31 | 0.12 | 5.44E-09 | 3.68E-13 |
| White Matter Tracts | iTBS | Tmem179 | transmembrane protein 179 | 104885 | 0.232612844 | 0.392 | 0.217 | 6.55E-06 | 4.43E-10 |
| White Matter Tracts | iTBS | Tceal9 | transcription elongation factor A like 9 | 22381 | 0.23551853 | 0.442 | 0.223 | 5.37E-10 | 3.64E-14 |
| White Matter Tracts | iTBS | Rpl9 | ribosomal protein L9 | 20005 | 0.235973109 | 1 | 0.998 | 3.12E-05 | 2.11E-09 |
| White Matter Tracts | iTBS | Pdp1 | pyruvate dehydrogenase phosphatase catalytic subunit 1 | 381511 | 0.236282519 | 0.323 | 0.127 | 6.58E-11 | 4.45E-15 |
| White Matter Tracts | iTBS | Lgmn | legumain | 19141 | 0.236652584 | 0.625 | 0.405 | 3.13E-06 | 2.12E-10 |
| White Matter Tracts | iTBS | Tmbim6 | transmembrane BAX inhibitor motif containing 6 | 110213 | 0.236939758 | 0.431 | 0.223 | 4.41E-06 | 2.98E-10 |
| White Matter Tracts | iTBS | Rps26 | ribosomal protein S26 | 27370 | 0.237632036 | 0.996 | 0.997 | 0.000484328 | 3.28E-08 |
| White Matter Tracts | iTBS | Commd3 | COMM domain containing 3 | 12238 | 0.241499887 | 0.399 | 0.185 | 5.69E-09 | 3.85E-13 |
| White Matter Tracts | iTBS | Psma4 | proteasome subunit alpha 4 | 26441 | 0.243990046 | 0.37 | 0.153 | 4.09E-10 | 2.77E-14 |
| White Matter Tracts | iTBS | Sec61b | Sec61 beta subunit | 66212 | 0.248009655 | 0.532 | 0.297 | 2.11E-09 | 1.43E-13 |
| White Matter Tracts | iTBS | Cdk16 | cyclin-dependent kinase 16 | 18555 | 0.248600197 | 0.493 | 0.283 | 8.07E-08 | 5.46E-12 |
| White Matter Tracts | iTBS | Rps12 | ribosomal protein S12 | 20042 | 0.250664118 | 1 | 0.998 | 0.000138898 | 9.40E-09 |
| White Matter Tracts | iTBS | Banf1 | BAF nuclear assembly factor 1 | 23825 | 0.252885736 | 0.493 | 0.295 | 1.16E-05 | 7.82E-10 |
| White Matter Tracts | iTBS | Cd47 | CD47 antigen (Rh-related antigen, integrin-associated signal transducer) | 16423 | 0.25602735 | 0.319 | 0.128 | 8.88E-09 | 6.01E-13 |
| White Matter Tracts | iTBS | Cct3 | chaperonin containing Tcp1, subunit 3 (gamma) | 12462 | 0.257764942 | 0.439 | 0.213 | 8.20E-10 | 5.55E-14 |
| White Matter Tracts | iTBS | Rpl30 | ribosomal protein L30 | 19946 | 0.267108506 | 1 | 1 | 9.13E-05 | 6.18E-09 |
| White Matter Tracts | iTBS | Hexb | hexosaminidase B | 15212 | 0.26767016 | 0.44 | 0.222 | 3.43E-07 | 2.32E-11 |
| White Matter Tracts | iTBS | Rpl23 | ribosomal protein L23 | 65019 | 0.26928313 | 1 | 1 | 8.69E-08 | 5.88E-12 |
| White Matter Tracts | iTBS | Cab39 | calcium binding protein 39 | 12283 | 0.269462269 | 0.307 | 0.108 | 4.10E-11 | 2.78E-15 |
| White Matter Tracts | iTBS | Crym | crystallin, mu | 12971 | 0.270578122 | 0.197 | 0.053 | 4.40E-09 | 2.98E-13 |
| White Matter Tracts | iTBS | Sf3b1 | splicing factor 3b, subunit 1 | 81898 | 0.273697497 | 0.384 | 0.173 | 1.53E-09 | 1.03E-13 |
| White Matter Tracts | iTBS | Rpl35a | ribosomal protein L35A | 57808 | 0.274475287 | 1 | 1 | 2.00E-05 | 1.35E-09 |
| White Matter Tracts | iTBS | Rps10 | ribosomal protein S10 | 67097 | 0.274498159 | 0.998 | 1 | 6.58E-06 | 4.46E-10 |
| White Matter Tracts | iTBS | Enpp2 | ectonucleotide pyrophosphatase/phosphodiesterase 2 | 18606 | 0.282580977 | 0.666 | 0.468 | 0.012315097 | 8.33E-07 |
| White Matter Tracts | iTBS | mt-Co2 | NA | NA | 0.296882088 | 1 | 1 | 8.97E-06 | 6.07E-10 |
| White Matter Tracts | iTBS | Snhg8 | small nucleolar RNA host gene 8 | 69895 | 0.308124984 | 0.5 | 0.237 | 6.81E-12 | 4.61E-16 |
| White Matter Tracts | iTBS | Rps24 | ribosomal protein S24 | 20088 | 0.312858966 | 1 | 1 | 1.35E-08 | 9.11E-13 |
| White Matter Tracts | iTBS | Rps21 | ribosomal protein S21 | 66481 | 0.327145778 | 1 | 1 | 3.90E-11 | 2.64E-15 |
| White Matter Tracts | iTBS | Rps28 | ribosomal protein S28 | 54127 | 0.334807146 | 0.987 | 0.995 | 1.79E-06 | 1.21E-10 |
| White Matter Tracts | iTBS | Gstp1 | glutathione S-transferase, pi 1 | 14870 | 0.538358018 | 0.825 | 0.597 | 1.39E-12 | 9.40E-17 |
| White Matter Tracts | iTBS | Uba52 | ubiquitin A-52 residue ribosomal protein fusion product 1 | 22186 | 0.57659486 | 0.991 | 0.958 | 1.98E-23 | 1.34E-27 |
| White Matter Tracts | iTBS | Ttr | transthyretin | 22139 | 0.609096865 | 0.3 | 0.078 | 3.26E-17 | 2.21E-21 |
| White Matter Tracts | iTBS | Gm10076 | ribosomal protein L41 pseudogene | 100126819 | 0.847062151 | 0.975 | 0.887 | 2.48E-46 | 1.68E-50 |
| Caudoputamen | iTBS | Dnm1 | dynamin 1 | 13429 | -0.303727691 | 0.849 | 0.944 | 1.04E-13 | 7.05E-18 |
| Caudoputamen | iTBS | Hpcal4 | hippocalcin-like 4 | 170638 | -0.289905788 | 0.525 | 0.596 | 1.98E-06 | 1.34E-10 |
| Caudoputamen | iTBS | Bc1 | brain cytoplasmic RNA 1 | 100568459 | -0.284200548 | 0.537 | 0.693 | 1.03E-08 | 6.97E-13 |
| Caudoputamen | iTBS | Vsnl1 | visinin-like 1 | 26950 | -0.281697653 | 0.333 | 0.544 | 1.47E-17 | 9.96E-22 |
| Caudoputamen | iTBS | Gm42418 | NA | NA | -0.277445049 | 0.926 | 0.963 | 1.13E-10 | 7.63E-15 |
| Caudoputamen | iTBS | Glul | glutamate-ammonia ligase (glutamine synthetase) | 14645 | -0.266010832 | 0.738 | 0.835 | 5.56E-09 | 3.77E-13 |
| Caudoputamen | iTBS | Olfm1 | olfactomedin 1 | 56177 | -0.262932448 | 0.6 | 0.764 | 1.48E-12 | 1.00E-16 |
| Caudoputamen | iTBS | Tuba1b | tubulin, alpha 1B | 22143 | -0.256390985 | 0.671 | 0.833 | 1.34E-11 | 9.08E-16 |
| Caudoputamen | iTBS | Sod1 | superoxide dismutase 1, soluble | 20655 | -0.246012289 | 0.749 | 0.812 | 2.35E-07 | 1.59E-11 |
| Caudoputamen | iTBS | Cplx2 | complexin 2 | 12890 | -0.244154702 | 0.878 | 0.899 | 1.49E-05 | 1.01E-09 |
| Caudoputamen | iTBS | Tubb4b | tubulin, beta 4B class IVB | 227613 | -0.233490235 | 0.538 | 0.625 | 3.16E-06 | 2.14E-10 |
| Caudoputamen | iTBS | Cdk5r2 | cyclin-dependent kinase 5, regulatory subunit 2 (p39) | 12570 | -0.230927658 | 0.335 | 0.433 | 8.57E-05 | 5.80E-09 |
| Caudoputamen | iTBS | Kif1a | kinesin family member 1A | 16560 | -0.227441191 | 0.639 | 0.695 | 0.000421954 | 2.86E-08 |
| Caudoputamen | iTBS | Atp1a3 | ATPase, Na+/K+ transporting, alpha 3 polypeptide | 232975 | -0.219844408 | 0.66 | 0.733 | 1.78E-05 | 1.20E-09 |
| Caudoputamen | iTBS | Ivns1abp | influenza virus NS1A binding protein | 117198 | -0.216426233 | 0.909 | 0.876 | 0.03353135 | 2.27E-06 |
| Caudoputamen | iTBS | Slc17a7 | solute carrier family 17 (sodium-dependent inorganic phosphate cotrans | 72961 | -0.216358944 | 0.217 | 0.361 | 2.65E-09 | 1.79E-13 |
| Caudoputamen | iTBS | Tubb2a | tubulin, beta 2A class IIA | 22151 | -0.214976534 | 0.582 | 0.713 | 2.56E-06 | 1.73E-10 |
| Caudoputamen | iTBS | Rab6b | RAB6B, member RAS oncogene family | 270192 | -0.210451714 | 0.879 | 0.917 | 1.14E-07 | 7.72E-12 |
| Caudoputamen | iTBS | Tuba1a | tubulin, alpha 1A | 22142 | -0.209871779 | 0.948 | 0.984 | 4.35E-09 | 2.94E-13 |
| Caudoputamen | iTBS | Slc1a2 | solute carrier family 1 (glial high affinity glutamate transporter), member | 20511 | -0.209030992 | 0.975 | 0.991 | 5.08E-08 | 3.44E-12 |
| Caudoputamen | iTBS | Acot7 | acyl-CoA thioesterase 7 | 70025 | -0.20437813 | 0.597 | 0.641 | 0.019396537 | 1.31E-06 |
| Caudoputamen | iTBS | Atp1a1 | ATPase, Na+/K+ transporting, alpha 1 polypeptide | 11928 | -0.201915106 | 0.886 | 0.954 | 9.56E-06 | 6.47E-10 |
| Caudoputamen | iTBS | Tuba4a | tubulin, alpha 4A | 22145 | -0.199776266 | 0.878 | 0.915 | 3.98E-05 | 2.69E-09 |
| Caudoputamen | iTBS | Aldoc | aldolase C, fructose-bisphosphate | 11676 | -0.199715765 | 0.981 | 0.997 | 5.30E-08 | 3.59E-12 |
| Caudoputamen | iTBS | Stxbp1 | syntaxin binding protein 1 | 20910 | -0.199575621 | 0.683 | 0.782 | 2.49E-05 | 1.69E-09 |
| Caudoputamen | iTBS | Gng3 | guanine nucleotide binding protein (G protein), gamma 3 | 14704 | -0.197950328 | 0.762 | 0.857 | 1.22E-05 | 8.29E-10 |
| Caudoputamen | iTBS | Atp5o | ATP synthase, H+ transporting, mitochondrial F1 complex, O subunit | 28080 | -0.197903568 | 0.926 | 0.951 | 5.17E-05 | 3.50E-09 |
| Caudoputamen | iTBS | Syn1 | synapsin I | 20964 | -0.197031388 | 0.614 | 0.669 | 0.022272264 | 1.51E-06 |
| Caudoputamen | iTBS | Kif5a | kinesin family member 5A | 16572 | -0.195500376 | 0.753 | 0.84 | 0.000176307 | 1.19E-08 |
| Caudoputamen | iTBS | Stmn1 | stathmin 1 | 16765 | -0.193601561 | 0.628 | 0.772 | 1.30E-05 | 8.82E-10 |
| Caudoputamen | iTBS | Gnas | GNAS (guanine nucleotide binding protein, alpha stimulating) complex lo | 14683 | -0.192941582 | 0.786 | 0.9 | 8.15E-07 | 5.52E-11 |
| Caudoputamen | iTBS | Prkar1b | protein kinase, cAMP dependent regulatory, type I beta | 19085 | -0.191330678 | 0.616 | 0.728 | 2.23E-05 | 1.51E-09 |
| Caudoputamen | iTBS | Eef1a2 | eukaryotic translation elongation factor 1 alpha 2 | 13628 | -0.191160339 | 0.97 | 0.988 | 0.009589435 | 6.49E-07 |
| Caudoputamen | iTBS | Syp | synaptophysin | 20977 | -0.189262044 | 0.551 | 0.642 | 0.006419589 | 4.34E-07 |
| Caudoputamen | iTBS | Actb | actin, beta | 11461 | -0.18789546 | 0.999 | 1 | 2.02E-06 | 1.36E-10 |
| Caudoputamen | iTBS | Epn1 | epsin 1 | 13854 | -0.186174956 | 0.68 | 0.705 | 0.013616759 | 9.21E-07 |
| Caudoputamen | iTBS | Apbb1 | amyloid beta (A4) precursor protein-binding, family B, member 1 | 11785 | -0.183486734 | 0.738 | 0.827 | 0.000569994 | 3.86E-08 |
| Caudoputamen | iTBS | Ywhag | tyrosine 3-monooxygenase/tryptophan 5-monooxygenase activation pro | 22628 | -0.183181764 | 0.756 | 0.844 | 0.000435323 | 2.95E-08 |
| Caudoputamen | iTBS | Sptbn2 | spectrin beta, non-erythrocytic 2 | 20743 | -0.181517182 | 0.707 | 0.734 | 0.042016382 | 2.84E-06 |
| Caudoputamen | iTBS | Eef2 | eukaryotic translation elongation factor 2 | 13629 | -0.179894337 | 0.95 | 0.979 | 0.000115754 | 7.83E-09 |
| Caudoputamen | iTBS | Ikbkb | inhibitor of kappaB kinase beta | 16150 | -0.179439433 | 0.225 | 0.32 | 0.001910955 | 1.29E-07 |
| Caudoputamen | iTBS | Cpe | carboxypeptidase E | 12876 | -0.171715894 | 1 | 1 | 1.88E-09 | 1.27E-13 |
| Caudoputamen | iTBS | Ndrg2 | N-myc downstream regulated gene 2 | 29811 | -0.166889928 | 0.891 | 0.932 | 0.001434138 | 9.70E-08 |
| Caudoputamen | iTBS | Clu | clusterin | 12759 | -0.161061623 | 0.982 | 0.993 | 0.010232243 | 6.92E-07 |
| Caudoputamen | iTBS | Tcf25 | transcription factor 25 (basic helix-loop-helix) | 66855 | -0.159615308 | 0.848 | 0.894 | 0.012521108 | 8.47E-07 |
| Caudoputamen | iTBS | Vamp2 | vesicle-associated membrane protein 2 | 22318 | -0.156580317 | 0.972 | 0.988 | 0.000270976 | 1.83E-08 |
| Caudoputamen | iTBS | Ywhah | tyrosine 3-monooxygenase/tryptophan 5-monooxygenase activation pro | 22629 | -0.155143131 | 0.915 | 0.958 | 0.004849813 | 3.28E-07 |
| Caudoputamen | iTBS | Rpl5 | ribosomal protein L5 | 100503670 | -0.154968169 | 0.945 | 0.962 | 0.022043917 | 1.49E-06 |
| Caudoputamen | iTBS | Thy1 | thymus cell antigen 1, theta | 21838 | -0.150630978 | 0.897 | 0.947 | 0.015265346 | 1.03E-06 |
| Caudoputamen | iTBS | Snhg9 | small nucleolar RNA host gene 9 | 73474 | -0.149087391 | 0.147 | 0.252 | 4.39E-05 | 2.97E-09 |
| Caudoputamen | iTBS | Hspa8 | heat shock protein 8 | 15481 | -0.137501767 | 1 | 1 | 4.21E-05 | 2.85E-09 |
| Caudoputamen | iTBS | Rpl4 | ribosomal protein L4 | 67891 | -0.136803021 | 0.986 | 0.996 | 0.021725848 | 1.47E-06 |
| Caudoputamen | iTBS | Rpl41 | ribosomal protein L41 | 67945 | -0.124300787 | 1 | 1 | 2.32E-09 | 1.57E-13 |
| Caudoputamen | iTBS | Cck | cholecystokinin | 12424 | -0.123452561 | 0.308 | 0.416 | 0.028189879 | 1.91E-06 |
| Caudoputamen | iTBS | Ywhaz | tyrosine 3-monooxygenase/tryptophan 5-monooxygenase activation pro | 22631 | -0.122034389 | 1 | 0.997 | 0.000760287 | 5.14E-08 |
| Caudoputamen | iTBS | Aldoa | aldolase A, fructose-bisphosphate | 11674 | -0.109230356 | 0.999 | 1 | 0.014494378 | 9.81E-07 |
| Caudoputamen | iTBS | Ndrg4 | N-myc downstream regulated gene 4 | 234593 | -0.108903091 | 0.999 | 0.999 | 0.037846313 | 2.56E-06 |
| Caudoputamen | iTBS | Gapdh | glyceraldehyde-3-phosphate dehydrogenase | 14433 | -0.099323276 | 1 | 1 | 0.000161654 | 1.09E-08 |
| Caudoputamen | iTBS | Zfp763 | zinc finger protein 763 | 73451 | 0.077897838 | 0.102 | 0.046 | 0.026290066 | 1.78E-06 |
| Caudoputamen | iTBS | Hmga1 | high mobility group AT-hook 1 | 15361 | 0.078127536 | 0.114 | 0.053 | 0.01167317 | 7.90E-07 |
| Caudoputamen | iTBS | Ddost | dolichyl-di-phosphooligosaccharide-protein glycotransferase | 13200 | 0.082293064 | 0.509 | 0.373 | 0.045826336 | 3.10E-06 |
| Caudoputamen | iTBS | Cpne2 | copine II | 234577 | 0.083188673 | 0.126 | 0.065 | 0.046409945 | 3.14E-06 |
| Caudoputamen | iTBS | Cog6 | component of oligomeric golgi complex 6 | 67542 | 0.08523283 | 0.192 | 0.115 | 0.03397837 | 2.30E-06 |
| Caudoputamen | iTBS | Ahcy | S-adenosylhomocysteine hydrolase | 269378 | 0.086149993 | 0.103 | 0.037 | 8.65E-05 | 5.85E-09 |
| Caudoputamen | iTBS | Golga4 | golgi autoantigen, golgin subfamily a, 4 | 54214 | 0.087169102 | 0.235 | 0.152 | 0.043637274 | 2.95E-06 |
| Caudoputamen | iTBS | Coro1b | coronin, actin binding protein 1B | 23789 | 0.089192604 | 0.1 | 0.035 | 0.000112076 | 7.58E-09 |
| Caudoputamen | iTBS | P4ha1 | procollagen-proline, 2-oxoglutarate 4-dioxygenase (proline 4-hydroxylase | 18451 | 0.090141434 | 0.144 | 0.079 | 0.042829052 | 2.90E-06 |
| Caudoputamen | iTBS | Tipin | timeless interacting protein | 66131 | 0.090544929 | 0.138 | 0.073 | 0.022986409 | 1.56E-06 |
| Caudoputamen | iTBS | Anapc10 | anaphase promoting complex subunit 10 | 68999 | 0.091669227 | 0.135 | 0.063 | 0.0008655 | 5.86E-08 |
| Caudoputamen | iTBS | Gpr139 | G protein-coupled receptor 139 | 209776 | 0.092920896 | 0.12 | 0.052 | 0.000842863 | 5.70E-08 |
| Caudoputamen | iTBS | Cdh9 | cadherin 9 | 12565 | 0.095266466 | 0.167 | 0.095 | 0.027263545 | 1.84E-06 |
| Caudoputamen | iTBS | Isca2 | iron-sulfur cluster assembly 2 | 74316 | 0.096919967 | 0.334 | 0.232 | 0.041702368 | 2.82E-06 |
| Caudoputamen | iTBS | Smad3 | SMAD family member 3 | 17127 | 0.098166277 | 0.228 | 0.139 | 0.006798992 | 4.60E-07 |
| Caudoputamen | iTBS | Ank | progressive ankylosis | 11732 | 0.098238902 | 0.356 | 0.254 | 0.0430586 | 2.91E-06 |
| Caudoputamen | iTBS | Fndc10 | fibronectin type III domain containing 10 | 230991 | 0.098277046 | 0.177 | 0.1 | 0.009143197 | 6.19E-07 |
| Caudoputamen | iTBS | Ppia | peptidylprolyl isomerase A | 268373 | 0.098815002 | 1 | 1 | 0.003784163 | 2.56E-07 |
| Caudoputamen | iTBS | Atp5l | ATP synthase, H+ transporting, mitochondrial F0 complex, subunit G | 27425 | 0.100010126 | 0.999 | 1 | 0.039733504 | 2.69E-06 |
| Caudoputamen | iTBS | Ppp5c | protein phosphatase 5, catalytic subunit | 19060 | 0.100195099 | 0.307 | 0.195 | 0.002120992 | 1.44E-07 |
| Caudoputamen | iTBS | Ankrd37 | ankyrin repeat domain 37 | 654824 | 0.100295449 | 0.243 | 0.15 | 0.005886759 | 3.98E-07 |
| Caudoputamen | iTBS | Galnt14 | polypeptide N-acetylgalactosaminyltransferase 14 | 71685 | 0.100303405 | 0.162 | 0.085 | 0.001655126 | 1.12E-07 |
| Caudoputamen | iTBS | Eif2b2 | eukaryotic translation initiation factor 2B, subunit 2 beta | 217715 | 0.100397824 | 0.226 | 0.142 | 0.019842673 | 1.34E-06 |
| Caudoputamen | iTBS | Rarb | retinoic acid receptor, beta | 218772 | 0.100676302 | 0.274 | 0.184 | 0.047086253 | 3.19E-06 |
| Caudoputamen | iTBS | Abhd11 | abhydrolase domain containing 11 | 68758 | 0.101265268 | 0.171 | 0.092 | 0.002224255 | 1.51E-07 |
| Caudoputamen | iTBS | Gm19410 | predicted gene, 19410 | 100502846 | 0.101655422 | 0.194 | 0.115 | 0.017292213 | 1.17E-06 |
| Caudoputamen | iTBS | Akap9 | A kinase (PRKA) anchor protein (yotiao) 9 | 100986 | 0.101860451 | 0.42 | 0.308 | 0.03987716 | 2.70E-06 |
| Caudoputamen | iTBS | Rplp1 | ribosomal protein, large, P1 | 56040 | 0.102400525 | 1 | 1 | 0.002393604 | 1.62E-07 |
| Caudoputamen | iTBS | Rab10os | RAB10, member RAS oncogene family, opposite strand | 74173 | 0.102769042 | 0.165 | 0.095 | 0.032747711 | 2.22E-06 |
| Caudoputamen | iTBS | Tiam2 | T cell lymphoma invasion and metastasis 2 | 24001 | 0.103161602 | 0.4 | 0.29 | 0.037208161 | 2.52E-06 |
| Caudoputamen | iTBS | Plvap | plasmalemma vesicle associated protein | 84094 | 0.10372679 | 0.165 | 0.093 | 0.014556594 | 9.85E-07 |
| Caudoputamen | iTBS | Rpl21 | ribosomal protein L21 | 19933 | 0.103985966 | 0.999 | 1 | 0.000693031 | 4.69E-08 |
| Caudoputamen | iTBS | Unc5a | unc-5 netrin receptor A | 107448 | 0.104156456 | 0.221 | 0.14 | 0.040569179 | 2.75E-06 |
| Caudoputamen | iTBS | Il17rc | interleukin 17 receptor C | 171095 | 0.104657102 | 0.156 | 0.076 | 0.00031268 | 2.12E-08 |
| Caudoputamen | iTBS | Tgfa | transforming growth factor alpha | 21802 | 0.104886794 | 0.495 | 0.366 | 0.009912524 | 6.71E-07 |
| Caudoputamen | iTBS | Zbtb8os | zinc finger and BTB domain containing 8 opposite strand | 67106 | 0.105259371 | 0.282 | 0.192 | 0.04892575 | 3.31E-06 |
| Caudoputamen | iTBS | Psme2 | proteasome (prosome, macropain) activator subunit 2 (PA28 beta) | 19188 | 0.105583709 | 0.211 | 0.13 | 0.018245344 | 1.23E-06 |
| Caudoputamen | iTBS | Ptrhd1 | peptidyl-tRNA hydrolase domain containing 1 | 69709 | 0.106508872 | 0.284 | 0.19 | 0.030893527 | 2.09E-06 |
| Caudoputamen | iTBS | Lrtm2 | leucine-rich repeats and transmembrane domains 2 | 211187 | 0.106780954 | 0.368 | 0.263 | 0.025091092 | 1.70E-06 |
| Caudoputamen | iTBS | Tmem80 | transmembrane protein 80 | 71448 | 0.106816712 | 0.152 | 0.08 | 0.004751909 | 3.22E-07 |
| Caudoputamen | iTBS | Ddx42 | DEAD box helicase 42 | 72047 | 0.107261537 | 0.246 | 0.156 | 0.01257811 | 8.51E-07 |
| Caudoputamen | iTBS | Dnal4 | dynein, axonemal, light chain 4 | 54152 | 0.107574629 | 0.414 | 0.306 | 0.034529875 | 2.34E-06 |
| Caudoputamen | iTBS | Plekhj1 | pleckstrin homology domain containing, family J member 1 | 78670 | 0.107576027 | 0.335 | 0.227 | 0.006370798 | 4.31E-07 |
| Caudoputamen | iTBS | Gadd45b | growth arrest and DNA-damage-inducible 45 beta | 17873 | 0.107930747 | 0.193 | 0.113 | 0.007339985 | 4.97E-07 |
| Caudoputamen | iTBS | Tpgs1 | tubulin polyglutamylase complex subunit 1 | 110012 | 0.108673417 | 0.409 | 0.296 | 0.024006571 | 1.62E-06 |
| Caudoputamen | iTBS | Klf9 | Kruppel-like factor 9 | 16601 | 0.108695548 | 0.537 | 0.409 | 0.042480767 | 2.87E-06 |
| Caudoputamen | iTBS | Mbd2 | methyl-CpG binding domain protein 2 | 17191 | 0.108892608 | 0.34 | 0.233 | 0.01345581 | 9.11E-07 |
| Caudoputamen | iTBS | Zfp580 | zinc finger protein 580 | 68992 | 0.109174152 | 0.269 | 0.178 | 0.022031222 | 1.49E-06 |
| Caudoputamen | iTBS | Sema7a | sema domain, immunoglobulin domain (Ig), and GPI membrane anchor, ( | 20361 | 0.109389318 | 0.187 | 0.1 | 0.000631621 | 4.27E-08 |
| Caudoputamen | iTBS | Bckdk | branched chain ketoacid dehydrogenase kinase | 12041 | 0.109529065 | 0.241 | 0.152 | 0.008726518 | 5.91E-07 |
| Caudoputamen | iTBS | Slc25a33 | solute carrier family 25, member 33 | 70556 | 0.109972859 | 0.448 | 0.333 | 0.04846185 | 3.28E-06 |
| Caudoputamen | iTBS | Acbd4 | acyl-Coenzyme A binding domain containing 4 | 67131 | 0.110255917 | 0.222 | 0.132 | 0.001893859 | 1.28E-07 |
| Caudoputamen | iTBS | Son | Son DNA binding protein | 20658 | 0.110260894 | 0.512 | 0.38 | 0.016601263 | 1.12E-06 |
| Caudoputamen | iTBS | Cox6a1 | cytochrome c oxidase subunit 6A1 | 12861 | 0.110352573 | 0.999 | 0.999 | 0.035053713 | 2.37E-06 |
| Caudoputamen | iTBS | Rp9 | retinitis pigmentosa 9 (human) | 55934 | 0.11039729 | 0.233 | 0.144 | 0.005583884 | 3.78E-07 |
| Caudoputamen | iTBS | Atg3 | autophagy related 3 | 67841 | 0.110412103 | 0.243 | 0.154 | 0.009818819 | 6.64E-07 |
| Caudoputamen | iTBS | Epdr1 | ependymin related protein 1 (zebrafish) | 105298 | 0.111502674 | 0.5 | 0.37 | 0.01652626 | 1.12E-06 |
| Caudoputamen | iTBS | Eef1a1 | eukaryotic translation elongation factor 1 alpha 1 | 13627 | 0.111781941 | 1 | 1 | 5.57E-05 | 3.77E-09 |
| Caudoputamen | iTBS | Rps11 | ribosomal protein S11 | 27207 | 0.111797549 | 0.986 | 0.989 | 0.011380849 | 7.70E-07 |
| Caudoputamen | iTBS | Tmco1 | transmembrane and coiled-coil domains 1 | 68944 | 0.111966261 | 0.317 | 0.21 | 0.005086142 | 3.44E-07 |
| Caudoputamen | iTBS | Mrpl36 | mitochondrial ribosomal protein L36 | 94066 | 0.112548952 | 0.376 | 0.27 | 0.013791998 | 9.33E-07 |
| Caudoputamen | iTBS | Osbpl2 | oxysterol binding protein-like 2 | 228983 | 0.112912137 | 0.274 | 0.177 | 0.006278747 | 4.25E-07 |
| Caudoputamen | iTBS | Rps5 | ribosomal protein S5 | 20103 | 0.113006038 | 0.969 | 0.957 | 0.008831814 | 5.98E-07 |
| Caudoputamen | iTBS | Hnrnpm | heterogeneous nuclear ribonucleoprotein M | 76936 | 0.113532446 | 0.526 | 0.379 | 0.002082768 | 1.41E-07 |
| Caudoputamen | iTBS | Adgrb3 | adhesion G protein-coupled receptor B3 | 210933 | 0.113802631 | 0.368 | 0.256 | 0.00867006 | 5.87E-07 |
| Caudoputamen | iTBS | Mettl23 | methyltransferase like 23 | 74319 | 0.113877869 | 0.332 | 0.226 | 0.009557195 | 6.47E-07 |
| Caudoputamen | iTBS | Tmem30a | transmembrane protein 30A | 69981 | 0.113963254 | 0.616 | 0.477 | 0.037234799 | 2.52E-06 |
| Caudoputamen | iTBS | Ckb | creatine kinase, brain | 12709 | 0.115592579 | 1 | 1 | 1.17E-07 | 7.95E-12 |
| Caudoputamen | iTBS | Dctn6 | dynactin 6 | 22428 | 0.115673336 | 0.6 | 0.463 | 0.036378754 | 2.46E-06 |
| Caudoputamen | iTBS | Selenos | selenoprotein S | 109815 | 0.116648163 | 0.314 | 0.215 | 0.007837009 | 5.30E-07 |
| Caudoputamen | iTBS | Acy1 | aminoacylase 1 | 109652 | 0.116648936 | 0.341 | 0.229 | 0.002970014 | 2.01E-07 |
| Caudoputamen | iTBS | Skap2 | src family associated phosphoprotein 2 | 54353 | 0.117718353 | 0.193 | 0.1 | 6.28E-05 | 4.25E-09 |
| Caudoputamen | iTBS | Lingo3 | leucine rich repeat and Ig domain containing 3 | 237403 | 0.118744235 | 0.497 | 0.385 | 0.028074716 | 1.90E-06 |
| Caudoputamen | iTBS | Fam174a | family with sequence similarity 174, member A | 67698 | 0.118910595 | 0.43 | 0.317 | 0.021849317 | 1.48E-06 |
| Caudoputamen | iTBS | Chd9 | chromodomain helicase DNA binding protein 9 | 109151 | 0.119514738 | 0.235 | 0.148 | 0.008978984 | 6.08E-07 |
| Caudoputamen | iTBS | Gcnt2 | glucosaminyl (N-acetyl) transferase 2, I-branching enzyme | 14538 | 0.119575922 | 0.421 | 0.309 | 0.013918204 | 9.42E-07 |
| Caudoputamen | iTBS | Tomm5 | translocase of outer mitochondrial membrane 5 | 68512 | 0.1202974 | 0.54 | 0.405 | 0.024975071 | 1.69E-06 |
| Caudoputamen | iTBS | Denr | density-regulated protein | 68184 | 0.120550652 | 0.492 | 0.369 | 0.00785197 | 5.31E-07 |
| Caudoputamen | iTBS | Slc6a6 | solute carrier family 6 (neurotransmitter transporter, taurine), member 6 | 21366 | 0.120702391 | 0.347 | 0.242 | 0.013368328 | 9.05E-07 |
| Caudoputamen | iTBS | Isca1 | iron-sulfur cluster assembly 1 | 69046 | 0.121411584 | 0.444 | 0.334 | 0.037742048 | 2.55E-06 |
| Caudoputamen | iTBS | Opalin | oligodendrocytic myelin paranodal and inner loop protein | 226115 | 0.121649235 | 0.3 | 0.205 | 0.030735762 | 2.08E-06 |
| Caudoputamen | iTBS | Cox5b | cytochrome c oxidase subunit 5B | 12859 | 0.122580722 | 1 | 0.999 | 0.000282201 | 1.91E-08 |
| Caudoputamen | iTBS | Sdhaf4 | succinate dehydrogenase complex assembly factor 4 | 68002 | 0.122626145 | 0.319 | 0.206 | 0.00040243 | 2.72E-08 |
| Caudoputamen | iTBS | Leprot | leptin receptor overlapping transcript | 230514 | 0.122762855 | 0.199 | 0.115 | 0.002333345 | 1.58E-07 |
| Caudoputamen | iTBS | Spint2 | serine protease inhibitor, Kunitz type 2 | 20733 | 0.123199722 | 0.254 | 0.157 | 0.001771142 | 1.20E-07 |
| Caudoputamen | iTBS | Lsm4 | LSM4 homolog, U6 small nuclear RNA and mRNA degradation associated | 50783 | 0.123472387 | 0.485 | 0.359 | 0.008412661 | 5.69E-07 |
| Caudoputamen | iTBS | Jkamp | JNK1/MAPK8-associated membrane protein | 104771 | 0.123817822 | 0.31 | 0.2 | 0.000904098 | 6.12E-08 |
| Caudoputamen | iTBS | Fut8 | fucosyltransferase 8 | 53618 | 0.124022148 | 0.245 | 0.151 | 0.002132619 | 1.44E-07 |
| Caudoputamen | iTBS | Chga | chromogranin A | 12652 | 0.124058929 | 0.379 | 0.265 | 0.012718576 | 8.61E-07 |
| Caudoputamen | iTBS | Rps14 | ribosomal protein S14 | 20044 | 0.12412127 | 0.981 | 0.975 | 0.011813394 | 7.99E-07 |
| Caudoputamen | iTBS | Chchd10 | coiled-coil-helix-coiled-coil-helix domain containing 10 | 103172 | 0.124584542 | 1 | 0.999 | 0.000185462 | 1.25E-08 |
| Caudoputamen | iTBS | Mrpl18 | mitochondrial ribosomal protein L18 | 67681 | 0.124748832 | 0.456 | 0.341 | 0.012569862 | 8.51E-07 |
| Caudoputamen | iTBS | Arpp21 | cyclic AMP-regulated phosphoprotein, 21 | 74100 | 0.125165285 | 0.988 | 0.97 | 0.010706689 | 7.25E-07 |
| Caudoputamen | iTBS | Cox4i1 | cytochrome c oxidase subunit 4I1 | 12857 | 0.125278231 | 1 | 1 | 5.94E-06 | 4.02E-10 |
| Caudoputamen | iTBS | Elk1 | ELK1, member of ETS oncogene family | 13712 | 0.12666731 | 0.303 | 0.193 | 0.000343117 | 2.32E-08 |
| Caudoputamen | iTBS | Spg21 | SPG21, maspardin | 27965 | 0.127338175 | 0.319 | 0.223 | 0.022159626 | 1.50E-06 |
| Caudoputamen | iTBS | Gpr158 | G protein-coupled receptor 158 | 241263 | 0.127686522 | 0.579 | 0.452 | 0.009110795 | 6.17E-07 |
| Caudoputamen | iTBS | Nop10 | NOP10 ribonucleoprotein | 66181 | 0.127694849 | 0.779 | 0.664 | 0.040442414 | 2.74E-06 |
| Caudoputamen | iTBS | Krt10 | keratin 10 | 16661 | 0.128129748 | 0.445 | 0.335 | 0.019156045 | 1.30E-06 |
| Caudoputamen | iTBS | Hagh | hydroxyacyl glutathione hydrolase | 14651 | 0.129555228 | 0.518 | 0.395 | 0.005670745 | 3.84E-07 |
| Caudoputamen | iTBS | Rpl29 | ribosomal protein L29 | 19944 | 0.129649113 | 0.986 | 0.983 | 0.000316408 | 2.14E-08 |
| Caudoputamen | iTBS | Trappc4 | trafficking protein particle complex 4 | 60409 | 0.130703692 | 0.41 | 0.289 | 0.001130745 | 7.65E-08 |
| Caudoputamen | iTBS | Krt12 | keratin 12 | 268482 | 0.131013165 | 0.181 | 0.087 | 9.65E-06 | 6.53E-10 |
| Caudoputamen | iTBS | Stub1 | STIP1 homology and U-Box containing protein 1 | 56424 | 0.131197451 | 0.625 | 0.479 | 0.02576908 | 1.74E-06 |
| Caudoputamen | iTBS | Tpm2 | tropomyosin 2, beta | 22004 | 0.131261686 | 0.221 | 0.122 | 4.89E-05 | 3.31E-09 |
| Caudoputamen | iTBS | Gpx4 | glutathione peroxidase 4 | 625249 | 0.132844921 | 0.993 | 0.996 | 0.000439738 | 2.98E-08 |
| Caudoputamen | iTBS | Rps20 | ribosomal protein S20 | 67427 | 0.133178848 | 0.998 | 0.997 | 2.33E-06 | 1.58E-10 |
| Caudoputamen | iTBS | Mrps16 | mitochondrial ribosomal protein S16 | 66242 | 0.133403333 | 0.438 | 0.301 | 7.80E-05 | 5.28E-09 |
| Caudoputamen | iTBS | Cd4 | CD4 antigen | 12504 | 0.134130177 | 0.383 | 0.281 | 0.008603104 | 5.82E-07 |
| Caudoputamen | iTBS | Gnb2 | guanine nucleotide binding protein (G protein), beta 2 | 14693 | 0.134546603 | 0.972 | 0.959 | 0.001638347 | 1.11E-07 |
| Caudoputamen | iTBS | Rps2 | ribosomal protein S2 | 16898 | 0.134972777 | 0.993 | 0.997 | 0.00016313 | 1.10E-08 |
| Caudoputamen | iTBS | Calm2 | calmodulin 2 | 12314 | 0.13525223 | 1 | 1 | 4.34E-11 | 2.94E-15 |
| Caudoputamen | iTBS | Uqcrb | ubiquinol-cytochrome c reductase binding protein | 67530 | 0.135557184 | 0.963 | 0.95 | 0.000420668 | 2.85E-08 |
| Caudoputamen | iTBS | Ndfip2 | Nedd4 family interacting protein 2 | 76273 | 0.135707297 | 0.51 | 0.376 | 0.000989939 | 6.70E-08 |
| Caudoputamen | iTBS | Elmod1 | ELMO/CED-12 domain containing 1 | 270162 | 0.13598525 | 0.923 | 0.845 | 0.009992017 | 6.76E-07 |
| Caudoputamen | iTBS | Guk1 | guanylate kinase 1 | 14923 | 0.136233495 | 0.662 | 0.517 | 0.006312181 | 4.27E-07 |
| Caudoputamen | iTBS | Tmbim4 | transmembrane BAX inhibitor motif containing 4 | 68212 | 0.136245958 | 0.337 | 0.211 | 2.78E-05 | 1.88E-09 |
| Caudoputamen | iTBS | Erbin | Erbb2 interacting protein | 59079 | 0.136341747 | 0.254 | 0.152 | 0.000148633 | 1.01E-08 |
| Caudoputamen | iTBS | Dtnbp1 | dystrobrevin binding protein 1 | 94245 | 0.136515096 | 0.477 | 0.35 | 0.004468356 | 3.02E-07 |
| Caudoputamen | iTBS | 2410006H16Rik | RIKEN cDNA 2410006H16 gene | 69221 | 0.137231477 | 0.55 | 0.407 | 0.000457138 | 3.09E-08 |
| Caudoputamen | iTBS | Rasl11b | RAS-like, family 11, member B | 68939 | 0.137578343 | 0.211 | 0.11 | 1.13E-05 | 7.67E-10 |
| Caudoputamen | iTBS | Rpl11 | ribosomal protein L11 | 67025 | 0.137663562 | 0.999 | 0.999 | 1.63E-05 | 1.10E-09 |
| Caudoputamen | iTBS | Hist1h1c | NA | NA | 0.137854384 | 0.349 | 0.244 | 0.004865876 | 3.29E-07 |
| Caudoputamen | iTBS | Sdhaf1 | succinate dehydrogenase complex assembly factor 1 | 68332 | 0.137997131 | 0.295 | 0.184 | 0.000146673 | 9.93E-09 |
| Caudoputamen | iTBS | H2afz | NA | NA | 0.138024265 | 0.965 | 0.944 | 0.002928391 | 1.98E-07 |
| Caudoputamen | iTBS | H2afj | NA | NA | 0.13831767 | 0.681 | 0.532 | 0.002303265 | 1.56E-07 |
| Caudoputamen | iTBS | Ndufa7 | NADH:ubiquinone oxidoreductase subunit A7 | 66416 | 0.139650783 | 0.906 | 0.86 | 0.012651277 | 8.56E-07 |
| Caudoputamen | iTBS | Mrpl57 | mitochondrial ribosomal protein L57 | 67840 | 0.140309686 | 0.441 | 0.323 | 0.006038727 | 4.09E-07 |
| Caudoputamen | iTBS | Polr2k | polymerase (RNA) II (DNA directed) polypeptide K | 17749 | 0.140890946 | 0.593 | 0.453 | 0.002529818 | 1.71E-07 |
| Caudoputamen | iTBS | Rplp2 | ribosomal protein, large P2 | 67186 | 0.14114778 | 0.99 | 0.984 | 0.000881642 | 5.97E-08 |

| | | | | | | | | | |
|---|---|---|---|---|---|---|---|---|---|
| Caudoputamen | iTBS | Neto2 | neuropilin (NRP) and tolloid (TLL)-like 2 | 74513 | 0.142864537 | 0.456 | 0.333 | 0.000752087 | 5.09E-08 |
| Caudoputamen | iTBS | Rps29 | ribosomal protein S29 | 20090 | 0.142919956 | 0.999 | 0.998 | 0.034581606 | 2.34E-06 |
| Caudoputamen | iTBS | mt-Cytb | NA | NA | 0.143204314 | 1 | 1 | 4.58E-17 | 3.10E-21 |
| Caudoputamen | iTBS | Dpm3 | dolichyl-phosphate mannosyltransferase polypeptide 3 | 68563 | 0.145135466 | 0.456 | 0.316 | 4.00E-05 | 2.70E-09 |
| Caudoputamen | iTBS | Tmsb4x | thymosin, beta 4, X chromosome | 19241 | 0.14532626 | 1 | 1 | 2.36E-11 | 1.60E-15 |
| Caudoputamen | iTBS | Cacna2d3 | calcium channel, voltage-dependent, alpha2/delta subunit 3 | 12294 | 0.145513371 | 0.596 | 0.457 | 0.000446389 | 3.02E-08 |
| Caudoputamen | iTBS | Mrpl17 | mitochondrial ribosomal protein L17 | 27397 | 0.14685864 | 0.377 | 0.251 | 4.00E-05 | 2.71E-09 |
| Caudoputamen | iTBS | 2310009A05Rik | NA | NA | 0.147912125 | 0.353 | 0.232 | 5.48E-05 | 3.71E-09 |
| Caudoputamen | iTBS | Sbsn | suprabasin | 282619 | 0.148124618 | 0.323 | 0.209 | 7.72E-05 | 5.23E-09 |
| Caudoputamen | iTBS | Cystm1 | cysteine-rich transmembrane module containing 1 | 66060 | 0.148875259 | 0.733 | 0.612 | 0.012622757 | 8.54E-07 |
| Caudoputamen | iTBS | Cox8a | cytochrome c oxidase subunit 8A | 12868 | 0.148932369 | 1 | 1 | 7.90E-10 | 5.35E-14 |
| Caudoputamen | iTBS | Pcp4 | Purkinje cell protein 4 | 18546 | 0.149493529 | 1 | 1 | 2.94E-06 | 1.99E-10 |
| Caudoputamen | iTBS | Fau | Finkel-Biskis-Reilly murine sarcoma virus (FBR-MuSV) ubiquitously expres | 14109 | 0.1508702 | 1 | 1 | 3.77E-08 | 2.55E-12 |
| Caudoputamen | iTBS | Sem1 | SEM1, 26S proteasome complex subunit | 20422 | 0.151008339 | 0.798 | 0.667 | 0.001803604 | 1.22E-07 |
| Caudoputamen | iTBS | Alkbh7 | alkB homolog 7 | 66400 | 0.151155099 | 0.341 | 0.21 | 2.47E-06 | 1.67E-10 |
| Caudoputamen | iTBS | Rpl38 | ribosomal protein L38 | 67671 | 0.151503819 | 1 | 1 | 2.62E-09 | 1.77E-13 |
| Caudoputamen | iTBS | Tesc | tescalcin | 57816 | 0.15184919 | 0.921 | 0.847 | 0.000147514 | 9.98E-09 |
| Caudoputamen | iTBS | 1500011B03Rik | RIKEN cDNA 1500011B03 gene | 66236 | 0.152142612 | 0.693 | 0.543 | 0.000838281 | 5.67E-08 |
| Caudoputamen | iTBS | Rpl23 | ribosomal protein L23 | 65019 | 0.153481126 | 0.997 | 0.996 | 2.91E-07 | 1.97E-11 |
| Caudoputamen | iTBS | Selenom | selenoprotein M | 114679 | 0.153689128 | 0.877 | 0.822 | 0.001342538 | 9.08E-08 |
| Caudoputamen | iTBS | Ubb | ubiquitin B | 22187 | 0.154058705 | 1 | 1 | 5.82E-05 | 3.94E-09 |
| Caudoputamen | iTBS | Ppp1r1a | protein phosphatase 1, regulatory inhibitor subunit 1A | 58200 | 0.155182596 | 0.901 | 0.797 | 0.004002154 | 2.71E-07 |
| Caudoputamen | iTBS | Bloc1s1 | biogenesis of lysosomal organelles complex-1, subunit 1 | 14533 | 0.155236462 | 0.358 | 0.227 | 5.45E-06 | 3.69E-10 |
| Caudoputamen | iTBS | Spock3 | sparc/osteonectin, cwcv and kazal-like domains proteoglycan 3 | 72902 | 0.155236462 | 0.909 | 0.812 | 0.00029273 | 1.98E-08 |
| Caudoputamen | iTBS | Mrpl53 | mitochondrial ribosomal protein L53 | 68499 | 0.156315883 | 0.596 | 0.44 | 2.24E-05 | 1.52E-09 |
| Caudoputamen | iTBS | Uqcr11 | ubiquinol-cytochrome c reductase, complex III subunit XI | 66594 | 0.157387129 | 1 | 1 | 9.18E-08 | 6.21E-12 |
| Caudoputamen | iTBS | Fth1 | ferritin heavy polypeptide 1 | 14319 | 0.157583588 | 1 | 1 | 1.42E-05 | 9.62E-10 |
| Caudoputamen | iTBS | Them6 | thioesterase superfamily member 6 | 223626 | 0.157955336 | 0.49 | 0.357 | 0.000171956 | 1.16E-08 |
| Caudoputamen | iTBS | Rps7 | ribosomal protein S7 | 20115 | 0.160084489 | 0.998 | 0.996 | 1.55E-08 | 1.05E-12 |
| Caudoputamen | iTBS | Sep-05 | NA | NA | 0.160434841 | 0.232 | 0.118 | 2.23E-07 | 1.51E-11 |
| Caudoputamen | iTBS | Rps12 | ribosomal protein S12 | 20042 | 0.160542342 | 0.993 | 0.985 | 7.09E-05 | 4.80E-09 |
| Caudoputamen | iTBS | Ndufb8 | NADH:ubiquinone oxidoreductase subunit B8 | 67264 | 0.160616367 | 0.965 | 0.948 | 1.30E-05 | 8.82E-10 |
| Caudoputamen | iTBS | Ppp2cb | protein phosphatase 2 (formerly 2A), catalytic subunit, beta isoform | 19053 | 0.161641854 | 0.181 | 0.06 | 1.01E-12 | 6.80E-17 |
| Caudoputamen | iTBS | Rpl19 | ribosomal protein L19 | 19921 | 0.163171573 | 0.997 | 0.995 | 1.69E-06 | 1.14E-10 |
| Caudoputamen | iTBS | Gpr155 | G protein-coupled receptor 155 | 68526 | 0.165938997 | 0.467 | 0.362 | 0.006532214 | 4.42E-07 |
| Caudoputamen | iTBS | Rpl36a | ribosomal protein L36A | 19982 | 0.169440959 | 0.938 | 0.906 | 3.50E-05 | 2.37E-09 |
| Caudoputamen | iTBS | Drd1 | dopamine receptor D1 | 13488 | 0.17050542 | 0.71 | 0.562 | 1.48E-05 | 9.99E-10 |
| Caudoputamen | iTBS | Rps10 | ribosomal protein S10 | 67097 | 0.171482183 | 0.99 | 0.99 | 6.54E-07 | 4.43E-11 |
| Caudoputamen | iTBS | Ano3 | anoctamin 3 | 228432 | 0.174217815 | 0.656 | 0.54 | 0.002853381 | 1.93E-07 |
| Caudoputamen | iTBS | Tmsb10 | thymosin, beta 10 | 19240 | 0.175295875 | 1 | 0.999 | 1.66E-07 | 1.12E-11 |
| Caudoputamen | iTBS | Scn4b | sodium channel, type IV, beta | 399548 | 0.175434314 | 0.856 | 0.762 | 0.007682914 | 5.20E-07 |
| Caudoputamen | iTBS | Rps27a | ribosomal protein S27A | 78294 | 0.175999941 | 1 | 1 | 6.87E-14 | 4.65E-18 |
| Caudoputamen | iTBS | Rps16 | ribosomal protein S16 | 20055 | 0.176155078 | 0.99 | 0.983 | 2.03E-07 | 1.38E-11 |
| Caudoputamen | iTBS | Cox6c | cytochrome c oxidase subunit 6C | 12864 | 0.176316061 | 1 | 1 | 8.10E-21 | 5.48E-25 |
| Caudoputamen | iTBS | Rasgef1b | RasGEF domain family, member 1B | 320292 | 0.176782876 | 0.439 | 0.31 | 1.36E-05 | 9.20E-10 |
| Caudoputamen | iTBS | St8sia3 | ST8 alpha-N-acetyl-neuraminide alpha-2,8-sialyltransferase 3 | 20451 | 0.177081673 | 0.863 | 0.765 | 0.000957541 | 6.48E-08 |
| Caudoputamen | iTBS | Rpl34 | ribosomal protein L34 | 68436 | 0.18012321 | 1 | 0.994 | 9.18E-08 | 6.21E-12 |
| Caudoputamen | iTBS | S100a10 | S100 calcium binding protein A10 (calpactin) | 20194 | 0.180961075 | 0.291 | 0.163 | 2.01E-07 | 1.36E-11 |
| Caudoputamen | iTBS | Rpl13 | ribosomal protein L13 | 270106 | 0.189188214 | 1 | 1 | 1.08E-15 | 7.32E-20 |
| Caudoputamen | iTBS | Rpl37 | ribosomal protein L37 | 67281 | 0.190933163 | 1 | 1 | 5.07E-16 | 3.43E-20 |
| Caudoputamen | iTBS | Sap18 | Sin3-associated polypeptide 18 | 20220 | 0.190968995 | 0.449 | 0.275 | 1.54E-10 | 1.04E-14 |
| Caudoputamen | iTBS | Sf3b5 | splicing factor 3b, subunit 5 | 66125 | 0.191377442 | 0.597 | 0.431 | 4.58E-06 | 3.10E-10 |
| Caudoputamen | iTBS | mt-Nd5 | NA | NA | 0.194157718 | 0.994 | 0.996 | 7.97E-08 | 5.39E-12 |
| Caudoputamen | iTBS | Ndufa1 | NADH:ubiquinone oxidoreductase subunit A1 | 54405 | 0.199500467 | 0.918 | 0.843 | 1.11E-07 | 7.54E-12 |
| Caudoputamen | iTBS | mt-Atp6 | NA | NA | 0.19985248 | 1 | 1 | 1.55E-24 | 1.05E-28 |
| Caudoputamen | iTBS | Malat1 | metastasis associated lung adenocarcinoma transcript 1 (non-coding RNA | 72289 | 0.202736245 | 0.773 | 0.645 | 0.000890041 | 6.02E-08 |
| Caudoputamen | iTBS | Rps24 | ribosomal protein S24 | 20088 | 0.210576663 | 1 | 1 | 1.02E-18 | 6.92E-23 |
| Caudoputamen | iTBS | Erh | ERH mRNA splicing and mitosis factor | 13877 | 0.215196624 | 0.452 | 0.278 | 8.69E-11 | 5.88E-15 |
| Caudoputamen | iTBS | Rpl30 | ribosomal protein L30 | 19946 | 0.216625612 | 1 | 0.999 | 6.69E-15 | 4.53E-19 |
| Caudoputamen | iTBS | Gpr88 | G-protein coupled receptor 88 | 64378 | 0.219196673 | 0.992 | 0.981 | 6.65E-14 | 4.50E-18 |
| Caudoputamen | iTBS | Rpl9 | ribosomal protein L9 | 20005 | 0.225953243 | 0.999 | 0.998 | 8.70E-17 | 5.89E-21 |
| Caudoputamen | iTBS | Cox7c | cytochrome c oxidase subunit 7C | 12867 | 0.228564363 | 1 | 1 | 1.24E-23 | 8.42E-28 |
| Caudoputamen | iTBS | mt-Nd4 | NA | NA | 0.228672926 | 1 | 1 | 9.59E-16 | 6.49E-20 |
| Caudoputamen | iTBS | Rps8 | ribosomal protein S8 | 20116 | 0.230594434 | 0.996 | 1 | 1.52E-19 | 1.03E-23 |
| Caudoputamen | iTBS | Ppp3ca | protein phosphatase 3, catalytic subunit, alpha isoform | 19055 | 0.233753474 | 1 | 0.999 | 1.39E-16 | 9.44E-21 |
| Caudoputamen | iTBS | mt-Nd1 | NA | NA | 0.234517462 | 1 | 1 | 8.02E-19 | 5.43E-23 |
| Caudoputamen | iTBS | Rps26 | ribosomal protein S26 | 27370 | 0.235225817 | 0.992 | 0.978 | 3.77E-14 | 2.55E-18 |
| Caudoputamen | iTBS | Rps28 | ribosomal protein S28 | 54127 | 0.2382301 | 0.95 | 0.935 | 2.04E-08 | 1.38E-12 |
| Caudoputamen | iTBS | Hmgn2 | high mobility group nucleosomal binding domain 2 | 15331 | 0.24812134 | 0.624 | 0.443 | 4.79E-11 | 3.24E-15 |
| Caudoputamen | iTBS | Rpl35a | ribosomal protein L35A | 57808 | 0.252392802 | 1 | 0.996 | 6.67E-24 | 4.51E-28 |
| Caudoputamen | iTBS | mt-Co3 | NA | NA | 0.260275142 | 1 | 1 | 9.34E-48 | 6.32E-52 |
| Caudoputamen | iTBS | Ppp1r1b | protein phosphatase 1, regulatory inhibitor subunit 1B | 19049 | 0.261919919 | 0.997 | 0.991 | 2.19E-21 | 1.48E-25 |
| Caudoputamen | iTBS | Rps21 | ribosomal protein S21 | 66481 | 0.283112319 | 1 | 1 | 4.45E-41 | 3.01E-45 |
| Caudoputamen | iTBS | mt-Co1 | NA | NA | 0.289256147 | 1 | 1 | 8.45E-52 | 5.72E-56 |
| Caudoputamen | iTBS | Cnr1 | cannabinoid receptor 1 (brain) | 12801 | 0.292881538 | 0.681 | 0.532 | 3.39E-09 | 2.30E-13 |
| Caudoputamen | iTBS | Adora2a | adenosine A2a receptor | 11540 | 0.300944336 | 0.78 | 0.614 | 1.49E-13 | 1.01E-17 |
| Caudoputamen | iTBS | Rpl35 | ribosomal protein L35 | 66489 | 0.36964948 | 0.985 | 0.965 | 1.36E-31 | 9.19E-36 |
| Caudoputamen | iTBS | mt-Co2 | NA | NA | 0.389695174 | 1 | 1 | 9.42E-38 | 6.37E-42 |
| Caudoputamen | iTBS | Ttr | transthyretin | 22139 | 0.615804318 | 0.258 | 0.141 | 4.31E-10 | 2.92E-14 |
| Caudoputamen | iTBS | Gstp1 | glutathione S-transferase, pi 1 | 14870 | 0.632915936 | 0.612 | 0.27 | 5.38E-56 | 3.64E-60 |
| Caudoputamen | iTBS | Uba52 | ubiquitin A-52 residue ribosomal protein fusion product 1 | 22186 | 0.904690012 | 0.883 | 0.78 | 4.63E-81 | 3.13E-85 |
| Caudoputamen | iTBS | Gm10076 | ribosomal protein L41 pseudogene | 100126819 | 1.504442045 | 0.811 | 0.463 | 8.87E-131 | 6.00E-135 |
| Lateral Septal Complex | iTBS | Ttr | transthyretin | 22139 | -1.617776097 | 0.174 | 0.321 | 0.023932756 | 1.62E-06 |
| Lateral Septal Complex | iTBS | Prkca | protein kinase C, alpha | 18750 | -0.338440643 | 0.264 | 0.426 | 0.017054536 | 1.15E-06 |
| Lateral Septal Complex | iTBS | Phyhip | phytanoyl-CoA hydroxylase interacting protein | 105653 | -0.317131003 | 0.289 | 0.483 | 0.000450812 | 3.05E-08 |
| Lateral Septal Complex | iTBS | Cdk5r2 | cyclin-dependent kinase 5, regulatory subunit 2 (p39) | 12570 | -0.29895976 | 0.232 | 0.441 | 5.13E-05 | 3.47E-09 |
| Lateral Septal Complex | iTBS | Luzp2 | leucine zipper protein 2 | 233271 | -0.287605523 | 0.402 | 0.561 | 0.016368979 | 1.11E-06 |
| Lateral Septal Complex | iTBS | Kcnq2 | potassium voltage-gated channel, subfamily Q, member 2 | 16536 | -0.276817724 | 0.46 | 0.621 | 0.044984929 | 3.04E-06 |
| Lateral Septal Complex | iTBS | Sgk1 | serum/glucocorticoid regulated kinase 1 | 20393 | -0.276265073 | 0.113 | 0.287 | 0.000143616 | 9.72E-09 |
| Lateral Septal Complex | iTBS | Parp6 | poly (ADP-ribose) polymerase family, member 6 | 67287 | -0.256255699 | 0.232 | 0.41 | 0.005951657 | 4.03E-07 |
| Lateral Septal Complex | iTBS | Hectd4 | HECT domain E3 ubiquitin protein ligase 4 | 269700 | -0.253133208 | 0.334 | 0.514 | 0.019764442 | 1.34E-06 |
| Lateral Septal Complex | iTBS | Kalrn | kalirin, RhoGEF kinase | 545156 | -0.252111211 | 0.27 | 0.449 | 0.016802031 | 1.14E-06 |
| Lateral Septal Complex | iTBS | Ppp1cb | protein phosphatase 1 catalytic subunit beta | 19046 | -0.241326741 | 0.482 | 0.658 | 0.033368969 | 2.26E-06 |
| Lateral Septal Complex | iTBS | Tuba1a | tubulin, alpha 1A | 22142 | -0.22386303 | 0.984 | 0.987 | 0.006064108 | 4.10E-07 |
| Lateral Septal Complex | iTBS | Cpe | carboxypeptidase E | 12876 | -0.20052739 | 1 | 1 | 0.000114487 | 7.75E-09 |
| Lateral Septal Complex | iTBS | Kcnip3 | Kv channel interacting protein 3, calsenilin | 56461 | -0.191869732 | 0.132 | 0.29 | 0.011281535 | 7.63E-07 |
| Lateral Septal Complex | iTBS | Fau | Finkel-Biskis-Reilly murine sarcoma virus (FBR-MuSV) ubiquitously expres | 14109 | 0.159567276 | 1 | 1 | 0.008123359 | 5.50E-07 |
| Lateral Septal Complex | iTBS | Rpl9 | ribosomal protein L9 | 20005 | 0.170399839 | 1 | 1 | 0.02034592 | 1.38E-06 |
| Lateral Septal Complex | iTBS | Atp5md | ATP synthase membrane subunit DAPIT | 66477 | 0.176142097 | 1 | 1 | 0.043148473 | 2.92E-06 |
| Lateral Septal Complex | iTBS | Rps20 | ribosomal protein S20 | 67427 | 0.17890355 | 1 | 0.997 | 0.003517822 | 2.38E-07 |
| Lateral Septal Complex | iTBS | mt-Nd1 | NA | NA | 0.202311763 | 1 | 1 | 0.002862704 | 1.94E-07 |
| Lateral Septal Complex | iTBS | Rps21 | ribosomal protein S21 | 66481 | 0.216266446 | 1 | 1 | 1.80E-07 | 1.22E-11 |
| Lateral Septal Complex | iTBS | Rpl38 | ribosomal protein L38 | 67671 | 0.228828493 | 1 | 1 | 1.06E-05 | 7.14E-10 |
| Lateral Septal Complex | iTBS | Rps29 | ribosomal protein S29 | 20090 | 0.310903719 | 1 | 1 | 0.00011808 | 7.99E-09 |
| Lateral Septal Complex | iTBS | Rps28 | ribosomal protein S28 | 54127 | 0.333394575 | 0.977 | 0.958 | 5.20E-05 | 3.52E-09 |
| Lateral Septal Complex | iTBS | Rpl35 | ribosomal protein L35 | 66489 | 0.444890654 | 0.997 | 0.948 | 3.81E-13 | 2.58E-17 |
| Lateral Septal Complex | iTBS | Uba52 | ubiquitin A-52 residue ribosomal protein fusion product 1 | 22186 | 0.752594811 | 0.833 | 0.705 | 7.13E-16 | 4.82E-20 |
| Lateral Septal Complex | iTBS | Gm10076 | ribosomal protein L41 pseudogene | 100126819 | 1.23638845 | 0.646 | 0.496 | 2.39E-15 | 1.62E-19 |
| Striatum Ventral Region | iTBS | Bc1 | brain cytoplasmic RNA 1 | 100568459 | -0.384592556 | 0.676 | 0.83 | 6.83E-07 | 4.62E-11 |
| Striatum Ventral Region | iTBS | Atp1a3 | ATPase, Na+/K+ transporting, alpha 3 polypeptide | 232975 | -0.266323597 | 0.735 | 0.818 | 6.50E-06 | 4.40E-10 |
| Striatum Ventral Region | iTBS | Actb | actin, beta | 11461 | -0.256477526 | 0.999 | 0.998 | 1.64E-07 | 1.11E-11 |
| Striatum Ventral Region | iTBS | Syn1 | synapsin I | 20964 | -0.239973357 | 0.615 | 0.712 | 0.022842128 | 1.55E-06 |
| Striatum Ventral Region | iTBS | Meg3 | maternally expressed 3 | 17263 | -0.229628851 | 0.986 | 0.992 | 0.002868654 | 1.94E-07 |
| Striatum Ventral Region | iTBS | Dnm1 | dynamin 1 | 13429 | -0.22820985 | 0.878 | 0.931 | 0.013451967 | 9.10E-07 |
| Striatum Ventral Region | iTBS | Glul | glutamate-ammonia ligase (glutamine synthetase) | 14645 | -0.220806292 | 0.745 | 0.819 | 0.008322698 | 5.63E-07 |
| Striatum Ventral Region | iTBS | Dynll2 | dynein light chain LC8-type 2 | 68097 | -0.216258769 | 0.915 | 0.949 | 0.006182429 | 4.18E-07 |
| Striatum Ventral Region | iTBS | Gnao1 | guanine nucleotide binding protein, alpha O | 14681 | -0.197577158 | 0.752 | 0.801 | 0.028294206 | 1.91E-06 |
| Striatum Ventral Region | iTBS | Eef2 | eukaryotic translation elongation factor 2 | 13629 | -0.195034193 | 0.953 | 0.97 | 0.033525655 | 2.27E-06 |
| Striatum Ventral Region | iTBS | Hspa8 | heat shock protein 8 | 15481 | -0.132345271 | 1 | 1 | 0.007522291 | 5.09E-07 |
| Striatum Ventral Region | iTBS | Eef1a1 | eukaryotic translation elongation factor 1 alpha 1 | 13627 | 0.095476485 | 1 | 1 | 0.014086426 | 9.53E-07 |
| Striatum Ventral Region | iTBS | Coro1b | coronin, actin binding protein 1B | 23789 | 0.102270482 | 0.121 | 0.044 | 0.005963946 | 4.04E-07 |
| Striatum Ventral Region | iTBS | Amd1 | S-adenosylmethionine decarboxylase 1 | 11702 | 0.113504408 | 0.168 | 0.075 | 0.005113521 | 3.46E-07 |
| Striatum Ventral Region | iTBS | Commd10 | COMM domain containing 10 | 69456 | 0.116287955 | 0.187 | 0.095 | 0.029554245 | 2.00E-06 |
| Striatum Ventral Region | iTBS | Rpl24 | ribosomal protein L24 | 68193 | 0.123032889 | 0.997 | 0.994 | 0.02754093 | 1.86E-06 |
| Striatum Ventral Region | iTBS | Rplp1 | ribosomal protein, large, P1 | 56040 | 0.124914551 | 1 | 1 | 0.003539279 | 2.39E-07 |
| Striatum Ventral Region | iTBS | mt-Cytb | NA | NA | 0.127201826 | 1 | 1 | 6.00E-05 | 4.06E-09 |
| Striatum Ventral Region | iTBS | Atp5md | ATP synthase membrane subunit DAPIT | 66477 | 0.135900316 | 1 | 1 | 0.047140952 | 3.19E-06 |
| Striatum Ventral Region | iTBS | 2310009A05Rik | NA | NA | 0.136915374 | 0.32 | 0.201 | 0.026764124 | 1.81E-06 |
| Striatum Ventral Region | iTBS | Rpl35a | ribosomal protein L35A | 57808 | 0.140828462 | 1 | 1 | 0.001125769 | 7.62E-08 |
| Striatum Ventral Region | iTBS | mt-Nd1 | NA | NA | 0.140855782 | 1 | 1 | 0.006589403 | 4.46E-07 |
| Striatum Ventral Region | iTBS | Rps19bp1 | ribosomal protein S19 binding protein 1 | 66538 | 0.143582824 | 0.363 | 0.238 | 0.041556109 | 2.81E-06 |
| Striatum Ventral Region | iTBS | Rps20 | ribosomal protein S20 | 67427 | 0.144449047 | 1 | 0.998 | 0.000390694 | 2.64E-08 |
| Striatum Ventral Region | iTBS | Rpl9 | ribosomal protein L9 | 20005 | 0.145143374 | 1 | 0.998 | 0.00075809 | 5.13E-08 |
| Striatum Ventral Region | iTBS | Bloc1s1 | biogenesis of lysosomal organelles complex-1, subunit 1 | 14533 | 0.145599063 | 0.369 | 0.237 | 0.007842228 | 5.31E-07 |
| Striatum Ventral Region | iTBS | Pdap1 | PDGFA associated protein 1 | 231887 | 0.146322156 | 0.439 | 0.31 | 0.039830082 | 2.70E-06 |
| Striatum Ventral Region | iTBS | Cox6c | cytochrome c oxidase subunit 6C | 12864 | 0.150062654 | 1 | 1 | 1.79E-05 | 1.21E-09 |
| Striatum Ventral Region | iTBS | mt-Nd4 | NA | NA | 0.150259595 | 1 | 1 | 0.003413319 | 2.31E-07 |
| Striatum Ventral Region | iTBS | Grcc10 | gene rich cluster, C10 gene | 14790 | 0.15541791 | 0.951 | 0.914 | 0.048171215 | 3.26E-06 |
| Striatum Ventral Region | iTBS | Tmsb4x | thymosin, beta 4, X chromosome | 19241 | 0.159733412 | 1 | 1 | 4.10E-06 | 2.77E-10 |
| Striatum Ventral Region | iTBS | Rps23 | ribosomal protein S23 | 66475 | 0.160366144 | 0.994 | 0.986 | 0.009850205 | 6.67E-07 |
| Striatum Ventral Region | iTBS | Cebpzos | CCAAT/enhancer binding protein (C/EBP), zeta, opposite strand | 68554 | 0.16133521 | 0.462 | 0.329 | 0.013544473 | 9.17E-07 |
| Striatum Ventral Region | iTBS | Chrac1 | chromatin accessibility complex 1 | 93696 | 0.161847675 | 0.326 | 0.192 | 0.000635848 | 4.30E-08 |
| Striatum Ventral Region | iTBS | Rpl34 | ribosomal protein L34 | 68436 | 0.163652 | 0.996 | 0.998 | 0.000220641 | 1.49E-08 |
| Striatum Ventral Region | iTBS | Sec61b | Sec61 beta subunit | 66212 | 0.166965415 | 0.532 | 0.416 | 0.048172761 | 3.26E-06 |
| Striatum Ventral Region | iTBS | Snhg8 | small nucleolar RNA host gene 8 | 69895 | 0.180106823 | 0.571 | 0.431 | 0.005217216 | 3.53E-07 |
| Striatum Ventral Region | iTBS | Ppp2cb | protein phosphatase 2 (formerly 2A), catalytic subunit, beta isoform | 19053 | 0.183863728 | 0.197 | 0.064 | 1.36E-08 | 9.20E-13 |
| Striatum Ventral Region | iTBS | Lrpap1 | low density lipoprotein receptor-related protein associated protein 1 | 16976 | 0.185620024 | 0.749 | 0.615 | 0.022212214 | 1.50E-06 |
| Striatum Ventral Region | iTBS | Rpl38 | ribosomal protein L38 | 67671 | 0.189170933 | 1 | 1 | 5.27E-09 | 3.56E-13 |
| Striatum Ventral Region | iTBS | mt-Atp6 | NA | NA | 0.194787098 | 1 | 1 | 5.38E-12 | 3.64E-16 |
| Striatum Ventral Region | iTBS | Ndufb4 | NADH:ubiquinone oxidoreductase subunit B4 | 68194 | 0.195124326 | 0.954 | 0.914 | 0.002290551 | 1.55E-07 |
| Striatum Ventral Region | iTBS | Rps27 | ribosomal protein S27 | 57294 | 0.198069547 | 0.991 | 0.991 | 0.016344221 | 1.11E-06 |
| Striatum Ventral Region | iTBS | Cox7c | cytochrome c oxidase subunit 7C | 12867 | 0.200940927 | 1 | 0.998 | 5.13E-09 | 3.47E-13 |
| Striatum Ventral Region | iTBS | Ubb | ubiquitin B | 22187 | 0.201471016 | 1 | 1 | 3.82E-10 | 2.58E-14 |
| Striatum Ventral Region | iTBS | Rps21 | ribosomal protein S21 | 66481 | 0.2017355 | 1 | 1 | 2.38E-10 | 1.61E-14 |
| Striatum Ventral Region | iTBS | Ppia | peptidylprolyl isomerase A | 268373 | 0.203162049 | 1 | 1 | 1.86E-12 | 1.26E-16 |
| Striatum Ventral Region | iTBS | Tmsb10 | thymosin, beta 10 | 19240 | 0.206648963 | 1 | 1 | 1.02E-05 | 6.89E-10 |
| Striatum Ventral Region | iTBS | Eno1 | enolase 1, alpha non-neuron | 13806 | 0.214430604 | 0.892 | 0.85 | 0.043445689 | 2.94E-06 |
| Striatum Ventral Region | iTBS | Setp5 | NA | NA | 0.214773317 | 0.23 | 0.087 | 1.86E-08 | 1.26E-12 |
| Striatum Ventral Region | iTBS | Rps29 | ribosomal protein S29 | 20090 | 0.217502852 | 0.999 | 0.998 | 0.011498336 | 7.78E-07 |
| Striatum Ventral Region | iTBS | mt-Co3 | NA | NA | 0.218541483 | 1 | 1 | 2.29E-16 | 1.55E-20 |
| Striatum Ventral Region | iTBS | Ndufb1-ps | NA | NA | 0.220992761 | 0.897 | 0.829 | 0.018149062 | 1.23E-06 |
| Striatum Ventral Region | iTBS | mt-Co1 | NA | NA | 0.234626116 | 1 | 1 | 1.57E-16 | 1.07E-20 |
| Striatum Ventral Region | iTBS | mt-Co2 | NA | NA | 0.254479414 | 1 | 1 | 9.80E-09 | 6.63E-13 |
| Striatum Ventral Region | iTBS | Rps28 | ribosomal protein S28 | 54127 | 0.270465407 | 0.96 | 0.964 | 0.00011676 | 7.90E-09 |
| Striatum Ventral Region | iTBS | Rpl35 | ribosomal protein L35 | 66489 | 0.305537969 | 0.984 | 0.972 | 2.08E-15 | 1.40E-19 |
| Striatum Ventral Region | iTBS | Gstp1 | glutathione S-transferase, pi 1 | 14870 | 0.478605517 | 0.584 | 0.368 | 7.52E-17 | 5.09E-21 |
| Striatum Ventral Region | iTBS | Uba52 | ubiquitin A-52 residue ribosomal protein fusion product 1 | 22186 | 0.801778525 | 0.879 | 0.832 | 9.45E-38 | 6.39E-42 |
| Striatum Ventral Region | iTBS | Gm10076 | ribosomal protein L41 pseudogene | 100126819 | 1.338143498 | 0.733 | 0.59 | 7.19E-48 | 4.87E-52 |
| White Matter Tracts | cTBS | Bc1 | brain cytoplasmic RNA 1 | 100568459 | -0.834899137 | 0.713 | 0.885 | 1.48E-22 | 9.99E-27 |
| White Matter Tracts | cTBS | Nrsn2 | neurensin 2 | 228777 | -0.284072589 | 0.339 | 0.538 | 8.05E-08 | 5.45E-12 |
| White Matter Tracts | cTBS | Tmem130 | transmembrane protein 130 | 243339 | -0.277167142 | 0.253 | 0.44 | 5.40E-07 | 3.66E-11 |
| White Matter Tracts | cTBS | Resp18 | regulated endocrine-specific protein 18 | 19711 | -0.252238902 | 0.594 | 0.73 | 0.000117832 | 7.97E-09 |
| White Matter Tracts | cTBS | Mgrn1 | mahogunin, ring finger 1 | 17237 | -0.241863763 | 0.355 | 0.518 | 0.002295737 | 1.55E-07 |
| White Matter Tracts | cTBS | Ncald | neurocalcin delta | 52589 | -0.232309316 | 0.43 | 0.592 | 0.003656749 | 2.47E-07 |
| White Matter Tracts | cTBS | mt-Nd2 | NA | NA | -0.228247148 | 1 | 1 | 0.014044266 | 9.50E-07 |
| White Matter Tracts | cTBS | Mapk8ip3 | mitogen-activated protein kinase 8 interacting protein 3 | 30957 | -0.223116101 | 0.48 | 0.625 | 0.035610238 | 2.41E-06 |
| White Matter Tracts | cTBS | Fam234b | family with sequence similarity 234, member B | 74525 | -0.223088611 | 0.329 | 0.505 | 0.001068056 | 7.23E-08 |
| White Matter Tracts | cTBS | Ahi1 | Abelson helper integration site 1 | 52906 | -0.22077216 | 0.165 | 0.298 | 0.006223014 | 4.21E-07 |
| White Matter Tracts | cTBS | Atp6v1a | ATPase, H+ transporting, lysosomal V1 subunit A | 11964 | -0.217659317 | 0.289 | 0.453 | 0.000734314 | 4.97E-08 |
| White Matter Tracts | cTBS | Gad2 | glutamic acid decarboxylase 2 | 14417 | -0.215179393 | 0.434 | 0.613 | 4.74E-05 | 3.21E-09 |
| White Matter Tracts | cTBS | Rnf157 | ring finger protein 157 | 217340 | -0.212327836 | 0.271 | 0.417 | 0.013796978 | 9.34E-07 |
| White Matter Tracts | cTBS | Nnat | neuronatin | 18111 | -0.209886442 | 0.462 | 0.625 | 0.035573989 | 2.41E-06 |
| White Matter Tracts | cTBS | Ubqln2 | ubiquilin 2 | 54609 | -0.209403346 | 0.271 | 0.422 | 0.01200918 | 8.13E-07 |
| White Matter Tracts | cTBS | Podxl2 | podocalyxin-like 2 | 319655 | -0.209344813 | 0.373 | 0.542 | 0.000173366 | 1.17E-08 |
| White Matter Tracts | cTBS | Ank3 | ankyrin 3, epithelial | 11735 | -0.206809269 | 0.359 | 0.523 | 0.009327571 | 6.31E-07 |
| White Matter Tracts | cTBS | Sucla2 | succinate-Coenzyme A ligase, ADP-forming, beta subunit | 20916 | -0.205068078 | 0.257 | 0.408 | 0.007131674 | 4.83E-07 |
| White Matter Tracts | cTBS | Kcnq2 | potassium voltage-gated channel, subfamily Q, member 2 | 16536 | -0.200137314 | 0.474 | 0.615 | 0.033376997 | 2.26E-06 |
| White Matter Tracts | cTBS | Spock2 | sparc/osteonectin, cwcv and kazal-like domains proteoglycan 2 | 94214 | -0.198843348 | 0.394 | 0.55 | 0.011626411 | 7.87E-07 |
| White Matter Tracts | cTBS | Lingo1 | leucine rich repeat and Ig domain containing 1 | 235402 | -0.198233321 | 0.361 | 0.518 | 0.002792435 | 1.89E-07 |
| White Matter Tracts | cTBS | Agpat3 | 1-acylglycerol-3-phosphate O-acyltransferase 3 | 28169 | -0.197760431 | 0.303 | 0.463 | 0.025502357 | 1.73E-06 |
| White Matter Tracts | cTBS | Laptm4a | lysosomal-associated protein transmembrane 4A | 17775 | -0.195376655 | 0.335 | 0.497 | 0.017378952 | 1.18E-06 |
| White Matter Tracts | cTBS | Svop | SV2 related protein | 68666 | -0.193769291 | 0.203 | 0.362 | 0.000358251 | 2.42E-08 |
| White Matter Tracts | cTBS | Nrxn3 | neurexin III | 18191 | -0.193694903 | 0.303 | 0.453 | 0.007312266 | 4.95E-07 |
| White Matter Tracts | cTBS | Nell2 | NEL-like 2 | 54003 | -0.193211199 | 0.171 | 0.298 | 0.01749695 | 1.18E-06 |
| White Matter Tracts | cTBS | Ndn | necdin, MAGE family member | 17984 | -0.192004768 | 0.38 | 0.53 | 0.032937367 | 2.23E-06 |
| White Matter Tracts | cTBS | Rnf208 | ring finger protein 208 | 68846 | -0.190986229 | 0.329 | 0.483 | 0.013495216 | 9.13E-07 |
| White Matter Tracts | cTBS | Oaz2 | ornithine decarboxylase antizyme 2 | 18247 | -0.190739685 | 0.273 | 0.428 | 0.004861078 | 3.29E-07 |
| White Matter Tracts | cTBS | Cbarp | calcium channel, voltage-dependent, beta subunit associated regulatory p | 100503659 | -0.189355487 | 0.51 | 0.662 | 0.018092201 | 1.22E-06 |
| White Matter Tracts | cTBS | Setp3 | NA | NA | -0.188668404 | 0.422 | 0.583 | 0.005538835 | 3.75E-07 |
| White Matter Tracts | cTBS | Ap2a2 | adaptor-related protein complex 2, alpha 2 subunit | 11772 | -0.187327725 | 0.444 | 0.602 | 0.00836411 | 5.66E-07 |
| White Matter Tracts | cTBS | Lamp5 | lysosomal-associated membrane protein family, member 5 | 76161 | -0.186393368 | 0.339 | 0.495 | 0.00458906 | 3.11E-07 |
| White Matter Tracts | cTBS | Impact | impact, RWD domain protein | 16210 | -0.185629156 | 0.546 | 0.698 | 0.011978469 | 8.11E-07 |
| White Matter Tracts | cTBS | Pfdn1 | prefoldin 1 | 67199 | -0.185465292 | 0.315 | 0.477 | 0.00938698 | 6.35E-07 |
| White Matter Tracts | cTBS | Abr | active BCR-related gene | 109934 | -0.184752 | 0.637 | 0.757 | 0.040492761 | 2.74E-06 |
| White Matter Tracts | cTBS | Klc1 | kinesin light chain 1 | 16593 | -0.184045107 | 0.972 | 0.98 | 0.021866516 | 1.48E-06 |
| White Matter Tracts | cTBS | Grik5 | glutamate receptor, ionotropic, kainate 5 (gamma 2) | 14809 | -0.183476527 | 0.53 | 0.693 | 0.003171957 | 2.15E-07 |
| White Matter Tracts | cTBS | Myl12b | myosin, light chain 12B, regulatory | 67938 | -0.180891292 | 0.952 | 0.968 | 0.013786298 | 9.33E-07 |
| White Matter Tracts | cTBS | Tmx4 | thioredoxin-related transmembrane protein 4 | 52837 | -0.180164061 | 0.231 | 0.37 | 0.029434876 | 1.99E-06 |
| White Matter Tracts | cTBS | Cadm3 | cell adhesion molecule 3 | 94332 | -0.179181346 | 0.321 | 0.475 | 0.006748344 | 4.57E-07 |
| White Matter Tracts | cTBS | Arpc4 | actin related protein 2/3 complex, subunit 4 | 68089 | -0.17766192 | 0.261 | 0.417 | 0.005852999 | 3.96E-07 |
| White Matter Tracts | cTBS | Usp14 | ubiquitin specific peptidase 14 | 59025 | -0.176679576 | 0.223 | 0.358 | 0.043315217 | 2.93E-06 |
| White Matter Tracts | cTBS | Peg13 | paternally expressed 13 | 353342 | -0.176667368 | 0.5 | 0.657 | 0.009057787 | 6.13E-07 |
| White Matter Tracts | cTBS | Ncs1 | neuronal calcium sensor 1 | 14299 | -0.174647799 | 0.345 | 0.507 | 0.002456908 | 1.66E-07 |
| White Matter Tracts | cTBS | Ly6h | lymphocyte antigen 6 complex, locus H | 23934 | -0.174613564 | 0.92 | 0.947 | 0.009880292 | 6.69E-07 |
| White Matter Tracts | cTBS | Gnal | guanine nucleotide binding protein, alpha stimulating, olfactory type | 14680 | -0.173359218 | 0.43 | 0.583 | 0.022761389 | 1.54E-06 |
| White Matter Tracts | cTBS | Usp5 | ubiquitin specific peptidase 5 (isopeptidase T) | 22225 | -0.172837683 | 0.271 | 0.427 | 0.011191193 | 7.57E-07 |
| White Matter Tracts | cTBS | Scg2 | secretogranin II | 20254 | -0.171455842 | 0.375 | 0.53 | 0.017674928 | 1.20E-06 |
| White Matter Tracts | cTBS | Snrpn | small nuclear ribonucleoprotein N | 20646 | -0.169969268 | 0.972 | 0.995 | 0.020180695 | 1.37E-06 |
| White Matter Tracts | cTBS | Atp1b1 | ATPase, Na+/K+ transporting, beta 1 polypeptide | 11931 | -0.167378839 | 0.984 | 0.993 | 0.046752577 | 3.16E-06 |
| White Matter Tracts | cTBS | Stxbp1 | syntaxin binding protein 1 | 20910 | -0.167100227 | 0.886 | 0.925 | 0.01432875 | 9.70E-07 |
| White Matter Tracts | cTBS | Rasgrf1 | RAS protein-specific guanine nucleotide-releasing factor 1 | 19417 | -0.166499029 | 0.478 | 0.632 | 0.008057517 | 5.45E-07 |
| White Matter Tracts | cTBS | Snap25 | synaptosomal-associated protein 25 | 20614 | -0.166119584 | 0.984 | 0.997 | 0.013510926 | 9.14E-07 |
| White Matter Tracts | cTBS | Vsnl1 | visinin-like 1 | 26950 | -0.16403496 | 0.936 | 0.945 | 0.003732621 | 2.53E-07 |
| White Matter Tracts | cTBS | Arhgef9 | CDC42 guanine nucleotide exchange factor (GEF) 9 | 236915 | -0.161932004 | 0.301 | 0.458 | 0.008566716 | 5.80E-07 |
| White Matter Tracts | cTBS | Tceal6 | transcription elongation factor A (SII)-like 6 | 66104 | -0.161113572 | 0.229 | 0.373 | 0.013361967 | 9.04E-07 |
| White Matter Tracts | cTBS | Pvalb | parvalbumin | 19293 | -0.160314451 | 0.319 | 0.468 | 0.017849253 | 1.21E-06 |
| White Matter Tracts | cTBS | Zwint | ZW10 interactor | 52696 | -0.15720523 | 0.924 | 0.947 | 0.014738814 | 9.97E-07 |
| White Matter Tracts | cTBS | Zfr | zinc finger RNA binding protein | 22763 | -0.155913938 | 0.392 | 0.55 | 0.035759334 | 2.42E-06 |
| White Matter Tracts | cTBS | Chga | chromogranin A | 12652 | -0.155506389 | 0.203 | 0.337 | 0.025794313 | 1.75E-06 |
| White Matter Tracts | cTBS | Ywhag | tyrosine 3-monooxygenase/tryptophan 5-monooxygenase activation pro | 22628 | -0.154151109 | 0.936 | 0.96 | 0.008799121 | 5.95E-07 |
| White Matter Tracts | cTBS | Slc22a17 | solute carrier family 22 (organic cation transporter), member 17 | 59049 | -0.153594729 | 0.964 | 0.99 | 0.010319595 | 6.98E-07 |
| White Matter Tracts | cTBS | Smim13 | small integral membrane protein 13 | 108934 | -0.153100985 | 0.418 | 0.582 | 0.018265882 | 1.24E-06 |
| White Matter Tracts | cTBS | Tmem132a | transmembrane protein 132A | 98170 | -0.146683403 | 0.496 | 0.648 | 0.046609006 | 3.15E-06 |
| White Matter Tracts | cTBS | Jph3 | junctophilin 3 | 57340 | -0.143461799 | 0.237 | 0.375 | 0.047528763 | 3.22E-06 |
| White Matter Tracts | cTBS | Rtn1 | reticulon 1 | 104001 | -0.141832055 | 0.982 | 0.99 | 0.011747176 | 7.95E-07 |
| White Matter Tracts | cTBS | Snhg11 | small nucleolar RNA host gene 11 | 319317 | -0.134986386 | 0.914 | 0.93 | 0.02253315 | 1.52E-06 |
| White Matter Tracts | cTBS | Rpl21 | ribosomal protein L21 | 19933 | 0.143483302 | 1 | 1 | 0.014644055 | 9.91E-07 |
| White Matter Tracts | cTBS | Cox8a | cytochrome c oxidase subunit 8A | 12868 | 0.152716569 | 1 | 1 | 0.026500607 | 1.79E-06 |
| White Matter Tracts | cTBS | Rps21 | ribosomal protein S21 | 66481 | 0.153368289 | 1 | 1 | 0.019612584 | 1.33E-06 |
| White Matter Tracts | cTBS | Cox6a1 | cytochrome c oxidase subunit 6A1 | 12861 | 0.189933891 | 1 | 1 | 0.004682218 | 3.17E-07 |
| White Matter Tracts | cTBS | Rpl23 | ribosomal protein L23 | 65019 | 0.200256715 | 1 | 1 | 0.000407168 | 2.76E-08 |
| White Matter Tracts | cTBS | Rpl35 | ribosomal protein L35 | 66489 | 0.224337442 | 0.998 | 1 | 0.010046001 | 6.80E-07 |
| White Matter Tracts | cTBS | Cdc37l1 | cell division cycle 37-like 1 | 67072 | 0.251715611 | 0.245 | 0.125 | 0.008505936 | 5.76E-07 |
| White Matter Tracts | cTBS | Plekhb1 | pleckstrin homology domain containing, family B (evectins) member 1 | 27276 | 0.261595664 | 0.982 | 0.978 | 0.019060061 | 1.29E-06 |
| White Matter Tracts | cTBS | Lars2 | leucyl-tRNA synthetase, mitochondrial | 102436 | 0.264064302 | 0.285 | 0.167 | 0.029131291 | 1.97E-06 |

| | | | | | | | | | |
|---|---|---|---|---|---|---|---|---|---|
| White Matter Tracts | cTBS | Ftl1 | ferritin light polypeptide 1 | 14325 | 0.305588024 | 1 | 1 | 3.24E-07 | 2.19E-11 |
| White Matter Tracts | cTBS | Kif5a | kinesin family member 5A | 16572 | 0.31720512 | 0.992 | 0.99 | 1.21E-05 | 8.22E-10 |
| White Matter Tracts | cTBS | Mobp | myelin-associated oligodendrocytic basic protein | 17433 | 0.317797139 | 0.998 | 1 | 7.71E-06 | 5.22E-10 |
| White Matter Tracts | cTBS | Mbp | myelin basic protein | 17196 | 0.338416104 | 1 | 1 | 3.39E-09 | 2.29E-13 |
| White Matter Tracts | cTBS | Bcas1 | brain enriched myelin associated protein 1 | 76960 | 0.348352484 | 0.695 | 0.545 | 0.000721123 | 4.88E-08 |
| White Matter Tracts | cTBS | Rapgef4 | Rap guanine nucleotide exchange factor (GEF) 4 | 56508 | 0.367187139 | 0.843 | 0.795 | 1.53E-05 | 1.03E-09 |
| White Matter Tracts | cTBS | Uba52 | ubiquitin A-52 residue ribosomal protein fusion product 1 | 22186 | 0.485056077 | 0.968 | 0.958 | 1.50E-12 | 1.02E-16 |
| White Matter Tracts | cTBS | Fth1 | ferritin heavy polypeptide 1 | 14319 | 0.517891522 | 1 | 1 | 5.79E-25 | 3.92E-29 |
| White Matter Tracts | cTBS | Gm10076 | ribosomal protein L41 pseudogene | 100126819 | 0.565014967 | 0.938 | 0.887 | 5.47E-20 | 3.70E-24 |
| White Matter Tracts | cTBS | Gstp1 | glutathione S-transferase, pi 1 | 14870 | 0.685173258 | 0.707 | 0.597 | 3.50E-11 | 2.37E-15 |
| Caudoputamen | cTBS | Bc1 | brain cytoplasmic RNA 1 | 100568459 | -0.704317676 | 0.342 | 0.693 | 2.11E-61 | 1.43E-65 |
| Caudoputamen | cTBS | Ttr | transthyretin | 22139 | -0.326153983 | 0.002 | 0.141 | 4.66E-33 | 3.15E-37 |
| Caudoputamen | cTBS | Vsnl1 | visinin-like 1 | 26950 | -0.309329542 | 0.314 | 0.544 | 1.38E-22 | 9.34E-27 |
| Caudoputamen | cTBS | Slc17a7 | solute carrier family 17 (sodium-dependent inorganic phosphate cotrans | 72961 | -0.272338813 | 0.196 | 0.361 | 1.47E-14 | 9.94E-19 |
| Caudoputamen | cTBS | Olfm1 | olfactomedin 1 | 56177 | -0.266129212 | 0.585 | 0.764 | 1.65E-13 | 1.12E-17 |
| Caudoputamen | cTBS | Stmn1 | stathmin 1 | 16765 | -0.252160997 | 0.614 | 0.772 | 7.28E-09 | 4.93E-13 |
| Caudoputamen | cTBS | Gng3 | guanine nucleotide binding protein (G protein), gamma 3 | 14704 | -0.250711456 | 0.733 | 0.857 | 1.40E-09 | 9.47E-14 |
| Caudoputamen | cTBS | Ptgds | prostaglandin D2 synthase (brain) | 19215 | -0.241162297 | 0.671 | 0.813 | 4.81E-06 | 3.25E-10 |
| Caudoputamen | cTBS | Tubb2a | tubulin, beta 2A class IIA | 22151 | -0.237262089 | 0.595 | 0.713 | 2.58E-07 | 1.74E-11 |
| Caudoputamen | cTBS | Nap1l5 | nucleosome assembly protein 1-like 5 | 58243 | -0.220257779 | 0.513 | 0.643 | 1.11E-05 | 7.51E-10 |
| Caudoputamen | cTBS | Tuba1b | tubulin, alpha 1B | 22143 | -0.212309698 | 0.72 | 0.833 | 2.01E-06 | 1.36E-10 |
| Caudoputamen | cTBS | Atp1a3 | ATPase, Na+/K+ transporting, alpha 3 polypeptide | 232975 | -0.20502498 | 0.653 | 0.733 | 1.92E-05 | 1.30E-09 |
| Caudoputamen | cTBS | Ywhah | tyrosine 3-monooxygenase/tryptophan 5-monooxygenase activation pro | 22629 | -0.203692008 | 0.918 | 0.958 | 1.32E-07 | 8.95E-12 |
| Caudoputamen | cTBS | Tuba1a | tubulin, alpha 1A | 22142 | -0.201513444 | 0.978 | 0.984 | 3.39E-08 | 2.29E-12 |
| Caudoputamen | cTBS | Cdk5r2 | cyclin-dependent kinase 5, regulatory subunit 2 (p39) | 12570 | -0.201139872 | 0.338 | 0.433 | 0.000573152 | 3.88E-08 |
| Caudoputamen | cTBS | Ikbkb | inhibitor of kappaB kinase beta | 16150 | -0.199450592 | 0.214 | 0.32 | 2.22E-05 | 1.50E-09 |
| Caudoputamen | cTBS | Ly6h | lymphocyte antigen 6 complex, locus H | 23934 | -0.198346836 | 0.692 | 0.779 | 8.21E-05 | 5.55E-09 |
| Caudoputamen | cTBS | Sncb | synuclein, beta | 104069 | -0.197357629 | 0.462 | 0.544 | 0.007066572 | 4.78E-07 |
| Caudoputamen | cTBS | Tubb4b | tubulin, beta 4B class IVB | 227613 | -0.19667221 | 0.545 | 0.625 | 0.000546345 | 3.70E-08 |
| Caudoputamen | cTBS | Cck | cholecystokinin | 12424 | -0.193032292 | 0.271 | 0.416 | 6.68E-08 | 4.52E-12 |
| Caudoputamen | cTBS | Tagln3 | transgelin 3 | 56370 | -0.188334531 | 0.648 | 0.725 | 0.001625043 | 1.10E-07 |
| Caudoputamen | cTBS | Gnas | GNAS (guanine nucleotide binding protein, alpha stimulating) complex lo | 14683 | -0.177807492 | 0.84 | 0.9 | 0.000220315 | 1.49E-08 |
| Caudoputamen | cTBS | Prkar1b | protein kinase, cAMP dependent regulatory, type I beta | 19085 | -0.174957273 | 0.609 | 0.728 | 0.000102757 | 6.95E-09 |
| Caudoputamen | cTBS | Atp6v1c1 | ATPase, H+ transporting, lysosomal V1 subunit C1 | 66335 | -0.166724918 | 0.63 | 0.69 | 0.034745517 | 2.35E-06 |
| Caudoputamen | cTBS | Atp6v1g2 | ATPase, H+ transporting, lysosomal V1 subunit G2 | 66237 | -0.159546273 | 0.779 | 0.848 | 0.008914432 | 6.03E-07 |
| Caudoputamen | cTBS | Zwint | ZW10 interactor | 52696 | -0.158745677 | 0.819 | 0.889 | 0.027501402 | 1.86E-06 |
| Caudoputamen | cTBS | Snhg9 | small nucleolar RNA host gene 9 | 73474 | -0.154812189 | 0.155 | 0.252 | 0.000160041 | 1.08E-08 |
| Caudoputamen | cTBS | Calm1 | calmodulin 1 | 12313 | -0.115803336 | 1 | 1 | 2.81E-05 | 1.90E-09 |
| Caudoputamen | cTBS | mt-Co3 | NA | NA | 0.073890073 | 1 | 1 | 0.046084267 | 3.12E-06 |
| Caudoputamen | cTBS | Gpr139 | G protein-coupled receptor 139 | 209776 | 0.078920768 | 0.112 | 0.052 | 0.00897228 | 6.07E-07 |
| Caudoputamen | cTBS | mt-Atp6 | NA | NA | 0.082285255 | 1 | 1 | 0.000337469 | 2.28E-08 |
| Caudoputamen | cTBS | Ngdn | neuroguidin, EIF4E binding protein | 68966 | 0.085301829 | 0.154 | 0.086 | 0.023254536 | 1.57E-06 |
| Caudoputamen | cTBS | Ppp2cb | protein phosphatase 2 (formerly 2A), catalytic subunit, beta isoform | 19053 | 0.086728982 | 0.118 | 0.06 | 0.045863103 | 3.10E-06 |
| Caudoputamen | cTBS | Slc7a10 | solute carrier family 7 (cationic amino acid transporter, y+ system), mem | 53896 | 0.087363309 | 0.174 | 0.1 | 0.019610079 | 1.33E-06 |
| Caudoputamen | cTBS | AI854703 | expressed sequence AI854703 | 243373 | 0.089868213 | 0.158 | 0.09 | 0.031563864 | 2.14E-06 |
| Caudoputamen | cTBS | Bmp1 | bone morphogenetic protein 1 | 12153 | 0.089911578 | 0.158 | 0.086 | 0.008170507 | 5.53E-07 |
| Caudoputamen | cTBS | Rpl13 | ribosomal protein L13 | 270106 | 0.090881783 | 1 | 1 | 0.017044571 | 1.15E-06 |
| Caudoputamen | cTBS | Ckb | creatine kinase, brain | 12709 | 0.091838068 | 1 | 1 | 3.93E-05 | 2.66E-09 |
| Caudoputamen | cTBS | Tmem80 | transmembrane protein 80 | 71448 | 0.092756731 | 0.146 | 0.08 | 0.027157097 | 1.84E-06 |
| Caudoputamen | cTBS | Me2 | malic enzyme 2, NAD(+)-dependent, mitochondrial | 107029 | 0.095856804 | 0.161 | 0.092 | 0.030245957 | 2.05E-06 |
| Caudoputamen | cTBS | Rplp1 | ribosomal protein, large, P1 | 56040 | 0.096395004 | 1 | 1 | 0.003704521 | 2.51E-07 |
| Caudoputamen | cTBS | Slc30a10 | solute carrier family 30, member 10 | 226781 | 0.097045834 | 0.197 | 0.122 | 0.041418405 | 2.80E-06 |
| Caudoputamen | cTBS | Rngtt | RNA guanylyltransferase and 5'-phosphatase | 24018 | 0.099527177 | 0.179 | 0.102 | 0.006782545 | 4.59E-07 |
| Caudoputamen | cTBS | Leprot | leptin receptor overlapping transcript | 230514 | 0.101956999 | 0.196 | 0.115 | 0.007072742 | 4.79E-07 |
| Caudoputamen | cTBS | Ubb | ubiquitin B | 22187 | 0.10383519 | 1 | 1 | 0.001155313 | 7.82E-08 |
| Caudoputamen | cTBS | Mapk6 | mitogen-activated protein kinase 6 | 50772 | 0.104936171 | 0.215 | 0.137 | 0.038981868 | 2.64E-06 |
| Caudoputamen | cTBS | Mgat5b | mannoside acetylglucosaminyltransferase 5, isoenzyme B | 268510 | 0.105717146 | 0.422 | 0.31 | 0.028703164 | 1.94E-06 |
| Caudoputamen | cTBS | Esam | endothelial cell-specific adhesion molecule | 69524 | 0.106097339 | 0.159 | 0.085 | 0.002548146 | 1.72E-07 |
| Caudoputamen | cTBS | Eml5 | echinoderm microtubule associated protein like 5 | 319670 | 0.107269628 | 0.301 | 0.205 | 0.017347444 | 1.17E-06 |
| Caudoputamen | cTBS | Arf3 | ADP-ribosylation factor 3 | 11842 | 0.107834086 | 0.999 | 0.994 | 0.027089307 | 1.83E-06 |
| Caudoputamen | cTBS | Cox5b | cytochrome c oxidase subunit 5B | 12859 | 0.110870975 | 1 | 0.999 | 0.001662978 | 1.13E-07 |
| Caudoputamen | cTBS | Btrc | beta-transducin repeat containing protein | 12234 | 0.110893671 | 0.266 | 0.174 | 0.009035169 | 6.11E-07 |
| Caudoputamen | cTBS | Hnrnpm | heterogeneous nuclear ribonucleoprotein M | 76936 | 0.111647034 | 0.512 | 0.379 | 0.013281261 | 8.99E-07 |
| Caudoputamen | cTBS | Rpl37 | ribosomal protein L37 | 67281 | 0.111932077 | 1 | 1 | 0.00036571 | 2.47E-08 |
| Caudoputamen | cTBS | Letmd1 | LETM1 domain containing 1 | 68614 | 0.11323381 | 0.27 | 0.172 | 0.002533393 | 1.71E-07 |
| Caudoputamen | cTBS | Slc22a23 | solute carrier family 22, member 23 | 73102 | 0.113779678 | 0.246 | 0.154 | 0.002504694 | 1.69E-07 |
| Caudoputamen | cTBS | Gm27032 | predicted gene, 27032 | 118568482 | 0.113793461 | 0.217 | 0.13 | 0.002385642 | 1.61E-07 |
| Caudoputamen | cTBS | Sema7a | sema domain, immunoglobulin domain (Ig), and GPI membrane anchor, ( | 20361 | 0.114142869 | 0.179 | 0.1 | 0.002879547 | 1.95E-07 |
| Caudoputamen | cTBS | Atp5j2 | ATP synthase, H+ transporting, mitochondrial F0 complex, subunit F2 | 57423 | 0.115780641 | 1 | 0.999 | 0.003896059 | 2.64E-07 |
| Caudoputamen | cTBS | Slc6a6 | solute carrier family 6 (neurotransmitter transporter, taurine), member 6 | 21366 | 0.116471156 | 0.35 | 0.242 | 0.005002295 | 3.38E-07 |
| Caudoputamen | cTBS | B3gnt2 | UDP-GlcNAc:betaGal beta-1,3-N-acetylglucosaminyltransferase 2 | 53625 | 0.117776775 | 0.463 | 0.335 | 0.002861953 | 1.94E-07 |
| Caudoputamen | cTBS | Inpp5j | inositol polyphosphate 5-phosphatase J | 170835 | 0.118319967 | 0.32 | 0.215 | 0.003198792 | 2.16E-07 |
| Caudoputamen | cTBS | Cox4i1 | cytochrome c oxidase subunit 4I1 | 12857 | 0.118694659 | 1 | 1 | 4.82E-06 | 3.26E-10 |
| Caudoputamen | cTBS | Actg1 | actin, gamma, cytoplasmic 1 | 11465 | 0.118837336 | 1 | 1 | 8.04E-06 | 5.44E-10 |
| Caudoputamen | cTBS | Actn1 | actinin, alpha 1 | 109711 | 0.119228558 | 0.489 | 0.378 | 0.014583345 | 9.87E-07 |
| Caudoputamen | cTBS | Rbfox1 | RNA binding protein, fox-1 homolog (C. elegans) 1 | 268859 | 0.119350096 | 0.229 | 0.135 | 0.000393743 | 2.66E-08 |
| Caudoputamen | cTBS | Fth1 | ferritin heavy polypeptide 1 | 14319 | 0.120959545 | 1 | 1 | 0.002501526 | 1.69E-07 |
| Caudoputamen | cTBS | Dach1 | dachshund family transcription factor 1 | 13134 | 0.121595099 | 0.415 | 0.312 | 0.03131483 | 2.12E-06 |
| Caudoputamen | cTBS | Ppp3ca | protein phosphatase 3, catalytic subunit, alpha isoform | 19055 | 0.121717324 | 0.999 | 0.999 | 0.000819373 | 5.54E-08 |
| Caudoputamen | cTBS | Atp5mpl | ATP synthase membrane subunit 6.8PL | 70257 | 0.123498108 | 0.996 | 0.997 | 0.000726318 | 4.91E-08 |
| Caudoputamen | cTBS | mt-Co1 | NA | NA | 0.124182091 | 1 | 1 | 1.31E-09 | 8.86E-14 |
| Caudoputamen | cTBS | Hist1h1c | NA | NA | 0.125851591 | 0.348 | 0.244 | 0.005506124 | 3.73E-07 |
| Caudoputamen | cTBS | Sipa1l3 | signal-induced proliferation-associated 1 like 3 | 74206 | 0.126011996 | 0.307 | 0.218 | 0.036496636 | 2.47E-06 |
| Caudoputamen | cTBS | Ncdn | neurochondrin | 26562 | 0.127191245 | 0.999 | 1 | 2.41E-05 | 1.63E-09 |
| Caudoputamen | cTBS | Mn1 | meningioma 1 | 433938 | 0.12965373 | 0.328 | 0.229 | 0.007459425 | 5.05E-07 |
| Caudoputamen | cTBS | Snd1 | staphylococcal nuclease and tudor domain containing 1 | 56463 | 0.129942611 | 0.279 | 0.19 | 0.014501776 | 9.81E-07 |
| Caudoputamen | cTBS | Celf3 | CUGBP, Elav-like family member 3 | 78784 | 0.131654606 | 0.41 | 0.303 | 0.011871816 | 8.03E-07 |
| Caudoputamen | cTBS | Rps27 | ribosomal protein S27 | 57294 | 0.134115747 | 0.982 | 0.977 | 0.006009891 | 4.07E-07 |
| Caudoputamen | cTBS | Spock3 | sparc/osteonectin, cwcv and kazal-like domains proteoglycan 3 | 72902 | 0.134371573 | 0.885 | 0.812 | 0.018842101 | 1.28E-06 |
| Caudoputamen | cTBS | Ppp1r1b | protein phosphatase 1, regulatory inhibitor subunit 1B | 19049 | 0.135207913 | 0.999 | 0.991 | 0.000603333 | 4.08E-08 |
| Caudoputamen | cTBS | Vat1l | vesicle amine transport protein 1 like | 270097 | 0.13587445 | 0.439 | 0.332 | 0.025713595 | 1.74E-06 |
| Caudoputamen | cTBS | Rpl36 | ribosomal protein L36 | 54217 | 0.136226239 | 0.997 | 0.999 | 6.53E-06 | 4.42E-10 |
| Caudoputamen | cTBS | Neto2 | neuropilin (NRP) and tolloid (TLL)-like 2 | 74513 | 0.13642654 | 0.446 | 0.333 | 0.002599465 | 1.76E-07 |
| Caudoputamen | cTBS | Myo1b | myosin IB | 17912 | 0.136964089 | 0.308 | 0.208 | 0.001503606 | 1.02E-07 |
| Caudoputamen | cTBS | Ftl1 | ferritin light polypeptide 1 | 14325 | 0.137730033 | 0.987 | 0.991 | 0.005039147 | 3.41E-07 |
| Caudoputamen | cTBS | Tmsb10 | thymosin, beta 10 | 19240 | 0.137739807 | 1 | 0.999 | 4.34E-08 | 2.94E-12 |
| Caudoputamen | cTBS | Plxnd1 | plexin D1 | 67784 | 0.13808934 | 0.371 | 0.273 | 0.024517659 | 1.66E-06 |
| Caudoputamen | cTBS | Adcy5 | adenylate cyclase 5 | 224129 | 0.139509589 | 0.96 | 0.917 | 0.001481751 | 1.00E-07 |
| Caudoputamen | cTBS | Foxo1 | forkhead box O1 | 56458 | 0.140020388 | 0.465 | 0.339 | 0.000424397 | 2.87E-08 |
| Caudoputamen | cTBS | Prkcb | protein kinase C, beta | 18751 | 0.140629723 | 0.966 | 0.945 | 0.0300527 | 2.03E-06 |
| Caudoputamen | cTBS | Ppp2r2a | protein phosphatase 2, regulatory subunit B, alpha | 71978 | 0.142169695 | 0.738 | 0.632 | 0.045274031 | 3.06E-06 |
| Caudoputamen | cTBS | Inf2 | inverted formin, FH2 and WH2 domain containing | 70435 | 0.142378088 | 0.916 | 0.84 | 0.004666846 | 3.16E-07 |
| Caudoputamen | cTBS | Rasd2 | RASD family, member 2 | 75141 | 0.142989887 | 0.968 | 0.935 | 0.004040561 | 2.73E-07 |
| Caudoputamen | cTBS | Stub1 | STIP1 homology and U-Box containing protein 1 | 56424 | 0.143304493 | 0.61 | 0.479 | 0.013303789 | 9.00E-07 |
| Caudoputamen | cTBS | Rpl35a | ribosomal protein L35A | 57808 | 0.143809726 | 1 | 0.996 | 3.22E-07 | 2.18E-11 |
| Caudoputamen | cTBS | Ppp1ca | protein phosphatase 1 catalytic subunit alpha | 19045 | 0.14496957 | 0.985 | 0.977 | 0.000326126 | 2.21E-08 |
| Caudoputamen | cTBS | Lrrc10b | leucine rich repeat containing 10B | 278795 | 0.145100247 | 0.815 | 0.707 | 0.024447141 | 1.65E-06 |
| Caudoputamen | cTBS | Fnbp1l | formin binding protein 1-like | 214459 | 0.14576677 | 0.421 | 0.307 | 0.001095202 | 7.41E-08 |
| Caudoputamen | cTBS | Nrgn | neurogranin | 64011 | 0.148216192 | 1 | 1 | 0.00729216 | 4.93E-07 |
| Caudoputamen | cTBS | Myh10 | myosin, heavy polypeptide 10, non-muscle | 77579 | 0.149483725 | 0.625 | 0.485 | 0.000648375 | 4.39E-08 |
| Caudoputamen | cTBS | Ndufa1 | NADH:ubiquinone oxidoreductase subunit A1 | 54405 | 0.151282016 | 0.887 | 0.843 | 0.030221999 | 2.05E-06 |
| Caudoputamen | cTBS | Rps29 | ribosomal protein S29 | 20090 | 0.152764044 | 0.999 | 0.998 | 3.04E-06 | 2.05E-10 |
| Caudoputamen | cTBS | Uqcr11 | ubiquinol-cytochrome c reductase, complex III subunit XI | 66594 | 0.155508476 | 1 | 1 | 1.67E-08 | 1.13E-12 |
| Caudoputamen | cTBS | Kcnip2 | Kv channel-interacting protein 2 | 80906 | 0.156058548 | 0.856 | 0.798 | 0.018835484 | 1.27E-06 |
| Caudoputamen | cTBS | Tesc | tescalcin | 57816 | 0.156513445 | 0.921 | 0.847 | 0.000742104 | 5.02E-08 |
| Caudoputamen | cTBS | Dlg4 | discs large MAGUK scaffold protein 4 | 13385 | 0.15679376 | 0.739 | 0.612 | 0.005891252 | 3.99E-07 |
| Caudoputamen | cTBS | Atp5md | ATP synthase membrane subunit DAPIT | 66477 | 0.15708449 | 1 | 1 | 6.05E-11 | 4.09E-15 |
| Caudoputamen | cTBS | Gas5 | growth arrest specific 5 | 14455 | 0.158489788 | 0.969 | 0.944 | 0.000122458 | 8.29E-09 |
| Caudoputamen | cTBS | Bcl11b | B cell leukemia/lymphoma 11B | 58208 | 0.160873103 | 0.738 | 0.621 | 0.002306853 | 1.56E-07 |
| Caudoputamen | cTBS | Rasgef1b | RasGEF domain family, member 1B | 320292 | 0.161114756 | 0.426 | 0.31 | 0.000304541 | 2.06E-08 |
| Caudoputamen | cTBS | Bex1 | brain expressed X-linked 1 | 19716 | 0.161306201 | 0.846 | 0.763 | 0.011186952 | 7.57E-07 |
| Caudoputamen | cTBS | Lmo2 | LIM domain only 2 | 16909 | 0.162697087 | 0.49 | 0.372 | 0.000412324 | 2.79E-08 |
| Caudoputamen | cTBS | Rpl23 | ribosomal protein L23 | 65019 | 0.162824741 | 0.998 | 0.996 | 6.81E-08 | 4.61E-12 |
| Caudoputamen | cTBS | Cox6c | cytochrome c oxidase subunit 6C | 12864 | 0.16360121 | 1 | 1 | 6.00E-16 | 4.06E-20 |
| Caudoputamen | cTBS | Hmgn2 | high mobility group nucleosomal binding domain 2 | 15331 | 0.164094691 | 0.558 | 0.443 | 0.004726878 | 3.20E-07 |
| Caudoputamen | cTBS | St8sia3 | ST8 alpha-N-acetyl-neuraminide alpha-2,8-sialyltransferase 3 | 20451 | 0.168704084 | 0.863 | 0.765 | 0.001258466 | 8.52E-08 |
| Caudoputamen | cTBS | Eno1 | enolase 1, alpha non-neuron | 13806 | 0.170108189 | 0.893 | 0.864 | 0.000372961 | 2.52E-08 |
| Caudoputamen | cTBS | Cox7c | cytochrome c oxidase subunit 7C | 12867 | 0.170578163 | 1 | 1 | 1.37E-13 | 9.28E-18 |
| Caudoputamen | cTBS | Kctd17 | potassium channel tetramerisation domain containing 17 | 72844 | 0.177165554 | 0.834 | 0.739 | 0.018578133 | 1.26E-06 |
| Caudoputamen | cTBS | Drd1 | dopamine receptor D1 | 13488 | 0.180049329 | 0.721 | 0.562 | 7.33E-07 | 4.96E-11 |
| Caudoputamen | cTBS | Atp5k | ATP synthase, H+ transporting, mitochondrial F1F0 complex, subunit E | 11958 | 0.184552498 | 0.998 | 0.999 | 2.18E-11 | 1.48E-15 |
| Caudoputamen | cTBS | Cnr1 | cannabinoid receptor 1 (brain) | 12801 | 0.185031018 | 0.654 | 0.532 | 0.001935068 | 1.31E-07 |
| Caudoputamen | cTBS | Gpr88 | G-protein coupled receptor 88 | 64378 | 0.185868182 | 0.993 | 0.981 | 7.70E-10 | 5.21E-14 |
| Caudoputamen | cTBS | Rps28 | ribosomal protein S28 | 54127 | 0.199990876 | 0.943 | 0.935 | 5.83E-05 | 3.94E-09 |
| Caudoputamen | cTBS | Rasgrp2 | RAS, guanyl releasing protein 2 | 19395 | 0.207947872 | 0.805 | 0.674 | 1.83E-06 | 1.24E-10 |
| Caudoputamen | cTBS | Polr2l | polymerase (RNA) II (DNA directed) polypeptide L | 66491 | 0.208696574 | 0.507 | 0.362 | 2.31E-06 | 1.57E-10 |
| Caudoputamen | cTBS | Rps21 | ribosomal protein S21 | 66481 | 0.209174389 | 1 | 1 | 1.42E-22 | 9.59E-27 |
| Caudoputamen | cTBS | Rgs9 | regulator of G-protein signaling 9 | 19739 | 0.211492162 | 0.953 | 0.896 | 6.05E-09 | 4.10E-13 |
| Caudoputamen | cTBS | Ptpn5 | protein tyrosine phosphatase, non-receptor type 5 | 19259 | 0.212627431 | 0.986 | 0.966 | 1.23E-10 | 8.31E-15 |
| Caudoputamen | cTBS | Pcp4 | Purkinje cell protein 4 | 18546 | 0.213168599 | 1 | 1 | 1.47E-12 | 9.97E-17 |
| Caudoputamen | cTBS | Rpl38 | ribosomal protein L38 | 67671 | 0.214068141 | 1 | 1 | 1.26E-19 | 8.51E-24 |
| Caudoputamen | cTBS | AC149090.1 | NA | NA | 0.234910805 | 0.727 | 0.563 | 1.62E-07 | 1.10E-11 |
| Caudoputamen | cTBS | Adora2a | adenosine A2a receptor | 11540 | 0.237865396 | 0.748 | 0.614 | 2.48E-07 | 1.68E-11 |
| Caudoputamen | cTBS | Gstp1 | glutathione S-transferase, pi 1 | 14870 | 0.355673161 | 0.472 | 0.27 | 4.87E-19 | 3.30E-23 |
| Caudoputamen | cTBS | Rpl35 | ribosomal protein L35 | 66489 | 0.404741398 | 0.983 | 0.965 | 1.09E-32 | 7.36E-37 |
| Caudoputamen | cTBS | Uba52 | ubiquitin A-52 residue ribosomal protein fusion product 1 | 22186 | 0.506746535 | 0.826 | 0.78 | 6.55E-29 | 4.43E-33 |
| Caudoputamen | cTBS | Gm10076 | ribosomal protein L41 pseudogene | 100126819 | 1.049698447 | 0.772 | 0.463 | 1.59E-94 | 1.07E-98 |
| Lateral Septal Complex | cTBS | Ttr | transthyretin | 22139 | -1.951188266 | 0.101 | 0.321 | 6.79E-10 | 4.60E-14 |
| Lateral Septal Complex | cTBS | Bc1 | brain cytoplasmic RNA 1 | 100568459 | -1.114948558 | 0.585 | 0.909 | 2.59E-32 | 1.75E-36 |
| Lateral Septal Complex | cTBS | Tmem158 | transmembrane protein 158 | 72309 | -0.43957192 | 0.735 | 0.838 | 1.26E-06 | 8.53E-11 |
| Lateral Septal Complex | cTBS | Bok | BCL2-related ovarian killer | 51800 | -0.406603618 | 0.477 | 0.608 | 0.025701243 | 1.74E-06 |
| Lateral Septal Complex | cTBS | Necab2 | N-terminal EF-hand calcium binding protein 2 | 117148 | -0.395191666 | 0.686 | 0.786 | 0.013799946 | 9.34E-07 |
| Lateral Septal Complex | cTBS | Gria1 | glutamate receptor, ionotropic, AMPA1 (alpha 1) | 14799 | -0.334501905 | 0.691 | 0.815 | 0.003208812 | 2.17E-07 |
| Lateral Septal Complex | cTBS | Lrrc17 | leucine rich repeat containing 17 | 74511 | -0.323409984 | 0.119 | 0.334 | 6.96E-09 | 4.71E-13 |
| Lateral Septal Complex | cTBS | Lmo4 | LIM domain only 4 | 16911 | -0.315996228 | 0.317 | 0.507 | 0.000422133 | 2.86E-08 |
| Lateral Septal Complex | cTBS | Cnih2 | cornichon family AMPA receptor auxiliary protein 2 | 12794 | -0.289620486 | 0.92 | 0.961 | 0.010041113 | 6.79E-07 |
| Lateral Septal Complex | cTBS | Calm1 | calmodulin 1 | 12313 | -0.258240594 | 1 | 1 | 0.000126483 | 8.56E-09 |
| Lateral Septal Complex | cTBS | Cdk5r2 | cyclin-dependent kinase 5, regulatory subunit 2 (p39) | 12570 | -0.257546971 | 0.273 | 0.441 | 0.004223403 | 2.86E-07 |
| Lateral Septal Complex | cTBS | Cadm3 | cell adhesion molecule 3 | 94332 | -0.21409173 | 0.387 | 0.564 | 0.038008236 | 2.57E-06 |
| Lateral Septal Complex | cTBS | Tuba1a | tubulin, alpha 1A | 22142 | -0.21235979 | 0.985 | 0.987 | 0.005714572 | 3.87E-07 |
| Lateral Septal Complex | cTBS | Leprotl1 | leptin receptor overlapping transcript-like 1 | 68192 | -0.200108197 | 0.268 | 0.433 | 0.044679033 | 3.02E-06 |
| Lateral Septal Complex | cTBS | Ndnf | neuron-derived neurotrophic factor | 68169 | -0.181717848 | 0.057 | 0.175 | 0.004308534 | 2.92E-07 |
| Lateral Septal Complex | cTBS | mt-Cytb | NA | NA | -0.153773853 | 1 | 1 | 0.000795164 | 5.38E-08 |
| Lateral Septal Complex | cTBS | Fau | Finkel-Biskis-Reilly murine sarcoma virus (FBR-MuSV) ubiquitously expres | 14109 | 0.143072066 | 1 | 1 | 0.012421034 | 8.41E-07 |
| Lateral Septal Complex | cTBS | Rplp1 | ribosomal protein, large, P1 | 56040 | 0.15327933 | 1 | 1 | 0.000852162 | 5.77E-08 |
| Lateral Septal Complex | cTBS | Rpl11 | ribosomal protein L11 | 67025 | 0.153736104 | 0.995 | 1 | 0.036488634 | 2.47E-06 |
| Lateral Septal Complex | cTBS | Rpl35a | ribosomal protein L35A | 57808 | 0.167978277 | 1 | 1 | 0.032313001 | 2.19E-06 |
| Lateral Septal Complex | cTBS | Atpif1 | ATPase inhibitory factor 1 | 11983 | 0.172157193 | 1 | 0.997 | 0.041395915 | 2.80E-06 |
| Lateral Septal Complex | cTBS | Uqcr11 | ubiquinol-cytochrome c reductase, complex III subunit XI | 66594 | 0.178209474 | 1 | 0.992 | 0.038462064 | 2.60E-06 |
| Lateral Septal Complex | cTBS | Rpl36 | ribosomal protein L36 | 54217 | 0.192380129 | 0.997 | 1 | 0.000603615 | 4.08E-08 |
| Lateral Septal Complex | cTBS | Cox7c | cytochrome c oxidase subunit 7C | 12867 | 0.195564926 | 1 | 1 | 0.00012135 | 8.21E-09 |
| Lateral Septal Complex | cTBS | Rpl37a | ribosomal protein L37a | 19981 | 0.20717611 | 1 | 0.997 | 0.000409264 | 2.77E-08 |
| Lateral Septal Complex | cTBS | Igsf1 | immunoglobulin superfamily, member 1 | 209268 | 0.210282896 | 0.245 | 0.11 | 0.013836464 | 9.36E-07 |
| Lateral Septal Complex | cTBS | Rpl39 | ribosomal protein L39 | 67248 | 0.216764026 | 0.997 | 0.987 | 0.007565678 | 5.12E-07 |
| Lateral Septal Complex | cTBS | Nts | neurotensin | 67405 | 0.22128398 | 0.541 | 0.337 | 0.000963977 | 6.52E-08 |
| Lateral Septal Complex | cTBS | Rpl37 | ribosomal protein L37 | 67281 | 0.230142733 | 1 | 1 | 3.02E-10 | 2.04E-14 |
| Lateral Septal Complex | cTBS | Ndufb1-ps | NA | NA | 0.239355893 | 0.954 | 0.898 | 0.008800166 | 5.95E-07 |
| Lateral Septal Complex | cTBS | Atp5k | ATP synthase, H+ transporting, mitochondrial F1F0 complex, subunit E | 11958 | 0.253587518 | 0.997 | 0.987 | 2.94E-06 | 1.99E-10 |
| Lateral Septal Complex | cTBS | Cox17 | cytochrome c oxidase assembly protein 17, copper chaperone | 12856 | 0.256922182 | 0.892 | 0.838 | 0.003805351 | 2.58E-07 |
| Lateral Septal Complex | cTBS | Rps21 | ribosomal protein S21 | 66481 | 0.258429401 | 1 | 1 | 4.75E-12 | 3.21E-16 |
| Lateral Septal Complex | cTBS | Rps29 | ribosomal protein S29 | 20090 | 0.268069665 | 1 | 1 | 3.97E-07 | 2.69E-11 |
| Lateral Septal Complex | cTBS | Rps27 | ribosomal protein S27 | 57294 | 0.273182668 | 0.997 | 0.995 | 4.76E-08 | 3.22E-12 |
| Lateral Septal Complex | cTBS | Polr2l | polymerase (RNA) II (DNA directed) polypeptide L | 66491 | 0.326951536 | 0.624 | 0.491 | 0.000841713 | 5.70E-08 |
| Lateral Septal Complex | cTBS | Rpl38 | ribosomal protein L38 | 67671 | 0.327903228 | 1 | 1 | 1.23E-21 | 8.32E-26 |
| Lateral Septal Complex | cTBS | Rps28 | ribosomal protein S28 | 54127 | 0.329211043 | 0.995 | 0.958 | 8.13E-06 | 5.50E-10 |
| Lateral Septal Complex | cTBS | Gstp1 | glutathione S-transferase, pi 1 | 14870 | 0.589590609 | 0.714 | 0.449 | 4.34E-16 | 2.94E-20 |
| Lateral Septal Complex | cTBS | Rpl35 | ribosomal protein L35 | 66489 | 0.654136538 | 0.995 | 0.948 | 5.72E-40 | 3.87E-44 |
| Lateral Septal Complex | cTBS | Uba52 | ubiquitin A-52 residue ribosomal protein fusion product 1 | 22186 | 0.784122465 | 0.894 | 0.705 | 2.92E-28 | 1.98E-32 |
| Lateral Septal Complex | cTBS | Gm10076 | ribosomal protein L41 pseudogene | 100126819 | 1.305172032 | 0.861 | 0.496 | 7.26E-56 | 4.92E-60 |
| Striatum Ventral Region | cTBS | Bc1 | brain cytoplasmic RNA 1 | 100568459 | -0.926359105 | 0.48 | 0.83 | 8.05E-52 | 5.45E-56 |
| Striatum Ventral Region | cTBS | Ptgds | prostaglandin D2 synthase (brain) | 19215 | -0.353222369 | 0.694 | 0.79 | 0.024112355 | 1.63E-06 |
| Striatum Ventral Region | cTBS | Malat1 | metastasis associated lung adenocarcinoma transcript 1 (non-coding RNA | 72289 | -0.307512671 | 0.573 | 0.643 | 0.017126764 | 1.16E-06 |
| Striatum Ventral Region | cTBS | Ttr | transthyretin | 22139 | -0.241571137 | 0.004 | 0.126 | 1.19E-17 | 8.06E-22 |
| Striatum Ventral Region | cTBS | Atp1a3 | ATPase, Na+/K+ transporting, alpha 3 polypeptide | 232975 | -0.240955314 | 0.73 | 0.818 | 0.000127402 | 8.62E-09 |
| Striatum Ventral Region | cTBS | Gm42418 | NA | NA | -0.197429186 | 0.92 | 0.958 | 0.000910304 | 6.16E-08 |
| Striatum Ventral Region | cTBS | Snhg9 | small nucleolar RNA host gene 9 | 73474 | -0.193927998 | 0.177 | 0.304 | 0.000111408 | 7.54E-09 |
| Striatum Ventral Region | cTBS | Tatdn1 | TatD DNase domain containing 1 | 69694 | -0.169123376 | 0.129 | 0.232 | 0.00278797 | 1.89E-07 |
| Striatum Ventral Region | cTBS | Lrrc17 | leucine rich repeat containing 17 | 74511 | -0.164650636 | 0.059 | 0.17 | 4.56E-07 | 3.09E-11 |
| Striatum Ventral Region | cTBS | Ywhah | tyrosine 3-monooxygenase/tryptophan 5-monooxygenase activation pro | 22629 | -0.164605595 | 0.959 | 0.974 | 0.049645714 | 3.36E-06 |
| Striatum Ventral Region | cTBS | 2900097C17Rik | NA | NA | -0.159355239 | 0.966 | 0.977 | 0.025927503 | 1.75E-06 |
| Striatum Ventral Region | cTBS | Calm1 | calmodulin 1 | 12313 | -0.125777514 | 1 | 1 | 0.010214258 | 6.91E-07 |
| Striatum Ventral Region | cTBS | mt-Atp6 | NA | NA | 0.110562868 | 1 | 1 | 0.012505683 | 8.46E-07 |
| Striatum Ventral Region | cTBS | Rpl35a | ribosomal protein L35A | 57808 | 0.12619547 | 1 | 1 | 0.014499159 | 9.81E-07 |
| Striatum Ventral Region | cTBS | Atp5mpl | ATP synthase membrane subunit 6.8PL | 70257 | 0.139449273 | 0.996 | 0.994 | 0.010923237 | 7.39E-07 |
| Striatum Ventral Region | cTBS | Rps15 | ribosomal protein S15 | 20054 | 0.140054271 | 0.999 | 0.988 | 0.024058385 | 1.63E-06 |
| Striatum Ventral Region | cTBS | Rps29 | ribosomal protein S29 | 20090 | 0.14129544 | 0.999 | 0.998 | 0.026959812 | 1.82E-06 |
| Striatum Ventral Region | cTBS | Uqcr11 | ubiquinol-cytochrome c reductase, complex III subunit XI | 66594 | 0.155853407 | 0.999 | 1 | 0.004157576 | 2.81E-07 |
| Striatum Ventral Region | cTBS | Cox8a | cytochrome c oxidase subunit 8A | 12868 | 0.161881783 | 1 | 1 | 2.49E-05 | 1.68E-09 |
| Striatum Ventral Region | cTBS | Rps21 | ribosomal protein S21 | 66481 | 0.167631544 | 1 | 1 | 4.35E-06 | 2.94E-10 |
| Striatum Ventral Region | cTBS | mt-Co1 | NA | NA | 0.170373687 | 1 | 1 | 3.11E-06 | 2.11E-10 |
| Striatum Ventral Region | cTBS | Ubb | ubiquitin B | 22187 | 0.172388618 | 1 | 1 | 2.45E-09 | 1.66E-13 |
| Striatum Ventral Region | cTBS | Cox6a1 | cytochrome c oxidase subunit 6A1 | 12861 | 0.17622877 | 0.997 | 0.992 | 0.001443039 | 9.76E-08 |
| Striatum Ventral Region | cTBS | Cox6c | cytochrome c oxidase subunit 6C | 12864 | 0.179055585 | 1 | 1 | 1.08E-10 | 7.34E-15 |
| Striatum Ventral Region | cTBS | Ndufa3 | NADH:ubiquinone oxidoreductase subunit A3 | 66091 | 0.180838083 | 0.979 | 0.984 | 0.000323531 | 2.19E-08 |
| Striatum Ventral Region | cTBS | Rps27 | ribosomal protein S27 | 57294 | 0.195996845 | 0.995 | 0.991 | 0.000491045 | 3.32E-08 |
| Striatum Ventral Region | cTBS | Atp5md | ATP synthase membrane subunit DAPIT | 66477 | 0.209517683 | 1 | 1 | 6.38E-13 | 4.32E-17 |
| Striatum Ventral Region | cTBS | Atp5k | ATP synthase, H+ transporting, mitochondrial F1F0 complex, subunit E | 11958 | 0.214402215 | 0.999 | 0.997 | 7.26E-08 | 4.91E-12 |
| Striatum Ventral Region | cTBS | Eno1 | enolase 1, alpha non-neuron | 13806 | 0.227569433 | 0.917 | 0.85 | 0.00111592 | 7.55E-08 |
| Striatum Ventral Region | cTBS | Cox7c | cytochrome c oxidase subunit 7C | 12867 | 0.246319387 | 1 | 0.998 | 3.05E-16 | 2.06E-20 |
| Striatum Ventral Region | cTBS | Rpl38 | ribosomal protein L38 | 67671 | 0.270207119 | 1 | 1 | 2.98E-24 | 2.02E-28 |
| Striatum Ventral Region | cTBS | Tmsb10 | thymosin, beta 10 | 19240 | 0.27410157 | 1 | 1 | 2.87E-17 | 1.94E-21 |
| Striatum Ventral Region | cTBS | Gstp1 | glutathione S-transferase, pi 1 | 14870 | 0.308468127 | 0.546 | 0.368 | 4.42E-09 | 2.99E-13 |
| Striatum Ventral Region | cTBS | Polr2l | polymerase (RNA) II (DNA directed) polypeptide L | 66491 | 0.386345439 | 0.558 | 0.341 | 4.63E-15 | 3.13E-19 |
| Striatum Ventral Region | cTBS | Rpl35 | ribosomal protein L35 | 66489 | 0.445589756 | 0.993 | 0.972 | 3.09E-33 | 2.09E-37 |
| Striatum Ventral Region | cTBS | Uba52 | ubiquitin A-52 residue ribosomal protein fusion product 1 | 22186 | 0.607126313 | 0.892 | 0.832 | 1.83E-30 | 1.24E-34 |
| Striatum Ventral Region | cTBS | Gm10076 | ribosomal protein L41 pseudogene | 100126819 | 1.120235596 | 0.866 | 0.59 | 2.62E-78 | 1.78E-82 |

**Data S3.** Differentially expressed genes (DEGs) identified in the motor (M1), and somatosensory (SS) cortex following iTBS, and cTBS. DEGs were identified using a cutoff of p.adj ≤ 0.05.

| Cortical Region | Cortical Layer | Stimulation | Gene Symbol | Gene Name | EntrezID | Avg log2FoldChange | % expressed in stimulated samples | % expressed in sham samples | p.adjust value (Bonferroni correction) | p.value |
|---|---|---|---|---|---|---|---|---|---|---|
| M1 | Layer 2/3 | iTBS | Bc1 | brain cytoplasmic RNA 1 | 100568459 | -0.930399046 | 0.468 | 0.78 | 1.53E-16 | 1.04E-20 |
| M1 | Layer 2/3 | iTBS | Nme7 | NME/NM23 family member 7 | 171567 | -0.633147575 | 0.266 | 0.461 | 0.000888303 | 6.01E-08 |
| M1 | Layer 2/3 | iTBS | Cdk5r2 | cyclin-dependent kinase 5, regulatory subunit 2 (p39) | 12570 | -0.532907036 | 0.365 | 0.688 | 7.93E-13 | 5.37E-17 |
| M1 | Layer 2/3 | iTBS | Dynll2 | dynein light chain LC8-type 2 | 68097 | -0.439269398 | 0.966 | 0.979 | 7.70E-06 | 5.21E-10 |
| M1 | Layer 2/3 | iTBS | Cplx2 | complexin 2 | 12890 | -0.436041517 | 0.792 | 0.869 | 0.000171026 | 1.16E-08 |
| M1 | Layer 2/3 | iTBS | Dnm1 | dynamin 1 | 13429 | -0.394078197 | 0.997 | 1 | 4.92E-08 | 3.33E-12 |
| M1 | Layer 2/3 | iTBS | Actb | actin, beta | 11461 | -0.392267123 | 1 | 0.993 | 2.04E-08 | 1.38E-12 |
| M1 | Layer 2/3 | iTBS | Syngr1 | synaptogyrin 1 | 20972 | -0.385827973 | 0.836 | 0.883 | 0.001770927 | 1.20E-07 |
| M1 | Layer 2/3 | iTBS | Glul | glutamate-ammonia ligase (glutamine synthetase) | 14645 | -0.368222064 | 0.727 | 0.865 | 0.003130555 | 2.12E-07 |
| M1 | Layer 2/3 | iTBS | Slc17a7 | solute carrier family 17 (sodium-dependent inorganic phosphate cotransporter), member 7 | 72961 | -0.351803462 | 1 | 1 | 0.000604668 | 4.09E-08 |
| M1 | Layer 2/3 | iTBS | Eef1a2 | eukaryotic translation elongation factor 1 alpha 2 | 13628 | -0.35158734 | 0.997 | 0.996 | 0.001908941 | 1.29E-07 |
| M1 | Layer 2/3 | iTBS | Syn1 | synapsin I | 20964 | -0.34867417 | 0.973 | 0.972 | 0.032540862 | 2.20E-06 |
| M1 | Layer 2/3 | iTBS | Epn1 | epsin 1 | 13854 | -0.344058651 | 0.874 | 0.915 | 0.002160859 | 1.46E-07 |
| M1 | Layer 2/3 | iTBS | Tuba1a | tubulin, alpha 1A | 22142 | -0.314281119 | 0.997 | 0.993 | 3.22E-07 | 2.18E-11 |
| M1 | Layer 2/3 | iTBS | Tuba1b | tubulin, alpha 1B | 22143 | -0.312792785 | 0.976 | 0.993 | 0.000152058 | 1.03E-08 |
| M1 | Layer 2/3 | iTBS | Rpl5 | ribosomal protein L5 | 100503670 | -0.305948502 | 0.922 | 0.943 | 0.001926034 | 1.30E-07 |
| M1 | Layer 2/3 | iTBS | R3hdm4 | R3H domain containing 4 | 109284 | -0.303133015 | 0.652 | 0.773 | 0.038279947 | 2.59E-06 |
| M1 | Layer 2/3 | iTBS | Syngap1 | synaptic Ras GTPase activating protein 1 homolog (rat) | 240057 | -0.286678646 | 0.215 | 0.433 | 0.000223026 | 1.51E-08 |
| M1 | Layer 2/3 | iTBS | Ikbkb | inhibitor of kappaB kinase beta | 16150 | -0.27390344 | 0.28 | 0.493 | 0.001960654 | 1.33E-07 |
| M1 | Layer 2/3 | iTBS | Ldha | lactate dehydrogenase A | 16828 | -0.265521606 | 0.908 | 0.943 | 0.014357214 | 9.72E-07 |
| M1 | Layer 2/3 | iTBS | Atp6v0e2 | ATPase, H+ transporting, lysosomal V0 subunit E2 | 76252 | -0.26004892 | 0.887 | 0.95 | 0.0048681 | 3.29E-07 |
| M1 | Layer 2/3 | iTBS | Dctn4 | dynactin 4 | 67665 | -0.237092875 | 0.283 | 0.475 | 0.024083992 | 1.63E-06 |
| M1 | Layer 2/3 | iTBS | Fosl2 | fos-like antigen 2 | 14284 | -0.225587125 | 0.181 | 0.351 | 0.045901191 | 3.11E-06 |
| M1 | Layer 2/3 | iTBS | Lrrc17 | leucine rich repeat containing 17 | 74511 | -0.194317382 | 0.109 | 0.259 | 0.047829164 | 3.24E-06 |
| M1 | Layer 2/3 | iTBS | Calm1 | calmodulin 1 | 12313 | -0.186409037 | 1 | 1 | 2.13E-05 | 1.44E-09 |
| M1 | Layer 2/3 | iTBS | Rpl41 | ribosomal protein L41 | 67945 | -0.164504145 | 1 | 1 | 0.002231487 | 1.51E-07 |
| M1 | Layer 2/3 | iTBS | Gnas | GNAS (guanine nucleotide binding protein, alpha stimulating) complex locus | 14683 | 0.156001223 | 1 | 1 | 0.040515152 | 2.74E-06 |
| M1 | Layer 2/3 | iTBS | Cox8a | cytochrome c oxidase subunit 8A | 12868 | 0.161527293 | 1 | 1 | 0.019583383 | 1.33E-06 |
| M1 | Layer 2/3 | iTBS | Atp1b1 | ATPase, Na+/K+ transporting, beta 1 polypeptide | 11931 | 0.174638194 | 1 | 1 | 0.023429173 | 1.59E-06 |
| M1 | Layer 2/3 | iTBS | Cox6c | cytochrome c oxidase subunit 6C | 12864 | 0.194249754 | 1 | 1 | 1.73E-06 | 1.17E-10 |
| M1 | Layer 2/3 | iTBS | Fth1 | ferritin heavy polypeptide 1 | 14319 | 0.197613997 | 1 | 1 | 3.04E-09 | 2.06E-13 |
| M1 | Layer 2/3 | iTBS | Rpl37 | ribosomal protein L37 | 67281 | 0.20874891 | 1 | 1 | 0.00635227 | 4.30E-07 |
| M1 | Layer 2/3 | iTBS | Rpl24 | ribosomal protein L24 | 68193 | 0.21520329 | 0.993 | 0.975 | 0.013278223 | 8.99E-07 |
| M1 | Layer 2/3 | iTBS | Rpl21 | ribosomal protein L21 | 19933 | 0.220984135 | 1 | 0.996 | 5.42E-05 | 3.67E-09 |
| M1 | Layer 2/3 | iTBS | Rpl11 | ribosomal protein L11 | 67025 | 0.221751834 | 0.997 | 0.979 | 0.001588468 | 1.07E-07 |
| M1 | Layer 2/3 | iTBS | Tmsb4x | thymosin, beta 4, X chromosome | 19241 | 0.224602885 | 1 | 1 | 5.33E-08 | 3.60E-12 |
| M1 | Layer 2/3 | iTBS | Rpl13 | ribosomal protein L13 | 270106 | 0.233635205 | 1 | 1 | 4.70E-06 | 3.18E-10 |
| M1 | Layer 2/3 | iTBS | Cox7c | cytochrome c oxidase subunit 7C | 12867 | 0.234243641 | 1 | 1 | 0.000221573 | 1.50E-08 |
| M1 | Layer 2/3 | iTBS | Rps7 | ribosomal protein S7 | 20115 | 0.243048301 | 0.997 | 0.979 | 0.007034415 | 4.76E-07 |
| M1 | Layer 2/3 | iTBS | Rps24 | ribosomal protein S24 | 20088 | 0.252146286 | 1 | 0.993 | 2.13E-05 | 1.44E-09 |
| M1 | Layer 2/3 | iTBS | Rps10 | ribosomal protein S10 | 67097 | 0.256160549 | 0.99 | 0.94 | 0.004997984 | 3.38E-07 |
| M1 | Layer 2/3 | iTBS | Rps27a | ribosomal protein S27A | 78294 | 0.262082632 | 1 | 0.986 | 0.001235155 | 8.36E-08 |
| M1 | Layer 2/3 | iTBS | Rpl30 | ribosomal protein L30 | 19946 | 0.266937423 | 0.993 | 0.936 | 0.00383604 | 2.60E-07 |
| M1 | Layer 2/3 | iTBS | Rpl23 | ribosomal protein L23 | 65019 | 0.273637269 | 1 | 0.975 | 3.09E-05 | 2.09E-09 |
| M1 | Layer 2/3 | iTBS | Rps21 | ribosomal protein S21 | 66481 | 0.281924237 | 1 | 0.996 | 5.16E-07 | 3.49E-11 |
| M1 | Layer 2/3 | iTBS | Rps8 | ribosomal protein S8 | 20116 | 0.289422095 | 0.99 | 0.982 | 9.05E-06 | 6.13E-10 |
| M1 | Layer 2/3 | iTBS | Junb | jun B proto-oncogene | 16477 | 0.298010412 | 0.659 | 0.457 | 0.001060932 | 7.18E-08 |
| M1 | Layer 2/3 | iTBS | Rpl9 | ribosomal protein L9 | 20005 | 0.323306121 | 1 | 0.979 | 2.01E-08 | 1.36E-12 |
| M1 | Layer 2/3 | iTBS | Nr4a1 | nuclear receptor subfamily 4, group A, member 1 | 15370 | 0.325322467 | 0.761 | 0.631 | 0.004682657 | 3.17E-07 |
| M1 | Layer 2/3 | iTBS | Rpl35 | ribosomal protein L35 | 66489 | 0.428949855 | 0.969 | 0.851 | 2.05E-08 | 1.39E-12 |
| M1 | Layer 2/3 | iTBS | Gstp1 | glutathione S-transferase, pi 1 | 14870 | 0.433633298 | 0.515 | 0.248 | 2.29E-08 | 1.55E-12 |
| M1 | Layer 2/3 | iTBS | Uba52 | ubiquitin A-52 residue ribosomal protein fusion product 1 | 22186 | 0.996141378 | 0.86 | 0.592 | 3.20E-29 | 2.17E-33 |
| M1 | Layer 2/3 | iTBS | Gm10076 | ribosomal protein L41 pseudogene | 100126819 | 1.454574365 | 0.819 | 0.301 | 2.76E-46 | 1.87E-50 |
| M1 | Layer 5 | iTBS | Bc1 | brain cytoplasmic RNA 1 | 100568459 | -0.556184087 | 0.49 | 0.73 | 1.43E-16 | 9.67E-21 |
| M1 | Layer 5 | iTBS | Cplx2 | complexin 2 | 12890 | -0.547573969 | 0.645 | 0.762 | 5.93E-12 | 4.01E-16 |
| M1 | Layer 5 | iTBS | Dynll2 | dynein light chain LC8-type 2 | 68097 | -0.361690156 | 0.959 | 0.977 | 1.27E-07 | 8.62E-12 |
| M1 | Layer 5 | iTBS | Cdk5r2 | cyclin-dependent kinase 5, regulatory subunit 2 (p39) | 12570 | -0.332047506 | 0.402 | 0.587 | 3.69E-08 | 2.50E-12 |
| M1 | Layer 5 | iTBS | Hpcal4 | hippocalcin-like 4 | 170638 | -0.326978757 | 0.656 | 0.805 | 5.63E-05 | 3.81E-09 |
| M1 | Layer 5 | iTBS | Eef1a2 | eukaryotic translation elongation factor 1 alpha 2 | 13628 | -0.313475648 | 1 | 0.998 | 4.66E-08 | 3.15E-12 |
| M1 | Layer 5 | iTBS | Dnm1 | dynamin 1 | 13429 | -0.309624637 | 0.998 | 1 | 5.74E-09 | 3.88E-13 |
| M1 | Layer 5 | iTBS | Tuba1b | tubulin, alpha 1B | 22143 | -0.304908539 | 0.967 | 0.987 | 8.58E-12 | 5.80E-16 |
| M1 | Layer 5 | iTBS | Glul | glutamate-ammonia ligase (glutamine synthetase) | 14645 | -0.293099767 | 0.815 | 0.872 | 5.70E-05 | 3.85E-09 |
| M1 | Layer 5 | iTBS | Stx1b | syntaxin 1B | 56216 | -0.293057921 | 0.828 | 0.874 | 4.40E-05 | 2.98E-09 |
| M1 | Layer 5 | iTBS | Ndrg2 | N-myc downstream regulated gene 2 | 29811 | -0.289289769 | 0.875 | 0.917 | 9.89E-05 | 6.70E-09 |
| M1 | Layer 5 | iTBS | Sptbn2 | spectrin beta, non-erythrocytic 2 | 20743 | -0.28839933 | 0.809 | 0.865 | 0.009945331 | 6.73E-07 |
| M1 | Layer 5 | iTBS | Actb | actin, beta | 11461 | -0.286079493 | 0.994 | 0.994 | 1.22E-06 | 8.25E-11 |
| M1 | Layer 5 | iTBS | Phyhip | phytanoyl-CoA hydroxylase interacting protein | 105653 | -0.284007447 | 0.427 | 0.583 | 2.62E-05 | 1.77E-09 |
| M1 | Layer 5 | iTBS | Kcnq2 | potassium voltage-gated channel, subfamily Q, member 2 | 16536 | -0.268063584 | 0.573 | 0.692 | 0.003602513 | 2.44E-07 |
| M1 | Layer 5 | iTBS | Slc17a7 | solute carrier family 17 (sodium-dependent inorganic phosphate cotransporter), member 7 | 72961 | -0.267703515 | 0.996 | 0.996 | 0.003462609 | 2.34E-07 |
| M1 | Layer 5 | iTBS | Rpl5 | ribosomal protein L5 | 100503670 | -0.264821637 | 0.923 | 0.962 | 6.73E-06 | 4.55E-10 |
| M1 | Layer 5 | iTBS | Ivns1abp | influenza virus NS1A binding protein | 117198 | -0.26005929 | 0.456 | 0.589 | 0.014667245 | 9.93E-07 |
| M1 | Layer 5 | iTBS | Epn1 | epsin 1 | 13854 | -0.257900631 | 0.846 | 0.893 | 0.009364102 | 6.34E-07 |
| M1 | Layer 5 | iTBS | Emc10 | ER membrane protein complex subunit 10 | 69683 | -0.253231918 | 0.815 | 0.863 | 0.000401949 | 2.72E-08 |
| M1 | Layer 5 | iTBS | Dusp26 | dual specificity phosphatase 26 (putative) | 66959 | -0.252448117 | 0.388 | 0.559 | 5.24E-05 | 3.54E-09 |
| M1 | Layer 5 | iTBS | Eno2 | enolase 2, gamma neuronal | 13807 | -0.247646972 | 0.969 | 0.983 | 0.000173344 | 1.17E-08 |
| M1 | Layer 5 | iTBS | Thy1 | thymus cell antigen 1, theta | 21838 | -0.243059102 | 0.969 | 0.996 | 1.46E-06 | 9.88E-11 |
| M1 | Layer 5 | iTBS | Brsk1 | BR serine/threonine kinase 1 | 381979 | -0.241335178 | 0.454 | 0.595 | 0.001171609 | 7.93E-08 |
| M1 | Layer 5 | iTBS | Add2 | adducin 2 (beta) | 11519 | -0.23948343 | 0.38 | 0.523 | 0.00162507 | 1.10E-07 |
| M1 | Layer 5 | iTBS | Ikbkb | inhibitor of kappaB kinase beta | 16150 | -0.236972258 | 0.292 | 0.445 | 0.000435446 | 2.95E-08 |
| M1 | Layer 5 | iTBS | Atp6v0a1 | ATPase, H+ transporting, lysosomal V0 subunit A1 | 11975 | -0.236710063 | 0.77 | 0.842 | 0.035703825 | 2.42E-06 |
| M1 | Layer 5 | iTBS | Lars2 | leucyl-tRNA synthetase, mitochondrial | 102436 | -0.233238855 | 0.257 | 0.426 | 3.95E-05 | 2.67E-09 |
| M1 | Layer 5 | iTBS | Pde1b | phosphodiesterase 1B, Ca2+-calmodulin dependent | 18574 | -0.230292745 | 0.313 | 0.452 | 0.008075804 | 5.46E-07 |
| M1 | Layer 5 | iTBS | Mt2 | metallothionein 2 | 17750 | -0.229777887 | 0.434 | 0.57 | 0.026209186 | 1.77E-06 |
| M1 | Layer 5 | iTBS | Tuba1a | tubulin, alpha 1A | 22142 | -0.229620706 | 0.983 | 0.998 | 4.60E-06 | 3.12E-10 |
| M1 | Layer 5 | iTBS | Rpl4 | ribosomal protein L4 | 67891 | -0.207869565 | 0.99 | 0.994 | 0.000111034 | 7.51E-09 |
| M1 | Layer 5 | iTBS | Reep5 | receptor accessory protein 5 | 13476 | -0.206299967 | 0.859 | 0.906 | 0.044111617 | 2.98E-06 |
| M1 | Layer 5 | iTBS | Dync1h1 | dynein cytoplasmic 1 heavy chain 1 | 13424 | -0.203017145 | 0.884 | 0.916 | 0.043027686 | 2.91E-06 |
| M1 | Layer 5 | iTBS | Agap2 | ArfGAP with GTPase domain, ankyrin repeat and PH domain 2 | 216439 | -0.202242686 | 0.216 | 0.37 | 0.000391815 | 2.65E-08 |
| M1 | Layer 5 | iTBS | Eef2 | eukaryotic translation elongation factor 2 | 13629 | -0.199167071 | 0.983 | 0.987 | 0.028233603 | 1.91E-06 |
| M1 | Layer 5 | iTBS | Gnb1 | guanine nucleotide binding protein (G protein), beta 1 | 14688 | -0.189714784 | 0.958 | 0.981 | 0.001067638 | 7.22E-08 |
| M1 | Layer 5 | iTBS | Clu | clusterin | 12759 | -0.187903639 | 0.961 | 0.981 | 0.030902846 | 2.09E-06 |
| M1 | Layer 5 | iTBS | Ppp1r1b | protein phosphatase 1, regulatory inhibitor subunit 1B | 19049 | -0.164769876 | 0.108 | 0.216 | 0.027527854 | 1.86E-06 |
| M1 | Layer 5 | iTBS | Aldoa | aldolase A, fructose-bisphosphate | 11674 | -0.144867399 | 1 | 1 | 0.000269084 | 1.82E-08 |
| M1 | Layer 5 | iTBS | Cpe | carboxypeptidase E | 12876 | -0.140047726 | 1 | 1 | 0.005531318 | 3.74E-07 |
| M1 | Layer 5 | iTBS | Hspa8 | heat shock protein 8 | 15481 | -0.115305427 | 1 | 1 | 0.03114526 | 2.11E-06 |
| M1 | Layer 5 | iTBS | Rpl41 | ribosomal protein L41 | 67945 | -0.114442923 | 1 | 1 | 0.018975084 | 1.28E-06 |
| M1 | Layer 5 | iTBS | Lhpp | phospholysine phosphohistidine inorganic pyrophosphate phosphatase | 76429 | 0.125199691 | 0.153 | 0.064 | 0.045745145 | 3.10E-06 |
| M1 | Layer 5 | iTBS | Calm2 | calmodulin 2 | 12314 | 0.13354441 | 1 | 1 | 0.000733871 | 4.97E-08 |
| M1 | Layer 5 | iTBS | Acyp2 | acylphosphatase 2, muscle type | 75572 | 0.142744163 | 0.27 | 0.152 | 0.041372707 | 2.80E-06 |
| M1 | Layer 5 | iTBS | Ppia | peptidylprolyl isomerase A | 268373 | 0.148260658 | 1 | 1 | 6.35E-05 | 4.29E-09 |
| M1 | Layer 5 | iTBS | Gnas | GNAS (guanine nucleotide binding protein, alpha stimulating) complex locus | 14683 | 0.148758834 | 1 | 1 | 0.00026691 | 1.81E-08 |
| M1 | Layer 5 | iTBS | Rpl11 | ribosomal protein L11 | 67025 | 0.149550676 | 1 | 0.989 | 0.012046139 | 8.15E-07 |
| M1 | Layer 5 | iTBS | Fth1 | ferritin heavy polypeptide 1 | 14319 | 0.153169848 | 1 | 1 | 6.19E-06 | 4.19E-10 |
| M1 | Layer 5 | iTBS | Atp1b1 | ATPase, Na+/K+ transporting, beta 1 polypeptide | 11931 | 0.15400753 | 1 | 1 | 0.0328506 | 2.22E-06 |
| M1 | Layer 5 | iTBS | Cox8a | cytochrome c oxidase subunit 8A | 12868 | 0.154790004 | 1 | 1 | 1.27E-05 | 8.56E-10 |
| M1 | Layer 5 | iTBS | Rpl21 | ribosomal protein L21 | 19933 | 0.162458094 | 1 | 1 | 1.86E-06 | 1.26E-10 |
| M1 | Layer 5 | iTBS | Tmsb4x | thymosin, beta 4, X chromosome | 19241 | 0.163983452 | 1 | 1 | 5.56E-07 | 3.76E-11 |
| M1 | Layer 5 | iTBS | Rpl37 | ribosomal protein L37 | 67281 | 0.165846849 | 1 | 1 | 0.00056436 | 3.82E-08 |
| M1 | Layer 5 | iTBS | Rtn1 | reticulon 1 | 104001 | 0.168887932 | 0.998 | 0.994 | 0.003120554 | 2.11E-07 |
| M1 | Layer 5 | iTBS | Ifit3 | interferon-induced protein with tetratricopeptide repeats 3 | 15959 | 0.171970043 | 0.124 | 0.024 | 9.10E-06 | 6.16E-10 |
| M1 | Layer 5 | iTBS | Rps7 | ribosomal protein S7 | 20115 | 0.172569875 | 0.992 | 0.977 | 0.001613082 | 1.09E-07 |
| M1 | Layer 5 | iTBS | Cox6c | cytochrome c oxidase subunit 6C | 12864 | 0.176906952 | 1 | 1 | 1.03E-07 | 6.98E-12 |
| M1 | Layer 5 | iTBS | Selenok | selenoprotein K | 80795 | 0.179591186 | 0.9 | 0.786 | 0.010586322 | 7.16E-07 |
| M1 | Layer 5 | iTBS | Rps27a | ribosomal protein S27A | 78294 | 0.180190549 | 1 | 0.998 | 0.000186452 | 1.26E-08 |
| M1 | Layer 5 | iTBS | Rpl36a | ribosomal protein L36A | 19982 | 0.18125957 | 0.898 | 0.824 | 0.009950042 | 6.73E-07 |
| M1 | Layer 5 | iTBS | Rpl13 | ribosomal protein L13 | 270106 | 0.187210724 | 1 | 0.998 | 7.40E-07 | 5.01E-11 |
| M1 | Layer 5 | iTBS | Dusp1 | dual specificity phosphatase 1 | 19252 | 0.190441152 | 0.523 | 0.381 | 0.025391256 | 1.72E-06 |
| M1 | Layer 5 | iTBS | Rps20 | ribosomal protein S20 | 67427 | 0.19284623 | 0.996 | 0.998 | 2.94E-05 | 1.99E-09 |
| M1 | Layer 5 | iTBS | Efhd2 | EF hand domain containing 2 | 27984 | 0.197545986 | 0.89 | 0.863 | 0.038633185 | 2.61E-06 |
| M1 | Layer 5 | iTBS | Ndufb9 | NADH:ubiquinone oxidoreductase subunit B9 | 66218 | 0.201520228 | 0.996 | 0.977 | 1.08E-05 | 7.29E-10 |
| M1 | Layer 5 | iTBS | Chd9 | chromodomain helicase DNA binding protein 9 | 109151 | 0.205439686 | 0.394 | 0.238 | 0.000557506 | 3.77E-08 |
| M1 | Layer 5 | iTBS | Rpl35a | ribosomal protein L35A | 57808 | 0.207090584 | 0.994 | 0.977 | 0.001319519 | 8.93E-08 |
| M1 | Layer 5 | iTBS | Rps21 | ribosomal protein S21 | 66481 | 0.210037112 | 1 | 0.996 | 6.99E-07 | 4.73E-11 |
| M1 | Layer 5 | iTBS | Eno1 | enolase 1, alpha non-neuron | 13806 | 0.236439726 | 0.965 | 0.904 | 0.000837028 | 5.66E-08 |
| M1 | Layer 5 | iTBS | Cox7c | cytochrome c oxidase subunit 7C | 12867 | 0.251816906 | 1 | 0.998 | 1.95E-11 | 1.32E-15 |
| M1 | Layer 5 | iTBS | Tmsb10 | thymosin, beta 10 | 19240 | 0.266888129 | 0.998 | 0.996 | 6.54E-10 | 4.42E-14 |
| M1 | Layer 5 | iTBS | Rpl9 | ribosomal protein L9 | 20005 | 0.28607227 | 0.998 | 0.983 | 1.75E-13 | 1.18E-17 |
| M1 | Layer 5 | iTBS | Pvalb | parvalbumin | 19293 | 0.29439878 | 0.838 | 0.762 | 0.021682193 | 1.47E-06 |
| M1 | Layer 5 | iTBS | Rps8 | ribosomal protein S8 | 20116 | 0.298601597 | 1 | 0.994 | 7.97E-15 | 5.39E-19 |
| M1 | Layer 5 | iTBS | Rpl35 | ribosomal protein L35 | 66489 | 0.366759307 | 0.981 | 0.901 | 3.65E-12 | 2.47E-16 |
| M1 | Layer 5 | iTBS | Gstp1 | glutathione S-transferase, pi 1 | 14870 | 0.389247318 | 0.537 | 0.31 | 2.86E-12 | 1.94E-16 |
| M1 | Layer 5 | iTBS | Uba52 | ubiquitin A-52 residue ribosomal protein fusion product 1 | 22186 | 0.779851038 | 0.861 | 0.664 | 5.64E-35 | 3.82E-39 |
| M1 | Layer 5 | iTBS | Gm10076 | ribosomal protein L41 pseudogene | 100126819 | 1.28664674 | 0.764 | 0.321 | 6.03E-70 | 4.08E-74 |
| M1 | Layer 6 | iTBS | Eef1a2 | eukaryotic translation elongation factor 1 alpha 2 | 13628 | -0.507722247 | 0.979 | 0.995 | 2.37E-05 | 1.60E-09 |
| M1 | Layer 6 | iTBS | Ppia | peptidylprolyl isomerase A | 268373 | 0.214189038 | 1 | 1 | 0.004501071 | 3.05E-07 |
| M1 | Layer 6 | iTBS | Cox6c | cytochrome c oxidase subunit 6C | 12864 | 0.222728066 | 1 | 1 | 0.001136761 | 7.69E-08 |

| | | | | | | | | | | |
|---|---|---|---|---|---|---|---|---|---|---|
| M1 | Layer 6 | iTBS | Cox5b | cytochrome c oxidase subunit 5B | 12859 | 0.240869199 | 1 | 1 | 0.007105391 | 4.81E-07 |
| M1 | Layer 6 | iTBS | Calm2 | calmodulin 2 | 12314 | 0.248976302 | 1 | 1 | 0.015082718 | 1.02E-06 |
| M1 | Layer 6 | iTBS | Rpl38 | ribosomal protein L38 | 67671 | 0.261461166 | 1 | 1 | 0.002561907 | 1.73E-07 |
| M1 | Layer 6 | iTBS | Ndufb9 | NADH:ubiquinone oxidoreductase subunit B9 | 66218 | 0.280285928 | 1 | 0.984 | 0.039632005 | 2.68E-06 |
| M1 | Layer 6 | iTBS | Rps8 | ribosomal protein S8 | 20116 | 0.293572287 | 0.995 | 0.995 | 0.00213008 | 1.44E-07 |
| M1 | Layer 6 | iTBS | Sytl2 | synaptotagmin-like 2 | 83671 | 0.31277338 | 0.351 | 0.141 | 0.017684192 | 1.20E-06 |
| M1 | Layer 6 | iTBS | Rpl35 | ribosomal protein L35 | 66489 | 0.337141937 | 0.995 | 0.979 | 0.001271736 | 8.61E-08 |
| M1 | Layer 6 | iTBS | Gstp1 | glutathione S-transferase, pi 1 | 14870 | 0.644680111 | 0.644 | 0.344 | 1.63E-07 | 1.10E-11 |
| M1 | Layer 6 | iTBS | Uba52 | ubiquitin A-52 residue ribosomal protein fusion product 1 | 22186 | 0.774490163 | 0.91 | 0.776 | 5.78E-12 | 3.91E-16 |
| M1 | Layer 6 | iTBS | Gm10076 | ribosomal protein L41 pseudogene | 100126819 | 1.224857727 | 0.793 | 0.49 | 4.21E-18 | 2.85E-22 |
| M1 | Layer 2/3 | cTBS | Bc1 | brain cytoplasmic RNA 1 | 100568459 | -1.093216524 | 0.382 | 0.78 | 1.21E-20 | 8.18E-25 |
| M1 | Layer 2/3 | cTBS | Ttr | transthyretin | 22139 | -0.737194838 | 0.013 | 0.234 | 1.61E-09 | 1.09E-13 |
| M1 | Layer 2/3 | cTBS | Cdk5r2 | cyclin-dependent kinase 5, regulatory subunit 2 (p39) | 12570 | -0.348726088 | 0.508 | 0.688 | 0.007100877 | 4.81E-07 |
| M1 | Layer 2/3 | cTBS | Tatdn1 | TatD DNase domain containing 1 | 69694 | -0.309512891 | 0.126 | 0.358 | 1.18E-05 | 7.97E-10 |
| M1 | Layer 2/3 | cTBS | Basp1 | brain abundant, membrane attached signal protein 1 | 70350 | -0.306187338 | 0.807 | 0.947 | 0.002924508 | 1.98E-07 |
| M1 | Layer 2/3 | cTBS | Tuba1a | tubulin, alpha 1A | 22142 | -0.300088442 | 0.992 | 0.993 | 2.38E-05 | 1.61E-09 |
| M1 | Layer 2/3 | cTBS | Tuba1b | tubulin, alpha 1B | 22143 | -0.29944023 | 0.987 | 0.993 | 0.00228783 | 1.55E-07 |
| M1 | Layer 2/3 | cTBS | Ikbkb | inhibitor of kappaB kinase beta | 16150 | -0.267576532 | 0.277 | 0.493 | 0.009503442 | 6.43E-07 |
| M1 | Layer 2/3 | cTBS | Fam207a | family with sequence similarity 207, member A | 108707 | -0.25483742 | 0.092 | 0.287 | 0.00039857 | 2.70E-08 |
| M1 | Layer 2/3 | cTBS | Lrrc17 | leucine rich repeat containing 17 | 74511 | -0.243106445 | 0.076 | 0.259 | 0.000559054 | 3.78E-08 |
| M1 | Layer 2/3 | cTBS | Cck | cholecystokinin | 12424 | -0.213469143 | 1 | 1 | 0.001061602 | 7.18E-08 |
| M1 | Layer 2/3 | cTBS | Calm1 | calmodulin 1 | 12313 | -0.182311629 | 1 | 1 | 0.000312125 | 2.11E-08 |
| M1 | Layer 2/3 | cTBS | Rpl38 | ribosomal protein L38 | 67671 | 0.202913413 | 1 | 1 | 0.001661876 | 1.12E-07 |
| M1 | Layer 2/3 | cTBS | Uqcr11 | ubiquinol-cytochrome c reductase, complex III subunit XI | 66594 | 0.26341501 | 1 | 0.996 | 0.000354382 | 2.40E-08 |
| M1 | Layer 2/3 | cTBS | Atp5k | ATP synthase, H+ transporting, mitochondrial F1F0 complex, subunit E | 11958 | 0.287120899 | 1 | 0.989 | 9.85E-05 | 6.67E-09 |
| M1 | Layer 2/3 | cTBS | Eno1 | enolase 1, alpha non-neuron | 13806 | 0.327195749 | 0.962 | 0.894 | 0.003632396 | 2.46E-07 |
| M1 | Layer 2/3 | cTBS | Gstp1 | glutathione S-transferase, pi 1 | 14870 | 0.344800544 | 0.458 | 0.248 | 0.001363282 | 9.23E-08 |
| M1 | Layer 2/3 | cTBS | Rpl35 | ribosomal protein L35 | 66489 | 0.586394679 | 0.966 | 0.851 | 8.56E-14 | 5.79E-18 |
| M1 | Layer 2/3 | cTBS | Uba52 | ubiquitin A-52 residue ribosomal protein fusion product 1 | 22186 | 0.671484509 | 0.79 | 0.592 | 9.79E-12 | 6.63E-16 |
| M1 | Layer 2/3 | cTBS | Gm10076 | ribosomal protein L41 pseudogene | 100126819 | 1.208843802 | 0.777 | 0.301 | 1.07E-33 | 7.24E-38 |
| M1 | Layer 5 | cTBS | Bc1 | brain cytoplasmic RNA 1 | 100568459 | -0.719464524 | 0.359 | 0.73 | 1.30E-31 | 8.80E-36 |
| M1 | Layer 5 | cTBS | Ttr | transthyretin | 22139 | -0.478572331 | 0.026 | 0.216 | 8.28E-18 | 5.60E-22 |
| M1 | Layer 5 | cTBS | Ikbkb | inhibitor of kappaB kinase beta | 16150 | -0.322632072 | 0.226 | 0.445 | 5.89E-11 | 3.99E-15 |
| M1 | Layer 5 | cTBS | Atp1a3 | ATPase, Na+/K+ transporting, alpha 3 polypeptide | 232975 | -0.319363151 | 0.804 | 0.919 | 1.66E-07 | 1.12E-11 |
| M1 | Layer 5 | cTBS | Cdk5r2 | cyclin-dependent kinase 5, regulatory subunit 2 (p39) | 12570 | -0.306716563 | 0.403 | 0.587 | 3.65E-07 | 2.47E-11 |
| M1 | Layer 5 | cTBS | Basp1 | brain abundant, membrane attached signal protein 1 | 70350 | -0.281820053 | 0.804 | 0.946 | 8.32E-08 | 5.63E-12 |
| M1 | Layer 5 | cTBS | Tuba1b | tubulin, alpha 1B | 22143 | -0.248692868 | 0.982 | 0.987 | 7.27E-08 | 4.92E-12 |
| M1 | Layer 5 | cTBS | Tuba1a | tubulin, alpha 1A | 22142 | -0.240953294 | 0.989 | 0.998 | 6.54E-08 | 4.43E-12 |
| M1 | Layer 5 | cTBS | Arpc4 | actin related protein 2/3 complex, subunit 4 | 68089 | -0.239991566 | 0.527 | 0.664 | 0.003285191 | 2.22E-07 |
| M1 | Layer 5 | cTBS | Trf | transferrin | 22041 | -0.232608624 | 0.379 | 0.522 | 0.02200046 | 1.49E-06 |
| M1 | Layer 5 | cTBS | Huwe1 | HECT, UBA and WWE domain containing 1 | 59026 | -0.228818034 | 0.351 | 0.512 | 0.000256833 | 1.74E-08 |
| M1 | Layer 5 | cTBS | Phyhip | phytanoyl-CoA hydroxylase interacting protein | 105653 | -0.226607491 | 0.436 | 0.583 | 0.002388576 | 1.62E-07 |
| M1 | Layer 5 | cTBS | Tubb4a | tubulin, beta 4A class IVA | 22153 | -0.2195663 | 0.845 | 0.906 | 0.01016501 | 6.88E-07 |
| M1 | Layer 5 | cTBS | Snhg9 | small nucleolar RNA host gene 9 | 73474 | -0.217932787 | 0.129 | 0.278 | 1.21E-05 | 8.19E-10 |
| M1 | Layer 5 | cTBS | Brsk1 | BR serine/threonine kinase 1 | 381979 | -0.212463507 | 0.455 | 0.595 | 0.008287477 | 5.61E-07 |
| M1 | Layer 5 | cTBS | Rnf141 | ring finger protein 141 | 67150 | -0.201042481 | 0.205 | 0.341 | 0.003097472 | 2.10E-07 |
| M1 | Layer 5 | cTBS | Spcs2 | signal peptidase complex subunit 2 homolog (S. cerevisiae) | 66624 | -0.192693091 | 0.329 | 0.473 | 0.011924516 | 8.07E-07 |
| M1 | Layer 5 | cTBS | Ogdhl | oxoglutarate dehydrogenase-like | 239017 | -0.185221114 | 0.211 | 0.347 | 0.005023209 | 3.40E-07 |
| M1 | Layer 5 | cTBS | Agap2 | ArfGAP with GTPase domain, ankyrin repeat and PH domain 2 | 216439 | -0.179837681 | 0.214 | 0.37 | 0.001035746 | 7.01E-08 |
| M1 | Layer 5 | cTBS | Tm9sf4 | transmembrane 9 superfamily member 4 | 99237 | -0.179108921 | 0.203 | 0.33 | 0.017742338 | 1.20E-06 |
| M1 | Layer 5 | cTBS | Satb2 | special AT-rich sequence binding protein 2 | 212712 | -0.178311074 | 0.222 | 0.36 | 0.014816336 | 1.00E-06 |
| M1 | Layer 5 | cTBS | Mrpl46 | mitochondrial ribosomal protein L46 | 67308 | -0.177401165 | 0.255 | 0.411 | 0.001741992 | 1.18E-07 |
| M1 | Layer 5 | cTBS | Tatdn1 | TatD DNase domain containing 1 | 69694 | -0.174666698 | 0.131 | 0.251 | 0.005190664 | 3.51E-07 |
| M1 | Layer 5 | cTBS | Snrpn | small nuclear ribonucleoprotein N | 20646 | -0.170364311 | 0.993 | 0.998 | 0.014978059 | 1.01E-06 |
| M1 | Layer 5 | cTBS | Snrpc | U1 small nuclear ribonucleoprotein C | 20630 | -0.166254775 | 0.287 | 0.43 | 0.039617137 | 2.68E-06 |
| M1 | Layer 5 | cTBS | Dapk3 | death-associated protein kinase 3 | 13144 | -0.16588297 | 0.17 | 0.296 | 0.013920735 | 9.42E-07 |
| M1 | Layer 5 | cTBS | Gm2000 | ribosomal protein L35 pseudogene | 100038991 | -0.164618003 | 0.172 | 0.3 | 0.013379906 | 9.05E-07 |
| M1 | Layer 5 | cTBS | Arxes2 | adipocyte-related X-chromosome expressed sequence 2 | 76976 | -0.159595338 | 0.194 | 0.321 | 0.034604954 | 2.34E-06 |
| M1 | Layer 5 | cTBS | Abca3 | ATP-binding cassette, sub-family A (ABC1), member 3 | 27410 | -0.153237316 | 0.305 | 0.45 | 0.046781662 | 3.17E-06 |
| M1 | Layer 5 | cTBS | Tmem131 | transmembrane protein 131 | 56030 | -0.149817158 | 0.107 | 0.212 | 0.032777275 | 2.22E-06 |
| M1 | Layer 5 | cTBS | Srrd | SRR1 domain containing | 70118 | -0.136970278 | 0.12 | 0.233 | 0.025889537 | 1.75E-06 |
| M1 | Layer 5 | cTBS | Atp6v0c | ATPase, H+ transporting, lysosomal V0 subunit C | 11984 | -0.128913468 | 1 | 1 | 0.020457049 | 1.38E-06 |
| M1 | Layer 5 | cTBS | Vwa1 | von Willebrand factor A domain containing 1 | 246228 | -0.108080746 | 0.063 | 0.152 | 0.048344574 | 3.27E-06 |
| M1 | Layer 5 | cTBS | Ubb | ubiquitin B | 22187 | 0.119925642 | 1 | 1 | 0.017822487 | 1.21E-06 |
| M1 | Layer 5 | cTBS | Gnas | GNAS (guanine nucleotide binding protein, alpha stimulating) complex locus | 14683 | 0.139352927 | 1 | 1 | 0.034704465 | 2.35E-06 |
| M1 | Layer 5 | cTBS | Rpl37 | ribosomal protein L37 | 67281 | 0.157092073 | 1 | 1 | 0.003991945 | 2.70E-07 |
| M1 | Layer 5 | cTBS | Rpl23 | ribosomal protein L23 | 65019 | 0.164047075 | 0.998 | 0.994 | 0.005270975 | 3.57E-07 |
| M1 | Layer 5 | cTBS | Cox6c | cytochrome c oxidase subunit 6C | 12864 | 0.169505587 | 1 | 1 | 1.82E-06 | 1.23E-10 |
| M1 | Layer 5 | cTBS | Atp5k | ATP synthase, H+ transporting, mitochondrial F1F0 complex, subunit E | 11958 | 0.175772193 | 1 | 0.996 | 0.005783073 | 3.91E-07 |
| M1 | Layer 5 | cTBS | Rpl35a | ribosomal protein L35A | 57808 | 0.185588177 | 0.994 | 0.977 | 0.006611856 | 4.47E-07 |
| M1 | Layer 5 | cTBS | Rpl26 | ribosomal protein L26 | 19941 | 0.185707853 | 0.987 | 0.977 | 0.013785048 | 9.33E-07 |
| M1 | Layer 5 | cTBS | Atp5mpl | ATP synthase membrane subunit 6.8PL | 70257 | 0.186919866 | 0.998 | 0.998 | 0.00147888 | 1.00E-07 |
| M1 | Layer 5 | cTBS | Rpl31 | ribosomal protein L31 | 114641 | 0.193842014 | 0.98 | 0.962 | 0.008484951 | 5.74E-07 |
| M1 | Layer 5 | cTBS | Gas5 | growth arrest specific 5 | 14455 | 0.213537528 | 0.935 | 0.88 | 0.002772999 | 1.88E-07 |
| M1 | Layer 5 | cTBS | Atp5md | ATP synthase membrane subunit DAPIT | 66477 | 0.226176535 | 1 | 1 | 5.81E-10 | 3.93E-14 |
| M1 | Layer 5 | cTBS | Rps21 | ribosomal protein S21 | 66481 | 0.226694256 | 1 | 0.996 | 5.19E-08 | 3.51E-12 |
| M1 | Layer 5 | cTBS | Rpl38 | ribosomal protein L38 | 67671 | 0.233839864 | 1 | 1 | 5.84E-13 | 3.95E-17 |
| M1 | Layer 5 | cTBS | Rps28 | ribosomal protein S28 | 54127 | 0.23759416 | 0.9 | 0.842 | 0.008353489 | 5.65E-07 |
| M1 | Layer 5 | cTBS | Cox7c | cytochrome c oxidase subunit 7C | 12867 | 0.255584577 | 0.998 | 0.998 | 5.10E-11 | 3.45E-15 |
| M1 | Layer 5 | cTBS | Rpl37a | ribosomal protein L37a | 19981 | 0.255790516 | 0.994 | 0.972 | 5.59E-06 | 3.78E-10 |
| M1 | Layer 5 | cTBS | Eno1 | enolase 1, alpha non-neuron | 13806 | 0.281712608 | 0.963 | 0.904 | 3.11E-06 | 2.11E-10 |
| M1 | Layer 5 | cTBS | Rps27 | ribosomal protein S27 | 57294 | 0.287594688 | 0.965 | 0.878 | 1.61E-07 | 1.09E-11 |
| M1 | Layer 5 | cTBS | Rps29 | ribosomal protein S29 | 20090 | 0.304107573 | 0.996 | 0.983 | 3.60E-13 | 2.43E-17 |
| M1 | Layer 5 | cTBS | Gstp1 | glutathione S-transferase, pi 1 | 14870 | 0.341951511 | 0.477 | 0.31 | 1.18E-06 | 7.99E-11 |
| M1 | Layer 5 | cTBS | Uba52 | ubiquitin A-52 residue ribosomal protein fusion product 1 | 22186 | 0.621487028 | 0.828 | 0.664 | 1.45E-24 | 9.79E-29 |
| M1 | Layer 5 | cTBS | Rpl35 | ribosomal protein L35 | 66489 | 0.629817196 | 0.987 | 0.901 | 4.94E-41 | 3.34E-45 |
| M1 | Layer 5 | cTBS | Gm10076 | ribosomal protein L41 pseudogene | 100126819 | 1.19205091 | 0.774 | 0.321 | 2.29E-69 | 1.55E-73 |
| M1 | Layer 6 | cTBS | Bc1 | brain cytoplasmic RNA 1 | 100568459 | -0.667902783 | 0.335 | 0.688 | 3.32E-09 | 2.24E-13 |
| M1 | Layer 6 | cTBS | Ttr | transthyretin | 22139 | -0.410667093 | 0.014 | 0.177 | 9.96E-05 | 6.74E-09 |
| M1 | Layer 6 | cTBS | Cox8a | cytochrome c oxidase subunit 8A | 12868 | 0.235235903 | 1 | 1 | 0.042436011 | 2.87E-06 |
| M1 | Layer 6 | cTBS | Fth1 | ferritin heavy polypeptide 1 | 14319 | 0.242242604 | 1 | 1 | 0.000538103 | 3.64E-08 |
| M1 | Layer 6 | cTBS | Cox6c | cytochrome c oxidase subunit 6C | 12864 | 0.257019884 | 1 | 1 | 2.94E-05 | 1.99E-09 |
| M1 | Layer 6 | cTBS | Atp5md | ATP synthase membrane subunit DAPIT | 66477 | 0.267543112 | 1 | 1 | 0.000177199 | 1.20E-08 |
| M1 | Layer 6 | cTBS | Rpl38 | ribosomal protein L38 | 67671 | 0.286612031 | 1 | 1 | 2.36E-05 | 1.60E-09 |
| M1 | Layer 6 | cTBS | Pcp4 | Purkinje cell protein 4 | 18546 | 0.325715802 | 1 | 1 | 0.011314731 | 7.66E-07 |
| M1 | Layer 6 | cTBS | Ftl1 | ferritin light polypeptide 1 | 14325 | 0.332260911 | 0.991 | 0.964 | 0.013163305 | 8.91E-07 |
| M1 | Layer 6 | cTBS | Rpl35 | ribosomal protein L35 | 66489 | 0.484942586 | 1 | 0.979 | 4.17E-08 | 2.82E-12 |
| M1 | Layer 6 | cTBS | Uba52 | ubiquitin A-52 residue ribosomal protein fusion product 1 | 22186 | 0.562418974 | 0.824 | 0.776 | 0.000970955 | 6.57E-08 |
| M1 | Layer 6 | cTBS | Gm10076 | ribosomal protein L41 pseudogene | 100126819 | 0.822078401 | 0.719 | 0.49 | 4.71E-10 | 3.19E-14 |
| SS | Layer 2/3 | iTBS | Bc1 | brain cytoplasmic RNA 1 | 100568459 | -0.680503187 | 0.551 | 0.726 | 3.47E-06 | 2.35E-10 |
| SS | Layer 2/3 | iTBS | Malat1 | metastasis associated lung adenocarcinoma transcript 1 (non-coding RNA) | 72289 | -0.420450595 | 0.508 | 0.627 | 0.033937586 | 2.30E-06 |
| SS | Layer 2/3 | iTBS | Nme7 | NME/NM23 family member 7 | 171567 | -0.401414215 | 0.195 | 0.363 | 0.030227141 | 2.05E-06 |
| SS | Layer 2/3 | iTBS | Calm1 | calmodulin 1 | 12313 | -0.212412049 | 1 | 1 | 6.81E-09 | 4.61E-13 |
| SS | Layer 2/3 | iTBS | Dele1 | DAP3 binding cell death enhancer 1 | 66839 | 0.194303666 | 0.215 | 0.076 | 0.01577846 | 1.07E-06 |
| SS | Layer 2/3 | iTBS | Rpl21 | ribosomal protein L21 | 19933 | 0.208596149 | 1 | 1 | 0.000145878 | 9.87E-09 |
| SS | Layer 2/3 | iTBS | Atp1a1 | ATPase, Na+/K+ transporting, alpha 1 polypeptide | 11928 | 0.221337962 | 1 | 1 | 0.037275638 | 2.52E-06 |
| SS | Layer 2/3 | iTBS | Gtdc1 | glycosyltransferase-like domain containing 1 | 227835 | 0.24961389 | 0.33 | 0.165 | 0.023433461 | 1.59E-06 |
| SS | Layer 2/3 | iTBS | Psd | pleckstrin and Sec7 domain containing | 73728 | 0.258455079 | 0.967 | 0.842 | 0.027841205 | 1.88E-06 |
| SS | Layer 2/3 | iTBS | Rpl35 | ribosomal protein L35 | 66489 | 0.278652367 | 0.974 | 0.947 | 0.002592361 | 1.75E-07 |
| SS | Layer 2/3 | iTBS | Bsg | basigin | 12215 | 0.287372453 | 0.977 | 0.931 | 0.000262858 | 1.78E-08 |
| SS | Layer 2/3 | iTBS | Adcy1 | adenylate cyclase 1 | 432530 | 0.320730341 | 0.825 | 0.726 | 0.001107199 | 7.49E-08 |
| SS | Layer 2/3 | iTBS | Gstp1 | glutathione S-transferase, pi 1 | 14870 | 0.362570079 | 0.446 | 0.244 | 0.000247185 | 1.67E-08 |
| SS | Layer 2/3 | iTBS | Uba52 | ubiquitin A-52 residue ribosomal protein fusion product 1 | 22186 | 0.710738094 | 0.848 | 0.64 | 5.15E-16 | 3.49E-20 |
| SS | Layer 2/3 | iTBS | Gm10076 | ribosomal protein L41 pseudogene | 100126819 | 1.068215366 | 0.756 | 0.419 | 1.36E-25 | 9.20E-30 |
| SS | Layer 4 | iTBS | Malat1 | metastasis associated lung adenocarcinoma transcript 1 (non-coding RNA) | 72289 | -0.575145043 | 0.466 | 0.681 | 1.05E-08 | 7.12E-13 |
| SS | Layer 4 | iTBS | Bc1 | brain cytoplasmic RNA 1 | 100568459 | -0.418345608 | 0.355 | 0.563 | 0.001108452 | 7.50E-08 |
| SS | Layer 4 | iTBS | Tuba1b | tubulin, alpha 1B | 22143 | -0.297419376 | 0.938 | 0.987 | 0.000397833 | 2.69E-08 |
| SS | Layer 4 | iTBS | Ndufa4 | Ndufa4, mitochondrial complex associated | 17992 | -0.266486561 | 1 | 1 | 0.003414701 | 2.31E-07 |
| SS | Layer 4 | iTBS | Snhg9 | small nucleolar RNA host gene 9 | 73474 | -0.264506064 | 0.098 | 0.271 | 0.000302944 | 2.05E-08 |
| SS | Layer 4 | iTBS | Tuba1a | tubulin, alpha 1A | 22142 | -0.2454036 | 0.99 | 0.983 | 0.003586035 | 2.43E-07 |
| SS | Layer 4 | iTBS | Fau | Finkel-Biskis-Reilly murine sarcoma virus (FBR-MuSV) ubiquitously expressed (fox derived) | 14109 | 0.210565538 | 1 | 1 | 0.008676153 | 5.87E-07 |
| SS | Layer 4 | iTBS | Rps21 | ribosomal protein S21 | 66481 | 0.217634338 | 1 | 1 | 0.001070903 | 7.25E-08 |
| SS | Layer 4 | iTBS | Camk2n1 | calcium/calmodulin-dependent protein kinase II inhibitor 1 | 66259 | 0.265511261 | 1 | 1 | 1.96E-06 | 1.32E-10 |
| SS | Layer 4 | iTBS | Atp1b1 | ATPase, Na+/K+ transporting, beta 1 polypeptide | 11931 | 0.277359798 | 1 | 1 | 1.43E-05 | 9.66E-10 |
| SS | Layer 4 | iTBS | Gstp1 | glutathione S-transferase, pi 1 | 14870 | 0.384704024 | 0.435 | 0.188 | 1.29E-05 | 8.70E-10 |
| SS | Layer 4 | iTBS | Rpl35 | ribosomal protein L35 | 66489 | 0.431538062 | 0.987 | 0.943 | 6.57E-11 | 4.45E-15 |
| SS | Layer 4 | iTBS | Uba52 | ubiquitin A-52 residue ribosomal protein fusion product 1 | 22186 | 0.794795851 | 0.909 | 0.721 | 2.43E-20 | 1.65E-24 |
| SS | Layer 4 | iTBS | Gm10076 | ribosomal protein L41 pseudogene | 100126819 | 1.192706911 | 0.806 | 0.393 | 2.58E-32 | 1.74E-36 |
| SS | Layer 5 | iTBS | Bc1 | brain cytoplasmic RNA 1 | 100568459 | -0.436738846 | 0.462 | 0.598 | 0.025236626 | 1.71E-06 |
| SS | Layer 5 | iTBS | Cdk5r2 | cyclin-dependent kinase 5, regulatory subunit 2 (p39) | 12570 | -0.279423366 | 0.294 | 0.481 | 0.002802952 | 1.90E-07 |
| SS | Layer 5 | iTBS | Aldoc | aldolase C, fructose-bisphosphate | 11676 | -0.241461448 | 1 | 1 | 0.000945711 | 6.40E-08 |
| SS | Layer 5 | iTBS | Rpl41 | ribosomal protein L41 | 67945 | -0.178229734 | 1 | 1 | 0.000275532 | 1.86E-08 |
| SS | Layer 5 | iTBS | Ppp2cb | protein phosphatase 2 (formerly 2A), catalytic subunit, beta isoform | 19053 | 0.167270377 | 0.189 | 0.059 | 0.001520583 | 1.03E-07 |
| SS | Layer 5 | iTBS | Ubb | ubiquitin B | 22187 | 0.244768106 | 1 | 1 | 4.90E-05 | 3.32E-09 |
| SS | Layer 5 | iTBS | Rpl35 | ribosomal protein L35 | 66489 | 0.274108668 | 0.991 | 0.96 | 0.000102629 | 6.94E-09 |
| SS | Layer 5 | iTBS | Uba52 | ubiquitin A-52 residue ribosomal protein fusion product 1 | 22186 | 0.52930806 | 0.869 | 0.753 | 4.14E-09 | 2.80E-13 |
| SS | Layer 5 | iTBS | Gm10076 | ribosomal protein L41 pseudogene | 100126819 | 0.965664168 | 0.747 | 0.41 | 2.39E-25 | 1.62E-29 |
| SS | Layer 6 | iTBS | Bc1 | brain cytoplasmic RNA 1 | 100568459 | -0.515050145 | 0.429 | 0.667 | 1.14E-08 | 7.74E-13 |
| SS | Layer 6 | iTBS | Tuba1b | tubulin, alpha 1B | 22143 | -0.283839882 | 0.945 | 0.989 | 0.000186732 | 1.26E-08 |

| | | | | | | | | | | |
|---|---|---|---|---|---|---|---|---|---|---|
| SS | Layer 6 | iTBS | Ppia | peptidylprolyl isomerase A | 268373 | 0.160489702 | 1 | 1 | 0.003705946 | 2.51E-07 |
| SS | Layer 6 | iTBS | Rpl21 | ribosomal protein L21 | 19933 | 0.176249392 | 1 | 1 | 0.007535071 | 5.10E-07 |
| SS | Layer 6 | iTBS | Stmn1 | stathmin 1 | 16765 | 0.182839472 | 1 | 1 | 0.02124185 | 1.44E-06 |
| SS | Layer 6 | iTBS | Tmsb10 | thymosin, beta 10 | 19240 | 0.197270893 | 1 | 1 | 0.028219683 | 1.91E-06 |
| SS | Layer 6 | iTBS | Rps24 | ribosomal protein S24 | 20088 | 0.204761193 | 1 | 1 | 0.000780749 | 5.28E-08 |
| SS | Layer 6 | iTBS | Rps7 | ribosomal protein S7 | 20115 | 0.205370161 | 1 | 0.997 | 0.0049521 | 3.35E-07 |
| SS | Layer 6 | iTBS | Fau | Finkel-Biskis-Reilly murine sarcoma virus (FBR-MuSV) ubiquitously expressed (fox derived) | 14109 | 0.208450971 | 1 | 1 | 0.001305517 | 8.83E-08 |
| SS | Layer 6 | iTBS | Ccm2 | cerebral cavernous malformation 2 | 216527 | 0.209186611 | 0.199 | 0.08 | 0.029351529 | 1.99E-06 |
| SS | Layer 6 | iTBS | Rpl13 | ribosomal protein L13 | 270106 | 0.221383199 | 1 | 1 | 8.76E-06 | 5.93E-10 |
| SS | Layer 6 | iTBS | Rpl9 | ribosomal protein L9 | 20005 | 0.228466136 | 0.995 | 0.992 | 0.000223451 | 1.51E-08 |
| SS | Layer 6 | iTBS | Gpx4 | glutathione peroxidase 4 | 625249 | 0.236944967 | 1 | 1 | 0.000298738 | 2.02E-08 |
| SS | Layer 6 | iTBS | Rps27a | ribosomal protein S27A | 78294 | 0.244873531 | 1 | 1 | 2.48E-05 | 1.68E-09 |
| SS | Layer 6 | iTBS | Rps10 | ribosomal protein S10 | 67097 | 0.250814978 | 0.997 | 0.979 | 0.003173887 | 2.15E-07 |
| SS | Layer 6 | iTBS | Chd9 | chromodomain helicase DNA binding protein 9 | 109151 | 0.268061223 | 0.325 | 0.144 | 9.88E-05 | 6.69E-09 |
| SS | Layer 6 | iTBS | Eno1 | enolase 1, alpha non-neuron | 13806 | 0.280086245 | 0.963 | 0.939 | 0.002393972 | 1.62E-07 |
| SS | Layer 6 | iTBS | Rps20 | ribosomal protein S20 | 67427 | 0.281960799 | 1 | 0.997 | 7.63E-08 | 5.16E-12 |
| SS | Layer 6 | iTBS | Rpl35 | ribosomal protein L35 | 66489 | 0.36946784 | 0.995 | 0.979 | 3.96E-10 | 2.68E-14 |
| SS | Layer 6 | iTBS | Gstp1 | glutathione S-transferase, pi 1 | 14870 | 0.53023204 | 0.513 | 0.28 | 1.95E-08 | 1.32E-12 |
| SS | Layer 6 | iTBS | Uba52 | ubiquitin A-52 residue ribosomal protein fusion product 1 | 22186 | 0.690898808 | 0.911 | 0.837 | 7.11E-17 | 4.81E-21 |
| SS | Layer 6 | iTBS | Gm10076 | ribosomal protein L41 pseudogene | 100126819 | 0.966468934 | 0.783 | 0.563 | 7.94E-22 | 5.37E-26 |
| SS | Layer 2/3 | cTBS | Bc1 | brain cytoplasmic RNA 1 | 100568459 | -0.934349372 | 0.348 | 0.726 | 2.36E-15 | 1.59E-19 |
| SS | Layer 2/3 | cTBS | Ikbkb | inhibitor of kappaB kinase beta | 16150 | -0.308652092 | 0.235 | 0.416 | 0.021137299 | 1.43E-06 |
| SS | Layer 2/3 | cTBS | Ttr | transthyretin | 22139 | -0.252681043 | 0 | 0.122 | 0.000202003 | 1.37E-08 |
| SS | Layer 2/3 | cTBS | Lrrc17 | leucine rich repeat containing 17 | 74511 | -0.216446063 | 0.045 | 0.185 | 0.009168588 | 6.20E-07 |
| SS | Layer 2/3 | cTBS | Camk2n1 | calcium/calmodulin-dependent protein kinase II inhibitor 1 | 66259 | 0.16716948 | 1 | 1 | 0.006776046 | 4.59E-07 |
| SS | Layer 2/3 | cTBS | Psd | pleckstrin and Sec7 domain containing | 73728 | 0.288558604 | 0.955 | 0.842 | 0.000261706 | 1.77E-08 |
| SS | Layer 2/3 | cTBS | Rpl35 | ribosomal protein L35 | 66489 | 0.3468535 | 0.98 | 0.947 | 0.001526864 | 1.03E-07 |
| SS | Layer 2/3 | cTBS | Uba52 | ubiquitin A-52 residue ribosomal protein fusion product 1 | 22186 | 0.49596171 | 0.737 | 0.64 | 0.001616433 | 1.09E-07 |
| SS | Layer 2/3 | cTBS | Gm10076 | ribosomal protein L41 pseudogene | 100126819 | 0.825540773 | 0.704 | 0.419 | 4.18E-15 | 2.83E-19 |
| SS | Layer 4 | cTBS | Pcp4 | Purkinje cell protein 4 | 18546 | -0.556809893 | 0.604 | 0.755 | 0.001595321 | 1.08E-07 |
| SS | Layer 4 | cTBS | Bc1 | brain cytoplasmic RNA 1 | 100568459 | -0.553197494 | 0.266 | 0.563 | 2.10E-08 | 1.42E-12 |
| SS | Layer 4 | cTBS | Ttr | transthyretin | 22139 | -0.294428991 | 0.015 | 0.162 | 7.06E-07 | 4.78E-11 |
| SS | Layer 4 | cTBS | Rplp1 | ribosomal protein, large, P1 | 56040 | 0.19543282 | 1 | 1 | 0.04727985 | 3.20E-06 |
| SS | Layer 4 | cTBS | Rps21 | ribosomal protein S21 | 66481 | 0.213574472 | 1 | 1 | 0.007008716 | 4.74E-07 |
| SS | Layer 4 | cTBS | Atp1b1 | ATPase, Na+/K+ transporting, beta 1 polypeptide | 11931 | 0.216766582 | 1 | 1 | 0.024394179 | 1.65E-06 |
| SS | Layer 4 | cTBS | Ncdn | neurochondrin | 26562 | 0.263969948 | 1 | 0.978 | 0.003990921 | 2.70E-07 |
| SS | Layer 4 | cTBS | Camk2n2 | calcium/calmodulin-dependent protein kinase II inhibitor 2 | 73047 | 0.275621869 | 0.908 | 0.843 | 0.015427158 | 1.04E-06 |
| SS | Layer 4 | cTBS | Camk2n1 | calcium/calmodulin-dependent protein kinase II inhibitor 1 | 66259 | 0.29066586 | 1 | 1 | 5.87E-10 | 3.97E-14 |
| SS | Layer 4 | cTBS | Rpl35 | ribosomal protein L35 | 66489 | 0.415216089 | 0.97 | 0.943 | 1.54E-06 | 1.04E-10 |
| SS | Layer 4 | cTBS | Gm10076 | ribosomal protein L41 pseudogene | 100126819 | 0.727480247 | 0.666 | 0.393 | 9.26E-12 | 6.27E-16 |
| SS | Layer 5 | cTBS | Bc1 | brain cytoplasmic RNA 1 | 100568459 | -0.553883679 | 0.364 | 0.598 | 1.01E-07 | 6.82E-12 |
| SS | Layer 5 | cTBS | Ttr | transthyretin | 22139 | -0.248566082 | 0.003 | 0.125 | 8.56E-07 | 5.79E-11 |
| SS | Layer 5 | cTBS | Rplp1 | ribosomal protein, large, P1 | 56040 | 0.171374173 | 1 | 1 | 0.041136227 | 2.78E-06 |
| SS | Layer 5 | cTBS | Rpl38 | ribosomal protein L38 | 67671 | 0.171794203 | 1 | 1 | 0.00326368 | 2.21E-07 |
| SS | Layer 5 | cTBS | Rpl35 | ribosomal protein L35 | 66489 | 0.392826794 | 0.977 | 0.96 | 1.30E-09 | 8.77E-14 |
| SS | Layer 5 | cTBS | Uba52 | ubiquitin A-52 residue ribosomal protein fusion product 1 | 22186 | 0.399234149 | 0.787 | 0.753 | 0.044643899 | 3.02E-06 |
| SS | Layer 5 | cTBS | Gm10076 | ribosomal protein L41 pseudogene | 100126819 | 0.762610332 | 0.685 | 0.41 | 2.01E-17 | 1.36E-21 |
| SS | Layer 6 | cTBS | Bc1 | brain cytoplasmic RNA 1 | 100568459 | -0.643015917 | 0.357 | 0.667 | 2.24E-14 | 1.51E-18 |
| SS | Layer 6 | cTBS | Rps21 | ribosomal protein S21 | 66481 | 0.187773003 | 1 | 1 | 0.020796938 | 1.41E-06 |
| SS | Layer 6 | cTBS | Fth1 | ferritin heavy polypeptide 1 | 14319 | 0.248986853 | 1 | 1 | 3.74E-06 | 2.53E-10 |
| SS | Layer 6 | cTBS | Rpl35 | ribosomal protein L35 | 66489 | 0.287112276 | 0.972 | 0.979 | 0.012213146 | 8.26E-07 |
| SS | Layer 6 | cTBS | Eno1 | enolase 1, alpha non-neuron | 13806 | 0.297044684 | 0.966 | 0.939 | 0.00125099 | 8.47E-08 |
| SS | Layer 6 | cTBS | Gstp1 | glutathione S-transferase, pi 1 | 14870 | 0.463396118 | 0.498 | 0.28 | 6.39E-06 | 4.32E-10 |
| SS | Layer 6 | cTBS | Gm10076 | ribosomal protein L41 pseudogene | 100126819 | 0.648563877 | 0.738 | 0.563 | 2.05E-12 | 1.39E-16 |

**Data S4.** Motor (M1) cortex layer-specific differentially expressed genes (DEGs) identified in aged mice 3h following iTBS, and cTBS.

| M1 layer | Stimulation | Gene Symbol | Gene Name | EntrezID | Avg log2FoldChange | % expressed in stimulated samples | % expressed in sham samples | p.adjust value (Bonferroni correction) | p.value | Higher-order groups |
|---|---|---|---|---|---|---|---|---|---|---|
| Layer 2/3 | iTBS | Gm10076 | ribosomal protein L41 pseudogene | 100126819 | 1.454574365 | 0.819 | 0.301 | 2.76E-46 | 1.87E-50 | |
| Layer 2/3 | iTBS | Uba52 | ubiquitin A-52 residue ribosomal protein fusion product 1 | 22186 | 0.996141378 | 0.86 | 0.592 | 3.20E-29 | 2.17E-33 | |
| Layer 2/3 | iTBS | Gstp1 | glutathione S-transferase, pi 1 | 14870 | 0.433633298 | 0.515 | 0.248 | 2.29E-08 | 1.55E-12 | |
| Layer 2/3 | iTBS | Rpl35 | ribosomal protein L35 | 66489 | 0.428949855 | 0.969 | 0.851 | 2.05E-08 | 1.39E-12 | |
| Layer 2/3 | iTBS | Nr4a1 | nuclear receptor subfamily 4, group A, member 1 | 15370 | 0.325322467 | 0.761 | 0.631 | 0.004682657 | 3.17E-07 | 1 |
| Layer 2/3 | iTBS | Rpl9 | ribosomal protein L9 | 20005 | 0.323306121 | 1 | 0.979 | 2.01E-08 | 1.36E-12 | |
| Layer 2/3 | iTBS | Junb | jun B proto-oncogene | 16477 | 0.298010412 | 0.659 | 0.457 | 0.001060932 | 7.18E-08 | |
| Layer 2/3 | iTBS | Rps8 | ribosomal protein S8 | 20116 | 0.289422095 | 0.99 | 0.982 | 9.05E-06 | 6.13E-10 | |
| Layer 2/3 | iTBS | Rps21 | ribosomal protein S21 | 66481 | 0.281924237 | 1 | 0.996 | 5.16E-07 | 3.49E-11 | |
| Layer 2/3 | iTBS | Rpl23 | ribosomal protein L23 | 65019 | 0.273637269 | 1 | 0.975 | 3.09E-05 | 2.09E-09 | |
| Layer 2/3 | iTBS | Rpl30 | ribosomal protein L30 | 19946 | 0.266937423 | 0.993 | 0.936 | 0.00383604 | 2.60E-07 | |
| Layer 2/3 | iTBS | Rps27a | ribosomal protein S27A | 78294 | 0.262082632 | 1 | 0.986 | 0.001235155 | 8.36E-08 | |
| Layer 2/3 | iTBS | Rps10 | ribosomal protein S10 | 67097 | 0.256160549 | 0.99 | 0.94 | 0.004997984 | 3.38E-07 | |
| Layer 2/3 | iTBS | Rps24 | ribosomal protein S24 | 20088 | 0.252146286 | 1 | 0.993 | 2.13E-05 | 1.44E-09 | |
| Layer 2/3 | iTBS | Atp6v0e2 | ATPase, H+ transporting, lysosomal V0 subunit E2 | 76252 | -0.26004892 | 0.887 | 0.95 | 0.0048681 | 3.29E-07 | |
| Layer 2/3 | iTBS | Ldha | lactate dehydrogenase A | 16828 | -0.265521606 | 0.908 | 0.943 | 0.014357214 | 9.72E-07 | |
| Layer 2/3 | iTBS | Ikbkb | inhibitor of kappaB kinase beta | 16150 | -0.27390344 | 0.28 | 0.493 | 0.001960654 | 1.33E-07 | 7 |
| Layer 2/3 | iTBS | Syngap1 | synaptic Ras GTPase activating protein 1 homolog (rat) | 240057 | -0.286678646 | 0.215 | 0.433 | 0.000223026 | 1.51E-08 | 1 |
| Layer 2/3 | iTBS | R3hdm4 | R3H domain containing 4 | 109284 | -0.303133015 | 0.652 | 0.773 | 0.038279947 | 2.59E-06 | |
| Layer 2/3 | iTBS | Rpl5 | ribosomal protein L5 | 100503670 | -0.305948502 | 0.922 | 0.943 | 0.001926034 | 1.30E-07 | |
| Layer 2/3 | iTBS | Tuba1b | tubulin, alpha 1B | 22143 | -0.312792785 | 0.976 | 0.993 | 0.000152058 | 1.03E-08 | |
| Layer 2/3 | iTBS | Tuba1a | tubulin, alpha 1A | 22142 | -0.314281119 | 0.997 | 0.993 | 3.22E-07 | 2.18E-11 | 1 |
| Layer 2/3 | iTBS | Epn1 | epsin 1 | 13854 | -0.344058651 | 0.874 | 0.915 | 0.002160859 | 1.46E-07 | |
| Layer 2/3 | iTBS | Syn1 | synapsin I | 20964 | -0.34867417 | 0.973 | 0.972 | 0.032540862 | 2.20E-06 | 1 |
| Layer 2/3 | iTBS | Eef1a2 | eukaryotic translation elongation factor 1 alpha 2 | 13628 | -0.35158734 | 0.997 | 0.996 | 0.001908941 | 1.29E-07 | |
| Layer 2/3 | iTBS | Slc17a7 | solute carrier family 17 (sodium-dependent inorganic phosphate cotransporter), member 7 | 72961 | -0.351803462 | 1 | 1 | 0.000604668 | 4.09E-08 | 1 |
| Layer 2/3 | iTBS | Glul | glutamate-ammonia ligase (glutamine synthetase) | 14645 | -0.368222064 | 0.727 | 0.865 | 0.003130555 | 2.12E-07 | 1 |
| Layer 2/3 | iTBS | Syngr1 | synaptogyrin 1 | 20972 | -0.385827973 | 0.836 | 0.883 | 0.001770927 | 1.20E-07 | 1 |
| Layer 2/3 | iTBS | Actb | actin, beta | 11461 | -0.392267123 | 1 | 0.993 | 2.04E-08 | 1.38E-12 | |
| Layer 2/3 | iTBS | Dnm1 | dynamin 1 | 13429 | -0.394078197 | 0.997 | 1 | 4.92E-08 | 3.33E-12 | 1 |
| Layer 2/3 | iTBS | Cplx2 | complexin 2 | 12890 | -0.436041517 | 0.792 | 0.869 | 0.000171026 | 1.16E-08 | 1 |
| Layer 2/3 | iTBS | Dynll2 | dynein light chain LC8-type 2 | 68097 | -0.439269398 | 0.966 | 0.979 | 7.70E-06 | 5.21E-10 | |
| Layer 2/3 | iTBS | Cdk5r2 | cyclin-dependent kinase 5, regulatory subunit 2 (p39) | 12570 | -0.532907036 | 0.365 | 0.688 | 7.93E-13 | 5.37E-17 | |
| Layer 2/3 | iTBS | Nme7 | NME/NM23 family member 7 | 171567 | -0.633147575 | 0.266 | 0.461 | 0.000888303 | 6.01E-08 | |
| Layer 2/3 | iTBS | Bc1 | brain cytoplasmic RNA 1 | 100568459 | -0.930399046 | 0.468 | 0.78 | 1.53E-16 | 1.04E-20 | 1 |
| Layer 5 | iTBS | Gm10076 | ribosomal protein L41 pseudogene | 100126819 | 1.28664674 | 0.764 | 0.321 | 6.03E-70 | 4.08E-74 | |
| Layer 5 | iTBS | Uba52 | ubiquitin A-52 residue ribosomal protein fusion product 1 | 22186 | 0.779851038 | 0.861 | 0.664 | 5.64E-35 | 3.82E-39 | |
| Layer 5 | iTBS | Gstp1 | glutathione S-transferase, pi 1 | 14870 | 0.389247318 | 0.537 | 0.31 | 2.86E-12 | 1.94E-16 | |
| Layer 5 | iTBS | Rpl35 | ribosomal protein L35 | 66489 | 0.366759307 | 0.981 | 0.901 | 3.65E-12 | 2.47E-16 | |
| Layer 5 | iTBS | Rps8 | ribosomal protein S8 | 20116 | 0.298601597 | 1 | 0.994 | 7.97E-15 | 5.39E-19 | |
| Layer 5 | iTBS | Pvalb | parvalbumin | 19293 | 0.29439878 | 0.838 | 0.762 | 0.021682193 | 1.47E-06 | 1 |
| Layer 5 | iTBS | Rpl9 | ribosomal protein L9 | 20005 | 0.28607227 | 0.998 | 0.983 | 1.75E-13 | 1.18E-17 | |
| Layer 5 | iTBS | Tmsb10 | thymosin, beta 10 | 19240 | 0.266888129 | 0.998 | 0.996 | 6.54E-10 | 4.42E-14 | |
| Layer 5 | iTBS | Cox7c | cytochrome c oxidase subunit 7C | 12867 | 0.251816906 | 1 | 0.998 | 1.95E-11 | 1.32E-15 | |
| Layer 5 | iTBS | Dusp26 | dual specificity phosphatase 26 (putative) | 66959 | -0.252448117 | 0.388 | 0.559 | 5.24E-05 | 3.54E-09 | |
| Layer 5 | iTBS | Emc10 | ER membrane protein complex subunit 10 | 69683 | -0.253231918 | 0.815 | 0.863 | 0.000401949 | 2.72E-08 | |
| Layer 5 | iTBS | Epn1 | epsin 1 | 13854 | -0.257900631 | 0.846 | 0.893 | 0.009364102 | 6.34E-07 | |
| Layer 5 | iTBS | Ivns1abp | influenza virus NS1A binding protein | 117198 | -0.26005929 | 0.456 | 0.589 | 0.014667245 | 9.93E-07 | |
| Layer 5 | iTBS | Rpl5 | ribosomal protein L5 | 100503670 | -0.264821637 | 0.923 | 0.962 | 6.73E-06 | 4.55E-10 | |
| Layer 5 | iTBS | Slc17a7 | solute carrier family 17 (sodium-dependent inorganic phosphate cotransporter), member 7 | 72961 | -0.267703515 | 0.996 | 0.996 | 0.003462609 | 2.34E-07 | 1 |
| Layer 5 | iTBS | Kcnq2 | potassium voltage-gated channel, subfamily Q, member 2 | 16536 | -0.268063584 | 0.573 | 0.692 | 0.003602513 | 2.44E-07 | 3 |
| Layer 5 | iTBS | Phyhip | phytanoyl-CoA hydroxylase interacting protein | 105653 | -0.284007447 | 0.427 | 0.583 | 2.62E-05 | 1.77E-09 | |
| Layer 5 | iTBS | Actb | actin, beta | 11461 | -0.286079493 | 0.994 | 0.994 | 1.22E-06 | 8.25E-11 | |
| Layer 5 | iTBS | Sptbn2 | spectrin beta, non-erythrocytic 2 | 20743 | -0.28839933 | 0.809 | 0.865 | 0.009945331 | 6.73E-07 | |
| Layer 5 | iTBS | Ndrg2 | N-myc downstream regulated gene 2 | 29811 | -0.289289769 | 0.875 | 0.917 | 9.89E-05 | 6.70E-09 | |
| Layer 5 | iTBS | Stx1b | syntaxin 1B | 56216 | -0.293057921 | 0.828 | 0.874 | 4.40E-05 | 2.98E-09 | 1 |
| Layer 5 | iTBS | Glul | glutamate-ammonia ligase (glutamine synthetase) | 14645 | -0.293099767 | 0.815 | 0.872 | 5.70E-05 | 3.85E-09 | 1 |
| Layer 5 | iTBS | Tuba1b | tubulin, alpha 1B | 22143 | -0.304908539 | 0.967 | 0.987 | 8.58E-12 | 5.80E-16 | |
| Layer 5 | iTBS | Dnm1 | dynamin 1 | 13429 | -0.309624637 | 0.998 | 1 | 5.74E-09 | 3.88E-13 | 1 |
| Layer 5 | iTBS | Eef1a2 | eukaryotic translation elongation factor 1 alpha 2 | 13628 | -0.313475648 | 1 | 0.998 | 4.66E-08 | 3.15E-12 | |
| Layer 5 | iTBS | Hpcal4 | hippocalcin-like 4 | 170638 | -0.326978757 | 0.656 | 0.805 | 5.63E-05 | 3.81E-09 | |
| Layer 5 | iTBS | Cdk5r2 | cyclin-dependent kinase 5, regulatory subunit 2 (p39) | 12570 | -0.332047506 | 0.402 | 0.587 | 3.69E-08 | 2.50E-12 | |
| Layer 5 | iTBS | Dynll2 | dynein light chain LC8-type 2 | 68097 | -0.361690156 | 0.959 | 0.977 | 1.27E-07 | 8.62E-12 | |
| Layer 5 | iTBS | Cplx2 | complexin 2 | 12890 | -0.547573969 | 0.645 | 0.762 | 5.93E-12 | 4.01E-16 | 1 |
| Layer 5 | iTBS | Bc1 | brain cytoplasmic RNA 1 | 100568459 | -0.556184087 | 0.49 | 0.73 | 1.43E-16 | 9.67E-21 | 1 |
| Layer 6 | iTBS | Gm10076 | ribosomal protein L41 pseudogene | 100126819 | 1.224857727 | 0.793 | 0.49 | 4.21E-18 | 2.85E-22 | |
| Layer 6 | iTBS | Uba52 | ubiquitin A-52 residue ribosomal protein fusion product 1 | 22186 | 0.774490163 | 0.91 | 0.776 | 5.78E-12 | 3.91E-16 | |
| Layer 6 | iTBS | Gstp1 | glutathione S-transferase, pi 1 | 14870 | 0.644680111 | 0.644 | 0.344 | 1.63E-07 | 1.10E-11 | |
| Layer 6 | iTBS | Rpl35 | ribosomal protein L35 | 66489 | 0.337141937 | 0.995 | 0.979 | 0.001271736 | 8.61E-08 | |
| Layer 6 | iTBS | Sytl2 | synaptotagmin-like 2 | 83671 | 0.31277338 | 0.351 | 0.141 | 0.017684192 | 1.20E-06 | |
| Layer 6 | iTBS | Rps8 | ribosomal protein S8 | 20116 | 0.293572287 | 0.995 | 0.995 | 0.00213008 | 1.44E-07 | |
| Layer 6 | iTBS | Ndufb9 | NADH:ubiquinone oxidoreductase subunit B9 | 66218 | 0.280285928 | 1 | 0.984 | 0.039632005 | 2.68E-06 | |
| Layer 6 | iTBS | Rpl38 | ribosomal protein L38 | 67671 | 0.261461166 | 1 | 1 | 0.002561907 | 1.73E-07 | |
| Layer 6 | iTBS | Eef1a2 | eukaryotic translation elongation factor 1 alpha 2 | 13628 | -0.507722247 | 0.979 | 0.995 | 2.37E-05 | 1.60E-09 | |
| Layer 2/3 | cTBS | Gm10076 | ribosomal protein L41 pseudogene | 100126819 | 1.208843802 | 0.777 | 0.301 | 1.07E-33 | 7.24E-38 | |
| Layer 2/3 | cTBS | Uba52 | ubiquitin A-52 residue ribosomal protein fusion product 1 | 22186 | 0.671484509 | 0.79 | 0.592 | 9.79E-12 | 6.63E-16 | |
| Layer 2/3 | cTBS | Rpl35 | ribosomal protein L35 | 66489 | 0.586394679 | 0.966 | 0.851 | 8.56E-14 | 5.79E-18 | |
| Layer 2/3 | cTBS | Gstp1 | glutathione S-transferase, pi 1 | 14870 | 0.344800544 | 0.458 | 0.248 | 0.001363282 | 9.23E-08 | |
| Layer 2/3 | cTBS | Eno1 | enolase 1, alpha non-neuron | 13806 | 0.327195749 | 0.962 | 0.894 | 0.003632396 | 2.46E-07 | |
| Layer 2/3 | cTBS | Atp5k | ATP synthase, H+ transporting, mitochondrial F1F0 complex, subunit E | 11958 | 0.287120899 | 1 | 0.989 | 9.85E-05 | 6.67E-09 | |
| Layer 2/3 | cTBS | Uqcr11 | ubiquinol-cytochrome c reductase, complex III subunit XI | 66594 | 0.26341501 | 1 | 0.996 | 0.000354382 | 2.40E-08 | |
| Layer 2/3 | cTBS | Fam207a | family with sequence similarity 207, member A | 108707 | -0.25483742 | 0.092 | 0.287 | 0.00039857 | 2.70E-08 | |
| Layer 2/3 | cTBS | Ikbkb | inhibitor of kappaB kinase beta | 16150 | -0.267576532 | 0.277 | 0.493 | 0.009503442 | 6.43E-07 | 7 |
| Layer 2/3 | cTBS | Tuba1b | tubulin, alpha 1B | 22143 | -0.29944023 | 0.987 | 0.993 | 0.00228783 | 1.55E-07 | |
| Layer 2/3 | cTBS | Tuba1a | tubulin, alpha 1A | 22142 | -0.300088442 | 0.992 | 0.993 | 2.38E-05 | 1.61E-09 | 1 |
| Layer 2/3 | cTBS | Basp1 | brain abundant, membrane attached signal protein 1 | 70350 | -0.306187338 | 0.807 | 0.947 | 0.002924508 | 1.98E-07 | 1 |
| Layer 2/3 | cTBS | Tatdn1 | TatD DNase domain containing 1 | 69694 | -0.309512891 | 0.126 | 0.358 | 1.18E-05 | 7.97E-10 | |
| Layer 2/3 | cTBS | Cdk5r2 | cyclin-dependent kinase 5, regulatory subunit 2 (p39) | 12570 | -0.348726088 | 0.508 | 0.688 | 0.007100877 | 4.81E-07 | |
| Layer 2/3 | cTBS | Ttr | transthyretin | 22139 | -0.737194838 | 0.013 | 0.234 | 1.61E-09 | 1.09E-13 | 1 |
| Layer 2/3 | cTBS | Bc1 | brain cytoplasmic RNA 1 | 100568459 | -1.093216524 | 0.382 | 0.78 | 1.21E-20 | 8.18E-25 | 1 |
| Layer 5 | cTBS | Gm10076 | ribosomal protein L41 pseudogene | 100126819 | 1.19205091 | 0.774 | 0.321 | 2.29E-69 | 1.55E-73 | |
| Layer 5 | cTBS | Rpl35 | ribosomal protein L35 | 66489 | 0.629817196 | 0.987 | 0.901 | 4.94E-41 | 3.34E-45 | |
| Layer 5 | cTBS | Uba52 | ubiquitin A-52 residue ribosomal protein fusion product 1 | 22186 | 0.621487028 | 0.828 | 0.664 | 1.45E-24 | 9.79E-29 | |
| Layer 5 | cTBS | Gstp1 | glutathione S-transferase, pi 1 | 14870 | 0.341951511 | 0.477 | 0.31 | 1.18E-06 | 7.99E-11 | |
| Layer 5 | cTBS | Rps29 | ribosomal protein S29 | 20090 | 0.304107573 | 0.996 | 0.983 | 3.60E-13 | 2.43E-17 | |
| Layer 5 | cTBS | Rps27 | ribosomal protein S27 | 57294 | 0.287594688 | 0.965 | 0.878 | 1.61E-07 | 1.09E-11 | |
| Layer 5 | cTBS | Eno1 | enolase 1, alpha non-neuron | 13806 | 0.281712608 | 0.963 | 0.904 | 3.11E-06 | 2.11E-10 | |
| Layer 5 | cTBS | Rpl37a | ribosomal protein L37a | 19981 | 0.255790516 | 0.994 | 0.972 | 5.59E-06 | 3.78E-10 | |
| Layer 5 | cTBS | Cox7c | cytochrome c oxidase subunit 7C | 12867 | 0.255584577 | 0.998 | 0.998 | 5.10E-11 | 3.45E-15 | |
| Layer 5 | cTBS | Basp1 | brain abundant, membrane attached signal protein 1 | 70350 | -0.281820053 | 0.804 | 0.946 | 8.32E-08 | 5.63E-12 | 1 |
| Layer 5 | cTBS | Cdk5r2 | cyclin-dependent kinase 5, regulatory subunit 2 (p39) | 12570 | -0.306716563 | 0.403 | 0.587 | 3.65E-07 | 2.47E-11 | |
| Layer 5 | cTBS | Atp1a3 | ATPase, Na+/K+ transporting, alpha 3 polypeptide | 232975 | -0.319363151 | 0.804 | 0.919 | 1.66E-07 | 1.12E-11 | 3 |
| Layer 5 | cTBS | Ikbkb | inhibitor of kappaB kinase beta | 16150 | -0.322632072 | 0.226 | 0.445 | 5.89E-11 | 3.99E-15 | 7 |
| Layer 5 | cTBS | Ttr | transthyretin | 22139 | -0.478572331 | 0.026 | 0.216 | 8.28E-18 | 5.60E-22 | 1 |
| Layer 5 | cTBS | Bc1 | brain cytoplasmic RNA 1 | 100568459 | -0.719464524 | 0.359 | 0.73 | 1.30E-31 | 8.80E-36 | 1 |
| Layer 6 | cTBS | Gm10076 | ribosomal protein L41 pseudogene | 100126819 | 0.822078401 | 0.719 | 0.49 | 4.71E-10 | 3.19E-14 | |
| Layer 6 | cTBS | Uba52 | ubiquitin A-52 residue ribosomal protein fusion product 1 | 22186 | 0.562418974 | 0.824 | 0.776 | 0.000970955 | 6.57E-08 | |
| Layer 6 | cTBS | Rpl35 | ribosomal protein L35 | 66489 | 0.484942586 | 1 | 0.979 | 4.17E-08 | 2.82E-12 | |
| Layer 6 | cTBS | Ftl1 | ferritin light polypeptide 1 | 14325 | 0.332260911 | 0.991 | 0.964 | 0.013163305 | 8.91E-07 | |
| Layer 6 | cTBS | Pcp4 | Purkinje cell protein 4 | 18546 | 0.325715802 | 1 | 1 | 0.011314731 | 7.66E-07 | 1 |
| Layer 6 | cTBS | Rpl38 | ribosomal protein L38 | 67671 | 0.286612031 | 1 | 1 | 2.36E-05 | 1.60E-09 | |
| Layer 6 | cTBS | Atp5md | ATP synthase membrane subunit DAPIT | 66477 | 0.267543112 | 1 | 1 | 0.000177199 | 1.20E-08 | |
| Layer 6 | cTBS | Cox6c | cytochrome c oxidase subunit 6C | 12864 | 0.257019884 | 1 | 1 | 2.94E-05 | 1.99E-09 | |
| Layer 6 | cTBS | Ttr | transthyretin | 22139 | -0.410667093 | 0.014 | 0.177 | 9.96E-05 | 6.74E-09 | 1 |
| Layer 6 | cTBS | Bc1 | brain cytoplasmic RNA 1 | 100568459 | -0.667902783 | 0.335 | 0.688 | 3.32E-09 | 2.24E-13 | 1 |

**Data S5.** Complete list of enriched Gene Ontology (GO) terms for the significant differentially expressed genes (DEGs) identified in the different motor (M1) cortex layers, following iTBS, and cTBS.

| M1 layer | Stimulation | GO ID | GO term description | Number of genes | Gene symbols | p.value | q.value | Ontology Source | Code |
|---|---|---|---|---|---|---|---|---|---|
| Layer 2/3 | iTBS | GO:0022626 | cytosolic ribosome | 11 | Uba52/Rpl9/Rp | 8.50E-19 | 5.19E-17 | CC | 8 |
| Layer 2/3 | iTBS | GO:0044391 | ribosomal subunit | 11 | Uba52/Rpl9/Rp | 4.05E-16 | 1.24E-14 | CC | 8 |
| Layer 2/3 | iTBS | GO:0005840 | ribosome | 11 | Uba52/Rpl9/Rp | 3.46E-15 | 7.04E-14 | CC | 8 |
| Layer 2/3 | iTBS | GO:0003735 | structural constituent of ribosome | 11 | Uba52/Rpl9/Rp | 4.88E-17 | 4.32E-15 | MF | 8 |
| Layer 2/3 | iTBS | GO:0043209 | myelin sheath | 9 | Uba52/Actb/Dr | 3.54E-12 | 5.40E-11 | CC | 4 |
| Layer 2/3 | iTBS | GO:0098793 | presynapse | 8 | Actb/Dnm1/Cpl | 1.00E-07 | 6.81E-07 | CC | 1 |
| Layer 2/3 | iTBS | GO:0042254 | ribosome biogenesis | 6 | Rpl35/Rps21/Rp | 1.13E-06 | 0.000579396 | BP | 8 |
| Layer 2/3 | iTBS | GO:0022613 | ribonucleoprotein complex biogenesis | 6 | Rpl35/Rps21/Rp | 7.60E-06 | 0.000776279 | BP | 8 |
| Layer 2/3 | iTBS | GO:0022627 | cytosolic small ribosomal subunit | 6 | Uba52/Rps21/R | 2.09E-11 | 2.55E-10 | CC | 8 |
| Layer 2/3 | iTBS | GO:0022625 | cytosolic large ribosomal subunit | 6 | Uba52/Rpl9/Rp | 1.30E-10 | 1.32E-09 | CC | 8 |
| Layer 2/3 | iTBS | GO:0015935 | small ribosomal subunit | 6 | Uba52/Rps21/R | 5.67E-10 | 4.95E-09 | CC | 8 |
| Layer 2/3 | iTBS | GO:0015934 | large ribosomal subunit | 6 | Uba52/Rpl9/Rp | 7.36E-09 | 5.62E-08 | CC | 8 |
| Layer 2/3 | iTBS | GO:0006364 | rRNA processing | 5 | Rpl35/Rps21/Rp | 4.87E-06 | 0.000776279 | BP | 8 |
| Layer 2/3 | iTBS | GO:0016072 | rRNA metabolic process | 5 | Rpl35/Rps21/Rp | 6.09E-06 | 0.000776279 | BP | 8 |
| Layer 2/3 | iTBS | GO:0099504 | synaptic vesicle cycle | 5 | Actb/Dnm1/Cpl | 6.63E-06 | 0.000776279 | BP | 1 |
| Layer 2/3 | iTBS | GO:0099003 | vesicle-mediated transport in synapse | 5 | Actb/Dnm1/Cpl | 1.14E-05 | 0.000970232 | BP | 1 |
| Layer 2/3 | iTBS | GO:0034470 | ncRNA processing | 5 | Rpl35/Rps21/Rp | 6.08E-05 | 0.003531833 | BP | |
| Layer 2/3 | iTBS | GO:0034660 | ncRNA metabolic process | 5 | Rpl35/Rps21/Rp | 0.000173986 | 0.006225221 | BP | |
| Layer 2/3 | iTBS | GO:0098978 | glutamatergic synapse | 5 | Actb/Dnm1/Dy | 3.94E-05 | 0.000240387 | CC | 1 |
| Layer 2/3 | iTBS | GO:0031625 | ubiquitin protein ligase binding | 5 | Uba52/Rpl23/T | 3.35E-05 | 0.001097492 | MF | |
| Layer 2/3 | iTBS | GO:0044389 | ubiquitin-like protein ligase binding | 5 | Uba52/Rpl23/T | 4.31E-05 | 0.001097492 | MF | |
| Layer 2/3 | iTBS | GO:0006836 | neurotransmitter transport | 4 | Cplx2/Slc17a7/I | 0.000185748 | 0.006225221 | BP | 2 |
| Layer 2/3 | iTBS | GO:0001505 | regulation of neurotransmitter levels | 4 | Cplx2/Slc17a7/I | 0.000226453 | 0.006267142 | BP | 2 |
| Layer 2/3 | iTBS | GO:0051348 | negative regulation of transferase activity | 4 | Actb/Gstp1/Rpl | 0.000233241 | 0.006267142 | BP | |
| Layer 2/3 | iTBS | GO:0008021 | synaptic vesicle | 4 | Dnm1/Slc17a7/ | 0.000108553 | 0.000602494 | CC | 1 |
| Layer 2/3 | iTBS | GO:0070382 | exocytic vesicle | 4 | Dnm1/Slc17a7/ | 0.000163341 | 0.000712314 | CC | |
| Layer 2/3 | iTBS | GO:0030133 | transport vesicle | 4 | Dnm1/Slc17a7/ | 0.000556414 | 0.001887251 | CC | |
| Layer 2/3 | iTBS | GO:0014069 | postsynaptic density | 4 | Actb/Syngap1/I | 0.000647358 | 0.002080153 | CC | 1 |
| Layer 2/3 | iTBS | GO:0032279 | asymmetric synapse | 4 | Actb/Syngap1/I | 0.0007162 | 0.002186295 | CC | 1 |
| Layer 2/3 | iTBS | GO:0099572 | postsynaptic specialization | 4 | Actb/Syngap1/I | 0.000833386 | 0.002312746 | CC | 1 |
| Layer 2/3 | iTBS | GO:0098984 | neuron to neuron synapse | 4 | Actb/Syngap1/I | 0.000906122 | 0.002405267 | CC | 1 |
| Layer 2/3 | iTBS | GO:0150034 | distal axon | 4 | Cdk5r2/Actb/Cp | 0.001023477 | 0.002603583 | CC | |
| Layer 2/3 | iTBS | GO:0005874 | microtubule | 4 | Dnm1/Tuba1a/ | 0.001715549 | 0.003879214 | CC | |
| Layer 2/3 | iTBS | GO:0003924 | GTPase activity | 4 | Dnm1/Tuba1a/ | 0.000406476 | 0.003267366 | MF | |
| Layer 2/3 | iTBS | GO:0048168 | regulation of neuronal synaptic plasticity | 3 | Syngap1/Syngr1 | 7.63E-05 | 0.003531833 | BP | 1 |
| Layer 2/3 | iTBS | GO:0048488 | synaptic vesicle endocytosis | 3 | Actb/Dnm1/Slc | 7.96E-05 | 0.003531833 | BP | 1 |
| Layer 2/3 | iTBS | GO:0140238 | presynaptic endocytosis | 3 | Actb/Dnm1/Slc | 7.96E-05 | 0.003531833 | BP | 1 |
| Layer 2/3 | iTBS | GO:0042274 | ribosomal small subunit biogenesis | 3 | Rps21/Rps8/Rp | 8.30E-05 | 0.003531833 | BP | 8 |
| Layer 2/3 | iTBS | GO:0036465 | synaptic vesicle recycling | 3 | Actb/Dnm1/Slc | 0.000122289 | 0.004802442 | BP | 1 |
| Layer 2/3 | iTBS | GO:0002181 | cytoplasmic translation | 3 | Rpl9/Rps21/Rpl | 0.0001951 | 0.006225221 | BP | |
| Layer 2/3 | iTBS | GO:0030672 | synaptic vesicle membrane | 3 | Slc17a7/Syngr1 | 0.000130306 | 0.000611965 | CC | 1 |
| Layer 2/3 | iTBS | GO:0099501 | exocytic vesicle membrane | 3 | Slc17a7/Syngr1 | 0.000130306 | 0.000611965 | CC | |
| Layer 2/3 | iTBS | GO:0030658 | transport vesicle membrane | 3 | Slc17a7/Syngr1 | 0.000405226 | 0.001509937 | CC | |
| Layer 2/3 | iTBS | GO:0043679 | axon terminus | 3 | Actb/Cplx2/Glu | 0.001523325 | 0.003577039 | CC | |
| Layer 2/3 | iTBS | GO:0044306 | neuron projection terminus | 3 | Actb/Cplx2/Glu | 0.002088393 | 0.004553639 | CC | |
| Layer 2/3 | iTBS | GO:0005200 | structural constituent of cytoskeleton | 3 | Actb/Tuba1a/T | 6.19E-05 | 0.001097492 | MF | |
| Layer 2/3 | iTBS | GO:0019843 | rRNA binding | 3 | Rpl9/Rpl23/Rpl | 9.94E-05 | 0.001098673 | MF | 8 |
| Layer 2/3 | iTBS | GO:1904667 | negative regulation of ubiquitin protein ligase activity | 2 | Rpl23/Rpl5 | 7.29E-05 | 0.003531833 | BP | |
| Layer 2/3 | iTBS | GO:0051444 | negative regulation of ubiquitin-protein transferase activity | 2 | Rpl23/Rpl5 | 0.000225185 | 0.006267142 | BP | |
| Layer 2/3 | iTBS | GO:0044305 | calyx of Held | 2 | Actb/Cplx2 | 0.000215569 | 0.000877405 | CC | |
| Layer 2/3 | iTBS | GO:0098563 | intrinsic component of synaptic vesicle membrane | 2 | Slc17a7/Syn1 | 0.000420439 | 0.001509937 | CC | 1 |
| Layer 2/3 | iTBS | GO:0042788 | polysomal ribosome | 2 | Rps21/Rpl30 | 0.000827554 | 0.002312746 | CC | 8 |
| Layer 2/3 | iTBS | GO:0030665 | clathrin-coated vesicle membrane | 2 | Epn1/Syn1 | 0.001306772 | 0.003191275 | CC | 1 |
| Layer 2/3 | iTBS | GO:1990948 | ubiquitin ligase inhibitor activity | 2 | Rpl23/Rpl5 | 6.21E-05 | 0.001097492 | MF | |
| Layer 2/3 | iTBS | GO:0055105 | ubiquitin-protein transferase inhibitor activity | 2 | Rpl23/Rpl5 | 7.58E-05 | 0.001098673 | MF | |
| Layer 2/3 | iTBS | GO:0031386 | protein tag | 2 | Uba52/Rps27a | 9.09E-05 | 0.001098673 | MF | |
| Layer 2/3 | iTBS | GO:0050998 | nitric-oxide synthase binding | 2 | Actb/Dnm1 | 0.000287237 | 0.002821977 | MF | |
| Layer 2/3 | iTBS | GO:0055106 | ubiquitin-protein transferase regulator activity | 2 | Rpl23/Rpl5 | 0.000376663 | 0.003267366 | MF | |
| Layer 5 | iTBS | GO:0098793 | presynapse | 8 | Cplx2/Dnm1/Ac | 2.00E-08 | 1.60E-06 | CC | 1 |
| Layer 5 | iTBS | GO:0099504 | synaptic vesicle cycle | 6 | Cplx2/Dnm1/Ac | 9.01E-08 | 3.11E-05 | BP | 1 |
| Layer 5 | iTBS | GO:0099003 | vesicle-mediated transport in synapse | 6 | Cplx2/Dnm1/Ac | 1.74E-07 | 3.11E-05 | BP | 1 |
| Layer 5 | iTBS | GO:0043209 | myelin sheath | 6 | Uba52/Tuba1b, | 6.53E-08 | 2.57E-06 | CC | 4 |
| Layer 5 | iTBS | GO:0022626 | cytosolic ribosome | 5 | Uba52/Rps8/Rp | 9.65E-08 | 2.57E-06 | CC | 8 |
| Layer 5 | iTBS | GO:0044391 | ribosomal subunit | 5 | Uba52/Rps8/Rp | 1.46E-06 | 2.33E-05 | CC | 8 |
| Layer 5 | iTBS | GO:0005840 | ribosome | 5 | Uba52/Rps8/Rp | 3.75E-06 | 5.01E-05 | CC | 8 |
| Layer 5 | iTBS | GO:0098978 | glutamatergic synapse | 5 | Cplx2/Dnm1/Dy | 1.52E-05 | 0.000152066 | CC | 1 |
| Layer 5 | iTBS | GO:0003735 | structural constituent of ribosome | 5 | Uba52/Rps8/Rp | 6.76E-07 | 6.05E-05 | MF | 8 |
| Layer 5 | iTBS | GO:0036465 | synaptic vesicle recycling | 4 | Dnm1/Actb/Stx | 1.33E-06 | 0.000158747 | BP | 1 |
| Layer 5 | iTBS | GO:0006836 | neurotransmitter transport | 4 | Cplx2/Stx1b/Slc | 9.83E-05 | 0.004385529 | BP | 2 |
| Layer 5 | iTBS | GO:0001505 | regulation of neurotransmitter levels | 4 | Cplx2/Stx1b/Slc | 0.000120077 | 0.004414519 | BP | 2 |
| Layer 5 | iTBS | GO:0051348 | negative regulation of transferase activity | 4 | Gstp1/Actb/Rpl | 0.000123711 | 0.004414519 | BP | |
| Layer 5 | iTBS | GO:0016050 | vesicle organization | 4 | Cplx2/Dnm1/St | 0.000176152 | 0.005525982 | BP | 1 |
| Layer 5 | iTBS | GO:0045055 | regulated exocytosis | 4 | Cplx2/Cdk5r2/S | 0.000185829 | 0.005525982 | BP | |
| Layer 5 | iTBS | GO:0042254 | ribosome biogenesis | 4 | Rps8/Rpl35/Rpl | 0.00020633 | 0.005663629 | BP | 8 |
| Layer 5 | iTBS | GO:0022625 | cytosolic large ribosomal subunit | 4 | Uba52/Rpl9/Rp | 4.68E-07 | 9.37E-06 | CC | 8 |
| Layer 5 | iTBS | GO:0015934 | large ribosomal subunit | 4 | Uba52/Rpl9/Rp | 6.63E-06 | 7.58E-05 | CC | 8 |
| Layer 5 | iTBS | GO:0008021 | synaptic vesicle | 4 | Dnm1/Stx1b/Sl | 5.09E-05 | 0.000452517 | CC | 1 |
| Layer 5 | iTBS | GO:0070382 | exocytic vesicle | 4 | Dnm1/Stx1b/Sl | 7.69E-05 | 0.000615148 | CC | |
| Layer 5 | iTBS | GO:0030133 | transport vesicle | 4 | Dnm1/Stx1b/Sl | 0.000265712 | 0.001635149 | CC | |
| Layer 5 | iTBS | GO:0150034 | distal axon | 4 | Cplx2/Cdk5r2/A | 0.000493266 | 0.002630753 | CC | |
| Layer 5 | iTBS | GO:1903421 | regulation of synaptic vesicle recycling | 3 | Dnm1/Stx1b/Sl | 4.80E-06 | 0.000428491 | BP | 1 |
| Layer 5 | iTBS | GO:0048488 | synaptic vesicle endocytosis | 3 | Dnm1/Actb/Slc | 4.90E-05 | 0.002917031 | BP | 1 |
| Layer 5 | iTBS | GO:0140238 | presynaptic endocytosis | 3 | Dnm1/Actb/Slc | 4.90E-05 | 0.002917031 | BP | 1 |
| Layer 5 | iTBS | GO:0070373 | negative regulation of ERK1 and ERK2 cascade | 3 | Gstp1/Dusp26/ | 7.82E-05 | 0.003987176 | BP | |
| Layer 5 | iTBS | GO:0030863 | cortical cytoskeleton | 3 | Actb/Sptbn2/Pv | 0.000247863 | 0.001635149 | CC | |
| Layer 5 | iTBS | GO:0042734 | presynaptic membrane | 3 | Dnm1/Stx1b/Ep | 0.000305531 | 0.00174589 | CC | 1 |
| Layer 5 | iTBS | GO:0043679 | axon terminus | 3 | Cplx2/Actb/Glu | 0.000875169 | 0.004375846 | CC | |
| Layer 5 | iTBS | GO:0044306 | neuron projection terminus | 3 | Cplx2/Actb/Glu | 0.001203869 | 0.005068921 | CC | |
| Layer 5 | iTBS | GO:0031629 | synaptic vesicle fusion to presynaptic active zone membrane | 2 | Cplx2/Stx1b | 0.000262734 | 0.006250311 | BP | 1 |
| Layer 5 | iTBS | GO:1900242 | regulation of synaptic vesicle endocytosis | 2 | Dnm1/Slc17a7 | 0.000262734 | 0.006250311 | BP | 1 |
| Layer 5 | iTBS | GO:0044305 | calyx of Held | 2 | Cplx2/Actb | 0.00014742 | 0.001072149 | CC | |
| Layer 5 | iTBS | GO:0022627 | cytosolic small ribosomal subunit | 2 | Uba52/Rps8 | 0.001022947 | 0.004741153 | CC | 8 |
| Layer 5 | iTBS | GO:0031201 | SNARE complex | 2 | Cplx2/Stx1b | 0.001066759 | 0.004741153 | CC | 1 |
| Layer 6 | iTBS | GO:0022626 | cytosolic ribosome | 4 | Uba52/Rpl35/R | 2.86E-08 | 3.91E-07 | CC | 8 |
| Layer 6 | iTBS | GO:0044391 | ribosomal subunit | 4 | Uba52/Rpl35/R | 2.57E-07 | 1.76E-06 | CC | 8 |
| Layer 6 | iTBS | GO:0005840 | ribosome | 4 | Uba52/Rpl35/R | 5.57E-07 | 2.54E-06 | CC | 8 |
| Layer 6 | iTBS | GO:0003735 | structural constituent of ribosome | 4 | Uba52/Rpl35/R | 7.70E-08 | 1.30E-06 | MF | 8 |
| Layer 6 | iTBS | GO:0022625 | cytosolic large ribosomal subunit | 3 | Uba52/Rpl35/R | 8.23E-07 | 2.81E-06 | CC | 8 |
| Layer 6 | iTBS | GO:0015934 | large ribosomal subunit | 3 | Uba52/Rpl35/R | 6.06E-06 | 1.66E-05 | CC | 8 |
| Layer 6 | iTBS | GO:0022627 | cytosolic small ribosomal subunit | 2 | Uba52/Rps8 | 9.26E-05 | 0.000211207 | CC | 8 |
| Layer 6 | iTBS | GO:0015935 | small ribosomal subunit | 2 | Uba52/Rps8 | 0.000269268 | 0.000526389 | CC | 8 |
| Layer 6 | iTBS | GO:0043209 | myelin sheath | 2 | Uba52/Eef1a2 | 0.001917959 | 0.003280718 | CC | 4 |
| Layer 2/3 | cTBS | GO:0043209 | myelin sheath | 4 | Uba52/Tuba1a, | 5.06E-06 | 9.59E-05 | CC | 4 |
| Layer 5 | cTBS | GO:0022626 | cytosolic ribosome | 5 | Rpl35/Uba52/R | 2.55E-09 | 6.71E-08 | CC | 8 |
| Layer 5 | cTBS | GO:0044391 | ribosomal subunit | 5 | Rpl35/Uba52/R | 3.98E-08 | 5.24E-07 | CC | 8 |
| Layer 5 | cTBS | GO:0005840 | ribosome | 5 | Rpl35/Uba52/R | 1.04E-07 | 9.16E-07 | CC | 8 |
| Layer 5 | cTBS | GO:0003735 | structural constituent of ribosome | 5 | Rpl35/Uba52/R | 1.83E-08 | 6.35E-07 | MF | 8 |
| Layer 5 | cTBS | GO:0022627 | cytosolic small ribosomal subunit | 3 | Uba52/Rps29/R | 1.81E-06 | 1.19E-05 | CC | 8 |
| Layer 5 | cTBS | GO:0022625 | cytosolic large ribosomal subunit | 3 | Rpl35/Uba52/R | 4.42E-06 | 2.32E-05 | CC | 8 |
| Layer 5 | cTBS | GO:0015935 | small ribosomal subunit | 3 | Uba52/Rps29/R | 9.09E-06 | 3.99E-05 | CC | 8 |
| Layer 5 | cTBS | GO:0015934 | large ribosomal subunit | 3 | Rpl35/Uba52/R | 3.22E-05 | 0.000121151 | CC | 8 |
| Layer 5 | cTBS | GO:0043209 | myelin sheath | 3 | Uba52/Atp1a3/ | 0.000173186 | 0.00056969 | CC | 4 |
| Layer 5 | cTBS | GO:1902554 | serine/threonine protein kinase complex | 2 | Ikbkb/Cdk5r2 | 0.000856061 | 0.002503103 | CC | |
| Layer 5 | cTBS | GO:1902911 | protein kinase complex | 2 | Ikbkb/Cdk5r2 | 0.001233831 | 0.003246923 | CC | |
| Layer 6 | cTBS | GO:0022625 | cytosolic large ribosomal subunit | 3 | Rpl35/Rpl38/Ub | 1.17E-06 | 9.88E-06 | CC | 8 |
| Layer 6 | cTBS | GO:0022626 | cytosolic ribosome | 3 | Rpl35/Rpl38/Ub | 7.04E-06 | 2.42E-05 | CC | 8 |
| Layer 6 | cTBS | GO:0015934 | large ribosomal subunit | 3 | Rpl35/Rpl38/Ub | 8.63E-06 | 2.42E-05 | CC | 8 |
| Layer 6 | cTBS | GO:0044391 | ribosomal subunit | 3 | Rpl35/Rpl38/Ub | 3.62E-05 | 7.62E-05 | CC | 8 |
| Layer 6 | cTBS | GO:0005840 | ribosome | 3 | Rpl35/Rpl38/Ub | 6.43E-05 | 0.000108297 | CC | 8 |
| Layer 6 | cTBS | GO:0003735 | structural constituent of ribosome | 3 | Rpl35/Rpl38/Ub | 2.27E-05 | 0.000239417 | MF | 8 |

| Code key | High order function or process |
|---|---|
| 1 | synapse and/or synaptic plasticity |
| 2 | Neurotransmitter |
| 3 | Intrinsic plasticity and/or membrane excitability |
| 4 | Oligodendrocyte and/or myelin related |
| 5 | neurogenesis |
| 6 | neurotrophin |
| 7 | Cell death |
| 8 | Ribosome |
| 9 | ATP metabolic pathway |

**Data S6.** Somatosensory (SS) cortex layer-specific differentially expressed genes (DEGs) identified in aged mice 3h following iTBS, and cTBS.

| SS layer | Stimulation | Gene Symbol | Gene Name | EntrezID | Avg log2FoldChange | % expressed in stimulated samples | % expressed in sham samples | p.adjust value (Bonferroni correction) | p.value | Higher-order groups |
|---|---|---|---|---|---|---|---|---|---|---|
| Layer 2/3 | iTBS | Gm10076 | ribosomal protein L41 pseudogene | 100126819 | 1.068215366 | 0.756 | 0.419 | 1.36E-25 | 9.20E-30 | |
| Layer 2/3 | iTBS | Uba52 | ubiquitin A-52 residue ribosomal protein fusion product 1 | 22186 | 0.710738094 | 0.848 | 0.64 | 5.15E-16 | 3.49E-20 | |
| Layer 2/3 | iTBS | Gstp1 | glutathione S-transferase, pi 1 | 14870 | 0.362570079 | 0.446 | 0.244 | 0.000247185 | 1.67E-08 | |
| Layer 2/3 | iTBS | Adcy1 | adenylate cyclase 1 | 432530 | 0.320730341 | 0.825 | 0.726 | 0.001107199 | 7.49E-08 | 1 |
| Layer 2/3 | iTBS | Bsg | basigin | 12215 | 0.287372453 | 0.977 | 0.931 | 0.000262858 | 1.78E-08 | |
| Layer 2/3 | iTBS | Rpl35 | ribosomal protein L35 | 66489 | 0.278652367 | 0.974 | 0.947 | 0.002592361 | 1.75E-07 | |
| Layer 2/3 | iTBS | Psd | pleckstrin and Sec7 domain containing | 73728 | 0.258455079 | 0.967 | 0.842 | 0.027841205 | 1.88E-06 | 1 |
| Layer 2/3 | iTBS | Nme7 | NME/NM23 family member 7 | 171567 | -0.401414215 | 0.195 | 0.363 | 0.030227141 | 2.05E-06 | |
| Layer 2/3 | iTBS | Malat1 | metastasis associated lung adenocarcinoma transcript 1 (non-coding RNA) | 72289 | -0.420450995 | 0.508 | 0.627 | 0.033937586 | 2.30E-06 | 1 |
| Layer 2/3 | iTBS | Bc1 | brain cytoplasmic RNA 1 | 100568459 | -0.680503187 | 0.551 | 0.726 | 3.47E-06 | 2.35E-10 | 1 |
| Layer 4 | iTBS | Gm10076 | ribosomal protein L41 pseudogene | 100126819 | 1.192706911 | 0.806 | 0.393 | 2.58E-32 | 1.74E-36 | |
| Layer 4 | iTBS | Uba52 | ubiquitin A-52 residue ribosomal protein fusion product 1 | 22186 | 0.794795851 | 0.909 | 0.721 | 2.43E-20 | 1.65E-24 | |
| Layer 4 | iTBS | Rpl35 | ribosomal protein L35 | 66489 | 0.431538062 | 0.987 | 0.943 | 6.57E-11 | 4.45E-15 | |
| Layer 4 | iTBS | Gstp1 | glutathione S-transferase, pi 1 | 14870 | 0.384704024 | 0.435 | 0.188 | 1.29E-05 | 8.70E-10 | |
| Layer 4 | iTBS | Atp1b1 | ATPase, Na+/K+ transporting, beta 1 polypeptide | 11931 | 0.277359798 | 1 | 1 | 1.43E-05 | 9.66E-10 | |
| Layer 4 | iTBS | Camk2n1 | calcium/calmodulin-dependent protein kinase II inhibitor 1 | 66259 | 0.265511261 | 1 | 1 | 1.96E-06 | 1.32E-10 | 1 |
| Layer 4 | iTBS | Snhg9 | small nucleolar RNA host gene 9 | 73474 | -0.264506064 | 0.098 | 0.271 | 0.000302944 | 2.05E-08 | 7 |
| Layer 4 | iTBS | Ndufa4 | Ndufa4, mitochondrial complex associated | 17992 | -0.266486561 | 1 | 1 | 0.003414701 | 2.31E-07 | |
| Layer 4 | iTBS | Tuba1b | tubulin, alpha 1B | 22143 | -0.297419376 | 0.938 | 0.987 | 0.000397833 | 2.69E-08 | |
| Layer 4 | iTBS | Bc1 | brain cytoplasmic RNA 1 | 100568459 | -0.418345608 | 0.355 | 0.563 | 0.001108452 | 7.50E-08 | 1 |
| Layer 4 | iTBS | Malat1 | metastasis associated lung adenocarcinoma transcript 1 (non-coding RNA) | 72289 | -0.575145043 | 0.466 | 0.681 | 1.05E-08 | 7.12E-13 | 1 |
| Layer 5 | iTBS | Gm10076 | ribosomal protein L41 pseudogene | 100126819 | 0.965664168 | 0.747 | 0.41 | 2.39E-25 | 1.62E-29 | |
| Layer 5 | iTBS | Uba52 | ubiquitin A-52 residue ribosomal protein fusion product 1 | 22186 | 0.52930806 | 0.869 | 0.753 | 4.14E-09 | 2.80E-13 | |
| Layer 5 | iTBS | Rpl35 | ribosomal protein L35 | 66489 | 0.274108668 | 0.991 | 0.96 | 0.000102629 | 6.94E-09 | |
| Layer 5 | iTBS | Cdk5r2 | cyclin-dependent kinase 5, regulatory subunit 2 (p39) | 12570 | -0.279423366 | 0.294 | 0.481 | 0.002802952 | 1.90E-07 | |
| Layer 5 | iTBS | Bc1 | brain cytoplasmic RNA 1 | 100568459 | -0.436738846 | 0.462 | 0.598 | 0.025236626 | 1.71E-06 | 1 |
| Layer 6 | iTBS | Gm10076 | ribosomal protein L41 pseudogene | 100126819 | 0.966468934 | 0.783 | 0.563 | 7.94E-22 | 5.37E-26 | |
| Layer 6 | iTBS | Uba52 | ubiquitin A-52 residue ribosomal protein fusion product 1 | 22186 | 0.690898808 | 0.911 | 0.837 | 7.11E-17 | 4.81E-21 | |
| Layer 6 | iTBS | Gstp1 | glutathione S-transferase, pi 1 | 14870 | 0.53023204 | 0.513 | 0.28 | 1.95E-08 | 1.32E-12 | |
| Layer 6 | iTBS | Rpl35 | ribosomal protein L35 | 66489 | 0.36946784 | 0.995 | 0.979 | 3.96E-10 | 2.68E-14 | |
| Layer 6 | iTBS | Rps20 | ribosomal protein S20 | 67427 | 0.281960799 | 1 | 0.997 | 7.63E-08 | 5.16E-12 | |
| Layer 6 | iTBS | Eno1 | enolase 1, alpha non-neuron | 13806 | 0.280086245 | 0.963 | 0.939 | 0.002393972 | 1.62E-07 | |
| Layer 6 | iTBS | Chd9 | chromodomain helicase DNA binding protein 9 | 109151 | 0.268061223 | 0.325 | 0.144 | 9.88E-05 | 6.69E-09 | |
| Layer 6 | iTBS | Rps10 | ribosomal protein S10 | 67097 | 0.250814978 | 0.997 | 0.979 | 0.003173887 | 2.15E-07 | |
| Layer 6 | iTBS | Tuba1b | tubulin, alpha 1B | 22143 | -0.283839882 | 0.945 | 0.989 | 0.000186732 | 1.26E-08 | |
| Layer 6 | iTBS | Bc1 | brain cytoplasmic RNA 1 | 100568459 | -0.515050145 | 0.429 | 0.667 | 1.14E-08 | 7.74E-13 | 1 |
| Layer 2/3 | cTBS | Gm10076 | ribosomal protein L41 pseudogene | 100126819 | 0.825540773 | 0.704 | 0.419 | 4.18E-15 | 2.83E-19 | |
| Layer 2/3 | cTBS | Uba52 | ubiquitin A-52 residue ribosomal protein fusion product 1 | 22186 | 0.49596171 | 0.737 | 0.64 | 0.001616433 | 1.09E-07 | |
| Layer 2/3 | cTBS | Rpl35 | ribosomal protein L35 | 66489 | 0.3468535 | 0.98 | 0.947 | 0.001526864 | 1.03E-07 | |
| Layer 2/3 | cTBS | Psd | pleckstrin and Sec7 domain containing | 73728 | 0.288558604 | 0.955 | 0.842 | 0.000261706 | 1.77E-08 | 1 |
| Layer 2/3 | cTBS | Ttr | transthyretin | 22139 | -0.252681043 | 0 | 0.122 | 0.000202003 | 1.37E-08 | 1 |
| Layer 2/3 | cTBS | Ikbkb | inhibitor of kappaB kinase beta | 16150 | -0.308652092 | 0.235 | 0.416 | 0.021137299 | 1.43E-06 | 7 |
| Layer 2/3 | cTBS | Bc1 | brain cytoplasmic RNA 1 | 100568459 | -0.934349372 | 0.348 | 0.726 | 2.36E-15 | 1.59E-19 | 1 |
| Layer 4 | cTBS | Gm10076 | ribosomal protein L41 pseudogene | 100126819 | 0.727480247 | 0.666 | 0.393 | 9.26E-12 | 6.27E-16 | |
| Layer 4 | cTBS | Rpl35 | ribosomal protein L35 | 66489 | 0.415216089 | 0.97 | 0.943 | 1.54E-06 | 1.04E-10 | |
| Layer 4 | cTBS | Camk2n1 | calcium/calmodulin-dependent protein kinase II inhibitor 1 | 66259 | 0.29066586 | 1 | 1 | 5.87E-10 | 3.97E-14 | 1 |
| Layer 4 | cTBS | Camk2n2 | calcium/calmodulin-dependent protein kinase II inhibitor 2 | 73047 | 0.275621869 | 0.908 | 0.843 | 0.015427158 | 1.04E-06 | 1 |
| Layer 4 | cTBS | Ncdn | neurochondrin | 26562 | 0.263969948 | 1 | 0.978 | 0.003990921 | 2.70E-07 | 1 |
| Layer 4 | cTBS | Ttr | transthyretin | 22139 | -0.294428991 | 0.015 | 0.162 | 7.06E-07 | 4.78E-11 | 1 |
| Layer 4 | cTBS | Bc1 | brain cytoplasmic RNA 1 | 100568459 | -0.553197494 | 0.266 | 0.563 | 2.10E-08 | 1.42E-12 | 1 |
| Layer 4 | cTBS | Pcp4 | Purkinje cell protein 4 | 18546 | -0.556809893 | 0.604 | 0.755 | 0.001595321 | 1.08E-07 | 1 |
| Layer 5 | cTBS | Gm10076 | ribosomal protein L41 pseudogene | 100126819 | 0.762610332 | 0.685 | 0.41 | 2.01E-17 | 1.36E-21 | |
| Layer 5 | cTBS | Uba52 | ubiquitin A-52 residue ribosomal protein fusion product 1 | 22186 | 0.399234149 | 0.787 | 0.753 | 0.044643899 | 3.02E-06 | |
| Layer 5 | cTBS | Rpl35 | ribosomal protein L35 | 66489 | 0.392826794 | 0.977 | 0.96 | 1.30E-09 | 8.77E-14 | |
| Layer 5 | cTBS | Bc1 | brain cytoplasmic RNA 1 | 100568459 | -0.553883679 | 0.364 | 0.598 | 1.01E-07 | 6.82E-12 | 1 |
| Layer 6 | cTBS | Gm10076 | ribosomal protein L41 pseudogene | 100126819 | 0.648563877 | 0.738 | 0.563 | 2.05E-12 | 1.39E-16 | |
| Layer 6 | cTBS | Gstp1 | glutathione S-transferase, pi 1 | 14870 | 0.463396118 | 0.498 | 0.28 | 6.39E-06 | 4.32E-10 | |
| Layer 6 | cTBS | Eno1 | enolase 1, alpha non-neuron | 13806 | 0.297044684 | 0.966 | 0.939 | 0.00125099 | 8.47E-08 | |
| Layer 6 | cTBS | Rpl35 | ribosomal protein L35 | 66489 | 0.287112276 | 0.972 | 0.979 | 0.012213146 | 8.26E-07 | |
| Layer 6 | cTBS | Bc1 | brain cytoplasmic RNA 1 | 100568459 | -0.643015917 | 0.357 | 0.667 | 2.24E-14 | 1.51E-18 | 1 |

**Data S7.** Complete list of enriched Gene Ontology (GO) terms for the significant differentially expressed genes (DEGs), identified in the different somatosensory (SS) cortex layers, following iTBS, and cTBS.

| SS layer | Stimulation | GO ID | GO term description | Number of genes | Gene symbols | p.value | q.value | Ontology Source | Code |
|---|---|---|---|---|---|---|---|---|---|
| Layer 4 | iTBS | GO:0022625 | cytosolic large ribosomal subunit | 2 | Uba52/Rpl35 | 4.66E-05 | 4.91E-05 | CC | 8 |
| Layer 4 | iTBS | GO:0022626 | cytosolic ribosome | 2 | Uba52/Rpl35 | 0.000153198 | 6.16E-05 | CC | 8 |
| Layer 4 | iTBS | GO:0015934 | large ribosomal subunit | 2 | Uba52/Rpl35 | 0.000175458 | 6.16E-05 | CC | 8 |
| Layer 4 | iTBS | GO:0044391 | ribosomal subunit | 2 | Uba52/Rpl35 | 0.00045655 | 0.000120145 | CC | 8 |
| Layer 4 | iTBS | GO:0005840 | ribosome | 2 | Uba52/Rpl35 | 0.000670615 | 0.000141182 | CC | 8 |
| Layer 4 | iTBS | GO:0003735 | structural constituent of ribosome | 2 | Uba52/Rpl35 | 0.000334805 | 0.001409707 | MF | 8 |
| Layer 5 | iTBS | GO:0043209 | myelin sheath | 3 | Uba52/Atp1b1/Tuba1b | 6.42E-05 | 0.001284023 | CC | 4 |
| Layer 5 | iTBS | GO:0022625 | cytosolic large ribosomal subunit | 2 | Uba52/Rpl35 | 0.000254233 | 0.002542327 | CC | 8 |
| Layer 6 | iTBS | GO:0022626 | cytosolic ribosome | 4 | Uba52/Rpl35/Rps20/Rps10 | 4.74E-08 | 3.50E-07 | CC | 8 |
| Layer 6 | iTBS | GO:0044391 | ribosomal subunit | 4 | Uba52/Rpl35/Rps20/Rps10 | 4.27E-07 | 1.18E-06 | CC | 8 |
| Layer 6 | iTBS | GO:0005840 | ribosome | 4 | Uba52/Rpl35/Rps20/Rps10 | 9.23E-07 | 1.70E-06 | CC | 8 |
| Layer 6 | iTBS | GO:0003735 | structural constituent of ribosome | 4 | Uba52/Rpl35/Rps20/Rps10 | 2.29E-07 | 3.61E-06 | MF | 8 |
| Layer 6 | iTBS | GO:0022627 | cytosolic small ribosomal subunit | 3 | Uba52/Rps20/Rps10 | 4.81E-07 | 1.18E-06 | CC | 8 |
| Layer 6 | iTBS | GO:0015935 | small ribosomal subunit | 3 | Uba52/Rps20/Rps10 | 2.42E-06 | 3.57E-06 | CC | 8 |
| Layer 6 | iTBS | GO:0043209 | myelin sheath | 3 | Uba52/Tuba1b/Eno1 | 4.69E-05 | 5.77E-05 | CC | 4 |
| Layer 6 | iTBS | GO:0022625 | cytosolic large ribosomal subunit | 2 | Uba52/Rpl35 | 0.000208302 | 0.000219265 | CC | 8 |
| Layer 6 | iTBS | GO:0015934 | large ribosomal subunit | 2 | Uba52/Rpl35 | 0.000778704 | 0.000717228 | CC | 8 |
| Layer 2/3 | cTBS | GO:0022625 | cytosolic large ribosomal subunit | 2 | Rpl35/Uba52 | 9.76e-05 | 0.001130322 | CC | 8 |
| Layer 2/3 | cTBS | GO:0022626 | cytosolic ribosome | 2 | Rpl35/Uba52 | 0.000320056 | 0.001414275 | CC | 8 |
| Layer 2/3 | cTBS | GO:0015934 | large ribosomal subunit | 2 | Rpl35/Uba52 | 0.000366426 | 0.001414275 | CC | 8 |
| Layer 2/3 | cTBS | GO:0044391 | ribosomal subunit | 2 | Rpl35/Uba52 | 0.000950161 | 0.002750465 | CC | 8 |
| Layer 2/3 | cTBS | GO:0005840 | ribosome | 2 | Rpl35/Uba52 | 0.001392962 | 0.003225807 | CC | 8 |
| Layer 4 | cTBS | GO:0006469 | negative regulation of protein kinase activity | 3 | Camk2n1/Pcp4/Camk2n2 | 1.84E-05 | 0.000775076 | BP | 9 |
| Layer 4 | cTBS | GO:0033673 | negative regulation of kinase activity | 3 | Camk2n1/Pcp4/Camk2n2 | 2.54E-05 | 0.000775076 | BP | 9 |
| Layer 4 | cTBS | GO:0051348 | negative regulation of transferase activity | 3 | Camk2n1/Pcp4/Camk2n2 | 3.81E-05 | 0.000775721 | BP | 9 |
| Layer 4 | cTBS | GO:0001933 | negative regulation of protein phosphorylation | 3 | Camk2n1/Pcp4/Camk2n2 | 9.25E-05 | 0.001411126 | BP | 9 |
| Layer 4 | cTBS | GO:0042326 | negative regulation of phosphorylation | 3 | Camk2n1/Pcp4/Camk2n2 | 0.000134375 | 0.001640788 | BP | 9 |
| Layer 4 | cTBS | GO:0010563 | negative regulation of phosphorus metabolic process | 3 | Camk2n1/Pcp4/Camk2n2 | 0.000199856 | 0.001743102 | BP | 9 |
| Layer 4 | cTBS | GO:0045936 | negative regulation of phosphate metabolic process | 3 | Camk2n1/Pcp4/Camk2n2 | 0.000199856 | 0.001743102 | BP | 9 |
| Layer 4 | cTBS | GO:0004860 | protein kinase inhibitor activity | 2 | Camk2n1/Camk2n2 | 0.000129846 | 0.000239546 | MF | 9 |
| Layer 4 | cTBS | GO:0019210 | kinase inhibitor activity | 2 | Camk2n1/Camk2n2 | 0.000151713 | 0.000239546 | MF | 9 |
| Layer 4 | cTBS | GO:0019887 | protein kinase regulator activity | 2 | Camk2n1/Camk2n2 | 0.00130069 | 0.001369148 | MF | 9 |
| Layer 4 | cTBS | GO:0019207 | kinase regulator activity | 2 | Camk2n1/Camk2n2 | 0.001750908 | 0.001382296 | MF | 9 |
| Layer 5 | cTBS | GO:0003735 | structural constituent of ribosome | 2 | Rpl35/Uba52 | 0.000201658 | 0.000212271 | MF | 8 |
| Layer 5 | cTBS | GO:0031386 | protein tag | 1 | Uba52 | 0.001686903 | 0.000887844 | MF | |

| Code key | High order function or process |
|---|---|
| 1 | synapse and/or synaptic plasticity |
| 2 | Neurotransmitter |
| 3 | Intrinsic plasticity and/or membrane excitability |
| 4 | Oligodendrocyte and/or myelin related |
| 5 | neurogenesis |
| 6 | neurotrophin |
| 7 | Cell death |
| 8 | Ribosome |
| 9 | ATP metabolic pathway |

**Data S8.** Differentially expressed genes (DEGs) identified in different cortical layers and brain regions in aged mice 3 h following intermittent (iTBS) or continuous theta burst stimulation (cTBS). Code for higher-order groups are as follows: 1 - synapse and/or synaptic plasticity; 2 - neurotransmitter; 3 - intrinsic plasticity and/or membrane excitability; 4 - oligodendrocyte and/or myelin related; 5 - neurogenesis; 6 - neurotrophin; 7 - inflammation/apoptosis.

| Brain reigon | Stimulation | Gene Symbol | Gene Name | EntrezID | Avg log2FoldChange | % expressed in stimulated samples | % expressed in sham samples | p.adjust value (Bonferroni correction) | p.value | Higher-order groups |
|---|---|---|---|---|---|---|---|---|---|---|
| Cortical Layer L2/3 | iTBS | Nr4a1 | nuclear receptor subfamily 4, group A, member 1 | 15370 | 0.334222124 | 0.553 | 0.42 | 0.000331119 | 2.2E-08 | 1 |
| Cortical Layer L2/3 | iTBS | Adcy1 | adenylate cyclase 1 | 432530 | 0.281144358 | 0.858 | 0.779 | 2.09E-05 | 1.4E-09 | 1 |
| Cortical Layer L2/3 | iTBS | Glul | glutamate-ammonia ligase (glutamine synthetase) | 14645 | -0.277222094 | 0.76 | 0.862 | 0.001298862 | 8.8E-08 | 1 |
| Cortical Layer L2/3 | iTBS | Cplx2 | complexin 2 | 12890 | -0.311164116 | 0.856 | 0.853 | 0.010851651 | 7.3E-07 | 1 |
| Cortical Layer L2/3 | iTBS | Cdk5r2 | cyclin-dependent kinase 5, regulatory subunit 2 (p39) | 12570 | -0.373759613 | 0.402 | 0.611 | 1.68E-10 | 1.1E-14 | 1 |
| Cortical Layer L2/3 | iTBS | Bc1 | brain cytoplasmic RNA 1 | 100568459 | -0.71989169 | 0.492 | 0.73 | 2.02E-18 | 1.4E-22 | 1 |
| Cortical Layer 4 | iTBS | Cdk5r2 | cyclin-dependent kinase 5, regulatory subunit 2 (p39) | 12570 | -0.261825698 | 0.372 | 0.517 | 0.033278761 | 2.3E-06 | 1 |
| Cortical Layer 4 | iTBS | Bc1 | brain cytoplasmic RNA 1 | 100568459 | -0.387897219 | 0.437 | 0.608 | 0.00019765 | 1.3E-08 | 1 |
| Cortical Layer 5 | iTBS | Pvalb | parvalbumin | 19293 | 0.291796575 | 0.896 | 0.818 | 2.56E-05 | 1.7E-09 | 1 |
| Cortical Layer 5 | iTBS | Syngr1 | synaptogyrin 1 | 20972 | -0.275389222 | 0.841 | 0.883 | 1.19E-05 | 8.0E-10 | 1 |
| Cortical Layer 5 | iTBS | Glul | glutamate-ammonia ligase (glutamine synthetase) | 14645 | -0.28424235 | 0.804 | 0.866 | 1.02E-08 | 6.9E-13 | 1 |
| Cortical Layer 5 | iTBS | Cdk5r2 | cyclin-dependent kinase 5, regulatory subunit 2 (p39) | 12570 | -0.309355067 | 0.37 | 0.553 | 2.55E-13 | 1.7E-17 | 1 |
| Cortical Layer 5 | iTBS | Cplx2 | complexin 2 | 12890 | -0.353548571 | 0.705 | 0.797 | 1.65E-10 | 1.1E-14 | 1 |
| Cortical Layer 5 | iTBS | Bc1 | brain cytoplasmic RNA 1 | 100568459 | -0.583355218 | 0.437 | 0.665 | 2.18E-24 | 1.5E-28 | 1 |
| Cortical Layer 6 | iTBS | Ttr | transthyretin | 22139 | 0.438710277 | 0.282 | 0.13 | 1.03E-06 | 6.9E-11 | 1 |
| Cortical Layer 6 | iTBS | Cplx2 | complexin 2 | 12890 | -0.263766222 | 0.851 | 0.899 | 0.002044561 | 1.4E-07 | 1 |
| Cortical Layer 6 | iTBS | Dnm1 | dynamin 1 | 13429 | -0.266230751 | 0.985 | 1 | 4.01E-05 | 2.7E-09 | 1 |
| Cortical Layer 6 | iTBS | Slc17a7 | solute carrier family 17 (sodium-dependent inorganic phosphate cotransporter), member 7 | 72961 | -0.268674902 | 0.995 | 0.994 | 0.000105398 | 7.1E-09 | 1 |
| Cortical Layer 6 | iTBS | Bc1 | brain cytoplasmic RNA 1 | 100568459 | -0.434433444 | 0.491 | 0.685 | 2.12E-09 | 1.4E-13 | 1 |
| Piriform Cortex | iTBS | Ttr | transthyretin | 22139 | 0.590664197 | 0.285 | 0.114 | 1.74E-08 | 1.2E-12 | 1 |
| Piriform Cortex | iTBS | Lynx1 | Ly6/neurotoxin 1 | 23936 | -0.268718044 | 0.291 | 0.413 | 0.038741175 | 2.6E-06 | 1 |
| Piriform Cortex | iTBS | Shank1 | SH3 and multiple ankyrin repeat domains 1 | 243961 | -0.269765778 | 0.386 | 0.539 | 0.001328322 | 9.0E-08 | 1 |
| Piriform Cortex | iTBS | Eef2 | eukaryotic translation elongation factor 2 | 13629 | -0.324199801 | 0.937 | 0.971 | 1.37E-06 | 9.3E-11 | 1 |
| Piriform Cortex | iTBS | Cplx2 | complexin 2 | 12890 | -0.361252302 | 0.816 | 0.844 | 0.000329245 | 2.2E-08 | 1 |
| Piriform Cortex | iTBS | Cdk5r2 | cyclin-dependent kinase 5, regulatory subunit 2 (p39) | 12570 | -0.385890004 | 0.338 | 0.554 | 1.01E-09 | 6.8E-14 | 1 |
| Piriform Cortex | iTBS | Syngr1 | synaptogyrin 1 | 20972 | -0.409655719 | 0.774 | 0.838 | 8.95E-06 | 6.1E-10 | 1 |
| Piriform Cortex | iTBS | Dnm1 | dynamin 1 | 13429 | -0.414065133 | 0.996 | 0.996 | 3.43E-14 | 2.3E-18 | 1 |
| Piriform Cortex | iTBS | Syn1 | synapsin I | 20964 | -0.422075828 | 0.933 | 0.947 | 4.84E-10 | 3.3E-14 | 1 |
| Piriform Cortex | iTBS | Bc1 | brain cytoplasmic RNA 1 | 100568459 | -0.431876265 | 0.629 | 0.773 | 0.000814112 | 5.5E-08 | 1 |
| Piriform Cortex | iTBS | Slc17a7 | solute carrier family 17 (sodium-dependent inorganic phosphate cotransporter), member 7 | 72961 | -0.445626054 | 0.996 | 0.998 | 2.56E-11 | 1.7E-15 | 1 |
| White Matter Tracts | iTBS | Ttr | transthyretin | 22139 | 0.609096865 | 0.3 | 0.078 | 3.26E-17 | 2.2E-21 | 1 |
| White Matter Tracts | iTBS | Bc1 | brain cytoplasmic RNA 1 | 100568459 | -0.433049208 | 0.816 | 0.885 | 0.017471008 | 1.2E-06 | 1 |
| Caudoputamen | iTBS | Ttr | transthyretin | 22139 | 0.615804318 | 0.258 | 0.141 | 4.31E-10 | 2.9E-14 | 1 |
| Caudoputamen | iTBS | Glul | glutamate-ammonia ligase (glutamine synthetase) | 14645 | -0.266010832 | 0.738 | 0.835 | 5.56E-09 | 3.8E-13 | 1 |
| Caudoputamen | iTBS | Bc1 | brain cytoplasmic RNA 1 | 100568459 | -0.284200548 | 0.537 | 0.693 | 1.03E-08 | 7.0E-13 | 1 |
| Caudoputamen | iTBS | Dnm1 | dynamin 1 | 13429 | -0.303727691 | 0.849 | 0.944 | 1.04E-13 | 7.1E-18 | 1 |
| Lateral Septal Complex | iTBS | Kalrr | kalirin, RhoGEF kinase | 545156 | -0.252111211 | 0.27 | 0.449 | 0.016802031 | 1.1E-06 | 1 |
| Lateral Septal Complex | iTBS | Cdk5r2 | cyclin-dependent kinase 5, regulatory subunit 2 (p39) | 12570 | -0.29895976 | 0.232 | 0.441 | 5.13E-05 | 3.5E-09 | 1 |
| Lateral Septal Complex | iTBS | Ttr | transthyretin | 22139 | -1.617776097 | 0.174 | 0.321 | 0.023932756 | 1.6E-06 | 1 |
| Striatum Ventral Region | iTBS | Bc1 | brain cytoplasmic RNA 1 | 100568459 | -0.384592556 | 0.676 | 0.83 | 6.83E-07 | 4.9E-52 | 1 |
| Cortical Layer L2/3 | cTBS | Bc1 | brain cytoplasmic RNA 1 | 100568459 | -0.888235077 | 0.367 | 0.73 | 1.87E-28 | 1.3E-32 | 1 |
| Cortical Layer L2/3 | cTBS | Ttr | transthyretin | 22139 | -1.38847059 | 0.007 | 0.216 | 5.67E-18 | 3.8E-22 | 1 |
| Cortical Layer 4 | cTBS | Camk2n1 | calcium/calmodulin-dependent protein kinase II inhibitor 1 | 66259 | 0.256554719 | 1 | 1 | 8.47E-12 | 5.7E-16 | 1 |
| Cortical Layer 4 | cTBS | Ttr | transthyretin | 22139 | -0.288348911 | 0.01 | 0.156 | 2.02E-12 | 1.4E-16 | 1 |
| Cortical Layer 4 | cTBS | Pcp4 | Purkinje cell protein 4 | 18546 | -0.399225391 | 0.681 | 0.789 | 0.007257433 | 4.9E-07 | 1 |
| Cortical Layer 4 | cTBS | Bc1 | brain cytoplasmic RNA 1 | 100568459 | -0.552946373 | 0.323 | 0.608 | 6.99E-13 | 4.7E-17 | 1 |
| Cortical Layer 5 | cTBS | Ttr | transthyretin | 22139 | -0.435559088 | 0.015 | 0.178 | 3.20E-26 | 2.2E-30 | 1 |
| Cortical Layer 5 | cTBS | Bc1 | brain cytoplasmic RNA 1 | 100568459 | -0.701902569 | 0.326 | 0.665 | 1.52E-43 | 1.0E-47 | 1 |
| Cortical Layer 6 | cTBS | Ttr | transthyretin | 22139 | -0.282802181 | 0.011 | 0.13 | 1.77E-10 | 1.2E-14 | 1 |
| Cortical Layer 6 | cTBS | Cdk5r2 | cyclin-dependent kinase 5, regulatory subunit 2 (p39) | 12570 | -0.287295226 | 0.29 | 0.4 | 0.041982987 | 2.8E-06 | 1 |
| Cortical Layer 6 | cTBS | Bc1 | brain cytoplasmic RNA 1 | 100568459 | -0.645763393 | 0.366 | 0.685 | 1.07E-24 | 7.2E-29 | 1 |
| Piriform Cortex | cTBS | Syn1 | synapsin I | 20964 | -0.2807775 | 0.946 | 0.947 | 0.017678469 | 1.2E-06 | 1 |
| Piriform Cortex | cTBS | Bc1 | brain cytoplasmic RNA 1 | 100568459 | -0.967071131 | 0.388 | 0.773 | 6.07E-32 | 4.1E-36 | 1 |
| White Matter Tracts | cTBS | Bc1 | brain cytoplasmic RNA 1 | 100568459 | -0.834899137 | 0.713 | 0.885 | 1.48E-22 | 1.0E-26 | 1 |
| Caudoputamen | cTBS | Slc17a7 | solute carrier family 17 (sodium-dependent inorganic phosphate cotransporter), member 7 | 72961 | -0.272338813 | 0.196 | 0.361 | 1.47E-14 | 9.9E-19 | 1 |
| Caudoputamen | cTBS | Ttr | transthyretin | 22139 | -0.326153983 | 0.002 | 0.141 | 4.66E-33 | 3.2E-37 | 1 |
| Caudoputamen | cTBS | Bc1 | brain cytoplasmic RNA 1 | 100568459 | -0.704317676 | 0.342 | 0.693 | 2.11E-61 | 1.4E-65 | 1 |
| Lateral Septal Complex | cTBS | Cnih2 | cornichon family AMPA receptor auxiliary protein 2 | 12794 | -0.289620486 | 0.92 | 0.961 | 0.010041113 | 6.8E-07 | 1 |
| Lateral Septal Complex | cTBS | Gria1 | glutamate receptor, ionotropic, AMPA1 (alpha 1) | 14799 | -0.334501905 | 0.691 | 0.815 | 0.003208812 | 2.2E-07 | 1 |
| Lateral Septal Complex | cTBS | Bc1 | brain cytoplasmic RNA 1 | 100568459 | -1.114948558 | 0.585 | 0.909 | 2.59E-32 | 1.8E-36 | 1 |
| Lateral Septal Complex | cTBS | Ttr | transthyretin | 22139 | -1.951188266 | 0.101 | 0.321 | 6.79E-10 | 4.6E-14 | 1 |
| Striatum Ventral Region | cTBS | Malat1 | metastasis associated lung adenocarcinoma transcript 1 (non-coding RNA) | 72289 | -0.307512671 | 0.573 | 0.643 | 0.017126764 | 1.2E-06 | 1 |
| Striatum Ventral Region | cTBS | Bc1 | brain cytoplasmic RNA 1 | 100568459 | -0.926359105 | 0.48 | 0.83 | 8.05E-52 | 5.4E-56 | 1 |
| Lateral Septal Complex | iTBS | Kcnq2 | potassium voltage-gated channel, subfamily Q, member 2 | 16536 | -0.276817724 | 0.46 | 0.621 | 0.044984929 | 3.0E-06 | 3 |
| Striatum Ventral Region | iTBS | Atp1a3 | ATPase, Na+/K+ transporting, alpha 3 polypeptide | 232975 | -0.266323597 | 0.735 | 0.818 | 6.50E-06 | 4.9E-52 | 3 |
| Piriform Cortex | cTBS | Atp1a3 | ATPase, Na+/K+ transporting, alpha 3 polypeptide | 232975 | -0.323809229 | 0.753 | 0.863 | 3.61E-06 | 2.4E-10 | 3 |
| Piriform Cortex | iTBS | Nat8l | N-acetyltransferase 8-like | 269642 | -0.261190961 | 0.583 | 0.684 | 0.036544555 | 2.5E-06 | 4 |
| White Matter Tracts | cTBS | Bcas1 | brain enriched myelin associated protein 1 | 76960 | 0.348352484 | 0.695 | 0.545 | 0.000721123 | 4.9E-08 | 4 |
| White Matter Tracts | cTBS | Mbp | myelin basic protein | 17196 | 0.338416104 | 1 | 1 | 3.39E-09 | 2.3E-13 | 4 |
| White Matter Tracts | cTBS | Mobp | myelin-associated oligodendrocytic basic protein | 17433 | 0.317797139 | 0.998 | 1 | 7.71E-06 | 5.2E-10 | 4 |
| Piriform Cortex | cTBS | Ptgds | prostaglandin D2 synthase (brain) | 19215 | -0.279582549 | 0.642 | 0.808 | 0.006737367 | 4.6E-07 | 6 |
| Striatum Ventral Region | cTBS | Ptgds | prostaglandin D2 synthase (brain) | 19215 | -0.353222369 | 0.694 | 0.79 | 0.024112355 | 1.6E-06 | 6 |
| Cortical Layer L2/3 | iTBS | Ndufa4 | Ndufa4, mitochondrial complex associated | 17992 | -0.267810222 | 1 | 1 | 7.10E-07 | 4.8E-11 | 7 |
| Cortical Layer 5 | iTBS | Tuba1b | tubulin, alpha 1B | 22143 | -0.301187772 | 0.967 | 0.985 | 1.71E-18 | 1.2E-22 | 7 |
| Piriform Cortex | iTBS | Ikbkb | inhibitor of kappaB kinase beta | 16150 | -0.307852202 | 0.245 | 0.432 | 1.03E-06 | 6.9E-11 | 7 |
| White Matter Tracts | iTBS | Snhg8 | small nucleolar RNA host gene 8 | 69895 | 0.308124984 | 0.5 | 0.237 | 6.81E-12 | 4.6E-16 | 7 |
| White Matter Tracts | iTBS | Banf1 | BAF nuclear assembly factor 1 | 23825 | 0.252885736 | 0.493 | 0.295 | 1.16E-05 | 7.8E-10 | 7 |
| Cortical Layer L2/3 | cTBS | Ikbkb | inhibitor of kappaB kinase beta | 16150 | -0.27364604 | 0.258 | 0.452 | 5.71E-06 | 3.9E-10 | 7 |
| Piriform Cortex | cTBS | Ikbkb | inhibitor of kappaB kinase beta | 16150 | -0.260378449 | 0.275 | 0.432 | 0.001712617 | 1.2E-07 | 7 |
| Cortical Layer L2/3 | iTBS | Gm10076 | ribosomal protein L41 pseudogene | 100126819 | 1.230542179 | 0.776 | 0.348 | 7.79E-63 | 5.3E-67 | |
| Cortical Layer L2/3 | iTBS | Uba52 | ubiquitin A-52 residue ribosomal protein fusion product 1 | 22186 | 0.823819101 | 0.841 | 0.62 | 1.14E-38 | 7.7E-43 | |
| Cortical Layer L2/3 | iTBS | Gstp1 | glutathione S-transferase, pi 1 | 14870 | 0.398236375 | 0.467 | 0.236 | 7.66E-13 | 5.2E-17 | |
| Cortical Layer L2/3 | iTBS | Rpl35 | ribosomal protein L35 | 66489 | 0.369907549 | 0.974 | 0.9 | 1.68E-13 | 1.1E-17 | |
| Cortical Layer L2/3 | iTBS | Phyhip | phytanoyl-CoA hydroxylase interacting protein | 105653 | -0.271420143 | 0.537 | 0.66 | 0.000878778 | 5.9E-08 | |
| Cortical Layer L2/3 | iTBS | Ivns1abp | influenza virus NS1A binding protein | 117198 | -0.300612084 | 0.451 | 0.558 | 0.017918876 | 1.2E-06 | |
| Cortical Layer L2/3 | iTBS | Nme7 | NME/NM23 family member 7 | 171567 | -0.539648195 | 0.24 | 0.416 | 5.40E-07 | 3.7E-11 | |
| Cortical Layer 4 | iTBS | Gm10076 | ribosomal protein L41 pseudogene | 100126819 | 1.093702705 | 0.821 | 0.406 | 3.67E-45 | 2.5E-49 | |
| Cortical Layer 4 | iTBS | Uba52 | ubiquitin A-52 residue ribosomal protein fusion product 1 | 22186 | 0.70093592 | 0.911 | 0.731 | 2.79E-23 | 1.9E-27 | |
| Cortical Layer 4 | iTBS | Rpl35 | ribosomal protein L35 | 66489 | 0.359278949 | 0.982 | 0.953 | 2.70E-11 | 1.8E-15 | |
| Cortical Layer 4 | iTBS | Gstp1 | glutathione S-transferase, pi 1 | 14870 | 0.346288988 | 0.419 | 0.203 | 4.67E-07 | 3.2E-11 | |
| Cortical Layer 4 | iTBS | Atp1b1 | ATPase, Na+/K+ transporting, beta 1 polypeptide | 11931 | 0.274732665 | 1 | 1 | 1.13E-08 | 7.6E-13 | |
| Cortical Layer 4 | iTBS | Lars2 | leucyl-tRNA synthetase, mitochondrial | 102436 | -0.285776713 | 0.256 | 0.394 | 0.024616403 | 1.7E-06 | |
| Cortical Layer 4 | iTBS | Nme7 | NME/NM23 family member 7 | 171567 | -0.467957477 | 0.276 | 0.436 | 0.000403714 | 2.7E-08 | |
| Cortical Layer 4 | iTBS | Malat1 | metastasis associated lung adenocarcinoma transcript 1 (non-coding RNA) | 72289 | -0.514632877 | 0.494 | 0.65 | 1.24E-08 | 8.4E-13 | |
| Cortical Layer 5 | iTBS | Gm10076 | ribosomal protein L41 pseudogene | 100126819 | 1.229509494 | 0.758 | 0.341 | 4.02E-100 | 3E-104 | |
| Cortical Layer 5 | iTBS | Uba52 | ubiquitin A-52 residue ribosomal protein fusion product 1 | 22186 | 0.738317653 | 0.865 | 0.685 | 1.40E-52 | 9.5E-57 | |
| Cortical Layer 5 | iTBS | Rpl35 | ribosomal protein L35 | 66489 | 0.370962746 | 0.987 | 0.918 | 6.52E-24 | 4.4E-28 | |
| Cortical Layer 5 | iTBS | Gstp1 | glutathione S-transferase, pi 1 | 14870 | 0.346645918 | 0.483 | 0.313 | 2.03E-12 | 1.4E-16 | |
| Cortical Layer 5 | iTBS | Chd9 | chromodomain helicase DNA binding protein 9 | 109151 | 0.25308307 | 0.385 | 0.207 | 4.57E-12 | 3.1E-16 | |
| Cortical Layer 5 | iTBS | Dynll2 | dynein light chain LC8-type 2 | 68097 | -0.252237304 | 0.967 | 0.975 | 1.01E-05 | 6.8E-10 | |
| Cortical Layer 5 | iTBS | Hpcal4 | hippocalcin-like 4 | 170638 | -0.267315518 | 0.582 | 0.703 | 1.98E-06 | 1.3E-10 | |
| Cortical Layer 5 | iTBS | Nme7 | NME/NM23 family member 7 | 171567 | -0.40447592 | 0.313 | 0.408 | 0.021411937 | 1.4E-06 | |
| Cortical Layer 6 | iTBS | Gm10076 | ribosomal protein L41 pseudogene | 100126819 | 0.97998495 | 0.782 | 0.559 | 1.13E-37 | 7.6E-42 | |
| Cortical Layer 6 | iTBS | Uba52 | ubiquitin A-52 residue ribosomal protein fusion product 1 | 22186 | 0.661968159 | 0.91 | 0.832 | 2.28E-27 | 1.5E-31 | |
| Cortical Layer 6 | iTBS | Gstp1 | glutathione S-transferase, pi 1 | 14870 | 0.518666373 | 0.53 | 0.285 | 4.13E-15 | 2.8E-19 | |
| Cortical Layer 6 | iTBS | Rpl35 | ribosomal protein L35 | 66489 | 0.315851569 | 0.992 | 0.985 | 2.36E-12 | 1.6E-16 | |
| Cortical Layer 6 | iTBS | Chd9 | chromodomain helicase DNA binding protein 9 | 109151 | 0.250144205 | 0.298 | 0.138 | 1.55E-06 | 1.0E-10 | |
| Cortical Layer 6 | iTBS | Map2 | microtubule-associated protein 2 | 17756 | -0.255229472 | 0.741 | 0.831 | 0.00060652 | 4.1E-08 | |
| Cortical Layer 6 | iTBS | Eef1a2 | eukaryotic translation elongation factor 1 alpha 2 | 13628 | -0.271544865 | 0.989 | 0.998 | 1.47E-05 | 9.9E-10 | |
| Cortical Layer 6 | iTBS | Hpcal4 | hippocalcin-like 4 | 170638 | -0.272302736 | 0.768 | 0.842 | 0.007162921 | 4.8E-07 | |
| Piriform Cortex | iTBS | Gm10076 | ribosomal protein L41 pseudogene | 100126819 | 1.377621314 | 0.751 | 0.423 | 1.48E-49 | 1.0E-53 | |
| Piriform Cortex | iTBS | Uba52 | ubiquitin A-52 residue ribosomal protein fusion product 1 | 22186 | 0.844989401 | 0.878 | 0.701 | 2.69E-35 | 1.8E-39 | |
| Piriform Cortex | iTBS | Rpl35 | ribosomal protein L35 | 66489 | 0.404752236 | 0.985 | 0.958 | 1.61E-17 | 1.1E-21 | |
| Piriform Cortex | iTBS | Rps28 | ribosomal protein S28 | 54127 | 0.399194726 | 0.908 | 0.857 | 5.16E-08 | 3.5E-12 | |
| Piriform Cortex | iTBS | Gstp1 | glutathione S-transferase, pi 1 | 14870 | 0.354232782 | 0.528 | 0.354 | 2.96E-06 | 2.0E-10 | |
| Piriform Cortex | iTBS | Rps27 | ribosomal protein S27 | 57294 | 0.312121998 | 0.948 | 0.905 | 1.68E-06 | 1.1E-10 | |
| Piriform Cortex | iTBS | Rps21 | ribosomal protein S21 | 66481 | 0.309049681 | 1 | 1 | 7.26E-17 | 4.9E-21 | |
| Piriform Cortex | iTBS | Rps29 | ribosomal protein S29 | 20090 | 0.306562748 | 0.994 | 0.985 | 4.30E-05 | 2.9E-09 | |
| Piriform Cortex | iTBS | Cox7c | cytochrome c oxidase subunit 7C | 12867 | 0.298461796 | 1 | 0.998 | 5.62E-14 | 3.8E-18 | |
| Piriform Cortex | iTBS | Rpl35a | ribosomal protein L35A | 57808 | 0.285960325 | 1 | 0.985 | 1.08E-09 | 7.3E-14 | |
| Piriform Cortex | iTBS | Rps26 | ribosomal protein S26 | 27370 | 0.269601214 | 0.977 | 0.954 | 2.84E-06 | 1.9E-10 | |
| Piriform Cortex | iTBS | Rpl9 | ribosomal protein L9 | 20005 | 0.259279456 | 1 | 0.987 | 2.92E-10 | 2.0E-14 | |
| Piriform Cortex | iTBS | Rps27a | ribosomal protein S27A | 78294 | 0.254026489 | 1 | 0.996 | 1.43E-10 | 9.7E-15 | |
| Piriform Cortex | iTBS | Atp6v0a1 | ATPase, H+ transporting, lysosomal V0 subunit A1 | 11975 | -0.250144227 | 0.786 | 0.846 | 0.01914904 | 1.3E-06 | |
| Piriform Cortex | iTBS | Emc10 | ER membrane protein complex subunit 10 | 69683 | -0.255475018 | 0.797 | 0.863 | 0.000122005 | 8.3E-09 | |
| Piriform Cortex | iTBS | Atp1a3 | ATPase, Na+/K+ transporting, alpha 3 polypeptide | 232975 | -0.256273819 | 0.795 | 0.863 | 0.005698402 | 3.9E-07 | |
| Piriform Cortex | iTBS | Eno2 | enolase 2, gamma neuronal | 13807 | -0.265189061 | 0.956 | 0.971 | 0.003479113 | 2.4E-07 | |
| Piriform Cortex | iTBS | Uqcrc1 | ubiquinol-cytochrome c reductase core protein 1 | 22273 | -0.26543376 | 0.778 | 0.861 | 0.002256156 | 1.5E-07 | |
| Piriform Cortex | iTBS | Golga7b | golgi autoantigen, golgin subfamily a, 7B | 71146 | -0.272646814 | 0.442 | 0.571 | 0.00616993 | 4.2E-07 | |
| Piriform Cortex | iTBS | Faim2 | Fas apoptotic inhibitory molecule 2 | 72393 | -0.280629639 | 0.734 | 0.785 | 0.001188851 | 8.0E-08 | |
| Piriform Cortex | iTBS | Epn1 | epsin 1 | 13854 | -0.282835387 | 0.826 | 0.874 | 0.00390282 | 2.6E-07 | |
| Piriform Cortex | iTBS | Dync1h1 | dynein cytoplasmic 1 heavy chain 1 | 13424 | -0.306294777 | 0.836 | 0.891 | 0.000116051 | 7.9E-09 | |
| Piriform Cortex | iTBS | Diras2 | DIRAS family, GTP-binding RAS-like 2 | 68203 | -0.325331392 | 0.755 | 0.815 | 0.002439183 | 1.7E-07 | |
| Piriform Cortex | iTBS | Phyhip | phytanoyl-CoA hydroxylase interacting protein | 105653 | -0.326178554 | 0.501 | 0.629 | 8.52E-05 | 5.8E-09 | |
| Piriform Cortex | iTBS | Sptbn2 | spectrin beta, non-erythrocytic 2 | 20743 | -0.33193069 | 0.805 | 0.844 | 1.30E-05 | 8.8E-10 | |
| Piriform Cortex | iTBS | Tuba1b | tubulin, alpha 1B | 22143 | -0.359977913 | 0.962 | 0.989 | 3.45E-13 | 2.3E-17 | |
| Piriform Cortex | iTBS | Dynll2 | dynein light chain LC8-type 2 | 68097 | -0.405864738 | 0.933 | 0.962 | 1.35E-12 | 9.1E-17 | |
| Piriform Cortex | iTBS | Eef1a2 | eukaryotic translation elongation factor 1 alpha 2 | 13628 | -0.454289962 | 0.983 | 0.994 | 3.90E-15 | 2.6E-19 | |
| White Matter Tracts | iTBS | Gm10076 | ribosomal protein L41 pseudogene | 100126819 | 0.847062151 | 0.975 | 0.887 | 2.48E-46 | 1.7E-50 | |
| White Matter Tracts | iTBS | Uba52 | ubiquitin A-52 residue ribosomal protein fusion product 1 | 22186 | 0.57659486 | 0.991 | 0.958 | 1.98E-23 | 1.3E-27 | |
| White Matter Tracts | iTBS | Gstp1 | glutathione S-transferase, pi 1 | 14870 | 0.538358018 | 0.825 | 0.597 | 1.39E-12 | 9.4E-17 | |
| White Matter Tracts | iTBS | Rps28 | ribosomal protein S28 | 54127 | 0.334807146 | 0.987 | 0.995 | 1.79E-06 | 1.2E-10 | |
| White Matter Tracts | iTBS | Rps21 | ribosomal protein S21 | 66481 | 0.327145778 | 1 | 1 | 3.90E-11 | 2.6E-15 | |
| White Matter Tracts | iTBS | Rps24 | ribosomal protein S24 | 20088 | 0.312858966 | 1 | 1 | 1.35E-08 | 9.1E-13 | |
| White Matter Tracts | iTBS | Enpp2 | ectonucleotide pyrophosphatase/phosphodiesterase 2 | 18606 | 0.282580977 | 0.666 | 0.468 | 0.012315097 | 8.3E-07 | |
| White Matter Tracts | iTBS | Rps10 | ribosomal protein S10 | 67097 | 0.274498159 | 0.998 | 1 | 6.58E-06 | 4.5E-10 | |
| White Matter Tracts | iTBS | Rpl35a | ribosomal protein L35A | 57808 | 0.274475287 | 1 | 1 | 2.00E-05 | 1.4E-09 | |
| White Matter Tracts | iTBS | Sf3b1 | splicing factor 3b, subunit 1 | 81898 | 0.273697497 | 0.384 | 0.173 | 1.53E-09 | 1.0E-13 | |
| White Matter Tracts | iTBS | Crym | crystallin, mu | 12971 | 0.270578122 | 0.197 | 0.053 | 4.40E-09 | 3.0E-13 | |
| White Matter Tracts | iTBS | Cab39 | calcium binding protein 39 | 12283 | 0.269462269 | 0.307 | 0.108 | 4.10E-11 | 2.8E-15 | |
| White Matter Tracts | iTBS | Rpl23 | ribosomal protein L23 | 65019 | 0.26928313 | 1 | 1 | 8.69E-08 | 5.9E-12 | |
| White Matter Tracts | iTBS | Hexb | hexosaminidase B | 15212 | 0.26767016 | 0.44 | 0.222 | 3.43E-07 | 2.3E-11 | |
| White Matter Tracts | iTBS | Rpl30 | ribosomal protein L30 | 19946 | 0.267108506 | 1 | 1 | 9.13E-05 | 6.2E-09 | |
| White Matter Tracts | iTBS | Cct3 | chaperonin containing Tcp1, subunit 3 (gamma) | 12462 | 0.257764942 | 0.439 | 0.213 | 8.20E-10 | 5.5E-14 | |
| White Matter Tracts | iTBS | Cd47 | CD47 antigen (Rh-related antigen, integrin-associated signal transducer) | 16423 | 0.25602735 | 0.319 | 0.128 | 8.88E-09 | 6.0E-13 | |
| White Matter Tracts | iTBS | Rps12 | ribosomal protein S12 | 20042 | 0.250664118 | 1 | 0.998 | 0.000138898 | 9.4E-09 | |
| White Matter Tracts | iTBS | Snx3 | sorting nexin 3 | 54198 | -0.256664141 | 0.791 | 0.877 | 0.002128645 | 1.4E-07 | |
| Caudoputamen | iTBS | Gm10076 | ribosomal protein L41 pseudogene | 100126819 | 1.504442045 | 0.811 | 0.463 | 8.87E-131 | 6E-135 | |
| Caudoputamen | iTBS | Uba52 | ubiquitin A-52 residue ribosomal protein fusion product 1 | 22186 | 0.904690012 | 0.883 | 0.78 | 4.63E-81 | 3.1E-85 | |
| Caudoputamen | iTBS | Gstp1 | glutathione S-transferase, pi 1 | 14870 | 0.632915936 | 0.612 | 0.27 | 5.38E-56 | 3.6E-60 | |
| Caudoputamen | iTBS | Rpl35 | ribosomal protein L35 | 66489 | 0.36964948 | 0.985 | 0.965 | 1.36E-31 | 9.2E-36 | |
| Caudoputamen | iTBS | Adora2a | adenosine A2a receptor | 11540 | 0.300944336 | 0.78 | 0.614 | 1.49E-13 | 1.0E-17 | |
| Caudoputamen | iTBS | Cnr1 | cannabinoid receptor 1 (brain) | 12801 | 0.292881538 | 0.681 | 0.532 | 3.39E-09 | 2.3E-13 | |
| Caudoputamen | iTBS | Rps21 | ribosomal protein S21 | 66481 | 0.283112319 | 1 | 1 | 4.45E-41 | 3.0E-45 | |
| Caudoputamen | iTBS | Ppp1r1b | protein phosphatase 1, regulatory inhibitor subunit 1B | 19049 | 0.261919919 | 0.997 | 0.991 | 2.19E-21 | 1.5E-25 | |
| Caudoputamen | iTBS | Rpl35a | ribosomal protein L35A | 57808 | 0.252392802 | 1 | 0.996 | 6.67E-24 | 4.5E-28 | |
| Caudoputamen | iTBS | Tuba1b | tubulin, alpha 1B | 22143 | -0.256390985 | 0.671 | 0.833 | 1.34E-11 | 9.1E-16 | |
| Caudoputamen | iTBS | Olfm1 | olfactomedin 1 | 56177 | -0.262932448 | 0.6 | 0.764 | 1.48E-12 | 1.0E-16 | |
| Caudoputamen | iTBS | Vsnl1 | visinin-like 1 | 26950 | -0.281697653 | 0.333 | 0.544 | 1.47E-17 | 1.0E-21 | |
| Caudoputamen | iTBS | Hpcal4 | hippocalcin-like 4 | 170638 | -0.289905788 | 0.525 | 0.596 | 1.98E-06 | 1.3E-10 | |
| Lateral Septal Complex | iTBS | Gm10076 | ribosomal protein L41 pseudogene | 100126819 | 1.23638845 | 0.646 | 0.496 | 2.39E-15 | 1.6E-19 | |
| Lateral Septal Complex | iTBS | Uba52 | ubiquitin A-52 residue ribosomal protein fusion product 1 | 22186 | 0.752594811 | 0.833 | 0.705 | 7.13E-16 | 4.8E-20 | |
| Lateral Septal Complex | iTBS | Rpl35 | ribosomal protein L35 | 66489 | 0.444890654 | 0.997 | 0.948 | 3.81E-13 | 2.6E-17 | |
| Lateral Septal Complex | iTBS | Rps28 | ribosomal protein S28 | 54127 | 0.333394575 | 0.977 | 0.958 | 5.20E-05 | 3.5E-09 | |
| Lateral Septal Complex | iTBS | Rps29 | ribosomal protein S29 | 20090 | 0.310903719 | 1 | 1 | 0.00011808 | 8.0E-09 | |
| Lateral Septal Complex | iTBS | Hectd4 | HECT domain E3 ubiquitin protein ligase 4 | 269700 | -0.253133208 | 0.334 | 0.514 | 0.019764442 | 1.3E-06 | |
| Lateral Septal Complex | iTBS | Parp6 | poly (ADP-ribose) polymerase family, member 6 | 67287 | -0.256255699 | 0.232 | 0.41 | 0.005951657 | 4.0E-07 | |
| Lateral Septal Complex | iTBS | Sgk1 | serum/glucocorticoid regulated kinase 1 | 20393 | -0.276265073 | 0.113 | 0.287 | 0.000143616 | 9.7E-09 | |
| Lateral Septal Complex | iTBS | Luzp2 | leucine zipper protein 2 | 233271 | -0.287605523 | 0.402 | 0.561 | 0.016368979 | 1.1E-06 | |
| Lateral Septal Complex | iTBS | Phyhip | phytanoyl-CoA hydroxylase interacting protein | 105653 | -0.317131003 | 0.289 | 0.483 | 0.000450812 | 3.1E-08 | |
| Lateral Septal Complex | iTBS | Prkca | protein kinase C, alpha | 18750 | -0.338440643 | 0.264 | 0.426 | 0.017054536 | 1.2E-06 | |
| Striatum Ventral Region | iTBS | Gm10076 | ribosomal protein L41 pseudogene | 100126819 | 1.338143498 | 0.733 | 0.59 | 7.19E-48 | 4.9E-52 | |
| Striatum Ventral Region | iTBS | Uba52 | ubiquitin A-52 residue ribosomal protein fusion product 1 | 22186 | 0.801778525 | 0.879 | 0.832 | 9.45E-38 | 4.9E-52 | |
| Striatum Ventral Region | iTBS | Gstp1 | glutathione S-transferase, pi 1 | 14870 | 0.478605517 | 0.584 | 0.368 | 7.52E-17 | 4.9E-52 | |
| Striatum Ventral Region | iTBS | Rpl35 | ribosomal protein L35 | 66489 | 0.305537969 | 0.984 | 0.972 | 2.08E-15 | 4.9E-52 | |
| Striatum Ventral Region | iTBS | Rps28 | ribosomal protein S28 | 54127 | 0.270465407 | 0.96 | 0.964 | 0.00011676 | 4.9E-52 | |
| Striatum Ventral Region | iTBS | Actb | actin, beta | 11461 | -0.256477526 | 0.999 | 0.998 | 1.64E-07 | 4.9E-52 | |
| Cortical Layer L2/3 | cTBS | Gm10076 | ribosomal protein L41 pseudogene | 100126819 | 1.072233836 | 0.749 | 0.348 | 2.82E-49 | 1.9E-53 | |
| Cortical Layer L2/3 | cTBS | Uba52 | ubiquitin A-52 residue ribosomal protein fusion product 1 | 22186 | 0.577826732 | 0.754 | 0.62 | 1.12E-14 | 7.6E-19 | |
| Cortical Layer L2/3 | cTBS | Rpl35 | ribosomal protein L35 | 66489 | 0.478474403 | 0.974 | 0.9 | 8.76E-19 | 5.9E-23 | |
| Cortical Layer L2/3 | cTBS | Gstp1 | glutathione S-transferase, pi 1 | 14870 | 0.353586231 | 0.445 | 0.236 | 4.81E-09 | 3.3E-13 | |
| Cortical Layer L2/3 | cTBS | Psd | pleckstrin and Sec7 domain containing | 73728 | 0.300529449 | 0.96 | 0.881 | 7.35E-10 | 5.0E-14 | |
| Cortical Layer L2/3 | cTBS | Eno1 | enolase 1, alpha non-neuron | 13806 | 0.253005294 | 0.915 | 0.892 | 0.005054135 | 3.4E-07 | |
| Cortical Layer L2/3 | cTBS | Tuba1a | tubulin, alpha 1A | 22142 | -0.267269854 | 0.986 | 0.994 | 1.25E-07 | 8.4E-12 | |
| Cortical Layer L2/3 | cTBS | Tatdn1 | TatD DNase domain containing 1 | 69694 | -0.269090494 | 0.111 | 0.306 | 6.28E-09 | 4.3E-13 | |
| Cortical Layer 4 | cTBS | Gm10076 | ribosomal protein L41 pseudogene | 100126819 | 0.745341144 | 0.705 | 0.406 | 5.11E-22 | 3.5E-26 | |
| Cortical Layer 4 | cTBS | Uba52 | ubiquitin A-52 residue ribosomal protein fusion product 1 | 22186 | 0.434269433 | 0.786 | 0.731 | 2.02E-05 | 1.4E-09 | |
| Cortical Layer 4 | cTBS | Rpl35 | ribosomal protein L35 | 66489 | 0.397464177 | 0.974 | 0.953 | 1.36E-11 | 9.2E-16 | |
| Cortical Layer 4 | cTBS | Selenop | selenoprotein P | 20363 | -0.294902507 | 0.412 | 0.558 | 0.022384033 | 1.5E-06 | |
| Cortical Layer 5 | cTBS | Gm10076 | ribosomal protein L41 pseudogene | 100126819 | 0.995523431 | 0.704 | 0.341 | 1.61E-73 | 1.1E-77 | |
| Cortical Layer 5 | cTBS | Uba52 | ubiquitin A-52 residue ribosomal protein fusion product 1 | 22186 | 0.533661504 | 0.787 | 0.685 | 7.44E-22 | 5.0E-26 | |
| Cortical Layer 5 | cTBS | Rpl35 | ribosomal protein L35 | 66489 | 0.516119763 | 0.981 | 0.918 | 1.51E-41 | 1.0E-45 | |
| Cortical Layer 5 | cTBS | Gstp1 | glutathione S-transferase, pi 1 | 14870 | 0.278587356 | 0.43 | 0.313 | 1.77E-05 | 1.2E-09 | |
| Cortical Layer 5 | cTBS | Polr2l | polymerase (RNA) II (DNA directed) polypeptide L | 66491 | 0.266333438 | 0.583 | 0.476 | 7.44E-05 | 5.0E-09 | |
| Cortical Layer 5 | cTBS | Cdk5r2 | cyclin-dependent kinase 5, regulatory subunit 2 (p39) | 12570 | -0.284455074 | 0.39 | 0.553 | 1.58E-10 | 1.1E-14 | |
| Cortical Layer 6 | cTBS | Gm10076 | ribosomal protein L41 pseudogene | 100126819 | 0.75879212 | 0.767 | 0.559 | 3.37E-30 | 2.3E-34 | |
| Cortical Layer 6 | cTBS | Gstp1 | glutathione S-transferase, pi 1 | 14870 | 0.474468605 | 0.512 | 0.285 | 5.93E-12 | 4.0E-16 | |
| Cortical Layer 6 | cTBS | Uba52 | ubiquitin A-52 residue ribosomal protein fusion product 1 | 22186 | 0.46715186 | 0.843 | 0.832 | 9.63E-11 | 6.5E-15 | |
| Cortical Layer 6 | cTBS | Rpl35 | ribosomal protein L35 | 66489 | 0.37508774 | 0.982 | 0.985 | 3.28E-14 | 2.2E-18 | |
| Piriform Cortex | cTBS | Gm10076 | ribosomal protein L41 pseudogene | 100126819 | 1.095206033 | 0.758 | 0.423 | 1.12E-41 | 7.6E-46 | |
| Piriform Cortex | cTBS | Uba52 | ubiquitin A-52 residue ribosomal protein fusion product 1 | 22186 | 0.61141369 | 0.779 | 0.701 | 1.70E-15 | 1.2E-19 | |
| Piriform Cortex | cTBS | Rpl35 | ribosomal protein L35 | 66489 | 0.547358357 | 0.972 | 0.958 | 1.05E-20 | 7.1E-25 | |
| Piriform Cortex | cTBS | Polr2l | polymerase (RNA) II (DNA directed) polypeptide L | 66491 | 0.36691555 | 0.607 | 0.429 | 5.11E-07 | 3.5E-11 | |
| Piriform Cortex | cTBS | Gstp1 | glutathione S-transferase, pi 1 | 14870 | 0.331980422 | 0.534 | 0.354 | 6.30E-06 | 4.3E-10 | |
| Piriform Cortex | cTBS | Rpl38 | ribosomal protein L38 | 67671 | 0.32644671 | 1 | 1 | 9.83E-21 | 6.6E-25 | |
| Piriform Cortex | cTBS | Cox7c | cytochrome c oxidase subunit 7C | 12867 | 0.323853069 | 1 | 0.998 | 1.34E-17 | 9.1E-22 | |
| Piriform Cortex | cTBS | Atp5md | ATP synthase membrane subunit DAPIT | 66477 | 0.317035604 | 1 | 1 | 1.43E-18 | 9.7E-23 | |
| Piriform Cortex | cTBS | Tmsb10 | thymosin, beta 10 | 19240 | 0.306765588 | 0.988 | 1 | 2.26E-06 | 1.5E-10 | |
| Piriform Cortex | cTBS | Rps21 | ribosomal protein S21 | 66481 | 0.288886951 | 1 | 1 | 3.95E-15 | 2.7E-19 | |
| Piriform Cortex | cTBS | Rps28 | ribosomal protein S28 | 54127 | 0.270532904 | 0.929 | 0.857 | 4.85E-05 | 3.3E-09 | |
| Piriform Cortex | cTBS | Ndufb4 | NADH:ubiquinone oxidoreductase subunit B4 | 68194 | 0.257858789 | 0.951 | 0.874 | 5.04E-05 | 3.4E-09 | |
| Piriform Cortex | cTBS | Agap2 | ArfGAP with GTPase domain, ankyrin repeat and PH domain 2 | 216439 | -0.25389471 | 0.228 | 0.385 | 0.000966429 | 6.5E-08 | |
| Piriform Cortex | cTBS | Cdk5r2 | cyclin-dependent kinase 5, regulatory subunit 2 (p39) | 12570 | -0.280578006 | 0.428 | 0.554 | 0.011840037 | 8.0E-07 | |
| Piriform Cortex | cTBS | Phyhip | phytanoyl-CoA hydroxylase interacting protein | 105653 | -0.294262305 | 0.527 | 0.629 | 0.006444959 | 4.4E-07 | |
| Piriform Cortex | cTBS | Dynll2 | dynein light chain LC8-type 2 | 68097 | -0.342342006 | 0.944 | 0.962 | 4.54E-08 | 3.1E-12 | |
| White Matter Tracts | cTBS | Gstp1 | glutathione S-transferase, pi 1 | 14870 | 0.685173258 | 0.707 | 0.597 | 3.50E-11 | 2.4E-15 | |
| White Matter Tracts | cTBS | Gm10076 | ribosomal protein L41 pseudogene | 100126819 | 0.565014967 | 0.938 | 0.887 | 5.47E-20 | 3.7E-24 | |
| White Matter Tracts | cTBS | Fth1 | ferritin heavy polypeptide 1 | 14319 | 0.517891522 | 1 | 1 | 5.79E-25 | 3.9E-29 | |
| White Matter Tracts | cTBS | Uba52 | ubiquitin A-52 residue ribosomal protein fusion product 1 | 22186 | 0.485056077 | 0.968 | 0.958 | 1.50E-12 | 1.0E-16 | |
| White Matter Tracts | cTBS | Rapgef4 | Rap guanine nucleotide exchange factor (GEF) 4 | 56508 | 0.367187139 | 0.843 | 0.795 | 1.53E-05 | 1.0E-09 | |
| White Matter Tracts | cTBS | Kif5a | kinesin family member 5A | 16572 | 0.31720512 | 0.992 | 0.99 | 1.21E-05 | 8.2E-10 | |
| White Matter Tracts | cTBS | Ftl1 | ferritin light polypeptide 1 | 14325 | 0.305588024 | 1 | 1 | 3.24E-07 | 2.2E-11 | |

**Data S9**

M1 iTBS L2/3

| DEGs Young sham vs Aged sham | iTBS DEGs unique to young adults | iTBS DEGs unique to young adults that are also ageing DEGs |
|---|---|---|
| Gapdh | Gstp1 | |
| Psd | Chd9 | |
| Mt1 | Ncald | |
| Ndufa4 | Tmsb10 | |
| Nme7 | Meg3 | |
| Actb | Cdk2ap1 | |
| Cox8a | Gm2000 | |
| Malat1 | | |
| Rpl4 | | |
| Rpl5 | | |
| Lars2 | | |
| Calm1 | | |
| Rps29 | | |
| Mbp | | |
| Rps28 | | |
| Cdk11b | | |
| Rbm3 | | |
| Dpysl2 | | |
| Ctxn1 | | |
| Ppia | | |
| Itm2c | | |
| Psap | | |
| Mt2 | | |
| Lrrc17 | | |
| Slc1a2 | | |
| Sptbn2 | | |
| Txndc15 | | |
| Ppp3r1 | | |
| Homer1 | | |
| Pfn1 | | |
| Eef1a2 | | |
| Arl8a | | |
| Ddn | | |
| Ewsr1 | | |
| Ppp2r1a | | |
| Stx1a | | |
| Cab39 | | |
| Stmn3 | | |
| Morf4l1 | | |
| Ptges3 | | |
| Pkm | | |
| Smim19 | | |
| Rps27 | | |
| Snrnp70 | | |
| Ogfrl1 | | |
| Rpl37a | | |
| Rps15a | | |
| Cspg5 | | |
| Ivns1abp | | |
| Abcf1 | | |
| Zfp365 | | |
| Arc | | |
| Lypla2 | | |
| Tuba1b | | |
| Set | | |
| Nrxn3 | | |
| Dynll2 | | |
| Cdc37 | | |
| Ilf3 | | |
| Capza2 | | |
| Pcp4 | | |
| AW047730 | | |
| Rpl3 | | |
| Ikbkb | | |
| Pja2 | | |
| Ccar2 | | |
| Ctsa | | |
| Gramd1a | | |
| Snhg6 | | |
| Bex2 | | |
| Pfdn5 | | |
| Gmpr | | |
| Kif5a | | |
| Insig1 | | |
| Atp1a1 | | |
| Dek | | |
| Anp32a | | |
| Rps24 | | |
| Nfkb2 | | |
| Smarcc2 | | |
| Ndn | | |
| Rpl9-ps6 | | |
| Gabbr1 | | |
| Alkbh7 | | |
| Cirbp | | |
| Ddx6 | | |
| Spint2 | | |
| Cox7b | | |
| Actr3 | | |
| Maco1 | | |
| Map1s | | |
| Eml2 | | |
| Rnf41 | | |
| St3gal5 | | |
| Sptbn1 | | |
| Tmem130 | | |
| Zwint | | |
| Maged1 | | |
| Rps20 | | |
| Bcl7a | | |
| Fam107a | | |
| Rps2 | | |
| Ptprj | | |
| Rac1 | | |
| Galnt16 | | |
| Rpn1 | | |
| Arfgap1 | | |
| Camk2b | | |
| Ctsb | | |
| Ppp2r5c | | |
| Smarcb1 | | |
| Tubb5 | | |
| Usp11 | | |
| Poglut1 | | |
| Phyhip | | |
| Pcbp2 | | |
| Bc1 | | |
| Puf60 | | |
| Spin1 | | |
| Ndrg2 | | |
| Cluh | | |
| Ssrp1 | | |
| Cort | | |
| Pantr1 | | |
| Usp50 | | |
| Ube3c | | |
| Gm10076 | | |
| Prdx6 | | |
| Mical2 | | |
| Eif6 | | |

M1 cTBS L2/3

| DEGs Young sham vs Aged sham | cTBS DEGs unique to young adults | cTBS DEGs unique to young adults that are also ageing DEGs |
|---|---|---|
| Uba52 | Lrrc17 | Bcl7a |
| Gapdh | Rps28 | Cox8a |
| Psd | Nrn1 | Ctsb |
| Mt1 | Tspyl4 | Ctxn1 |
| Ndufa4 | Snca | Ddn |
| Nme7 | Rps29 | Eef1a2 |
| Actb | Pacsin1 | Gapdh |
| Cox8a | Snap25 | Lrrc17 |
| Malat1 | Fam131a | Mical2 |
| Rpl4 | Ost4 | Mt1 |
| Rpl5 | Cox17 | Mt2 |
| Lars2 | Araf | Pfdn5 |
| Calm1 | Tomm7 | Phyhip |
| Rps29 | Ddn | Psd |
| Mbp | Snrpf | Rpl37a |
| Rps28 | Ndufa1 | Rpl4 |
| Cdk11b | Cox7c | Rpl5 |
| Rbm3 | Isca1 | Rpl9-ps6 |
| Dpysl2 | Dmtn | Rps15a |
| Ctxn1 | Rps27 | Rps20 |
| Ppia | Snhg8 | Rps24 |
| Itm2c | Txn1 | Rps27 |
| Psap | Actr2 | Rps28 |
| Mt2 | Snhg6 | Rps29 |
| Lrrc17 | Gng13 | Snhg6 |
| Slc1a2 | Kctd13 | Sptbn2 |
| Rpl35 | Slc6a1 | Stx1a |
| Sptbn2 | Nptx2 | Uba52 |
| Txndc15 | Atp6v1a | Usp50 |
| Ppp3r1 | Sdhd | Zfp365 |
| Homer1 | Atp5md | Zwint |
| Pfn1 | Elmo2 | |
| Eef1a2 | Snhg9 | |
| Arl8a | Zwint | |
| Ddn | Agap2 | |
| Ewsr1 | Usp50 | |
| Ppp2r1a | Lmo4 | |
| Stx1a | Dnajb5 | |
| Cab39 | Sema7a | |
| Stmn3 | Arpp21 | |
| Morf4l1 | Hpcal4 | |
| Ptges3 | Naa20 | |
| Pkm | Sem1 | |
| Smim19 | Nr4a1 | |
| Rps27 | Schip1 | |
| Snrnp70 | Wsb2 | |
| Ogfrl1 | Megf9 | |
| Rpl37a | Atp1a3 | |
| Rps15a | Smim26 | |
| Cspg5 | Sipa1l1 | |
| Ivns1abp | Zfp365 | |
| Abcf1 | Slc25a22 | |
| Zfp365 | Camk2a | |
| Arc | Stxbp1 | |
| Lypla2 | App | |
| Tuba1b | Ttc7b | |
| Set | Shank1 | |
| Nrxn3 | Plppr4 | |
| Dynll2 | Atn1 | |
| Cdc37 | Rpl37a | |
| Ilf3 | Ppp6r1 | |
| Capza2 | Cox8a | |
| Pcp4 | Vxn | |
| AW047730 | Fam131b | |

| | |
|---|---|
| Rpl3 | Stum |
| Ikbkb | Pam |
| Pja2 | Slc4a10 |
| Ccar2 | Fbxl16 |
| Ctsa | Rab3c |
| Gramd1a | Gas7 |
| Snhg6 | Lmtk2 |
| Bex2 | Phyhip |
| Pfdn5 | Cops7a |
| Gmpr | Ppm1h |
| Kif5a | Gucy1b1 |
| Insig1 | Kcnh3 |
| Atp1a1 | Nell2 |
| Dek | Rnf5 |
| Anp32a | Dusp7 |
| Rps24 | Vgf |
| Nfkb2 | Lpcat4 |
| Smarcc2 | Huwe1 |
| Ndn | Jph4 |
| Rpl9-ps6 | Mgat3 |
| Gabbr1 | Sptbn2 |
| Alkbh7 | Sprn |
| Cirbp | Rusc2 |
| Ddx6 | F3 |
| Spint2 | Cacna2d1 |
| Cox7b | Rpl9-ps6 |
| Actr3 | Spag9 |
| Maco1 | Cinp |
| Map1s | Dnm1 |
| Eml2 | Psd |
| Rnf41 | Ncald |
| St3gal5 | Syngap1 |
| Sptbn1 | Senp3 |
| Tmem130 | Mat2b |
| Zwint | Inpp4a |
| Maged1 | Dnajc21 |
| Rps20 | Tsnax |
| Bcl7a | Comt |
| Fam107a | Rgs7bp |
| Rps2 | Cmpk1 |
| Ptprj | Cs |
| Rac1 | Atp6v1h |
| Galnt16 | Gnai1 |
| Rpn1 | Ccnb1 |
| Arfgap1 | Phpt1 |
| Camk2b | Gm1673 |
| Ctsb | Dkk3 |
| Ppp2r5c | Chst2 |
| Smarcb1 | Pip5k1c |
| Tubb5 | Chd9 |
| Usp11 | Kif3c |
| Poglut1 | Dnajb4 |
| Phyhip | Bloc1s1 |
| Pcbp2 | Csnk2a2 |
| Bc1 | Junb |
| Puf60 | 1810037I17Rik |
| Spin1 | Dab2ip |
| Ndrg2 | Eif4a2 |
| Cluh | Dnajc6 |
| Ssrp1 | Slc8a2 |
| Cort | Psma5 |
| Pantr1 | Hectd4 |
| Usp50 | Mical2 |
| Ube3c | Cadm4 |
| Gm10076 | Klc2 |
| Prdx6 | Eef1a2 |
| Mical2 | Fgfr1op2 |

| | |
|---|---|
| Eif6 | Car2 |
| | Phyhipl |
| | Unc5a |
| | Ank2 |
| | Rap2a |
| | Usp19 |
| | Arpc4 |
| | Rheb |
| | Abhd8 |
| | Tsc22d4 |
| | Dact3 |
| | Hivep2 |
| | Numb |
| | Mt1 |
| | Itpka |
| | Ndufa3 |
| | Ctxn1 |
| | Atxn2l |
| | Zmiz1 |
| | Bhlhe40 |
| | Brms1l |
| | Rtn4r |
| | Glul |
| | Nrxn2 |
| | Cux2 |
| | Ndrg4 |
| | Cck |
| | Cfap20 |
| | Erh |
| | Gpr26 |
| | Sv2b |
| | Acyp2 |
| | Nfix |
| | Tmem196 |
| | Spred1 |
| | Camk1g |
| | Frrs1l |
| | Bccip |
| | Sft2d1 |
| | Ppp1r9b |
| | Stx1a |
| | B230334C09Rik |
| | Ncan |
| | Bzw1 |
| | Cacng3 |
| | Atxn2 |
| | Rai1 |
| | Rps27rt |
| | Lzts3 |
| | Pofut2 |
| | Higd2a |
| | Ubqln2 |
| | Prkaca |
| | Pou3f3 |
| | Insyn1 |
| | Cnksr2 |
| | Wwc1 |
| | Enc1 |
| | Med31 |
| | Dusp6 |
| | Nutf2-ps1 |
| | Ppme1 |
| | Apba1 |
| | Spcs3 |
| | Tacc1 |
| | Spry2 |
| | Rimbp2 |

Clcn2
Lnpep
Gja1
U2af2
Cd47
Tmeff1
Ano3
Add2
Bcl2l2
Jdp2
Arhgap32
Phactr1
Hprt
Slc30a3
Rapgef2
Btbd10
Psip1
Tiprl
B3gat1
Unc50
Cacng7
Herc3
Rgcc
Ndfip2
Cd302
Myadm
Yme1l1
Arhgap39
Klf13
Rnd1
Arhgef12
Prrc2a
Ank
Ppp2cb
Sptan1
Lsm8
Rab28
Kcna2
Mif
Ogdhl
Spop
Rraga
Prex1
Golph3
Mt2
Frmpd4
Ptms
Trappc2
Pde4dip
Yaf2
Ndufb3
Stard8
Arhgap1
Coa6
Rgs6
Mt3
Cntnap1
Skil
Nbl1
Nrgn
Mrps28
Sncb
Ryr2
Mapre2
Hopx
Med25
Romo1

Bcl7a
Prpf8
Fus
Pmpca
Irak1bp1
Cyth1
Nptxr
Trrap
Srsf5
Thra
Mgat5b
Scn2a
Gnao1
Rpl39
Lrrc57
Ccn3
Chrm3
Usp7
Etv5
Syt17
Dgkz
Fam81a
Ccnh
Ahcyl2
Car10
Usp9x
Acap2
Ppp2r2c
Rapgefl1
Myrip
Grk6
Trp53i11
Samd8
Pin1
Tmed7
L1cam
Rasgef1c
Pcdh10
Lingo1
Luzp2
Dpf1
Nova2
Lamtor3
Crip2
Arpp19
Ssbp4
Cds2
Exoc1
Abhd2
Usp12
Kitl
Atxn1
Msi2
Irs2
Rragd
Acat1
Mthfsl
Rcn2
Mpv17
Rnasek
Rpl10-ps3
Rbbp6
Cnnm1
Lrfn5
Syn1
Insig2
Prkar1a

Cpeb2
Klhl9
Mfsd14a
Polr3k
Hmgb1
Fam126b
Zfr
Kmt2d
Ap1ar
Yipf4
Kdm7a
Cnep1r1
Eif4g1
Slc17a7
Gfra4
Scn1a
Wdr89
Kcnh1
Arnt2
Rnf114
Sgsm2
Homer2
Kcnv1
Npas2
Bpnt1
Ccdc32
Nae1
Olfm2
5730455P16Rik
Gtpbp2
Pde4b
Rabggtb
Stx4a
Ramp2
Tmem178b
Gabra3
Katnal1
Atp6v1b2
Cggbp1
Clip1
Map6d1
Usp25
Fjx1
Cdc42se2
Fmn2
Mtln
Cstb
Mtmr9
Atp5mpl
Rtn4rl1
Pde6d
Gas6
Spred2
Smim4
Kctd1
Pld3
Stim2
Cadm1
Mpp3
Socs7
Lsm3
Llgl1
Orc6
Thrb
Copb1
Prkci
Rock2

Pgls
Klhl29
Ifngr2
Abhd3
Cul2
Ulk1
Gm20300
Ptprf
Idi1
Pmpcb
Stx1b
Ctnnd1
Gramd4
Dvl1
Lst1
Meis2
Ttyh3
Fam13c
Rgs20
Shank3
Pank4
Ehd1
Ak4
Dagla
Shd
Hes5
Acvr1b
Slc4a4
Pcsk1
Rnf141
Erg28
Trim8
Tmem184c
Ube2q1
Ift57
Ccdc6
Mrpl27
Srrm1
Sdf2
Lin52
Arhgef10l
Smim15
Rabl6
Cebpb
Cxcl12
C130071C03Rik
Sfxn1
Atp6v1g2
Khsrp
Dhx9
Rgs14
Rnf146
Myef2
Nme3
Zdhhc14
Tmed2
Iars2
Nmt2
Dcakd
Satb2
Mtx2
Raph1
Sh3bp5l
Rnf123
Tnnc1
Fam241b
Tcea1

Setd5
Coq5
Zfp445
Nol4
Neurl1b
Rpl7l1
Hspa14
Alas1
Spats2l
Hnrnpf
Clasp2
Lix1
Rap1a
Pitpnm3
Tbc1d10a
Polr2h
Cebpg
Cd200
Gspt1
Tbl1xr1
Shprh
Jmjd1c
Hba-a2
Gapdh
Rps5
Ndufa13
Eef1a1
Rpl31
Rps17
Rpsa
Rpl13
Gabarap
Rpl35a
Rpl30
Hsp90aa1
Pfdn5
Ctsb
Rpl9
Rps11
Fkbp2
Rps4x
Chgb
Rps12
Rpl26
Ppp3ca
Gabarapl2
Gpr88
Rpl6
Naca
Rpl11
Rps15a
Fau
Rps3a1
Rpl7
Mrfap1
Nenf
Rpl13a
Rpl14
Pink1
Rpl24
Mobp
Rps7
Rpl4
Rpl21
Nefm
Rpl10
Rps20

Rps8
Cryab
Rpl5
Rps27a
Rps24
Hbb-bs

M1 iTBS L5

| DEGs Young sham vs Aged sham | iTBS DEGs unique to young adults | iTBS DEGs unique to young adults that are also ageing DEGs |
|---|---|---|
| Uba52 | Uba52 | Bc1 |
| Rpl4 | Gstp1 | Chd9 |
| Gapdh | Mobp | Gm10076 |
| Rps29 | Chd9 | Mbp |
| Mt1 | Mt2 | Mobp |
| Cox8a | Dbi | Mt1 |
| Malat1 | Eno1 | Mt2 |
| Snhg6 | Slc1a3 | Rps27 |
| Rps28 | Mbp | Uba52 |
| Lrrc17 | Mt1 | |
| Psd | Fbxl16 | |
| Mt2 | Dbndd2 | |
| Slc1a2 | Rps27 | |
| Rps24 | Pitpnm1 | |
| Rpl5 | Sncb | |
| Cox7b | Gm2000 | |
| Nme7 | Bc1 | |
| Usp50 | | |
| Dpysl2 | | |
| Rph3a | | |
| Fam107a | | |
| Ndufa4 | | |
| Actb | | |
| Rpl9-ps6 | | |
| Lmtk2 | | |
| Sema7a | | |
| Fau | | |
| Zwint | | |
| Ctxn1 | | |
| Mfge8 | | |
| Arc | | |
| Itm2c | | |
| Mbp | | |
| Homer1 | | |
| Rpl24 | | |
| Rpl21 | | |
| Lars2 | | |
| Cspg5 | | |
| Eef1a1 | | |
| Pcp4 | | |
| Rps27a | | |
| Sst | | |
| Zfp365 | | |
| Mobp | | |
| Mical2 | | |
| Ndufa3 | | |
| Stmn2 | | |
| Kif5a | | |
| Eef1a2 | | |
| Rpl37a | | |
| Ddn | | |
| Ivns1abp | | |
| Rps2 | | |
| Rps27 | | |
| Gm10076 | | |
| Gnb2 | | |

Plcxd2
Bc1
Egr3
Dclk1
Kifc2
Gpr26
Rpl35
Pip5k1c
Pfn1
Ndufv1
Eno1b
Myadm
Mif
Tomm7
Atp6v1g2
Ypel3
Rpl8
Cplx1
Tspyl4
Ndrg2
Scn1a
Pak1
Rbm3
Gtf3c3
Rims1
Cyb5r4
Rps7
Ina
Rnasek
Sptbn1
Rpl9
Rps8
Cinp
Adcy1
Dusp7
Hspa5
Ost4
Hecw1
Atxn2l
Sptbn2
Fxyd7
Rundc3a
Camk1g
Utrn
Ndufa7
Kif5c
Nptx2
Rpsa
Ptma
Rapgef4
Rpl3
Tmem160
Pfdn5
Capzb
Atp1a1
Oaz1
Nrxn3
Napb
Kcnj10

Stac2
Sptan1
Galnt9
Asap2
Rab31
Kif21a
Ewsr1
Stx1a
Ids
Stxbp1
Rpl14
Pfdn2
Gde1
Hpcal4
Dctn5
Camk2a
Rps4x
Vgf
Eif4a1
Dynll2
Arl8a
Hs3st2
Dok6
Chd9
Hivep2
Saxo2
Acsl5
Aff3
Kif21b
Hsph1
Agap2
Ntsr2
Rps3a1
Gfod1
Csf1r
BC005537
Stradb
Asap1
Ubap2l
Ly6e
Tfrc
Tlcd1
Slc32a1
Atxn2
Tpp1
Hnrnpa2b1
Washc5
Jund
Sgsm1
Fkbp1b
Nucks1
Nme2
Rpl6
Car10
Plk2
Aldh1a1
Man1c1
Ipo7
Kcnv1

Bex1

M1 cTBS L5

| DEGs Young sham vs Aged sham | cTBS DEGs unique to young adults | cTBS DEGs unique to young adults that are also ageing DEGs |
|---|---|---|
| Uba52 | Lrrc17 | Agap2 |
| Rpl4 | Rps28 | Atxn2 |
| Gapdh | Tspyl4 | Atxn2l |
| Rps29 | Snap25 | Bex1 |
| Mt1 | Pacsin1 | Camk2a |
| Cox8a | Rps29 | Chd9 |
| Malat1 | Actr2 | Cinp |
| Snhg6 | Snca | Cox8a |
| Rps28 | Nrn1 | Cplx1 |
| Lrrc17 | Glul | Ddn |
| Psd | Tomm7 | Dusp7 |
| Mt2 | Camk2a | Eef1a1 |
| Slc1a2 | Atp6v1a | Eno1b |
| Rps24 | Slc25a22 | Ewsr1 |
| Rpl5 | Dmtn | Fau |
| Cox7b | Ndufa1 | Gapdh |
| Nme7 | App | Gpr26 |
| Usp50 | Agap2 | Hnrnpa2b1 |
| Dpysl2 | Zwint | Hpcal4 |
| Rph3a | Tatdn1 | Ids |
| Fam107a | Ddn | Jund |
| Ndufa4 | Cox17 | Kif5a |
| Actb | Ost4 | Kif5c |
| Rpl9-ps6 | Araf | Kifc2 |
| Lmtk2 | Huwe1 | Lmtk2 |
| Sema7a | Snhg9 | Lrrc17 |
| Fau | Rap1gds1 | Mical2 |
| Zwint | Snrpf | Mt1 |
| Ctxn1 | Cplx2 | Mt2 |
| Mfge8 | Ndrg4 | Myadm |
| Arc | Snhg6 | Ndufa3 |
| Itm2c | Phpt1 | Ndufa7 |
| Mbp | Sdhd | Nptx2 |
| Homer1 | Cops7a | Oaz1 |
| Rpl24 | Stxbp1 | Ost4 |
| Rpl21 | Ank | Pak1 |
| Lars2 | Slc6a1 | Pfdn5 |
| Cspg5 | Txn1 | Pip5k1c |
| Eef1a1 | Chd9 | Psd |
| Pcp4 | Usp50 | Rapgef4 |
| Rps27a | Mt2 | Rnasek |
| Sst | Isca1 | Rph3a |
| Zfp365 | Cntnap1 | Rpl14 |
| Mobp | Eif4a2 | Rpl21 |
| Mical2 | Hpcal4 | Rpl24 |
| Ndufa3 | Frrs1l | Rpl3 |
| Stmn2 | Rapgef4 | Rpl4 |
| Kif5a | Rab3c | Rpl5 |
| Eef1a2 | Arpp21 | Rpl6 |
| Rpl37a | Snhg8 | Rpl8 |
| Ddn | Kctd13 | Rpl9 |
| Ivns1abp | Ppp1r9b | Rpl9-ps6 |
| Rps2 | Fbxl16 | Rps2 |
| Rps27 | Zfp365 | Rps24 |
| Gm10076 | Phactr1 | Rps27a |
| Gnb2 | Rgs7bp | Rps28 |

Plcxd2
Bc1
Egr3
Dclk1
Kifc2
Gpr26
Rpl35
Pip5k1c
Pfn1
Ndufv1
Eno1b
Myadm
Mif
Tomm7
Atp6v1g2
Ypel3
Rpl8
Cplx1
Tspyl4
Ndrg2
Scn1a
Pak1
Rbm3
Gtf3c3
Rims1
Cyb5r4
Rps7
Ina
Rnasek
Sptbn1
Rpl9
Rps8
Cinp
Adcy1
Dusp7
Hspa5
Ost4
Hecw1
Atxn2l
Sptbn2
Fxyd7
Rundc3a
Camk1g
Utrn
Ndufa7
Kif5c
Nptx2
Rpsa
Ptma
Rapgef4
Rpl3
Tmem160
Pfdn5
Capzb
Atp1a1
Oaz1
Nrxn3
Napb
Kcnj10
Schip1
Ank2
Smim26
Rpl9-ps6
Rnf5
Lmo4
Dnajc6
Fus
Naa20
Sst
Atn1
Wsb2
Pip5k1c
Rph3a
Sem1
Cs
Sema7a
Nptx2
Gja1
Prrc2b
Slc1a3
Unc5a
Kif3c
Fam131a
Atp5g3
Sptbn1
Ppp1r16b
Spag9
Psip1
Prrc2a
Ids
Atp5md
Lpcat4
Mical2
Slc4a10
Klc2
R3hdm4
Ewsr1
Sptbn2
Fam131b
Rusc2
Eps15
Ndufa3
1810037I17Rik
Brms1l
Spop
Gucy1b1
Phyhipl
Shank1
Nrxn1
Psd3
Bloc1s1
Sipa1l1
Atp6v1h
Luzp2
Kif5c
Map2
Phyhip
Syn1
Rps29
Rps3a1
Rps4x
Rps7
Rps8
Rpsa
Saxo2
Scn1a
Sema7a
Slc1a2
Snhg6
Sptan1
Sptbn1
Sptbn2
Sst
Stxbp1
Tomm7
Tspyl4
Ubap2l
Usp50
Vgf
Zfp365
Zwint

| | |
|---|---|
| Stac2 | Ahcyl2 |
| Sptan1 | Mgat3 |
| Galnt9 | Herc3 |
| Asap2 | Rcn2 |
| Rab31 | Ndfip2 |
| Kif21a | Chst2 |
| Ewsr1 | Paip2 |
| Stx1a | Car2 |
| Ids | Sptan1 |
| Stxbp1 | Kif5a |
| Rpl14 | Gnao1 |
| Pfdn2 | Srsf5 |
| Gde1 | Atxn2l |
| Hpcal4 | Inpp4a |
| Dctn5 | Msi2 |
| Camk2a | Atxn2 |
| Rps4x | Lmtk2 |
| Vgf | Dusp7 |
| Eif4a1 | Ubqln2 |
| Dynll2 | Ttc7b |
| Arl8a | Arhgef12 |
| Hs3st2 | Comt |
| Dok6 | Psma5 |
| Chd9 | Ccnb1 |
| Hivep2 | Lynx1 |
| Saxo2 | Gng13 |
| Acsl5 | Scn1a |
| Aff3 | Fgfr1op2 |
| Kif21b | Syt1 |
| Hsph1 | Mapk1 |
| Agap2 | Apba1 |
| Ntsr2 | Rabgap1l |
| Rps3a1 | Arrb1 |
| Gfod1 | Spock2 |
| Csf1r | Add2 |
| BC005537 | Rheb |
| Stradb | Gnai1 |
| Asap1 | Tacc1 |
| Ubap2l | Tsc22d3 |
| Ly6e | Sparcl1 |
| Tfrc | Eno1b |
| Tlcd1 | Ptprz1 |
| Slc32a1 | Panx2 |
| Atxn2 | Syngr1 |
| Tpp1 | Mapre2 |
| Hnrnpa2b1 | Ptn |
| Washc5 | Hectd4 |
| Jund | Bpnt1 |
| Sgsm1 | Romo1 |
| Fkbp1b | Bhlhe40 |
| Nucks1 | Rabggtb |
| Nme2 | Kcna2 |
| Rpl6 | Gpr26 |
| Car10 | Slc4a4 |
| Plk2 | Slc8a2 |
| Aldh1a1 | Kifc2 |
| Man1c1 | Cstb |
| Ipo7 | Scg2 |
| Kcnv1 | Pdpk1 |

| | |
|---|---|
| Bex1 | Apc |
| | Stx1b |
| | Erc2 |
| | Lamp1 |
| | Psd |
| | Tcf4 |
| | Klf13 |
| | Cd47 |
| | Nell2 |
| | Ctnnb1 |
| | Ssb |
| | Ppm1h |
| | Cmpk1 |
| | Tmeff1 |
| | Syngap1 |
| | Dgkz |
| | Ncald |
| | Insyn1 |
| | Dusp6 |
| | Gria2 |
| | Sprn |
| | Klf9 |
| | Megf9 |
| | Idi1 |
| | Cox8a |
| | Senp3 |
| | Acyp2 |
| | Churc1 |
| | Myadm |
| | Wdr89 |
| | S1pr1 |
| | Dnm1 |
| | Arhgap39 |
| | Selenof |
| | Ppme1 |
| | Cds2 |
| | Numb |
| | Stum |
| | Ttyh1 |
| | Iars2 |
| | Kcnh1 |
| | Prkar1a |
| | Pde4dip |
| | Tmed7 |
| | Tef |
| | Dnajb4 |
| | Ap1ar |
| | Sft2d1 |
| | Reep1 |
| | Lrp11 |
| | Nedd4 |
| | Dnajb5 |
| | Brsk1 |
| | Elavl3 |
| | Clasp2 |
| | Actn4 |
| | Ogdhl |
| | Fam126b |
| | Slc12a5 |

Rnf157
Vxn
Cinp
Arpc4
Cadm4
Acat1
Ssr3
Dab2ip
Dbndd2
Camk2n1
Pik3r1
Mfsd14a
Hopx
Plppr4
Scn4b
Oxr1
Tiprl
Hnrnpa2b1
Gpm6b
Rnf114
Pcdh1
F3
Pitpnm3
Socs7
Ccser2
Kcnh3
B230312C02Rik
Cpeb2
Bcl2l2
Snx5
Arhgef4
Usp31
Sdf2
Rab28
Atf2
Ndufa7
Dhx9
Rpl10-ps3
Dnal1
Vgf
Myrip
Cfap20
Cplx1
Hibadh
Aftph
Add3
Arhgef10l
Trrap
Rps27rt
Rpl39
Ppp2cb
Slc1a2
Rnasek
Fmn2
Kif1a
Slc36a1
Tmem128
Pmpcb
Gas6

Unc50
Cyth1
Fam120a
Epb41l1
Ldb1
Cd302
Gtpbp2
Ube2d1
Mrps28
Samd8
Mrpl10
Ppp1r12b
Alas1
Pdcd6ip
Ccng2
Ubap2l
Rnf19b
Mt1
Abhd2
Hace1
Dock4
Vstm2b
Fbrsl1
Ppp2r2c
Ttbk2
Nae1
Npc1
Ifngr2
Adgrl2
Spry2
Pank4
Stox2
Spns2
Saxo2
Aagab
A830018L16Rik
Pak1
Arpp19
Nadk
Atp5d
Mpc2
Rpl3
Actg1
Eef1a1
Ftl1
Rpl23a
Atp5o
Chchd2
Rps13
Rpl31
Rps5
Oaz1
Rpl23
Ubb
Ndufa2
Ndufb11
Rpl19
Rpl35a
Pebp1

Jund
Tpt1
Rps16
Dynll1
Rps3
Hba-a1
Rpl27a
Atp9a
Rps14
Gabarapl2
Rps15a
Ndufb10
Rpl32
Rps15
Rps3a1
Selenom
Rpsa
Fkbp2
Smdt1
Rps4x
Bex1
Hras
Rpl17
Rps10
Rps11
Ndufb9
Gapdh
Eno1
Rpl7
Rpl6
Ndufb8
Ndufa13
Rpl26
Rpl18
Rpl8
Rpl30
Rpl5
Rpl11
Rps12
Rpl18a
Rpl14
Nenf
Rpl24
Pfdn5
Rpl13a
Rpl13
Rps20
Rpl9
Rpl4
Myl4
Fau
Rps2
Rps7
Rpl21
Rps8
Rps27a
Rps24
Rpl10
Hbb-bs

M1 iTBS L6

| DEGs Young sham vs Aged sham | iTBS DEGs unique to young adults | iTBS DEGs unique to young adults that are also ageing DEGs |
|---|---|---|
| Mobp | Penk | Gapdh |
| Uba52 | Dbi | Rps27 |
| Mbp | Ptn | |
| Pcp4 | Gpr88 | |
| Gapdh | Rps27 | |
| Rps29 | Stmn3 | |
| Lrrc17 | Gapdh | |
| Fau | Akt1s1 | |
| Rpl9-ps6 | Fbxl19 | |
| Rps24 | Gprin1 | |
| Ost4 | Hapln4 | |
| Kif5a | Ttc9b | |
| Ighm | | |
| Rpl4 | | |
| Epas1 | | |
| Ccn2 | | |
| Cox8a | | |
| Dpysl2 | | |
| Slc1a2 | | |
| Rps28 | | |
| Eef1a1 | | |
| Rpl5 | | |
| Ro60 | | |
| Tmem167 | | |
| Pacsin1 | | |
| Hivep2 | | |
| Ctxn1 | | |
| Rph3a | | |
| Rps27a | | |
| Rpl21 | | |
| Rps2 | | |
| Far1 | | |
| Mtmr9 | | |
| Gadd45a | | |
| Zdhhc14 | | |
| Tatdn1 | | |
| Trp53inp2 | | |
| Asap1 | | |
| Mical2 | | |
| Flcn | | |
| Unc80 | | |
| Kctd1 | | |
| Usp50 | | |
| Pitpnc1 | | |
| Cldn11 | | |
| Setd7 | | |
| Snhg12 | | |
| Arhgap39 | | |
| Snx6 | | |
| Arc | | |
| Stac2 | | |
| Ndfip2 | | |
| Kcnk1 | | |
| Man1a2 | | |
| Micu1 | | |
| Tsc22d3 | | |

Camkk1
Myo6
Ccp110
Nrbp2
Tnk2
Nme7
Gnai1
Tspan2
Large1
Hacd1
Psd
Gapvd1
Rapgef1
Rps27
Synrg
Cisd2
Efna3
Mal
Dvl3
Vps37a
Rps7
Sf3b4
Ptp4a3
Prss23
Tnks2
Tbc1d20
Sesn1
Adcy1
Evi2a
Itgb1
Ppp2r5b
Agap2

M1 cTBS L6

| DEGs Young sham vs Aged sham | cTBS DEGs unique to young adults | cTBS DEGs unique to young adults that are also ageing DEGs |
|---|---|---|
| Mobp | Lrrc17 | Agap2 |
| Uba52 | Basp1 | Ccn2 |
| Mbp | Rps28 | Eef1a1 |
| Pcp4 | Pacsin1 | Fau |
| Gapdh | Tspyl4 | Gapdh |
| Rps29 | Ccn2 | Kif5a |
| Lrrc17 | Rps29 | Lrrc17 |
| Fau | Ost4 | Mtmr9 |
| Rpl9-ps6 | Snca | Ndfip2 |
| Rps24 | Araf | Ost4 |
| Ost4 | Snap25 | Pacsin1 |
| Kif5a | Ndufa1 | Rpl21 |
| Ighm | Glul | Rpl4 |
| Rpl4 | Agap2 | Rpl5 |
| Epas1 | Atp1a3 | Rpl9-ps6 |
| Ccn2 | Dmtn | Rps2 |
| Cox8a | Tatdn1 | Rps24 |
| Dpysl2 | Rapgef4 | Rps27 |
| Slc1a2 | Tomm7 | Rps27a |
| Rps28 | Snhg9 | Rps28 |
| Eef1a1 | Cox17 | Rps29 |
| Rpl5 | Phactr1 | Rps7 |
| Ro60 | Gas7 | Tatdn1 |
| Tmem167 | App | Usp50 |
| Pacsin1 | Isca1 | |
| Hivep2 | Slc6a1 | |
| Ctxn1 | Car2 | |
| Rph3a | Rps27 | |
| Rps27a | Snrpf | |
| Rpl21 | Zwint | |
| Rps2 | Ndfip2 | |
| Far1 | Ank2 | |
| Mtmr9 | Rpl9-ps6 | |
| Gadd45a | Smim26 | |
| Zdhhc14 | Kctd13 | |
| Tatdn1 | Srsf5 | |
| Trp53inp2 | Slc25a22 | |
| Asap1 | Ewsr1 | |
| Mical2 | Ddn | |
| Flcn | Hpcal4 | |
| Unc80 | Ndrg4 | |
| Kctd1 | Sem1 | |
| Usp50 | Phpt1 | |
| Pitpnc1 | Fam131b | |
| Cldn11 | Cox7c | |
| Setd7 | Huwe1 | |
| Snhg12 | Ank | |
| Arhgap39 | Arpp21 | |
| Snx6 | Dnajc6 | |
| Arc | Sdhd | |
| Stac2 | Fam131a | |
| Ndfip2 | Klf13 | |
| Kcnk1 | Rap1gds1 | |
| Man1a2 | Fam168a | |
| Micu1 | Snhg8 | |
| Tsc22d3 | Eif4a2 | |

| | |
|---|---|
| Camkk1 | Naa20 |
| Myo6 | Jph4 |
| Ccp110 | Stxbp1 |
| Nrbp2 | Mt2 |
| Tnk2 | Atn1 |
| Nme7 | Cadm4 |
| Gnai1 | Mtmr9 |
| Tspan2 | Ppp1r9b |
| Large1 | Rnf5 |
| Hacd1 | Camk2a |
| Psd | Tef |
| Gapvd1 | Prkar1a |
| Rapgef1 | Kif5a |
| Rps27 | Rpl39 |
| Synrg | Hnrnpa2b1 |
| Cisd2 | Rusc2 |
| Efna3 | Add3 |
| Mal | Rgs7bp |
| Dvl3 | Vxn |
| Vps37a | Lpcat4 |
| Rps7 | Gja1 |
| Sf3b4 | Sptbn1 |
| Ptp4a3 | Rab3c |
| Prss23 | Fam219a |
| Tnks2 | Atp5g3 |
| Tbc1d20 | Syngr1 |
| Sesn1 | Sptan1 |
| Adcy1 | Gpr88 |
| Evi2a | Nsf |
| Itgb1 | Usp50 |
| Ppp2r5b | Comt |
| Agap2 | Rps27rt |
| | Rab28 |
| | Atxn2 |
| | Spry2 |
| | Arhgap32 |
| | Senp3 |
| | Rpl37a |
| | Sdk2 |
| | Cyth1 |
| | Kcnh1 |
| | Cpox |
| | Idi1 |
| | Mrps28 |
| | Cpe |
| | Ubb |
| | Rpl23 |
| | Dynll1 |
| | Cox4i1 |
| | Rpl15 |
| | Uqcrq |
| | Rpl27a |
| | Rps16 |
| | Rps15 |
| | Rpl29 |
| | Eef1a1 |
| | Rps15a |
| | Cst3 |
| | Rps12 |

Uqcrh
Pfdn5
Rpl35a
Rpsa
Rpl30
Rps11
Rps3a1
Tpt1
Fis1
Rpl8
Rps5
Rps13
Rpl18
Rpl18a
Rpl4
Rpl17
Ttc9b
Hba-a1
Rps2
Rpl7
Rps3
Ndufa13
Rpl26
Rpl5
Rpl9
Gapdh
Ndufb8
Nme2
Rps4x
Rpl24
Rpl13a
Cryab
Fau
Rpl14
Myl4
Rpl11
Ndufb9
Rpl21
Rps20
Rpl13
Rps8
Hbb-bs
Rpl6
Rps7
Rps27a
Rps24
Rpl10

SS iTBS L2/3

| DEGs Young sham vs Aged sham | iTBS DEGs unique to young adults | iTBS DEGs unique to young adults that are also ageing DEGs |
|---|---|---|
| Gapdh | Gm10076 | |
| Uba52 | Rpl35 | |
| Rpl4 | Pcp4 | |
| Malat1 | Gstp1 | |
| Nme7 | Tmsb10 | |
| Psd | Rps27 | |
| Calm1 | Camk2n1 | |
| Cox8a | Aldoc | |
| Ndufa4 | Ddit4l | |
| Actb | Gm2000 | |
| Dpysl2 | Ppme1 | |
| Ost4 | Kcnj10 | |
| Lrrc17 | Glul | |
| Slc1a2 | | |
| Snhg6 | | |
| Bc1 | | |
| Lars2 | | |
| Pgam1 | | |
| Rps2 | | |
| Penk | | |
| Eef1a1 | | |
| Ddn | | |
| Rpl9-ps6 | | |
| Rpl14 | | |
| Mt1 | | |
| Pfn1 | | |
| Rps28 | | |
| Rps24 | | |
| Hsp90ab1 | | |
| Comt | | |
| Ndufa1 | | |
| Txn1 | | |
| Ppp2r1a | | |
| Arl8a | | |
| Rps27a | | |
| Eef1a2 | | |
| Rpl5 | | |
| Rpl6 | | |
| Fkbp8 | | |
| Usp50 | | |
| Tspyl4 | | |
| Gnb2 | | |
| Rps8 | | |
| Rps29 | | |
| Ddx5 | | |
| Tubb3 | | |
| Ewsr1 | | |
| Mical2 | | |
| Dlg4 | | |

Cab39
Gpr26
Stub1
Rpl3
BC005537
Arc
Rps15a
Maged1
Ppp3r1
Nap1l5
Cinp
Rgs20
Lmtk2
A830009L08Rik
Ptma
Atox1
Cox7b
Ctxn1
Junb
Pfdn5
Ptk2b

SS cTBS L2/3

| DEGs Young sham vs Aged sham | cTBS DEGs unique to young adults | cTBS DEGs unique to young adults that are also ageing DEGs |
| --- | --- | --- |
| Gapdh | Lrrc17 | Cinp |
| Uba52 | Basp1 | Comt |
| Rpl4 | Rps28 | Cox8a |
| Malat1 | Tspyl4 | Ddn |
| Nme7 | Snap25 | Eef1a1 |
| Psd | Snca | Ewsr1 |
| Calm1 | Dmtn | Fkbp8 |
| Cox8a | Rps29 | Gapdh |
| Ndufa4 | Cox17 | Gpr26 |
| Actb | Ndufa1 | Lmtk2 |
| Dpysl2 | Nrn1 | Lrrc17 |
| Ost4 | Tomm7 | Ndufa1 |
| Lrrc17 | Isca1 | Ost4 |
| Slc1a2 | Tatdn1 | Pfdn5 |
| Snhg6 | Snrpf | Ppp2r1a |
| Bc1 | Pacsin1 | Rpl14 |
| Lars2 | Actr2 | Rpl4 |
| Pgam1 | Arpp21 | Rpl6 |
| Rps2 | Araf | Rpl9-ps6 |
| Penk | Atp5md | Rps2 |
| Eef1a1 | Rps27 | Rps24 |
| Ddn | Sdhd | Rps27a |
| Rpl9-ps6 | Ost4 | Rps28 |
| Rpl14 | Atp6v1a | Rps29 |
| Mt1 | Sem1 | Rps8 |
| Pfn1 | Cdk5r2 | Snhg6 |
| Rps28 | Atp1a3 | Tspyl4 |
| Rps24 | Cox7c | Txn1 |
| Hsp90ab1 | Snhg9 | Usp50 |
| Comt | Snhg6 | |
| Ndufa1 | Eif4a2 | |
| Txn1 | Slc6a1 | |
| Ppp2r1a | Spag9 | |
| Arl8a | App | |
| Rps27a | Slc25a22 | |
| Eef1a2 | Snhg8 | |
| Rpl5 | Schip1 | |
| Rpl6 | Ddn | |
| Fkbp8 | Rgs7bp | |
| Usp50 | Comt | |
| Tspyl4 | Txn1 | |
| Gnb2 | Ndrg4 | |
| Rps8 | Smim26 | |
| Rps29 | Rnf5 | |
| Ddx5 | Zwint | |
| Tubb3 | Agap2 | |
| Ewsr1 | Naa20 | |
| Mical2 | Fam131b | |
| Dlg4 | Stxbp1 | |

| | |
|---|---|
| Cab39 | Bloc1s1 |
| Gpr26 | Wsb2 |
| Stub1 | Gucy1b1 |
| Rpl3 | Sema7a |
| BC005537 | Cops7a |
| Arc | Selenof |
| Rps15a | Acyp2 |
| Maged1 | Cdk17 |
| Ppp3r1 | Rheb |
| Nap1l5 | Huwe1 |
| Cinp | Srsf5 |
| Rgs20 | Cmpk1 |
| Lmtk2 | Lpcat4 |
| A830009L08Rik | 1810037I17Rik |
| Ptma | Pam |
| Atox1 | Usp50 |
| Cox7b | Ndufa3 |
| Ctxn1 | Mrpl33 |
| Junb | Rpl39 |
| Pfdn5 | Lmtk2 |
| Ptk2b | Fus |
| | Lmo4 |
| | B2m |
| | Arhgap39 |
| | Hmgb1 |
| | Psip1 |
| | Atn1 |
| | Psd3 |
| | Kcna2 |
| | Glul |
| | Atp6v1h |
| | Ndufb3 |
| | Hnrnpa2b1 |
| | Rab28 |
| | Unc13a |
| | Snhg3 |
| | Prrc2a |
| | Ank2 |
| | Fgfr1op2 |
| | Camk2a |
| | Ewsr1 |
| | Hprt |
| | Kctd13 |
| | Ubqln2 |
| | Slc4a10 |
| | Qk |
| | Ndfip2 |
| | Zfp365 |
| | Prkar1a |
| | Phyhipl |
| | Brms1l |
| | Oxr1 |

Cntnap1
Mgat3
Paip2
Ppp1r16b
Tmem258
Numb
Cd302
Serinc3
Atp5mpl
Shank1
Dnajc6
Megf9
Atp8a1
Btbd10
R3hdm4
Phpt1
Azin1
Unc5a
Gng13
Mapre2
Fbxl16
Herc3
Hlf
Cnksr2
Rpl37a
Capza2
Rab1a
Pde2a
Gas7
Golph3
Gria2
Rcn2
Mt2
Fam131a
Tmed7
Spry2
Cinp
Gja1
Chd9
Ank
Ptprz1
Vps29
Sec14l1
Anks1b
Ppm1h
Atp6ap2
Sft2d1
Gabra4
Ids
Dnajb5
Auts2
Bccip

Slc12a5
Kalrn
Gas5
Abhd8
Vxn
Btf3l4
Senp3
Slc4a4
Arhgap32
Wdr89
Rpl9-ps6
Stk25
Irf2bpl
Fez1
Rsrp1
Arhgef12
Mpv17
Cacng7
Lzts3
Rab3c
Scn1a
Rps27rt
Ogdhl
Arpp19
Hopx
Nedd4
Tma7
Ppp2cb
Dusp7
Plxna2
Sptbn2
Ccn3
Abhd3
Ssr3
Rap2a
Atxn2
Cox8a
Kitl
Tmx3
Car2
Tmem184b
Ncoa2
Gpr26
Deaf1
Saxo2
Dnajb4
Pmpcb
Atp5k
Ssbp1
Rai1
Nme3
Coa6

Cnnm1
Vstm2b
Atp8b2
Syngap1
Dmtf1
B230334C09Rik
Mrps28
Gtpbp2
Kif22
Vps50
Cyth1
Rps16
Hspa8
Rpl27a
Edf1
Nrsn1
Cst3
Tubb5
Mrfap1
Rps15
Fkbp8
Ndufb10
Clstn1
Nme2
Pkm
Saraf
Rps5
Ppp3ca
Rps11
Ctsd
Flywch1
Rpl30
Chgb
Ndufb8
Syp
Smdt1
Ppp2r1a
Rps12
Eef1a1
Rpl28
Pink1
Ubb
Rpl9
Rpl8
Rps10
Ndufa13
Eif3k
Psap
Rpl6
Rpl29
Nenf
Rps3a1

Rpl26
Fkbp2
Rps3
Rpsa
Rpl11
Rpl18a
Ap2b1
Pebp1
Pfdn5
Rps20
Rps7
Hbb-bs
Rps2
Rpl13a
Rpl13
Rpl24
Rpl14
Fau
Ndufb9
Rpl4
Eno1
Gapdh
Rpl21
Nefm
Rpl10
Rps27a
Rps8
Rps24

SS iTBS L4

| DEGs Young sham vs Aged sham | iTBS DEGs unique to young adults | iTBS DEGs unique to young adults that are also ageing DEGs |
|---|---|---|
| Gapdh | Rps27 | Rps27 |
| Uba52 | Pcp4 | Rps29 |
| Rpl4 | Tusc3 | |
| Camk2n1 | Rps29 | |
| Rps29 | Rpl10 | |
| Malat1 | Tmsb10 | |
| Nme7 | Tmsb4x | |
| Rps28 | Atp5md | |
| Lrrc17 | Ppia | |
| Lmtk2 | | |
| Dpysl2 | | |
| Rpl35 | | |
| Ost4 | | |
| Sema7a | | |
| Gpr26 | | |
| Bex2 | | |
| Lsm8 | | |
| Ndufa4 | | |
| Cacna1g | | |
| Rgs4 | | |
| Tspyl4 | | |
| Plcxd2 | | |
| Psd | | |
| Snhg6 | | |
| Rpl9-ps6 | | |
| Mical2 | | |
| Maged1 | | |
| Kcnab3 | | |
| Sel1l3 | | |
| Itm2c | | |
| Rps27rt | | |
| Dab2ip | | |
| Rraga | | |
| Banp | | |
| Schip1 | | |
| Bbs4 | | |
| Pkm | | |
| Actg1 | | |
| Mcee | | |
| Txn1 | | |
| Cd302 | | |
| Scn1a | | |
| Mtpn | | |
| Syndig1 | | |
| Medag | | |
| Rps27 | | |
| Car7 | | |
| Gstp1 | | |
| Mtch1 | | |

Calb1
Nrn1
Akt3
Grm2
Reep2
Ptpn5
Vcp
Cinp
Sdhd
Plec
Mbp
Ly6e
Gm10076
Krt10
Slc17a6
Ptpn4
Stx1a
Fam241b
Vegfa
Actr2
Rps2
Gaa
Stmn3
Lrrc57
Lrp11
Fam107a
Pantr1
Caprin1
Rabggta
Prkaca
Dynll2
Cnot2
Kcnj10
Dact2
Rnf181
Zfr
Pdia3
Dcdc2a
Serinc3
0610010K14Rik
Unc50
Tubb5
Stradb
Kansl1
Tln1
Bag5
Rims1
Inpp5a
Nisch
Usp50
Isca1
Slc25a23

SS cTBS L4

| DEGs Young sham vs Aged sham | cTBS DEGs unique to young adults | cTBS DEGs unique to young adults that are also ageing DEGs |
|---|---|---|
| Gapdh | Uba52 | Actr2 |
| Uba52 | Lrrc17 | Bbs4 |
| Rpl4 | Tspyl4 | Cacna1g |
| Camk2n1 | Rps28 | Cd302 |
| Rps29 | Basp1 | Dab2ip |
| Malat1 | Nrn1 | Gapdh |
| Nme7 | Rps29 | Gpr26 |
| Rps28 | Isca1 | Isca1 |
| Lrrc17 | Araf | Kansl1 |
| Lmtk2 | Snap25 | Lmtk2 |
| Dpysl2 | Snca | Lrrc17 |
| Rpl35 | Actr2 | Lrrc57 |
| Ost4 | Rps27 | Lsm8 |
| Sema7a | Tomm7 | Nrn1 |
| Gpr26 | Cox17 | Ost4 |
| Bex2 | Slc6a1 | Pkm |
| Lsm8 | Sdhd | Rpl4 |
| Ndufa4 | Ost4 | Rpl9-ps6 |
| Cacna1g | Ndufa1 | Rps2 |
| Rgs4 | Sem1 | Rps27 |
| Tspyl4 | Spag9 | Rps27rt |
| Plcxd2 | Snrpf | Rps28 |
| Psd | Cops7a | Rps29 |
| Snhg6 | Pacsin1 | Rraga |
| Rpl9-ps6 | Arpp21 | Schip1 |
| Mical2 | Snhg9 | Scn1a |
| Maged1 | 1810037I17Rik | Sdhd |
| Kcnab3 | Lmo4 | Sema7a |
| Sel1l3 | Dmtn | Snhg6 |
| Itm2c | Kcna2 | Stmn3 |
| Rps27rt | Phpt1 | Tspyl4 |
| Dab2ip | Snhg6 | Txn1 |
| Rraga | Snhg8 | Uba52 |
| Banp | Ank | Unc50 |
| Schip1 | Selenof | Usp50 |
| Bbs4 | Atp1a3 | Vegfa |
| Pkm | Lmtk2 | Zfr |
| Actg1 | Eif4a2 | |
| Mcee | App | |
| Txn1 | Paip2 | |
| Cd302 | Ppp1r16b | |
| Scn1a | Sema7a | |
| Mtpn | Rcn2 | |
| Syndig1 | Mgat3 | |
| Medag | Ddn | |
| Rps27 | Tatdn1 | |
| Car7 | Cox7c | |
| Gstp1 | Atp6v1a | |
| Mtch1 | Srsf5 | |

| | |
|---|---|
| Calb1 | Naa20 |
| Nrn1 | Gja1 |
| Akt3 | Car2 |
| Grm2 | Atp5md |
| Reep2 | Bloc1s1 |
| Ptpn5 | Schip1 |
| Vcp | Smim26 |
| Cinp | Rbis |
| Sdhd | Klf9 |
| Plec | Rnf5 |
| Mbp | Huwe1 |
| Ly6e | Txn1 |
| Gm10076 | Mat2b |
| Krt10 | Ndrg4 |
| Slc17a6 | Ewsr1 |
| Ptpn4 | Cd302 |
| Stx1a | Ank2 |
| Fam241b | Kif3c |
| Vegfa | Cacna2d1 |
| Actr2 | Oxr1 |
| Rps2 | Atp6v1h |
| Gaa | Inpp4a |
| Stmn3 | Zwint |
| Lrrc57 | Slc25a22 |
| Lrp11 | Rnf7 |
| Fam107a | Rgs7bp |
| Pantr1 | Cdk5r2 |
| Caprin1 | Smim10l1 |
| Rabggta | Nrxn1 |
| Prkaca | Scg2 |
| Dynll2 | Ndfip2 |
| Cnot2 | Ndufa3 |
| Kcnj10 | Scn1a |
| Dact2 | Arpc2 |
| Rnf181 | Gria2 |
| Zfr | Slc4a10 |
| Pdia3 | Vps29 |
| Dcdc2a | Zfr |
| Serinc3 | Comt |
| 0610010K14Rik | Ids |
| Unc50 | Rap1gds1 |
| Tubb5 | Unc5a |
| Stradb | Fus |
| Kansl1 | Nedd4 |
| Tln1 | Rusc2 |
| Bag5 | Dbndd2 |
| Rims1 | Gpr26 |
| Inpp5a | Fmc1 |
| Nisch | Camk2a |
| Usp50 | Ttc7b |
| Isca1 | Cntnap1 |
| Slc25a23 | Apba1 |

Sdcbp
Glul
Rpl39
Herc3
F3
Tmem258
Rheb
Rbfox3
Zfp365
Capza2
Eif5
Dpm3
Csnk1a1
Cs
Trappc6b
Tafa2
Rpl9-ps6
Phactr1
Acyp2
Hectd4
Hypk
Shank1
Csnk2a2
Atf2
Snn
Tsc22d4
Lsm8
Lrrc57
Ndufb3
Uba3
Cacna1g
Iscu
Ppp1r2
Slc12a5
Prrc2b
Hopx
Mapre2
Pet100
Dusp6
Rraga
Mapk1
Wsb2
Unc50
Fbxo34
Dab2ip
Usp50
Lpcat4
Foxk2
Senp3
Sgip1
Cystm1
Fbxl16

Cdk17
Gnai1
Cacng7
Rapgef4
Vegfa
Vstm2b
Trappc2
Atn1
Atxn2
Impact
Gas7
Fam120a
Bbs4
Brms1l
Car11
Kansl1
Chd9
Mir124a-1hg
Cds2
Hpcal4
Slc4a4
Rps27rt
Rpl37a
Arpp19
Idi1
Samd8
Btf3l4
Tceal1
Myrip
Mrpl3
Acat1
Med30
Churc1
Atxn2l
Hibadh
Klf6
Cox8a
Syt1
Gtpbp2
Ccnh
C130071C03Rik
Myadm
Clcn2
Atp5k
Acap2
Shd
Agk
Dusp7
Cspp1
Kif22
Fam98b
Cpeb2

Adam9
Taf1d
Cep104
Tac1
Eef1a1
Selenow
Rpl32
Psap
Rpl26
Stmn3
Rpl9
Cox4i1
Rpl30
Rps3
Tpt1
Cst3
Syp
Rps3a1
Rpl31
Hspa8
Tuba4a
Rpl17
Rpl18
Rps15
Rps10
Bsg
Aplp1
Atp5d
Rpl10
Rpl6
Pebp1
Rps7
Rpl28
Eef1b2
Rps11
Rps2
Rpl8
Ubb
Rpl11
Rpl13a
Ndufa13
Fkbp2
Pfdn5
Pkm
Rpl24
Rpsa
Nenf
Rpl18a
Chgb
Rpl4
Ndufb9
Rpl13

Rpl21
Eno1
Fau
Rps20
Rps8
Gapdh
Rpl14
Rps24
Rps27a

SS iTBS L5

| DEGs Young sham vs Aged sham | iTBS DEGs unique to young adults | iTBS DEGs unique to young adults that are also ageing DEGs |
|---|---|---|
| Gapdh | Penk | Penk |
| Uba52 | Eno1 | |
| Nme7 | Rps27 | |
| Rps20 | Tusc3 | |
| Rpl4 | Gpr88 | |
| Plcxd2 | Tac1 | |
| Ndufa4 | Gas5 | |
| Lrrc17 | Tmsb4x | |
| Rps24 | Gap43 | |
| Snhg6 | H3f3a | |
| Lmtk2 | Rpl17 | |
| Fmnl1 | Nr1d1 | |
| Fth1 | Tecr | |
| Zwint | Tatdn1 | |
| Malat1 | Nop53 | |
| Ost4 | Gm2000 | |
| Fau | Bc1 | |
| Rpl21 | | |
| Gpr26 | | |
| Itm2c | | |
| Rps27a | | |
| Rps2 | | |
| Pak1 | | |
| Reep1 | | |
| Ly6e | | |
| Aldoc | | |
| Panx2 | | |
| Dab2ip | | |
| Rps28 | | |
| Satb1 | | |
| Rpl9 | | |
| Rtn1 | | |
| Mical2 | | |
| Rtn4r | | |
| Ifngr2 | | |
| Usp50 | | |
| Vapa | | |
| Eef1a1 | | |
| St3gal3 | | |
| Nrep | | |
| Ptma | | |
| Rps8 | | |
| Fxyd7 | | |
| Fam107a | | |
| Sst | | |
| Kcnab3 | | |
| Sema7a | | |
| Shisa4 | | |
| Mef2c | | |

Oaz1
Prkaca
Atp1a3
Vsnl1
Atp6v1g2
Stx1a
Pfdn5
Homer1
Dock4
Napb
Arhgap32
Nlgn2
Dbi
Plxnd1
Atp2a2
Pacsin1
Scn1a
Rps4x
Syt1
Pip5k1c
Neurod6
Coro2b
Cinp
Schip1
Adcy1
Ccdc136
Btbd3
Slc6a7
Prdx1
Mir124a-1hg
Maz
Zfp938
Txn1
Tspyl4
Rpl5
Rpl14
Rps7
Sncb
Gnai1
Actb
Syn1
Ttll11
Lrp11
Chgb
Sdhd
Ppp6r1
Rpl18a
Nrip3
Sptbn2
Dkkl1
Atf4
Atp6v1a

Dok6
Rab6b
Cox7b
Nudt4
Shank1
Neurod2
Nlk
Sel1l3
Myo5a
Syndig1
Aldh1l1
Rps10
Stxbp1
Arc
Prrt1
Tmem145
Agap2
Rps29
Adora1
Kcnj10
Galnt9
Atg13
Atxn7l3
Dnajb2
Map6
Camk2n1
Nrn1
Igsf9b
Rph3a
Tln1
Dclk1
Cx3cl1
Slc30a3
R3hdm1
Eif3c
Psmg4
Cnih1
Arap2
Whrn
Hsd11b1
Ndfip1
Slc25a5
Opcml
Ppm1b
Atp8a1
Jdp2
Rpl22
Ablim1
Gtdc1
Eif4a2
Nrbp2
Bmpr2

Eif5a2
Ulk2
Dnm1
Penk
Esd
Gfod1
Mb21d2
Sv2b
Ptprd
Gpr158
Ephb6
Ppp1r13b
Porcn
Ptprk
Gria4
Myrip
Usp15
Car7
Pafah1b1
Kifc2
Nmt2
Dpysl4
Mpp3
Rgs4
Cpne9
Morf4l1
Tafa2
Atxn2
Luzp1
Syngr1
Ids
Comt
Mrpl55
Myl4
Rgs17
Sfxn3
Mpi
Tmx3
Senp2
Rubcn
Fgf13
Cacnb1
Chmp4b
Ddn
Camk2n2
Psd
Adnp
Ypel4
Nptxr
Shfl
Arf2
Rabl6

Arhgap20
Rae1
Lingo1
Ncald
Uba3
Tmbim6
Mfge8
Reep2
Hlf
Shank2
Lrpprc
Gm20300
Clip2
Ddost
Plekhj1
Il34
Rab3gap1
Pamr1
Hgs
Stac2
Hspa5
Arhgap44
Hacd3
Brsk2
Eipr1
Ttl
Extl2
Casc3
Rpl24
Gfra4
Slc35a4
Unc50
Arpc4
Tmem151a
Ube2ql1
Mcf2l
Asph
Prrc2a
Nsf
Snrpd2
Ncan
Naa60
Fkbp1b
Atxn1
Dnajb14
Twf2
Ankrd24
Cnbp
Chd9
Dpysl2
Ptprn
Pdhx

Brinp1
D430041D05Rik
Zdhhc2
Arxes2
Vps9d1
Ttc4
Sez6l
Rims4
Gbf1
Hecw1
Mapk8
Kif5c
Zranb2
Higd2a

SS cTBS L5

| DEGs Young sham vs Aged sham | cTBS DEGs unique to young adults | cTBS DEGs unique to young adults that are also ageing DEGs |
|---|---|---|
| Gapdh | Lrrc17 | Agap2 |
| Uba52 | Basp1 | Arhgap32 |
| Nme7 | Tspyl4 | Arpc4 |
| Rps20 | Rps28 | Atp1a3 |
| Rpl4 | Snca | Atp6v1a |
| Plcxd2 | Snap25 | Atxn1 |
| Ndufa4 | Pacsin1 | Atxn2 |
| Lrrc17 | Araf | Chd9 |
| Rps24 | Nrn1 | Cinp |
| Snhg6 | Ndufa1 | Comt |
| Lmtk2 | Ost4 | Dab2ip |
| Fmnl1 | Rps29 | Ddn |
| Fth1 | Dmtn | Dnm1 |
| Zwint | Atp6v1a | Dok6 |
| Malat1 | Tomm7 | Eif4a2 |
| Ost4 | Rps27 | Fau |
| Fau | Isca1 | Gapdh |
| Rpl21 | Sdhd | Gnai1 |
| Gpr26 | Cox17 | Gpr26 |
| Itm2c | App | Higd2a |
| Rps27a | Slc6a1 | Ifngr2 |
| Rps2 | Slc25a22 | Kifc2 |
| Pak1 | Actr2 | Lmtk2 |
| Reep1 | Snrpf | Lrrc17 |
| Ly6e | Zwint | Myo5a |
| Aldoc | Cops7a | Nrn1 |
| Panx2 | Smim26 | Ost4 |
| Dab2ip | Hprt | Pacsin1 |
| Rps28 | Snhg6 | Pfdn5 |
| Satb1 | Naa20 | Pip5k1c |
| Rpl9 | Atp1a3 | Prrc2a |
| Rtn1 | Wsb2 | Reep1 |
| Mical2 | Snhg9 | Rpl14 |
| Rtn4r | Atp5md | Rpl18a |
| Ifngr2 | Stxbp1 | Rpl21 |
| Usp50 | Cox7c | Rpl24 |
| Vapa | Sem1 | Rpl4 |
| Eef1a1 | Arpp21 | Rpl9 |
| St3gal3 | Ddn | Rps10 |
| Nrep | Spag9 | Rps2 |
| Ptma | Dnajc6 | Rps20 |
| Rps8 | Glul | Rps24 |
| Fxyd7 | Rap1gds1 | Rps27a |
| Fam107a | Rapgef4 | Rps28 |
| Sst | Eif4a2 | Rps29 |
| Kcnab3 | Cdk5r2 | Rps4x |
| Sema7a | Rgs7bp | Rps7 |
| Shisa4 | Rnf5 | Rps8 |
| Mef2c | Schip1 | Schip1 |

| | | |
|---|---|---|
| Oaz1 | Kcna2 | Scn1a |
| Prkaca | Tatdn1 | Sdhd |
| Atp1a3 | Ndrg4 | Shank1 |
| Vsnl1 | Car2 | Snhg6 |
| Atp6v1g2 | 1810037I17Rik | Sptbn2 |
| Stx1a | Frrs1l | Stxbp1 |
| Pfdn5 | Lmtk2 | Syngr1 |
| Homer1 | Phpt1 | Tmbim6 |
| Dock4 | Hpcal4 | Tspyl4 |
| Napb | Txn1 | Txn1 |
| Arhgap32 | Srsf5 | Usp50 |
| Nlgn2 | Snhg8 | Zwint |
| Dbi | Fus | |
| Plxnd1 | Klc2 | |
| Atp2a2 | Ank2 | |
| Pacsin1 | Rab3c | |
| Scn1a | Bloc1s1 | |
| Rps4x | Brms1l | |
| Syt1 | Cntnap1 | |
| Pip5k1c | Agap2 | |
| Neurod6 | Ndfip2 | |
| Coro2b | Rheb | |
| Cinp | Add2 | |
| Schip1 | Huwe1 | |
| Adcy1 | Tef | |
| Ccdc136 | Slc4a10 | |
| Btbd3 | Cadm4 | |
| Slc6a7 | Camk2a | |
| Prdx1 | Fam131a | |
| Mir124a-1hg | Mgat3 | |
| Maz | Rcn2 | |
| Zfp938 | Prrc2a | |
| Txn1 | Kctd13 | |
| Tspyl4 | Psd3 | |
| Rpl5 | Lmo4 | |
| Rpl14 | Syngr1 | |
| Rps7 | Herc3 | |
| Sncb | Kif3c | |
| Gnai1 | Zfp365 | |
| Actb | Smim10l1 | |
| Syn1 | Ppp1r2 | |
| Ttll11 | Usp50 | |
| Lrp11 | Tmbim6 | |
| Chgb | Ndufa3 | |
| Sdhd | Scn1a | |
| Ppp6r1 | Chd9 | |
| Rpl18a | Nedd4 | |
| Nrip3 | Phyhipl | |
| Sptbn2 | Acyp2 | |
| Dkkl1 | Erc2 | |
| Atf4 | Sptbn2 | |
| Atp6v1a | Selenof | |

| | |
|---|---|
| Dok6 | Fam131b |
| Rab6b | Chmp5 |
| Cox7b | Tsc22d4 |
| Nudt4 | Atp6v1h |
| Shank1 | Atp5mpl |
| Neurod2 | Oxr1 |
| Nlk | Zfr |
| Sel1l3 | Hectd4 |
| Myo5a | Atn1 |
| Syndig1 | Atp1b2 |
| Aldh1l1 | Paip2 |
| Rps10 | Csnk1e |
| Stxbp1 | R3hdm4 |
| Arc | Lpcat4 |
| Prrt1 | Gng13 |
| Tmem145 | Rpl39 |
| Agap2 | Kif5a |
| Rps29 | Ppp1r16b |
| Adora1 | Shank1 |
| Kcnj10 | Atxn2l |
| Galnt9 | Rnf157 |
| Atg13 | Smim14 |
| Atxn7l3 | Gnai1 |
| Dnajb2 | Tmem258 |
| Map6 | Ewsr1 |
| Camk2n1 | Ogdhl |
| Nrn1 | Rpl37a |
| Igsf9b | Hypk |
| Rph3a | Ifngr2 |
| Tln1 | Sec61g |
| Dclk1 | Scn2a |
| Cx3cl1 | F3 |
| Slc30a3 | Psip1 |
| R3hdm1 | Luzp2 |
| Eif3c | Cyth1 |
| Psmg4 | Ppp2r2c |
| Cnih1 | Cinp |
| Arap2 | Fbxl16 |
| Whrn | Prkar1a |
| Hsd11b1 | Unc5a |
| Ndfip1 | Apba1 |
| Slc25a5 | Ubqln2 |
| Opcml | Trappc2 |
| Ppm1b | B2m |
| Atp8a1 | Arpc4 |
| Jdp2 | Sft2d1 |
| Rpl22 | Phactr1 |
| Ablim1 | Gja1 |
| Gtdc1 | Emc7 |
| Eif4a2 | Jph4 |
| Nrbp2 | Pet100 |
| Bmpr2 | Rps27rt |

| | |
|---|---|
| Eif5a2 | Tppp |
| Ulk2 | Megf9 |
| Dnm1 | Arrb1 |
| Penk | Psma5 |
| Esd | Slc12a5 |
| Gfod1 | Gas7 |
| Mb21d2 | Rian |
| Sv2b | Gm14305 |
| Ptprd | Bcl2l2 |
| Gpr158 | Dnajb5 |
| Ephb6 | Tmed7 |
| Ppp1r13b | Atxn2 |
| Porcn | Azin1 |
| Ptprk | Reep1 |
| Gria4 | Scn4b |
| Myrip | Sucla2 |
| Usp15 | Rsrp1 |
| Car7 | Erh |
| Pafah1b1 | Idi1 |
| Kifc2 | Kifc2 |
| Nmt2 | Capza2 |
| Dpysl4 | Ank |
| Mpp3 | Mapk1 |
| Rgs4 | Gpr26 |
| Cpne9 | Scg2 |
| Morf4l1 | Mrps28 |
| Tafa2 | Atxn1 |
| Atxn2 | Sod2 |
| Luzp1 | Peg3 |
| Syngr1 | Necap1 |
| Ids | Higd2a |
| Comt | Gria2 |
| Mrpl55 | Insyn1 |
| Myl4 | Kcnh1 |
| Rgs17 | Arhgap32 |
| Sfxn3 | Senp3 |
| Mpi | Comt |
| Tmx3 | Cd302 |
| Senp2 | Rpl9-ps6 |
| Rubcn | Tmem59 |
| Fgf13 | Rora |
| Cacnb1 | Sparcl1 |
| Chmp4b | Plppr4 |
| Ddn | Ptprz1 |
| Camk2n2 | Sgip1 |
| Psd | Rusc2 |
| Adnp | Gas5 |
| Ypel4 | Tiprl |
| Nptxr | Lrrc4c |
| Shfl | Pip5k1c |
| Arf2 | Dab2ip |
| Rabl6 | Dnajb4 |

Arhgap20
Rae1
Lingo1
Ncald
Uba3
Tmbim6
Mfge8
Reep2
Hlf
Shank2
Lrpprc
Gm20300
Clip2
Ddost
Plekhj1
Il34
Rab3gap1
Pamr1
Hgs
Stac2
Hspa5
Arhgap44
Hacd3
Brsk2
Eipr1
Ttl
Extl2
Casc3
Rpl24
Gfra4
Slc35a4
Unc50
Arpc4
Tmem151a
Ube2ql1
Mcf2l
Asph
Prrc2a
Nsf
Snrpd2
Ncan
Naa60
Fkbp1b
Atxn1
Dnajb14
Twf2
Ankrd24
Cnbp
Chd9
Dpysl2
Ptprn
Pdhx

Vps29
Cystm1
Arpc2
Rap2a
Abhd8
Tac1
Slc4a4
Dnm1
Necab2
Aftph
Gnao1
Fez1
Map2k1
Fmc1
Dynlt3
Trrap
Frmpd4
Qk
Hnrnpk
Nme3
Vma21
Dok6
Ppip5k1
Fjx1
Btf3l4
Snn
Anks1b
Myo5a
Wdr89
Rps16
Rplp1
Rps19
Rps3a1
Pebp1
Rpl36al
Rpl27a
Pkm
Ctsd
Ndufb11
Rrp1
Rpl6
Ppp2r1a
Rpl7
Rpsa
Rps15
Tubb3
Calr
Rpl23
Eef1b2
Ndufb8
Tpt1
Smdt1

| | |
|---|---|
| Brinp1 | Rpl30 |
| D430041D05Rik | Atp5d |
| Zdhhc2 | Rpl28 |
| Arxes2 | Rps11 |
| Vps9d1 | Rps12 |
| Ttc4 | Syp |
| Sez6l | Ndufa13 |
| Rims4 | Fkbp2 |
| Gbf1 | Rpl11 |
| Hecw1 | Rpl4 |
| Mapk8 | Rpl9 |
| Kif5c | Tmem160 |
| Zranb2 | Eno1 |
| Higd2a | Rps4x |
| | Ttr |
| | Fkbp8 |
| | Rpl8 |
| | Edf1 |
| | Rpl26 |
| | Gapdh |
| | Rpl13 |
| | Rps7 |
| | Rps2 |
| | Ndufb9 |
| | Ndufb10 |
| | Rpl21 |
| | Rpl18a |
| | Rpl14 |
| | Rps20 |
| | Rps10 |
| | Pfdn5 |
| | Rpl24 |
| | Rps8 |
| | Fau |
| | Rpl13a |
| | Rpl10 |
| | Pvalb |
| | Rps27a |
| | Rps24 |
| | Hbb-bs |

SS iTBS L6

| DEGs Young sham vs Aged sham | iTBS DEGs unique to young adults | iTBS DEGs unique to young adults that are also ageing DEGs |
|---|---|---|
| Gapdh | Rps29 | Gapdh |
| Uba52 | Rps27 | Gas5 |
| Pcp4 | Gas5 | Gstp1 |
| Mbp | Rps28 | Pcp4 |
| Lrrc17 | Pcp4 | Penk |
| Cox8a | Rpl37a | Rpl35 |
| Mobp | Tmsb4x | Rps27 |
| Snhg6 | Rpl37a | Rps28 |
| Rpl4 | Ftl1 | Rps29 |
| Rps28 | Penk | Sem1 |
| Rps29 | Tma7 | Tmsb4x |
| Tmsb4x | Dbi | |
| Psd | Tac1 | |
| Ost4 | Rpl22l1 | |
| Pkm | Atp5md | |
| Klf9 | Ndufb3 | |
| Rpl21 | Rpl39 | |
| Tatdn1 | Gpr88 | |
| Slc1a2 | Sem1 | |
| Kif5a | Rps15a | |
| Bcas1 | Mrpl42 | |
| Eef1a1 | Tmsb10 | |
| Cox7b | Tuba1a | |
| Usp50 | Igfbp6 | |
| Clic4 | Rpusd1 | |
| Lpcat4 | Lynx1 | |
| Dpysl2 | Myl4 | |
| Dusp6 | Gapdh | |
| Ndfip2 | Cplx2 | |
| Ndufa1 | Apba1 | |
| Chd9 | Tubb4a | |
| Rps24 | Cdk5r2 | |
| Arc | Klc2 | |
| Isca1 | | |
| Mical2 | | |
| Twf2 | | |
| Ighm | | |
| Polr3e | | |
| Txn1 | | |
| Cldn11 | | |
| Fau | | |
| Pfdn5 | | |
| Atraid | | |
| Mal | | |
| Gadd45a | | |
| Hivep2 | | |
| Rps27 | | |
| Comt | | |
| Actg1 | | |

Rpl14
Schip1
Cspg5
Agap2
Smim26
Ptp4a3
Ctxn1
Gm10076
Shisa4
Gstp1
Tubb5
Tspyl4
Mrpl27
Rps2
Malat1
Hs3st2
Dlg2
Ccm2
Sema7a
Rpl37a
Sgsm1
Bloc1s1
Ccdc12
6330403K07Rik
Eef1akmt1
Stmn3
Nbea
Ina
Atxn7l3b
Exoc6
Ddn
Tsc22d3
Arhgap33
Eif3c
Eef1a2
Unc50
Chn2
Zfp365
Slc4a10
Acap3
Actb
Lrp11
Uba3
Pacsin1
Epas1
Nae1
S1pr1
Rasl10a
Coa6
Srcin1
Snhg9
Jagn1

Syt6
Fgf13
Tmbim4
Ly6e
Hpcal4
Golga7
Grb14
Fxyd7
Homer1
Arhgef7
Ncald
Immp1l
Ube2j1
Vxn
Bex2
Lsm8
Fut9
Fmnl1
Tmem151b
Tomm7
Kif5c
Trp53inp2
Nsg2
Ntsr2
Acyp2
Slirp
Snrpf
Kcnj10
Rims2
B3galt2
Tmem167
Rpl5
Gnai1
Sdr39u1
Prrc2a
Mog
Slc8a1
Emc3
Grsf1
Cplx3
Fyn
Ddah1
Phldb1
Gas5
Cfap36
Mrpl35
Kcnip4
Fkbp8
Rmnd5a
Epn2
Nxf1
Ttc9b

Wdr13
Cdc123
Cd200
Cdc37l1
Usp31
Rab15
Actl6b
Tnfrsf21
Abhd16a
Eif4a2
Tbc1d9b
Polr2f
Crmp1
Polr2i
Prkaca
Cops7a
Reep1
Sdhd
Plxna2
Sipa1l1
Rnmt
Sdhaf2
Lamtor3
Hipk3
Ak1
Tspan2
Chrm3
Tmeff2
Trak2
Rap1gap2
Rnasek
Maged1
Mtx1
Kcnh7
Itm2c
Vps50
Zfand2a
Nr1h2
Rraga
Cox16
Rpl35
Dnajb4
Ube2w
Necab3
Snhg8
Abhd17a
Golph3
Igsf21
Tmem135
Psip1
Zfp445
Ppp2r5a

Ugp2
Slc32a1
Cox20
Zfas1
Thra
Ermn
Sem1
Elmo2
Ntrk3
Slc2a1
Mfsd6
Wnk1
Pcna
Btbd1
Aopep
Zdhhc17
Tmem208
Abracl
Basp1
Nisch
Snhg12
Brwd1
Setd7
Rps8
9130401M01Rik
Ccnh
Plpbp
Naa20
Nsa2
Pnpla8
Pdlim2
Ubl7
Gfod1
Plppr4
Kcnc4
Tfg
Usp32
Limk1
R3hdm4
Rer1
Rbm3
Pomgnt1
Slc12a2
Trbc2
Taf13
Serpinb6a
Fez2
Pitpnc1
Adora1
Ndufv1
Vcp
Large1

Rala
Epb41l1
Clec2l
Pou3f3
Pdpk1
Cisd3
Tmem126a
Penk
Gjb6
Necab1
Frrs1l
Maz
Igsf9b
Paqr4
AI593442
Commd6
Otud5
Spop
Fa2h
Abce1
Arhgef12
Elof1
Trpc4ap
Dok6
Bcap31
Mpv17
Tmem50b
Snrpc
Clns1a
Tubb3
Slmap
Wdr47
Rprd1a
Shank2
Map4k3
A830018L16Rik
Vstm2l
Pltp
Ptprz1
Ube3a
Pdcd10
Tsg101
Ubl4a
Ly6c1
Snapc5
St8sia5
Tln1
Cckbr
Becn1
Panx2
Kcnc1
Acsl6

Src
Fxyd1
Neurod6
Mxi1
Arhgap23
Carm1
Nme7
Zmynd8
Rab5if
Anp32b
Rps27l
Mgat4b
Asrgl1
Etv5
Arxes2
Rab28
Rida
Mir124a-1hg
Clstn2
Trim28
Srsf10
Dop1b
Dctn4
Gpd1l
Tnk2
Rcn2
Pik3r3
Slc27a4
Med10
Rab3gap1
Dagla
Manf
Zyx
Enpp2
BC004004
Amz2
Gaa
Uba2
Tacc1
B3gat1
Slc25a23
Eid2
Ppib
Scarb2
Supt6
Gpd1
Dtnb
Arl6ip1
Mxd4
Negr1
Pcmtd2
Emc2

Slc25a18
Higd2a
Hdhd2
Chka
Sprn
Cryab
Prpf8
Slc8a2
Atp2c1
Speg
Arfgef1
Cstb
Klf13
Atg4b
Acyp1
Ddx1

SS cTBS L6

| DEGs Young sham vs Aged sham | cTBS DEGs unique to young adults | cTBS DEGs unique to young adults that are also ageing DEGs |
|---|---|---|
| Gapdh | Uba52 | Acyp2 |
| Uba52 | Lrrc17 | Agap2 |
| Pcp4 | Rps28 | Basp1 |
| Mbp | Basp1 | Bloc1s1 |
| Lrrc17 | Tspyl4 | Chd9 |
| Cox8a | Rps29 | Coa6 |
| Mobp | Cox17 | Comt |
| Snhg6 | Ndufa1 | Cops7a |
| Rpl4 | Tomm7 | Cox8a |
| Rps28 | Snap25 | Cryab |
| Rps29 | Araf | Ddn |
| Tmsb4x | Rps27 | Dnajb4 |
| Psd | Pacsin1 | Eef1a1 |
| Ost4 | Snca | Eif4a2 |
| Pkm | Snrpf | Fau |
| Klf9 | Isca1 | Fkbp8 |
| Rpl21 | Slc6a1 | Gapdh |
| Tatdn1 | Arpp21 | Gas5 |
| Slc1a2 | Sem1 | Hpcal4 |
| Kif5a | Ost4 | Isca1 |
| Bcas1 | Car2 | Itm2c |
| Eef1a1 | Rapgef4 | Kif5a |
| Cox7b | App | Kif5c |
| Usp50 | Snhg9 | Klf9 |
| Clic4 | Tatdn1 | Lpcat4 |
| Lpcat4 | Ndrg4 | Lrp11 |
| Dpysl2 | Atp6v1a | Lrrc17 |
| Dusp6 | Nrn1 | Lsm8 |
| Ndfip2 | Glul | Mbp |
| Ndufa1 | Txn1 | Naa20 |
| Chd9 | Smim26 | Nae1 |
| Rps24 | Atp1a3 | Ndfip2 |
| Arc | Dmtn | Ndufa1 |
| Isca1 | Actr2 | Ost4 |
| Mical2 | Eif4a2 | Pacsin1 |
| Twf2 | Ddn | Pfdn5 |
| Ighm | Srsf5 | Pkm |
| Polr3e | Selenof | Plppr4 |
| Txn1 | Cops7a | Prrc2a |
| Cldn11 | Cox7c | R3hdm4 |
| Fau | Rgs7bp | Rab28 |
| Pfdn5 | Bloc1s1 | Rcn2 |
| Atraid | Naa20 | Rpl14 |
| Mal | Atp5md | Rpl21 |
| Gadd45a | Ndfip2 | Rpl37a |
| Hivep2 | Slc25a22 | Rpl4 |
| Rps27 | Wsb2 | Rpl5 |
| Comt | Snhg6 | Rps2 |
| Actg1 | Camk2a | Rps24 |

Rpl14
Schip1
Cspg5
Agap2
Smim26
Ptp4a3
Ctxn1
Gm10076
Shisa4
Gstp1
Tubb5
Tspyl4
Mrpl27
Rps2
Malat1
Hs3st2
Dlg2
Ccm2
Sema7a
Rpl37a
Sgsm1
Bloc1s1
Ccdc12
6330403K07Rik
Eef1akmt1
Stmn3
Nbea
Ina
Atxn7l3b
Exoc6
Ddn
Tsc22d3
Arhgap33
Eif3c
Eef1a2
Unc50
Chn2
Zfp365
Slc4a10
Acap3
Actb
Lrp11
Uba3
Pacsin1
Epas1
Nae1
S1pr1
Rasl10a
Coa6
Srcin1
Snhg9
Jagn1

Phpt1
Zfp365
Rap1gds1
Rpl39
Rnf5
Sdhd
Agap2
Schip1
Zwint
Ppp1r2
Klf9
Cdk5r2
Slc4a10
Spag9
Ccn2
Kif5a
Fus
Snhg8
Lpcat4
Kctd13
Ndufa3
Rcn2
Nedd4
Fbxl16
Hprt
Gpm6b
Comt
Unc5a
Tsc22d3
Paip2
Ank2
Hmgb1
Vxn
Sipa1l1
Usp50
1810037I17Rik
Gria2
Chd9
Slc12a5
Ids
Sparcl1
Sptbn1
Ewsr1
Apc
R3hdm4
Acyp2
Ppp1r16b
Fam131b
Slc1a3
Gnao1
Plppr4
Gpr88

Rps27
Rps28
Rps29
Rps8
S1pr1
Schip1
Sdhd
Sem1
Sema7a
Sipa1l1
Slc4a10
Smim26
Snhg6
Snhg8
Snhg9
Snrpf
Tatdn1
Tomm7
Tsc22d3
Tspyl4
Ttc9b
Tubb5
Txn1
Uba52
Ugp2
Usp50
Vxn
Zfp365
Zfp445

| | |
|---|---|
| Syt6 | Scn4b |
| Fgf13 | Stxbp1 |
| Tmbim4 | Lrp11 |
| Ly6e | Mapk1 |
| Hpcal4 | Rpl37a |
| Golga7 | Ndufb3 |
| Grb14 | Gad1 |
| Fxyd7 | Cmpk1 |
| Homer1 | Fez1 |
| Arhgef7 | Gas5 |
| Ncald | Inpp4a |
| Immp1l | Romo1 |
| Ube2j1 | Huwe1 |
| Vxn | Phyhipl |
| Bex2 | Rheb |
| Lsm8 | Ppp2r2c |
| Fut9 | Tmem258 |
| Fmnl1 | Cs |
| Tmem151b | Cadm4 |
| Tomm7 | Arpc2 |
| Kif5c | Rab3c |
| Trp53inp2 | Gng13 |
| Nsg2 | Hnrnpa2b1 |
| Ntsr2 | Atp1b2 |
| Acyp2 | Ttyh1 |
| Slirp | Ubqln2 |
| Snrpf | Atp5mpl |
| Kcnj10 | Kcna2 |
| Rims2 | Psma5 |
| B3galt2 | Nap1l2 |
| Tmem167 | Ctnnb1 |
| Rpl5 | Ppp1r1b |
| Gnai1 | Selenop |
| Sdr39u1 | Vps29 |
| Prrc2a | Dnajc6 |
| Mog | Rusc2 |
| Slc8a1 | Rps27rt |
| Emc3 | Sema7a |
| Grsf1 | Rgs4 |
| Cplx3 | Sec61g |
| Fyn | Ank |
| Ddah1 | Rsrp1 |
| Phldb1 | Hpcal4 |
| Gas5 | Tac1 |
| Cfap36 | Prkar1a |
| Mrpl35 | Kif5c |
| Kcnip4 | Atxn2 |
| Fkbp8 | Lsm8 |
| Rmnd5a | Abhd8 |
| Epn2 | Impact |
| Nxf1 | Idi1 |
| Ttc9b | Sod2 |

| | |
|---|---|
| Wdr13 | Rpl9-ps6 |
| Cdc123 | Tmem256 |
| Cd200 | Lmtk2 |
| Cdc37l1 | Mapre2 |
| Usp31 | Anks1b |
| Rab15 | Ccnb1 |
| Actl6b | Cox8a |
| Tnfrsf21 | Herc3 |
| Abhd16a | Bcl2l2 |
| Eif4a2 | Pet100 |
| Tbc1d9b | Smim10l1 |
| Polr2f | Mrpl33 |
| Crmp1 | Rasd2 |
| Polr2i | Dnajb5 |
| Prkaca | Oxr1 |
| Cops7a | Gng7 |
| Reep1 | Ppp1r9b |
| Sdhd | Hectd4 |
| Plxna2 | Sgip1 |
| Sipa1l1 | Prrc2a |
| Rnmt | Coa6 |
| Sdhaf2 | Phactr1 |
| Lamtor3 | Ugp2 |
| Hipk3 | Zfp445 |
| Ak1 | Sst |
| Tspan2 | Laptm4a |
| Chrm3 | Enpp5 |
| Tmeff2 | Atn1 |
| Trak2 | F3 |
| Rap1gap2 | Camk2n1 |
| Rnasek | Capza2 |
| Maged1 | Adcy5 |
| Mtx1 | Aco2 |
| Kcnh7 | Ppp2cb |
| Itm2c | Nptx2 |
| Vps50 | Slc4a4 |
| Zfand2a | Ube2d1 |
| Nr1h2 | Rpl22l1 |
| Rraga | Sft2d1 |
| Cox16 | Nae1 |
| Rpl35 | Morf4l1 |
| Dnajb4 | Cystm1 |
| Ube2w | Rab28 |
| Necab3 | S1pr1 |
| Snhg8 | Dnajb4 |
| Abhd17a | Med30 |
| Golph3 | Map2k1 |
| Igsf21 | Bpnt1 |
| Tmem135 | Mthfsl |
| Psip1 | Senp3 |
| Zfp445 | Rpl10-ps3 |
| Ppp2r5a | Mbp |

| | |
|---|---|
| Ugp2 | Wdr89 |
| Slc32a1 | Mrps28 |
| Cox20 | Necab2 |
| Zfas1 | Nme3 |
| Thra | Atxn2l |
| Ermn | Dync1h1 |
| Sem1 | Pcsk1n |
| Elmo2 | Eif3k |
| Ntrk3 | Rps14 |
| Slc2a1 | Ndufb8 |
| Mfsd6 | Atp5d |
| Wnk1 | Ndufs6 |
| Pcna | Tubb5 |
| Btbd1 | Cd81 |
| Aopep | Ubb |
| Zdhhc17 | Nrgn |
| Tmem208 | Yars |
| Abracl | Rps16 |
| Basp1 | Rangap1 |
| Nisch | Cst3 |
| Snhg12 | Eef1a1 |
| Brwd1 | Ndufb10 |
| Setd7 | Fkbp2 |
| Rps8 | Tpt1 |
| 9130401M01Rik | Pomp |
| Ccnh | Uchl1 |
| Plpbp | Smdt1 |
| Naa20 | Ppp2r1a |
| Nsa2 | Cryab |
| Pnpla8 | Ndufb11 |
| Pdlim2 | Rpl18 |
| Ubl7 | Rpl30 |
| Gfod1 | Hspa8 |
| Plppr4 | Rplp1 |
| Kcnc4 | Tle5 |
| Tfg | Rpl29 |
| Usp32 | Ctsd |
| Limk1 | Rpl5 |
| R3hdm4 | Rpl17 |
| Rer1 | Rpl32 |
| Rbm3 | Clstn1 |
| Pomgnt1 | Rps3a1 |
| Slc12a2 | Gnaz |
| Trbc2 | Atp9a |
| Taf13 | Psap |
| Serpinb6a | Rps4x |
| Fez2 | Rps12 |
| Pitpnc1 | Myl4 |
| Adora1 | Rpl7 |
| Ndufv1 | Chgb |
| Vcp | Rpl26 |
| Large1 | Cox4i1 |

Rala
Epb41l1
Clec2l
Pou3f3
Pdpk1
Cisd3
Tmem126a
Penk
Gjb6
Necab1
Frrs1l
Maz
Igsf9b
Paqr4
AI593442
Commd6
Otud5
Spop
Fa2h
Abce1
Arhgef12
Elof1
Trpc4ap
Dok6
Bcap31
Mpv17
Tmem50b
Snrpc
Clns1a
Tubb3
Slmap
Wdr47
Rprd1a
Shank2
Map4k3
A830018L16Rik
Vstm2l
Pltp
Ptprz1
Ube3a
Pdcd10
Tsg101
Ubl4a
Ly6c1
Snapc5
St8sia5
Tln1
Cckbr
Becn1
Panx2
Kcnc1
Acsl6
Rps11
Ppp1r1a
Nenf
Tmem160
Rpl6
Map1lc3a
Pebp1
Rpl28
Rpl11
Ndufa13
Rps5
Pkm
Itm2c
Fkbp8
Cck
Rpsa
Rps3
Rpl24
Ndufb9
Rps10
Rps7
Rps2
Rpl8
Rpl9
Ttc9b
Rps15
Rps20
Rpl13
Rps27a
Pfdn5
Hbb-bs
Rpl4
Rpl13a
Rpl18a
Rpl21
Ttr
Fau
Gapdh
Rpl14
Rps8
Rpl10
Rps24

Src
Fxyd1
Neurod6
Mxi1
Arhgap23
Carm1
Nme7
Zmynd8
Rab5if
Anp32b
Rps27l
Mgat4b
Asrgl1
Etv5
Arxes2
Rab28
Rida
Mir124a-1hg
Clstn2
Trim28
Srsf10
Dop1b
Dctn4
Gpd1l
Tnk2
Rcn2
Pik3r3
Slc27a4
Med10
Rab3gap1
Dagla
Manf
Zyx
Enpp2
BC004004
Amz2
Gaa
Uba2
Tacc1
B3gat1
Slc25a23
Eid2
Ppib
Scarb2
Supt6
Gpd1
Dtnb
Arl6ip1
Mxd4
Negr1
Pcmtd2
Emc2

Slc25a18
Higd2a
Hdhd2
Chka
Sprn
Cryab
Prpf8
Slc8a2
Atp2c1
Speg
Arfgef1
Cstb
Klf13
Atg4b
Acyp1
Ddx1

CP iTBS

| DEGs Young sham vs Aged sham | iTBS DEGs unique to young adults | iTBS DEGs unique to young adults that are also ageing DEGs |
|---|---|---|
| Gm10076 | Rps27 | Lrrc17 |
| Uba52 | Rps29 | Rps27 |
| Rpl35 | Tmsb4x | Rps29 |
| Gstp1 | Pcp4 | Tatdn1 |
| Rps27 | Tusc3 | |
| Rps29 | Rpl10 | |
| Tmsb4x | Gas5 | |
| Pcp4 | Hmgn2 | |
| Tusc3 | Ftl1 | |
| Rpl10 | Tatdn1 | |
| Gas5 | Gm2000 | |
| Hmgn2 | Lrrc17 | |
| Ftl1 | | |
| Tatdn1 | | |
| Gm2000 | | |
| Lrrc17 | | |
| Bc1 | | |

CP cTBS

| DEGs Young sham vs Aged sham | cTBS DEGs unique to young adults | cTBS DEGs unique to young adults that are also ageing DEGs |
|---|---|---|
| Gapdh | Lrrc17 | AI593442 |
| Uba52 | Pde10a | Acat1 |
| Rpl4 | Rps28 | Acyp2 |
| Rps29 | Scn4b | Add3 |
| Lrrc17 | Rps29 | Agap2 |
| Mbp | Ndufa1 | Anapc16 |
| Rps28 | Rps27 | Ank |
| Lzts3 | Arpp21 | Anks1b |
| Atp1a1 | Tatdn1 | Ano3 |
| Slc17a7 | Araf | Ap1ar |
| Cox8a | Ost4 | Apbb1 |
| Rpl21 | Tspyl4 | Araf |
| Snhg6 | Snca | Arhgef12 |
| Mobp | Actr2 | Arpp21 |
| Stmn3 | Snrpf | Atp1b2 |
| Fau | Snhg9 | B2m |
| Rps24 | Cox17 | Basp1 |
| Rps27 | Ddn | Bloc1s1 |
| Rpl5 | Isca1 | Bzw1 |
| Ywhah | Sem1 | Cacna2d3 |
| Calm1 | Snap25 | Cbr3 |
| Ndufa4 | Ppp1r16b | Cck |
| Gng7 | Rgs7bp | Ccng2 |
| Eef1a2 | Tomm7 | Cd302 |
| Malat1 | Agap2 | Cdk5r2 |
| Ppp1r1b | Snhg8 | Cdr1os |
| Olfm1 | Plppr4 | Cfap36 |
| Ivns1abp | Ndrg4 | Chst2 |
| Bcas1 | Bloc1s1 | Churc1 |
| Bex2 | Cox7c | Cinp |
| Vsnl1 | Sdhd | Clstn1 |
| Pkm | Snhg6 | Cnp |
| Dpysl2 | Atp5md | Comt |
| Tuba1b | Slc6a1 | Cox16 |
| Ost4 | Dmtn | Cox8a |
| Usp50 | Basp1 | Crym |
| Rpl9-ps6 | Glul | Cystm1 |
| Ddn | Rnf5 | Ddn |
| Tubb5 | Ndfip2 | Dhx15 |
| Rasd2 | Eif4a2 | Dnajb4 |
| Itm2c | Spag9 | Dock4 |
| Tmsb10 | App | Dync1i2 |
| Lmtk2 | Klf9 | Eno1 |
| Psap | Cdk5r2 | Erh |
| Scn4b | Pacsin1 | Ewsr1 |
| Slc1a2 | Lzts3 | Fam120a |
| Ctxn1 | Fbxl16 | Fam174a |
| Gng3 | Smim26 | Fau |
| Ly6c1 | Car2 | Fgfr1op2 |

| | | |
|---|---|---|
| Mal | Comt | Fis1 |
| Zdhhc14 | Naa20 | Fkbp8 |
| Comt | Rpl39 | Fth1 |
| Rps20 | Lpcat4 | Gapdh |
| Rgs20 | Meis2 | Gja1 |
| Cck | Ppp1r2 | Gm33651 |
| Psd | Ido1 | Gng7 |
| Ndufa1 | Atp1a3 | Gpm6b |
| Tatdn1 | Slc4a4 | Hdhd2 |
| Rarb | Huwe1 | Hnrnpa2b1 |
| Rpl35 | Gucy1b1 | Id4 |
| Rps4x | Phactr1 | Ido1 |
| Unc13c | Wsb2 | Ids |
| Gstp1 | Ewsr1 | Insyn1 |
| Rps27rt | Rab3c | Isca1 |
| Bloc1s1 | Cd302 | Jph4 |
| Rpl3 | Phpt1 | Kazn |
| Mog | Txn1 | Klf13 |
| Pfdn5 | Psip1 | Klf9 |
| Cldn11 | Fus | Klhl7 |
| Rpl37a | Zwint | L1cam |
| Pgam1 | Cops7a | Lmtk2 |
| P4ha1 | Srsf5 | Lpcat4 |
| Trak2 | 1810037I17Rik | Lrp11 |
| Sema7a | Cntnap1 | Lrrc17 |
| Pde10a | Caln1 | Lrrk2 |
| Rgs4 | Rgs14 | Lsm8 |
| Actb | Spry2 | Luzp2 |
| Ppp2r1a | Ap1ar | Lzts3 |
| Crym | Usp50 | Mapre2 |
| Prkar1b | Kctd13 | Meis2 |
| Gpm6b | Ank2 | Morf4l1 |
| Trp53inp2 | Rapgef4 | Mrpl27 |
| Tubb3 | Rps27rt | Mrpl33 |
| Epas1 | Atp6v1a | Mrps28 |
| Cinp | Sipa1l1 | Mt2 |
| Acyp2 | Tsc22d4 | Naa20 |
| Grb14 | Rpl9-ps6 | Ndfip2 |
| Gm10076 | Rgs20 | Ndufa1 |
| Dnm1 | Cs | Nenf |
| Hspa5 | Rarb | Neto1 |
| Ido1 | Slc25a22 | Ost4 |
| Gnas | Id4 | Pacsin1 |
| Gpr88 | Ndufa3 | Pcdh17 |
| Adcy5 | Zfp365 | Pdcd10 |
| Slc2a1 | Gabra4 | Pde10a |
| Vcp | Mapre2 | Pde4b |
| Rps2 | Chst2 | Pde4dip |
| Adora2a | Prrc2a | Pet100 |
| Pdlim2 | Jph4 | Pfdn5 |
| Arl6ip1 | Selenof | Phactr1 |
| Plppr4 | Dnajb4 | Pkm |

| | | |
|---|---|---|
| Tspyl4 | Acyp2 | Plp1 |
| Rpl24 | Cacna2d3 | Plppr4 |
| Mt1 | Arf4 | Ppp1r16b |
| Tspan2 | Cd47 | Ppp2r1a |
| Cap1 | Spop | Pptc7 |
| Cd302 | Tmem258 | Prpf8 |
| Eno1 | Cdk17 | Prrc2a |
| Ablim2 | Gng7 | Psap |
| Rraga | Rpl37a | Psd3 |
| Ndufa7 | Churc1 | Psma5 |
| Txn1 | Rab28 | Ptprz1 |
| Akap9 | Rcn2 | Rab28 |
| Ly6a | Ubqln2 | Rai1 |
| 6330403K07Rik | Insyn1 | Rapgef2 |
| Rps7 | Ano3 | Rapgef4 |
| Gamt | Ptprz1 | Rarb |
| Sirt2 | Scg2 | Reep1 |
| Dynll2 | Camk2a | Rgs14 |
| Tmeff2 | Tmed7 | Rgs20 |
| Rabac1 | Sdcbp | Rian |
| Polb | Hprt | Rnf5 |
| Cdr1os | Acat1 | Rpl14 |
| Fkbp8 | Ids | Rpl21 |
| Agap2 | Psd3 | Rpl24 |
| Acot7 | Shank3 | Rpl37a |
| Apbb1 | Rheb | Rpl39 |
| Tubb4b | B230334C09Rik | Rpl4 |
| Mag | Mt2 | Rpl5 |
| Nptxr | Atp5mpl | Rpl6 |
| Slc22a23 | Ppp1r9b | Rpl9-ps6 |
| Slc41a1 | Sparcl1 | Rps2 |
| Mir124a-1hg | Lmtk2 | Rps20 |
| Lingo3 | Mapk1 | Rps24 |
| Arhgap33 | Gja1 | Rps27 |
| Rgs14 | Paip2 | Rps27rt |
| Golga7 | Dock4 | Rps28 |
| Dock4 | Spock3 | Rps29 |
| Rpl39 | Atxn2l | Rps3a1 |
| Pet100 | Cfap20 | Rps4x |
| Hnrnpa2b1 | Gria2 | Rps7 |
| Kcnab1 | Sowaha | Rraga |
| Ppp3r1 | Strn | Rrp1 |
| Tlnrd1 | Slc4a10 | Scn4b |
| Wnk1 | Phyhip | Sdhd |
| Syp | Anks1b | Sec14l1 |
| Prnp | Sft2d1 | Sem1 |
| Actn1 | Smim14 | Sgip1 |
| Dlg2 | Lmo4 | Sipa1l1 |
| Fam120a | Pet100 | Slc4a4 |
| Josd2 | Romo1 | Smim26 |
| Lpcat4 | Fam174a | Snca |
| Stmn1 | Erh | Snhg6 |

Coro2b
Nme7
Mrpl33
Snhg9
Timm22
Nisch
Car12
Gng5
Tiam2
Slc32a1
Ano3
Clstn1
Arc
Pdzd2
Ugp2
Acat1
Sbk1
Dnajb11
Pde4dip
Cacna2d3
Mbnl2
Npdc1
Rasgef1b
Reep1
Atp6v1b2
Snca
Psenen
Klf9
Rpl6
Synpr
Sdhd
Smim27
Neto2
Slain1
Rpl14
Fa2h
Chst2
Ctsb
Nap1l5
Prkar2b
Gnb1
Slco1c1
Clic4
Sncb
L1cam
Ntsr2
Chordc1
Kcnq3
Ppm1b
Syn1
Fxyd2
Ostc
Hnrnpa2b1
Sgip1
Neto1
Rapgef2
Wdr89
Psma5
Tmeff1
Gas7
Stxbp1
Arpc2
Kazn
Gm33651
Mrpl27
Tgfa
Mrps28
Atp6v1h
Tacc1
Atp5g3
Brms1l
Fgfr1op2
Senp3
Ccng2
Pde4dip
Gad1
F3
Rragd
Crym
Chmp5
Frrs1l
Usp9x
R3hdm4
Klhl9
Cdr1os
Prkar1a
Trappc6b
Sec14l1
Atp1b2
Tmbim6
L1cam
Bzw1
Mrpl33
Dgkb
AI593442
Nedd4
Cmpk1
Gpm6b
Klf13
Add2
Morf4l1
Lmo3
Rai1
Arhgef12
Snhg8
Snhg9
Sowaha
Spop
Spry2
Ssr3
Stmn3
Strn
Tatdn1
Tgfa
Tmbim6
Tmeff1
Trf
Trpc1
Tspyl4
Txn1
U2af2
Unc50
Unc5a
Usp50
Usp9x
Vgf
Zfr

| | |
|---|---|
| Atg3 | Mat2b |
| Csf1r | Zcchc14 |
| Thy1 | Rian |
| Hmgcs1 | Ndufb3 |
| Meis2 | Mgat3 |
| Plekhj1 | Rpl22l1 |
| Tle4 | Luzp2 |
| Ly6h | Rap1gds1 |
| Ppp1r16b | Gnao1 |
| Btbd1 | Map2k1 |
| Insyn1 | Dnajc6 |
| Dpy19l3 | Fam168a |
| Churc1 | Plp1 |
| Rps3a1 | Cstb |
| Nenf | Lrrk2 |
| Htr1b | Trrap |
| Gpr6 | Atn1 |
| Ryr3 | Golph3 |
| Unc80 | Hpcal4 |
| Sdhaf4 | Hdhd2 |
| Ntrk3 | Inpp4a |
| Smpd3 | Lars2 |
| Mmd2 | Prpf8 |
| Sfxn1 | Add3 |
| Mt2 | Meg3 |
| Kctd12 | Klhl2 |
| Shisa4 | Rraga |
| Asrgl1 | Fam131b |
| Kcnj10 | Azin1 |
| Erbin | Atp8b2 |
| Dnm2 | Ppp2r2c |
| Cmtm5 | Pptc7 |
| Etv1 | Erc2 |
| Eif4a1 | Cbx3 |
| Gja1 | Unc5a |
| Tmbim4 | Cox8a |
| Ppdpf | Ssr3 |
| Ttll11 | Cystm1 |
| Crocc | Wdr17 |
| Ap1ar | Hnrnpa0 |
| Mrpl10 | Phyhipl |
| Scn3b | Nrxn1 |
| Pon2 | Unc50 |
| Leprot | Yipf4 |
| Dock10 | Cinp |
| Hivep2 | Prdx3 |
| Eef2 | Lrp11 |
| Pts | Nr4a1 |
| Gpr158 | Nus1 |
| Gpr155 | Cacnb4 |
| Syndig1 | Kcna2 |
| Htra1 | Dcaf17 |

Robo2
Elovl5
Snhg1
Ift57
Srpk1
Ttc9b
Ezr
Sptbn1
Syngr1
Ddit3
B3gnt2
Gnptg
Snrpa
Tmem33
Calb1
Tmem229a
Cnp
Taf13
Dguok
Rnf5
Tbc1d9b
Abce1
B230219D22Rik
Itm2a
Gsn
Pigp
Lsm8
Msra
Tmco1
Btrc
Gabarap
Capzb
Cacna1h
Pllp
Ppp1r14a
Scn1b
Epn1
C1qtnf12
Psat1
Homer1
Mrpl17
Lyrm2
Isca1
Eps15
Naca
Enpp2
Atp6v1e1
Sowaha
Abcd3
Smim11
Rapgef2
Dmkn
Dync1i2
Klhl7
Zfr
Arpp19
Reep1
Prrc2b
Ldb1
Capza2
Tmed2
Slc1a3
Ctnnb1
B230312C02Rik
Hmgb1
Abat
B2m
Dhx15
Dnajb5
Lsm8
Tiprl
Hlf
Vgf
Cfap36
Ank
Abhd3
Rpl10-ps3
Hypk
Rabggtb
Trpc1
Pcdh17
Anapc16
Fam120a
U2af2
Ssb
Pde4b
Ube2d1
Cbr3
Cox16
Pdcd10
Pam
Rps15a
Ctsd
Ubb
Cd81
Rps14
Trf
Rpl18
Clstn1
Ndufb8
Rpl23
Fkbp8
Nefm
Rpl30

| | |
|---|---|
| Ncoa2 | Actg1 |
| Far1 | Psap |
| St8sia3 | Rpl6 |
| Snrpe | Stmn3 |
| Eml5 | Fis1 |
| Ppid | Hras |
| Stard3 | Rpl32 |
| Dusp6 | Ftl1 |
| Gnb2 | Apbb1 |
| Ccdc88c | Rps11 |
| Plp1 | Rpl9 |
| Drd2 | Hba-a1 |
| Sgip1 | Ndufb10 |
| Mapre2 | Ndufb9 |
| Ss18l2 | Rpl11 |
| Cbr3 | Rrp1 |
| Inpp5j | Fth1 |
| Unc50 | Rps5 |
| Ndfip2 | Ppp2r1a |
| Gltp | Rpl26 |
| Cdk5r2 | Ptgds |
| Tubb4a | Pkm |
| Myl12b | Rpl7 |
| Myo6 | Rps12 |
| Hexb | Atp5d |
| Dag1 | Smdt1 |
| Fnip1 | Pvalb |
| Zswim6 | Rpl5 |
| Epb41l2 | Rps15 |
| Igfbp7 | Atp9a |
| Cspg5 | Rpsa |
| Lrp11 | Rps3a1 |
| Vps29 | Pebp1 |
| Dync1h1 | Fkbp2 |
| Atf2 | Rps3 |
| Trappc4 | Rpl28 |
| Lmo7 | Rpl8 |
| Atp6v1c1 | Cnp |
| Ube2g1 | Ndufa13 |
| Zfp445 | Cck |
| Tpd52l1 | Gapdh |
| Atraid | Rpl24 |
| Phgdh | Rps10 |
| Rbm3 | Rpl18a |
| Max | Nenf |
| Dmac1 | Rps4x |
| Cnst | Cryab |
| Anks1b | Rps2 |
| Zfr | Rpl4 |
| Gadd45g | Rps7 |
| Sem1 | Rpl14 |
| Abhd17a | Rpl13 |

Pdia3
Scrg1
Inpp5a
Gcc2
Ndufv1
Capn2
Scamp3
Ipo7
Luzp2
Bhlhb9
Mark1
Usp33
Phactr1
Lcmt1
Sptb
Plpbp
Phldb1
Med4
Trf
Aldh6a1
Mphosph8
Syt6
Dhx15
Rfk
Wipf3
Mbd2
Kctd3
Prpf8
Csnk2a2
Rrp1
Fxyd1
Ubc
Cnot6
Jak1
Rab28
Gphn
Snapc5
Arpp21
Rnf166
Zfp385a
Isca2
Fez1
Mrps36
Ankrd10
Ramp1
Fmnl1
Hectd1
Mrps28
Pcdh17
Npas2
Prkch
Eml1
Rpl13a
Eno1
Fau
Rps27a
Pfdn5
Rps20
Rps8
Rpl10
Rpl21
Apod
Rps24
Hbb-bs
Ttr

Tubb2a
Rbis
Epb41l4aos
Kpnb1
Phlpp1
Maged1
Fnbp1l
Ina
Scg5
Hadhb
Rph3a
Jund
Nebl
Araf
Hs3st4
Dmxl2
Pfn1
Clk3
Park7
Gba2
Tppp3
Fhl1
Gng11
Olig1
Mn1
9330159F19Rik
Ctnna1
Zcchc24
Nexn
Lrrk2
Sec14l1
Tcf25
Rap1a
Eif4a3
Vbp1
Slc9a1
Ptprz1
Pmepa1
Pycr2
2010204K13Rik
Ccdc106
Phyh
Slc39a3
Grik2
Pnisr
Grm4
Tmod1
Wbp2
Gas2l1
Xbp1
Pitpnc1
3110039M20Rik

Acap2
Plppr5
Car11
Ssr3
Clvs1
Utrn
Setd7
Macf1
Snhg8
Agrn
Cenpx
Cacng4
Coa6
Api5
D5Ertd579e
Pdp1
Ssbp1
Selenop
Schip1
Higd2a
Lap3
Opalin
Efcab14
Gprc5b
Mapk1ip1
Ogt
Fads1
Sez6l2
Slirp
Chmp2b
Rnf13
Grn
Luc7l
Mrpl57
Lage3
Clip1
Brwd1
Marcks
Snrpc
Atp6v1g2
Tbc1d8
Neat1
Tln1
Ythdf1
Pcdh10
B2m
Ak3
Snapin
Pptc7
Mbtps1
Myh10
Srrt

Kif17
Dusp11
Vps52
Smarca5
Spata2l
Vps35
Ppp4r2
Sec23a
Ermn
Arhgap21
Tuba4a
Kdm4b
Stmn2
Ppp2r5a
2510009E07Rik
Lrrtm1
Tagln3
Xpo1
Apc
Prrc2a
Atp6v0e
Scd1
Atg12
Aggf1
Rapgef4
Tmem167
Isyna1
Ctss
Gramd1b
Psmg4
Ankrd13a
Abhd4
Rbp4
Nipa1
Rgs8
Usp3
Fxyd7
Pdk3
Arhgef2
Letmd1
Xpo7
Tmem47
Gtf3c2
Map7
Mcf2l
Ccnh
Got1
Tmem183a
AI593442
Mpv17l
Vgf
Ric8b

Cldn5
Sh3rf2
Myo1b
Serinc5
Snhg20
Ifrd1
Hax1
Id4
Cdk2ap2
Yipf3
Mpped2
Prkacb
Spry2
Unc5a
Tsc22d3
Rabggta
Tyrobp
Smim26
Zfas1
Pbx3
Gng12
Tmem170b
Arhgdib
Fam216a
Dtna
Ryr1
Kdm3a
Ciao2b
Cd9
Sesn1
Foxk2
Add3
Nsd2
Ralgapa1
Cobl
Fcho1
Acy1
Pttg1ip
Dab2ip
Pacsin1
Tra2a
Dusp3
Fyn
Paqr8
Tnk2
Rian
Dip2c
Mxd4
Chka
Gdap1
Dld
Rbp1

Mrps11
Il33
Dpf2
Foxo1
Ccdc85c
Six3
Mrpl40
Uqcrc1
Polr2f
Degs1
Mzt1
Cfap36
Aldh5a1
Polr2i
Vat1l
Tmem208
S1pr1
Ankrd13c
Nme2
Slc35a4
Cope
Ldah
Surf1
Ephx1
Phf24
Apba1
Pou3f3
Plpp6
Caprin1
Vps9d1
Gaa
Med27
Tafa5
Neto1
Nat8f1
Fnta
Golga4
Cdk19
Sh3kbp1
Azi2
Ube2e3
Rcn1
Acvr1c
Mcee
Smim20
Dnajb4
Fgf13
Ube2m
Mrpl44
Smad3
Rtf2
Ralgds

Srsf6
Ccnl2
Epb41l3
Ppp6r1
Nomo1
P4hb
Fut8
Csnk1d
Fam120b
Ddx50
U2af2
Hsd17b12
Grk6
Cstf2t
Pdxp
Dpy30
Limch1
Tram1l1
Srr
Agtpbp1
Polr2j
Stip1
Eif2s1
Faim2
Nrep
Sh3gl3
Immp1l
Hmgcl
Ccm2
Rubcn
Gnb5
Taf1b
Dpm3
Chrm1
Kansl2
Mrpl13
Fnbp1
Fbxl3
Mettl23
Fabp3
Chgb
Ddx1
Emc3
Ahcyl2
Srsf7
Rabl6
Arl8a
Gpsm1
Gm10419
Arhgap23
Ppp4r4
Zfp871

Fis1
Slitrk5
Rnf181
Lsm7
Sccpdh
Bcr
Lrrc10b
Kazn
Dhrs7
Sec11a
Armc1
Polr2k
Ubqln1
Actn2
Relch
Ktn1
Dnaja4
Manf
Fam174a
Adgrl2
Sumo2
Rgs2
Saraf
Plxna2
Sh3gl2
Tbc1d16
Trappc13
Dclk2
Fcer1g
Smg1
Hspd1
Zmym2
Wscd1
Tardbp
Pcmtd2
Dixdc1
Slc35f3
Mctp1
Stk32a
Sbds
Ube2i
Sf3b5
Asb13
Pmm1
C2cd2l
Rhot1
Cdk16
Asah1
Galnt13
Mink1
Cdo1
Junb

Abcc5
Rab10
Acyp1
Ap1s2
Rprd1a
Nme1
Nxf1
Bud23
Rmdn3
Sppl3
Chn2
Snx2
Ccdc47
Ttc9
Sv2a
Fcor
Mtus1
Nin
Trim9
Osbpl2
Atp8a1
Polr2g
Cox20
Dnajc2
Rab40c
Pltp
Dmtf1
Mapkap1
Maz
Mdh2
Anapc16
Cyb5a
Papss1
Tomm34
Phf20l1
9130401M01Rik
Adcy1
Slc35c2
Vma21
Stub1
Akap8
Pdcd10
Kbtbd2
Fth1
Dctpp1
Crat
Aph1a
Prkaca
1110038F14Rik
Maneal
Snrnp27
Coro7

Tomm5
Lrrc4b
Shisa9
Gnpat
Ndufaf5
Txndc16
Commd6
Tmem134
Ptpn4
Uba5
Rpp21
Scarb2
Lpl
Wbp1
Rps6ka5
Sri
Rbfa
Cacng3
Ino80e
Jak2
Hnrnpc
Dpm2
Sipa1l3
Ppp2r2d
Lamtor3
Slc35d3
Sel1l
Pdzd8
Timm10
Scn3a
Tmem201
Spop
Ripor2
Sdr39u1
Akap5
Myl12a
Aopep
Vta1
Cystm1
Ndrg1
Yme1l1
Mtmr9
Stk19
Sod1
Slc1a1
Srgap1
Naa20
Dmwd
Phf5a
Dop1b
Skil
Timm9

Cstf3
Bcl11a
Shb
Cd4
Abl2
Emc6
Large1
Stau1
Akt3
Son
Gmpr
Prr7
Wwc1
Secisbp2l
Iah1
Pantr1
Dctn5
Ap3b1
Dut
Cdc34
Ubr2
Cacybp
Casc3
Rala
Cd99l2
Bap1
Hikeshi
Ilk
Arih1
Pde4b
Rhobtb2
Dctn4
Mrpl32
Usp9x
Osbpl1a
Gpcpd1
Ewsr1
Orc4
Esd
Mrpl16
Hspa4l
Senp2
Jtb
Ankrd37
Chtop
Dagla
Morf4l1
Chd9
Rgs5
Cnn3
Gcsh
Jkamp

Ankrd40
Chmp4b
Klhl7
Cox16
Tmem185a
Rmnd5b
Plekhb1
Gga3
Lgi1
Sfpq
Slc49a4
Cstf2
Pxdn
Plcxd1
Fzd3
Mrpl3
Pip5k1a
Ube2a
Tpgs2
Prelid3b
Maea
Apbb3
Grsf1
Hnrnph3
Uba3
Stam
Bud31
Dda1
Arel1
Mien1
Slk
Enah
Gabrb3
Pcna
Timp3
Nedd4l
Pdia6
Fbxw11
Bptf
C1qb
Dtymk
Carm1
Lrtm2
Cacnb1
Ube2f
Otulinl
Kcnk1
Rnf6
Smpdl3a
Cdh13
Dcaf7
Snd1

Sipa1l1
Scg3
Plekhn1
Fabp5
Itgb1bp1
Serp1
Clta
Serinc3
Tor1aip2
Ank3
Ankmy2
Rad23a
Cacna1c
Atp1b2
Kmt2e
Asic4
Gpatch8
Appl2
Rc3h2
Tbc1d20
Ankrd46
Ncbp1
Irf2bpl
Ubald2
Clk1
Il18
Ncor2
Trp53bp1
Hsd17b10
St6galnac5
Tex264
Fgfr1op2
Atxn10
Iffo1
Hbs1l
Fubp1
Chst15
Ppib
Emc2
Spred1
Dnal4
Gcnt2
B3galt5
Vps53
Sh3bgrl
Jph4
Slc44a1
Sae1
Ap3m1
Gsg1l
Acaa2
Psmd13

Snap47
Etv5
Slc25a44
Rai1
Gpd1
Zdhhc18
Ado
Mbnl1
Slc25a46
Dst
Ranbp3
1110059G10Rik
Ctdsp2
Ipo11
Tm2d2
Tprkb
Ccng2
Ppa2
Pip4p1
Abhd16a
Usp32
Srm
Krt10
Syn2
Tax1bp1
Abhd17c
Gm3764
Ppp1r12a
Ccdc32
Snx1
Timm29
Rell2
Ubp1
Zfp180
Necab1
Cnot7
Rnf114
Tmub2
Mat2a
Cnih1
Fam8a1
Rab5if
Micu1
Rbm7
Epn2
Gm33651
C1qc
Hmbs
Taf11
Rdh14
Slc25a25
Plec

Arhgap5
Srsf1
Pnrc1
Stau2
Slc30a9
Mpp1
Mocs2
Rnf4
Sybu
Atg4b
Ncoa1
Dnajb9
Rnf34
Ran
Sorcs2
Chchd3
Slc35b4
Ddit4l
Dhx36
Ppp5c
Krtcap2
Malsu1
Arfrp1
Rbm18
Nktr
Lrpprc
Naxd
Nsf
Cdc42ep3
Wdr26
Cisd2
S100a13
Ap3s1
Hnrnpll
Dhx32
Dync1i2
Luc7l2
Sst
Nipsnap1
Ank
Ppp6c
Ubr3
Ndufaf3
Mrpl41
Nt5c3
Pfkm
B4galnt1
Becn1
Trim2
Prickle2
Sun2
Cdadc1

Thyn1
Leprotl1
Afdn
Cnr1
Kmt5a
Ptgr2
Prpf4b
Hadha
Cdc42bpa
Psd3
Lypd1
Tmem131
Dnajb2
Msmo1
Gm2a
Pik3c3
Lrrtm2
Nsd1
Rbm33
Endod1
Ccdc28b
Mageh1
Zeb1
Alkbh7
Foxp2
Lman2
M6pr
Cbx4
Msrb1
Ulk2
Pten
Cnih4
Zdhhc2
Med21
Supt6
Rin1
Zfp637
Scrt1
Fibp
Ubr5
Adk
Thsd7a
Adcyap1r1
Mrps12
Elp5
Opa1
Mtdh
Kif3c
Cnot2
Ubac1
Ndst1
Impdh1

Rgs10
Ctnna2
Trpc4ap
Syt16
Fkbp4
Ythdf3
Pigs
Xiap
Tmeff1
Yipf5
Polr2l
Plxnd1
Pgs1
Usf2
Pomgnt1
Zranb2
Leng8
Sugt1
Sh3glb1
Camk4
Sirt7
Tsr3
Gpam
Mpp3
Eef1g
Psmc6
Stt3b
Mrpl27
Gskip
Hpf1
Gna12
Desi1
Plppr1
Ube2h
Ubl4a
Arsb
Zhx1
Uso1
Gtpbp6
Pclo
Trap1
Efna3
Cacnb2
Ncbp2
Ralbp1
Bri3
Exoc5
Atxn1
Tia1
Ica1
Irak1bp1
Akt1

Tcf4
Ubr4
Cdc123
Mlec
Ubxn2b
Otud6b
U2af1l4
Ndel1
Bckdha
Ggt7
Kbtbd11
Npy
Tomm40
Med9
Abracl
Zfp638
Gosr2
Tmbim6
Eif1a
Gpr27
Ilkap
Trim11
Erg28
B4galt3
Pi4k2a
Tug1
Babam2
Tmem68
Elp1
Slmap
Dipk1b
Sacm1l
Klf13
Tmem14a
Usp7
Atp6v0a2
Picalm
Med25
Tmem126a
Pum1
Rnf20
Lrrc8b
Snhg12
Ube3c
Eif3e
Eef1akmt1
Sap18
Strn
Fgf12
Adgrb3
Ddx42
Mal2

Basp1
Acvrl1
Gpbp1
Tgfa
Lyrm4
Rnf7
Ckmt1
Jagn1
Dctn2
Rab26
Man1a2
Tmx3
Wasl
Arl6
Igfbp4
Evi5l
Tceal8
Mapk9
Scrt2
Eif4g1
Seh1l
Fbxo11
Fundc1
Gap43
Snap91
Shroom2
Cttnbp2
Ylpm1
BC004004
Cfl2
Oaz2
Elof1
Vps11
Tmem181a
Top1
Manbal
Cacna1e
Kcns2
Acbd5
Kctd1
Syne1
Anapc5
Clns1a
Magi2
Chst11
Ddah1
Abhd14a
Tmed4
Nfx1
Mrpl50
Nicn1
Dgke

Magoh
Rab11fip3
Mtor
Tmem184b
Etf1
Txnl4a
Cox19
B9d2
Akt1s1
Aup1
Ehbp1l1
B4galt2
Zfp580
Emg1
Adam15
Trpc1
Glcci1
Mdp1
Mrps7
Nat8l
Sult4a1
Aldh7a1
Sdc2
Blcap
Nt5c2
Srsf10
Slc25a1
Anapc2
Slc4a4
Hexa
Nudt4
Got2
Mrpl49
Scmh1
Pdzd11
Cipc
Lsm1
Gtf2a2
Bzw1
Ids
Tmem263
Snw1
Sec61a1
Zbtb8os
Mia3
Sgcb
Med10
Bphl
Snx27
Adra2c
Dhx30
Ptpa

Morn4
Pde8b
Vezt
Rnpep
Zmynd8
Coa3
Vtn
Rgs7
Plekhm2
Ckap5
Ebna1bp2
Kif5c
Traf7
Naa50
Gclm
Trappc11
Plekha3
Ddit4
Myo9a
Adcy3
Rad21
Snx5
Ankrd17
Slc9a5
Kat7
Ufd1
Pla2g12a
Caml
Chst1
Mark2
Pnn
Pacsin2
Mrpl46
Psma5
Psen1
Amer2
Trim3
Ube2j1
Arhgef12
Coq10b
Copa
Ube3a
Tango2
Slc38a3
Ncoa7
2300009A05Rik
Tmem59l
Osgep
Ptpn11
Pkp4
Peli1
Ipo13

Flot1
Ajap1
Ttc14
Tmem205
Plxnb1
Agap1
Cop1
Abcb9
Hdhd2
Limk2
Fam91a1
Cyfip1
Med14
Rbbp4
Prkca
Srgap2
Sf3b3
Mib2
Lman2l
Rit1
Dalrd3
Celf3
Exosc7
Nmt2
Trip12
Ski
Stx7
Ufm1
Pdk2
Arid1a
Mfap1b
Commd1
Hsd17b4
Tpst2
B3galt1
Mxi1
Rab3ip
Cdk5r1
Nrbp1
Ppip5k1
Yaf2
Ppil4
Mfsd13a
Cuedc2
Ecpas
Lsm14a
Rsrc2
Nfasc
Znhit3
Trappc12
Eif3c
Prkcsh

Sf3b4
Wdr45b
Trmt1
Fam107a
Nipa2
Tppp
Upf1
0610010K14Rik
Arf2
Nr2c2
Syt3
Ruvbl1
Pls3
Metap2
Pigx
Aldh1l1
Calcoco1
Kcna4
Morn2
Mob4
Fam117b
Erh
Zdhhc3
Pcbp3
Ache
Ube2q1
Mrpl36
Rbm28
Mrpl4
Ganab
Map3k10
Dapk3
Nipsnap2
Agpat5
Slc6a8
Mydgf
Tmem70
Parp1
R3hdm1
Gbf1
Ifi27
Naa10
Coq9
Oxa1l
Gfra4
Gdpd5
Ap1g1
Mrpl14
Aff3
Cyth2
Fech
Rpl7l1

Med19
Ociad2
Hdgf
Sdhaf1
Hmgcr
Mrtfa
Fbll1
Otud5
Arfgef1
Foxo3
Ralgapb
B230217C12Rik
Ogfr

Lat Septal Complex iTBS

| DEGs Young sham vs Aged sham | iTBS DEGs unique to young adults | iTBS DEGs unique to young adults that are also ageing DEGs |
|---|---|---|
| Cpe | Gstp1 | Bc1 |
| Uba52 | Ppp1r1b | Cartpt |
| Mt1 | Camk2n1 | Gpm6b |
| Rpl4 | Cck | Lrrc17 |
| Slc1a2 | Eno1 | Pcp4 |
| Rps24 | Pde1b | Penk |
| Mt2 | Pcp4 | Rpl37 |
| Bc1 | Penk | |
| Ndufa4 | Gpm6b | |
| Rpl5 | Itpr1 | |
| Thrsp | Fbxl16 | |
| Rpl37 | Canx | |
| Rpl14 | Rgs4 | |
| Gapdh | Arpp21 | |
| Cox8a | Ndn | |
| Fau | Syt5 | |
| Rps27rt | Rpl37 | |
| Rpl21 | Lrrc17 | |
| Malat1 | Tatdn1 | |
| Ppia | Gm2000 | |
| Ddn | Nts | |
| Fam107a | Cartpt | |
| Cnih2 | Bc1 | |
| Nme7 | | |
| Rps7 | | |
| Rps10 | | |
| Pcp4 | | |
| Rps29 | | |
| Rpl9-ps6 | | |
| Rps20 | | |
| Camk2a | | |
| Ezr | | |
| Dbi | | |
| Bok | | |
| Rps27a | | |
| Dbn1 | | |
| Zcchc24 | | |
| Fads1 | | |
| Rpl9 | | |
| Agap2 | | |
| Epas1 | | |
| Calm2 | | |
| Mlc1 | | |
| Whrn | | |
| Myo6 | | |
| Pfdn5 | | |
| Synm | | |
| Pfn1 | | |
| Usp50 | | |

Slc1a3
Rps28
Rps8
Paqr8
Zic1
Stmn1
Neurl1a
Lrrc17
Rpl35
Itm2c
Igfbpl1
Gng4
Chst2
Kcnj10
Atp1a1
Penk
Rpsa
Kctd12
Rpl35a
Lats2
Doc2g
Ctnnb1
Scn1b
Cacna1g
Cacng5
Pde2a
Necab2
Klhl5
Rpl3
Dlg2
Lars2
Gap43
Rbm3
Hspa14
Rps4x
Tprkb
Rpl24
Tspyl4
Rps2
Timp3
Nbl1
Rps11
Ppm1e
Ndrg2
Aldh5a1
Sptbn1
Aldh7a1
Cartpt
Gfap
Shank1
Slc8a1
Ldah

Snhg6
Arhgef12
Dpysl2
Ewsr1
Fhl1
Dag1
Prkcd
Rps3
Stac2
Slc39a12
Lrrfip1
Ndufa7
Slco1c1
Prdx1
Virma
Pitpnm3
Trim8
Atp1a2
Atp2a2
Kansl2
Ivns1abp
Psph
Tcf4
Mvb12b
Timm22
Ifngr2
Kif5c
Hs3st2
Tnfrsf21
Maneal
Rims2
Nfia
Cdk2ap2
Tubb5
Chd3
Map4k3
Slc6a1
Il6st
Pygb
Nenf
Mink1
Pea15a
Vim
Setd7
Atxn1
Tbc1d9b
Plekha1
Eef1a2
Stmn4
Ube2ql1
Ost4
Lsm2

Chordc1
Chd9
Tmem47
Clu
Prmt5
Klc1
Jund
Nfib
Sulf2
Twf2
Zic4
Clic4
Gpr137
Trpm3
Lix1
Map9
Mir100hg
Sh3bgrl
Elfn2
Rph3a
Jak1
Epb41l2
Zfr
Rgs10
Plcg1
Enah
Nfkbia
Ppp1r9b
Ly6a
Arhgap1
Txndc16
Tssc4
Zfp521
Parva
Gpm6b
Acss1
Gpam
Sacm1l
Hdac6
Gldc
Map4
Rab31
Gpd2
Sox1ot
Ythdf1
Cnot6l
Zfr2
Ids
Brinp2
Nwd1
Map7
Epn2

Btbd1
Zdhhc14
Abhd17b
Araf
Apc
Api5
Mmd2
Gnb2
Ahcyl2
Fbxo41
Cep120
Ap1s2
Tpd52l1
Rapgefl1
Scrt1
Foxk2
Shd
Exoc1
Trappc5
Eif4a1
Epha4
Pdzd2
Adk
Pacsin1
Mat2a
Fam120a
Ywhah
Sp9
Dok6
Iqsec3
Megf9
Atg4b
Prxl2b
Sgip1
Gadd45g
Crat
Zeb2
Sptan1
Kcng1
Luc7l2
Cdr1os
Ormdl3
Ak3
Abcb7
Rc3h2
Fam210b
Nif3l1
Prdm16
D430019H16Rik
Acyp2
Srgap3
Adarb1

Homer2
Nubp2
Plppr4
Glul
Ddx17
Mark4
Pmpca
Vma21
Klhl24
Eps15
Syt1
Bud23
Serp1
Nek6
Prelp
Clmn
Immt
Slc6a11
Tfdp1
Arhgef2
Osbpl9
Ttyh1
Zfp423
Nf2
Dlx6os1
Dvl1
Calb1
Mrtfa
Cnn3
Ptpn4
Camsap1
Idh2

Lat Septal Complex cTBS

| DEGs Young sham vs Aged sham | cTBS DEGs unique to young adults | cTBS DEGs unique to young adults that are also ageing DEGs |
|---|---|---|
| Cpe | Tatdn1 | Acyp2 |
| Uba52 | Araf | Adk |
| Mt1 | Tspyl4 | Agap2 |
| Rpl4 | Actr2 | Ahcyl2 |
| Slc1a2 | Agap2 | Aldh5a1 |
| Rps24 | Slc6a1 | Apc |
| Mt2 | Prkcd | Araf |
| Bc1 | Basp1 | Arhgef12 |
| Ndufa4 | Snap25 | Atxn1 |
| Rpl5 | Rasd1 | Cacna1g |
| Thrsp | Glul | Camk2a |
| Rpl37 | Snhg9 | Cdr1os |
| Rpl14 | Ddn | Chd3 |
| Gapdh | Tomm7 | Chd9 |
| Cox8a | Isca1 | Chst2 |
| Fau | Ndufa1 | Clic4 |
| Rps27rt | Camk2a | Cpe |
| Rpl21 | Atp1a3 | Ctnnb1 |
| Malat1 | Snca | D430019H16Rik |
| Ppia | Hopx | Ddn |
| Ddn | Car2 | Dlg2 |
| Fam107a | Ost4 | Elfn2 |
| Cnih2 | Snrpf | Eps15 |
| Nme7 | Pacsin1 | Ewsr1 |
| Rps7 | Ppp1r9b | Fads1 |
| Rps10 | App | Fam120a |
| Pcp4 | Phpt1 | Fau |
| Rps29 | Ptprz1 | Gapdh |
| Rpl9-ps6 | Ank2 | Glul |
| Rps20 | Bloc1s1 | Gpm6b |
| Camk2a | Snhg8 | Ids |
| Ezr | Sem1 | Kctd12 |
| Dbi | Mbp | Map4 |
| Bok | Ndrg4 | Megf9 |
| Rps27a | Naa20 | Mt2 |
| Dbn1 | Eif4a2 | Nenf |
| Zcchc24 | Sdhd | Ost4 |
| Fads1 | Plppr4 | Pacsin1 |
| Rpl9 | Slc4a4 | Paqr8 |
| Agap2 | Atp6v1a | Pfdn5 |
| Epas1 | Gja1 | Plppr4 |
| Calm2 | Zwint | Pmpca |
| Mlc1 | Wsb2 | Ppp1r9b |
| Whrn | Atn1 | Prkcd |
| Myo6 | Rnf5 | Rpl14 |
| Pfdn5 | Snhg6 | Rpl21 |
| Synm | Slc25a22 | Rpl24 |
| Pfn1 | Cdk5r2 | Rpl3 |
| Usp50 | Spag9 | Rpl35a |

Slc1a3
Rps28
Rps8
Paqr8
Zic1
Stmn1
Neurl1a
Lrrc17
Rpl35
Itm2c
Igfbpl1
Gng4
Chst2
Kcnj10
Atp1a1
Penk
Rpsa
Kctd12
Rpl35a
Lats2
Doc2g
Ctnnb1
Scn1b
Cacna1g
Cacng5
Pde2a
Necab2
Klhl5
Rpl3
Dlg2
Lars2
Gap43
Rbm3
Hspa14
Rps4x
Tprkb
Rpl24
Tspyl4
Rps2
Timp3
Nbl1
Rps11
Ppm1e
Ndrg2
Aldh5a1
Sptbn1
Aldh7a1
Cartpt
Gfap
Shank1
Slc8a1
Ldah

Luzp2
Rps27rt
Arpp21
Chst2
Rpl9-ps6
Prrc2a
Slc1a2
F3
Ppp1r1b
Cox7c
Map2
Rab3c
Mt2
Tmed7
Kctd13
Gpm6b
Acyp2
Plp1
Tsc22d4
Sipa1l1
Ctnnb1
Rcn2
Prpf8
Usp50
Atp5md
Mgat3
Gnai1
Phactr1
1810037I17Rik
Slc4a10
Psip1
Ubqln2
Huwe1
Ids
Fam131b
Fbxl16
Cpe
Megf9
Ahcyl2
Fus
Slc1a3
Cs
Id4
Dmtn
Jph4
Sparcl1
Man1a
Cops7a
Prrc2b
Ppp1r16b
Hpcal4
Phyhip

Rpl4
Rpl5
Rpl9
Rpl9-ps6
Rps10
Rps11
Rps2
Rps20
Rps24
Rps27a
Rps27rt
Rps3
Rps4x
Rps7
Rps8
Rpsa
Setd7
Sgip1
Shank1
Slc1a2
Slc1a3
Slc39a12
Slc6a1
Snhg6
Sptan1
Srgap3
Syt1
Tbc1d9b
Tmem47
Trim8
Tspyl4
Ttyh1
Usp50
Zcchc24

| | |
|---|---|
| Snhg6 | Txn1 |
| Arhgef12 | Rpl39 |
| Dpysl2 | Rusc2 |
| Ewsr1 | Camk2n1 |
| Fhl1 | Atp6v1h |
| Dag1 | Kctd12 |
| Prkcd | Peg10 |
| Rps3 | Ap1ar |
| Stac2 | Scd1 |
| Slc39a12 | Lmtk3 |
| Lrrfip1 | Stxbp1 |
| Ndufa7 | Klf13 |
| Slco1c1 | Tmbim6 |
| Prdx1 | Ndfip2 |
| Virma | Dnajb5 |
| Pitpnm3 | Ndufa3 |
| Trim8 | Rap1gds1 |
| Atp1a2 | B2m |
| Atp2a2 | Brms1l |
| Kansl2 | Dock4 |
| Ivns1abp | Ncan |
| Psph | Cfap20 |
| Tcf4 | B230312C02Rik |
| Mvb12b | Gad1 |
| Timm22 | Tbc1d9b |
| Ifngr2 | Gnao1 |
| Kif5c | Rab28 |
| Hs3st2 | Spop |
| Tnfrsf21 | Kif3c |
| Maneal | Crk |
| Rims2 | Rheb |
| Nfia | Peg3 |
| Cdk2ap2 | Selenof |
| Tubb5 | Eps15 |
| Chd3 | Cadm4 |
| Map4k3 | Cd9 |
| Slc6a1 | Shank1 |
| Il6st | Khsrp |
| Pygb | Arpc4 |
| Nenf | Paip2 |
| Mink1 | Plekhb1 |
| Pea15a | Ank |
| Vim | Prkar1a |
| Setd7 | Rerg |
| Atxn1 | Pde10a |
| Tbc1d9b | Add3 |
| Plekha1 | Senp3 |
| Eef1a2 | Stum |
| Stmn4 | Abhd3 |
| Ube2ql1 | Hectd4 |
| Ost4 | Cds2 |
| Lsm2 | Atp5g3 |

| | |
|---|---|
| Chordc1 | Cdk17 |
| Chd9 | Fam168a |
| Tmem47 | Qk |
| Clu | Apc |
| Prmt5 | Wdr89 |
| Klc1 | Hprt |
| Jund | Pou3f3 |
| Nfib | Cd47 |
| Sulf2 | Elfn2 |
| Twf2 | Sptan1 |
| Zic4 | Clic4 |
| Clic4 | Bzw1 |
| Gpr137 | Fam219a |
| Trpm3 | Necap1 |
| Lix1 | Fads1 |
| Map9 | Usp19 |
| Mir100hg | Fam102a |
| Sh3bgrl | Smim4 |
| Elfn2 | Kif5a |
| Rph3a | Cbx3 |
| Jak1 | Dlx1 |
| Epb41l2 | Smim26 |
| Zfr | Sft2d1 |
| Rgs10 | Tuba1a |
| Plcg1 | Syngap1 |
| Enah | Ccdc85c |
| Nfkbia | Fam171b |
| Ppp1r9b | Arhgef12 |
| Ly6a | Tmem47 |
| Arhgap1 | Chd9 |
| Txndc16 | Csrnp3 |
| Tssc4 | Hnrnpa2b1 |
| Zfp521 | Rapgef4 |
| Parva | Ewsr1 |
| Gpm6b | Ttyh3 |
| Acss1 | Gsk3b |
| Gpam | Setd7 |
| Sacm1l | Tm9sf2 |
| Hdac6 | Arhgap39 |
| Gldc | Atp1b2 |
| Map4 | Slc39a12 |
| Rab31 | Rpl10-ps3 |
| Gpd2 | Gria2 |
| Sox1ot | Atxn2 |
| Ythdf1 | Psma5 |
| Cnot6l | Yipf4 |
| Zfr2 | Cntnap1 |
| Ids | Hnrnpa3 |
| Brinp2 | Nol4l |
| Nwd1 | Atxn2l |
| Map7 | Dnajc6 |
| Epn2 | Dnajb4 |

| | |
|---|---|
| Btbd1 | Chmp5 |
| Zdhhc14 | Acsl3 |
| Abhd17b | Ttc7b |
| Araf | Usp9x |
| Apc | Anks1b |
| Api5 | Gng7 |
| Mmd2 | Mrps28 |
| Gnb2 | Ripor2 |
| Ahcyl2 | Trrap |
| Fbxo41 | Ssr3 |
| Cep120 | Kif1b |
| Ap1s2 | Map4 |
| Tpd52l1 | Cyth1 |
| Rapgefl1 | Tsc22d3 |
| Scrt1 | Klf9 |
| Foxk2 | Zcchc24 |
| Shd | Ccni |
| Exoc1 | U2af2 |
| Trappc5 | Mtmr9 |
| Eif4a1 | Abat |
| Epha4 | Aldoc |
| Pdzd2 | Sgip1 |
| Adk | Lamp1 |
| Pacsin1 | Aldh5a1 |
| Mat2a | Trim8 |
| Fam120a | Dact3 |
| Ywhah | Mapre2 |
| Sp9 | Larp1 |
| Dok6 | Tmem258 |
| Iqsec3 | Syt1 |
| Megf9 | Chd3 |
| Atg4b | Dlg2 |
| Prxl2b | Rraga |
| Sgip1 | Gucy1b1 |
| Gadd45g | Dhx15 |
| Crat | Srsf5 |
| Zeb2 | Cdr1os |
| Sptan1 | Comt |
| Kcng1 | Ogdhl |
| Luc7l2 | Acat1 |
| Cdr1os | Morf4l1 |
| Ormdl3 | Fbrsl1 |
| Ak3 | Bmpr2 |
| Abcb7 | Msi2 |
| Rc3h2 | Hectd2 |
| Fam210b | Ttyh1 |
| Nif3l1 | Nova2 |
| Prdm16 | Hacd2 |
| D430019H16Rik | Arel1 |
| Acyp2 | Nus1 |
| Srgap3 | Tnk2 |
| Adarb1 | Serinc3 |

Homer2
Nubp2
Plppr4
Glul
Ddx17
Mark4
Pmpca
Vma21
Klhl24
Eps15
Syt1
Bud23
Serp1
Nek6
Prelp
Clmn
Immt
Slc6a11
Tfdp1
Arhgef2
Osbpl9
Ttyh1
Zfp423
Nf2
Dlx6os1
Dvl1
Calb1
Mrtfa
Cnn3
Ptpn4
Camsap1
Idh2

Max
Mapk1
Pde4b
Frmpd3
Sbds
Pdpk1
B230334C09Rik
Umad1
Fgfr1op2
Fam120a
Shank3
Atp5mpl
9330159F19Rik
Rragd
Mrpl10
Abhd8
Lzts1
Rasd2
Pde4dip
Gabra4
Bhlhe40
Adk
Prkcb
Cacna1g
Srgap3
Lin52
Slc8a2
Cdc42se2
Rpl37a
Mpv17l
Pmpca
Dlgap1
Srrm1
Hnrnpu
Nutf2-ps1
Rnf141
Lpcat4
D430019H16Rik
Smim10l1
Xiap
Zcchc14
Gm20594
Unc80
Lsm8
Unc50
Paqr8
Spry2
Eno1b
Atxn1
Hibadh
Egln1
Rgs7bp

Rap1a
Dazap2
Miga1
Thra
Tiprl
Pmm1
Rps17
Ndufb8
Rpl3
Rpl27a
Psmc5
Rpl23a
Rpl7
Gpx4
Rps14
Ndufb11
Rpl29
Fdx2
Uqcrh
Timm8b
Hint1
Rpl28
Rpl23
Naca
Rpl36al
Rpsa
Ndn
Rplp1
Rps19
Eif3k
Nme2
Ubb
Edf1
Ftl1
Rpl5
Rpl35a
Rpl8
Ndufb9
Rps15
Rpl30
Eno1
Dynll1
Rps3a1
Rpl19
Rpl17
Rpl4
Rpl11
Pebp1
Rps4x
Rps16
Rpl32
Rps5

Rps2
Rpl18a
Gapdh
Rpl26
Rpl6
Rpl13
Rps15a
Rpl9
Rps3
Rpl18
Rps12
Pfdn5
Rpl24
Nenf
Rps11
Rps10
Rps20
Rpl13a
Ndufa13
Rps7
Rps27a
Fau
Rpl21
Rpl14
Hbb-bs
Rps8
Rpl10
Rps24

Striatum iTBS

| DEGs Young sham vs Aged sham | iTBS DEGs unique to young adults | iTBS DEGs unique to young adults that are also ageing DEGs |
|---|---|---|
| Bc1 | Eno1 | Lrrc17 |
| Cox8a | Pcp4 | Mbp |
| Actb | Hmgn2 | Pcp4 |
| Rps24 | Mbp | |
| Malat1 | Nrgn | |
| Rpl4 | Agap2 | |
| Sst | Snhg9 | |
| Fau | Lrrc17 | |
| Dpysl2 | | |
| Pcp4 | | |
| Mt1 | | |
| Rpl3 | | |
| Rpl5 | | |
| Bloc1s1 | | |
| Rasgrf2 | | |
| Snca | | |
| Rpl37 | | |
| Rpl13 | | |
| Eef1a1 | | |
| Ly6c1 | | |
| Usp50 | | |
| Nts | | |
| Gamt | | |
| Otof | | |
| Rpl14 | | |
| Rps10 | | |
| Ntsr2 | | |
| Hspa5 | | |
| Snhg6 | | |
| Ddn | | |
| Lingo3 | | |
| Cpe | | |
| Itm2c | | |
| Rps8 | | |
| Ptgds | | |
| Uba52 | | |
| Cacna1h | | |
| Calm1 | | |
| Tsc22d1 | | |
| Them6 | | |
| Atp1b1 | | |
| Polr2l | | |
| Ndrg4 | | |
| Cox7b | | |
| Rpl13a | | |
| Rps2 | | |
| Psd | | |
| Rpl9 | | |
| Polb | | |

Kcnip2
Strip2
Snhg11
Rps7
Josd2
Ankrd13b
Ndufa4
Nefm
Unc13c
Tac1
Pfdn5
Polr2j
Mbp
Snhg20
Necab1
Cck
Pacsin2
Arhgap33
Maneal
Mal
Higd2a
Coro7
Ccm2
Zdhhc14
Trak2
Dalrd3
Rps3a1
Jsrp1
Arpp21
Lrpap1
Camk1g
Snhg1
Srsf11
Mrpl41
Pitpnm2
Eef1a2
Gucy1a1
Sdhaf2
Chordc1
Rac3
Acvr2a
Rpl9-ps6
Rnf20
Srp72
Smpd3
Rps14
Yipf1
Morf4l1
Acsbg1
Rps27a
Mapk1ip1
Ppid

Oga
Sirt2
Mat2a
Pfn1
Btbd3
Agrn
Elp2
Ralgds
Sirt3
Scn4b
Limd2
Mrpl21
Slc2a1
Emc2
Lyst
Ly6a
Rps4x
Epas1
Taf13
Tmem230
Ost4
Kansl2
Amz2
Filip1
Arel1
Slc35d3
Rgl2
Acyp2
Tmub2
Rpl30
Nedd4l
Tubb5
Tbc1d9b
Pttg1ip
Prxl2a
Brinp1
Nlgn1
Trmt1
Cbx4
Inf2
Rasd2
Pvalb
Ccdc28b
Timm9
Grm4
Ubap2l
Adcy5
Stmn1
Trmt2a
Snhg3
Myo5b
Mydgf

Phldb1
Ahcyl2
Arhgef2
Tmem126a
Sptbn4
Kif21b
Cdo1
Adam15
Ctxn1
Ndufa7
Hip1r
Mrps12
Panx2
Zmat5
Ptpro
Cinp
Rps6ka3
Ypel4
Mirg
Txndc16
Mir124a-1hg
Rps20
Adora2a
Cspg5
Fam210b
Ywhah
Rps5
Ephx1
Cyp51
Mcee
Pds5b
Ppp5c
Mog
Cdk4
Chn2
Tmem208
Carm1
Degs1
Fxyd2
Creg1
Fyco1
Large1
Cox19
Slc17a7
Ddit3
Hspe1
Pop7
Kcnk1
Luc7l
Dlx6os1
Tmem167
Fhl1

Rasgrp2
Mcf2l
Adarb1
Senp2
Aldh7a1
R3hdm4
Rpl24
Slco1c1
Dpf2
Tmod1
Ythdf1
Pdia6
Eml5
Ldah
Scg2
Zmym3
Polr3e
Tspan2
Kcnd3
Mobp
Dmac1
Sirt7
Plekhj1
Coq2
Cldn5
Ilkap
Nt5c3b
Mdk
Gadd45g
Chst2
Wfs1
Mmp17
Il18
Ube2f
Rell2
Dnajb11
Gap43
Lamp5
Rbm10
Trib2
Rpl27a
Zfp580
Rnf166
Ppp1r2
Mcts1
Lrrc10b
Ina
Dtx3
Rpp21
Egfl7
Otud4
Rhobtb2

Impdh1
Npy
Cyp46a1
Myo6
Shroom2
Anapc16
Rpl35a
Rpl8
Psmg4
Mrpl22
Rps19
Pgs1
Atp6v0a2
Pofut2
Dop1b
Cope
Ajap1
Coro2b
Ddx50
Ak3
Plcg1
Fgfr1op2
Pip4p1
Lage3
Cdk2ap2
Ndufa1
Cpsf1
Rprm
Med25
Gga3
Ppp6r3
3110039M20Rik
Elmo2
Rgs20
Ryr1
Eif4a3
Inha
Cdk16
Eloc
Olig1
Ptcd3
Mef2c
Snrpd3
Dgkd
Abhd14a
Fcor
Lrrc17
B9d2
Tram1l1
Adra2c
Rpl32
Gpr83

Tbc1d20
Med15
Ubp1
Pex6
Ahsa1
Mt2
Ttc4
Mrps18a
Pon2
Rps27rt
Ubr2
Rpsa
Ets2
Dclk2
Srm
Mrpl57
Rbis
Wipf3
Shisa4
Enah
Ccdc106
Rgs8
St3gal3
Capn2
Parp6
Cacna1c
Hs6st1
Ipo8
Adk
Akap9
Rpl36a
Mrtfa
Cops5
Reep2
Ssbp4
Plpbp
Rilpl1
Isyna1
Rida
Fndc5
Cfap410
Ino80e
Gm3764
Stk16
Clvs1
Aldh5a1
Rnf40
Ksr1
Dlg4
Traip
Xpo7
Tmem33

Tprkb
Vps4a
Smpdl3a
Tiam1
Mpi
Tmem126b
Cldn11
Dvl1
Clec2l
Ttc19
Klhl23
Igfbp7
Dpm2
Acy1
Mbnl2
Srrt
Lgi3
Psenen
Zcchc7
Psmg2
Nudt19
Nt5dc3
Pdia3
Abcc5
Eif2s3y
Dab1
Dnal4
Krt10
Rhoq
Stip1
Gpsm1
Adcy3
Cxxc1
Tmem185a
Mier2
Lrpprc
Map4k5
Smad3
Ankrd45
Surf4
Shisa7
Pdap1
Pcp4l1
Max
Bcas1
Fam120a
Tyro3
Ormdl3
Dnm2
Tars2
Dot1l
Orc4

Flna
Disp2
Sel1l
Zfr2
Rexo2
Gramd1b
Diras1
Foxo3
Prmt5
Cygb
Smox
Arih1
Tln1
Rnft2
Myh10
Tceal6
Smim27
Neat1
Ppp1r1b
Map1a
Speg
Prr7
Comt
Tmem70
Pcmtd1
Aup1
Ppp2r5a
Vrk1
Nicn1
Chid1
Tmem175
Chka
Chmp1a
Hivep2
Mpp3
Gng7
1500004A13Rik
Kcnab1
C1qb
Lratd1
Rapgefl1
Tmem234
Fbxw2
Rpl19
Cnih1
1110038F14Rik
Ppp1r13b
Il6st
Pde6d
Acap2
Ankrd46
Nubp2

Tenm1
Mrpl33
Fam193b
Gdpd1
Klhl13
Prpf8
Pip5k1c
Ramp2
Atad3a
Creld1
Rps19bp1
Magi2
Zfp511
Dagla
Mcm3ap
Med21
U2af2
Gfra4
Asic1
Mark1
Pigs
Cnot2
Usp32
Gnb2
Cmtm5
Sfxn3
Fnbp1l
Smg5
Cstf2
Ptcd2
Stub1
Lxn
Shank2
Slc25a23
Nae1
Isca2
Neu1
Limk2
Cxcl14
Nup50
Zfp871
Cbx7
Gramd1a
Syt3
Dlst
Nat8f1
Elovl5
Dnajc2
Dcaf7
Map7
Fam8a1
Vps13b

Kif17
Plpp1
Plppr3
Ints10
Rdh14
Slirp
Yipf3
Ppp6r1
Man1a2
Wasl
Rtf2
Gpr88
Jkamp
Rasgef1c
Selenos
Ndufaf5
Tango2
Stim2
Dlk2
Smim20
Baz1b
Gjb6
Czib
Ing4
Abhd17c
Comtd1
Chchd7
Opa1
Snrpa1
Anp32b
Rtn2
Trim46
Snx15
Hsph1
Mtx1
Begain
Sdr39u1
Ankrd10
Fam20c
Nr1h2
Dpm1
Krt12
Tsen15
Vcl
Abcb9
Cat
Cyb5r4
Timp3
Mocs2
Neto1
Immp1l
Car11

Sh3gl3
Psme2
Tmem223
Kansl3
Mlycd
Trak1
Ppp1cc
Mrpl19
Snapc5
Cdr1os
Rab4b
Stx8
Mrps36
Rubcn
Tomm5
Phf12
Hpcal1
Arhgap21
Cdc123
AU040320
Fads2
Ppp2r5b
Inpp1
Utrn
Polr2i
Htra1
Miga2
Inpp5j
Plxnb1
Ptma
Tsr3
Bcat1
Lrrc4b
Gtf3c2
Borcs6
Ydjc
Snx19
Timm10
Dhrs1
Naxd
Smim12
Wscd1
Scrt2
Sugt1
Ech1
Gba2
Cops2
Htr1b
Gm33651
Rap1a
Ssbp1
Tmem184b

Creld2
Ndufaf3
Kctd12
Ylpm1
Grin2a
Spsb3
Acap3
Rpl6
Pdzd2
Ppp1r37
Pik3c3
Actr1b
Ccndbp1
Syncrip
Ankrd13a
Smim26
Rtkn
Rraga
Nrbp2
Ablim1
Kifc2
Ccdc88c
Grik2
Ndufs1
Sec31a
Osbpl9
Trp53bp1
Gmpr2
Scrg1
Gnptg
Pbx2
Ss18l2
Erp29
Mrps11
Mir124-2hg
Srrd
Tmem240
Emc6
Pla2g7
Usp20
Mpv17l
Epb41l2
Nrbp1
Dctn3
Lgi2
Rrbp1
Letmd1
Fam110b
Mxd4
Dcun1d4
Med23
Tomm7

Dmtf1
Preb
Rala
Phf5a
Txnl1
Snx2
Ezr
Sucla2
Armc1
Tsc22d3
Uqcc3
Pdp1
Otud5
Zfp598
Hnrnpa2b1
Brwd1
Cuta
Phyhipl
Magoh
Lamtor3
Gapvd1
Cstf3
Dph3
Tomm34
Map4k2
Ranbp9
Rem2
Arf2
Leprot
Uchl3
Tpgs2
Pccb
Tmem258
Eif2b5
Pcdh9
Spire1
Tmem91
Smim18
Rab36
Hnrnpm
Dock10
Fn3k
Slc12a6
Eif4g3
Epb41l4aos
Pet100
Tra2a
Ncor2
Lmo7
Drd2
Zfp664
Lrrc45

Endog
Noc2l
Lars2
Rmdn3
Wdr83os
Pdzd11
Fcho1
Pno1
Kirrel3
Flot2
Zfp445
Zfp207
Csf1r
Cnot7
Btg3
Apoo
Med8
Exosc10
Asb13
Secisbp2l
Gm27032
Cnp
Pclo
Amph
Sec61b
Fut8
Cenpx
Ubac1
Lrrc42
Gt(ROSA)26Sor
Itgb1bp1
Jagn1
Eef1akmt1
Fam3c
Rin1
Stxbp6
Tssc4
Arhgef18
Slc6a11
Tspan9
Lrrtm3
Ttll7
Sacm1l
Emc8
Slain1
Rtl8a
Bap1
Tiam2
Mrpl46
Ryr3
Spata2l
Ndufaf2

Trim41
Kpna6
Slc50a1
Mrps14
Rprml
Cop1
Gtpbp6
Mctp1
Tbce
Spns1
Scn3b
Rpl10a
Trappc5
Tmem68
Gsk3a
Slc2a3
Entpd6
Gpr162
Smap2
Ccdc167
Cept1
Fbxo7
2310061I04Rik
Lysmd2
Rbm28
Bloc1s6
Mdp1
Rnf25
Ccser2
Tufm
B9d1
Tm2d2
Impdh2
Rcan1
Lrrfip1
Lyrm2
Suclg1
Vps52
Rbfa
Rimbp2
Osbpl8
Josd1
Ift22
Crabp1
Snn
Paip2b
Scamp3
Surf2
Hnrnph3
Atrx
Ubr4
Coa6

Abcd3
Ccdc97
Mb21d2
Zfp532
Sh3bgrl
Fam91a1
Rnmt
Prkab2
Srsf1
Cend1
Pnisr
Fem1b
L1cam
Serp1
Rab5c
Mark2
Ccng1
Tpp1
Kpnb1
Slc38a3
Lrrk2
Cog4
Sdhaf4
C1qtnf12
Emc4
Erbin
Nme7
Cntnap2
Armcx1
Cnot8
Ephx4
Nacc2
Stx16
Unc5a
Wac
Adam11
Csnk2a2
Ppil2
Ttll1
Dhrs7b
Cacna1b
Wtap
Leng8
Zfp385a
Dtnbp1
Mrps25
Timm29
Gga1
Ccs
Tyrobp
Zranb2
Iffo1

Kat7
Dcun1d5
Pop4
Gnb1
Chchd5
Col6a1
Rgs9
Gm10419
Homer1
Aldh1l1
Ptp4a3
Atat1
Gdpd5
Mapk6
Gmpr
Rapgef6
Herc2
Nrsn1
Cdk19
Ap3m1
Asrgl1
Mlec
Rab5if
Mrps6
Zcchc17
Ppp1r14a
Dhrs7
Rer1
Izumo4
Aip
C1qtnf4
Chrac1
Dnaja4
Usp33
Gsg1l
Dld
Gsto1
Mfap3l
Rnf181
Tmed3
Gbf1
Plppr4
Borcs8
Tmem14a
Jtb
Znhit3
Rasgef1b
Mthfsl
Wdr45
Sgcb
Hsd17b12
Kcnip4

Ramp1
Yeats4
Phyh
Dynll2
Slc25a12
Mrpl36
Ciao2b
Brd9
Fuom
2410002F23Rik
Tnks2
Wdr45b
Bax
Fam163b
Tmem38a
Cisd2
Nfyc
Tmem214
Mrpl15
Iah1
Acat1
Dnajc18
2010204K13Rik
Pnn
Mdm2
Hspbp1
Mrps17
Dtna
Trim3
Dnajb2
Dmac2
Pnpo
Isca1
Snd1
Gpr158
Chmp2b
Sp3os
Dpy30
Selenop
Nfasc
Ttbk1
Cox20
Snw1
Cacnb1
Vrk3
Foxk2
Churc1
Pim2
Pllp
Dpm3
Slc27a4
Robo2

Nr2c2
Cadm2
Ulk2
Parp1
Gng11
Vps29
Mrpl40
Med10
Copz1
Tafa5
Itch
Lamtor5
Akap5
Zpr1
Calcoco1
Bpgm
Alcam
Tmem201
Lman2
Txlna
Wdr13
Unc80
Kmt2e
Snx32
Apex1
Manf
Ube2i
Pstk
Fam107a
Ppm1b
Mtus1
Tmem127
Cnot6
BC004004
Sprn
Vps9d1
Trappc1
Tlk2
Tmbim4
Ube2h
Ccdc107
Pum1
Atp6v0e
Pin4
Paqr9
Men1
Pnrc1
Arfgef1
Dzip3
Rab24
Pkia
Clasrp

Mms19
Rgs17
Dhdds
Cyth2
Slc1a1
Hdgfl2
Dpysl4
Abca3
Txn1
Pdhb
Bcr
Atg3
Paqr4
Akap8l
Ndfip2
Rrn3
Ncbp2
Cacna1e
Mrps18b
Hectd1
Gtf2h5
Higd1a
Tpst2
Map9
C1qc
Ehd3
Capzb
Ptrhd1
Ccnh
Zmynd8
Sap30bp
Matk
Cep19
Cog1
Src
Cox14
Ppm1f
Itm2a
Api5
Zhx1
Myl12a
Myo9b
Ppip5k1
Tia1
Acadvl
Slc35e1
Ola1
Cebpzos
Med14
Emg1
Acp2
Lrtm2

1110059G10Rik
Ypel5
Agpat3
Fem1a
Prpf31
Srrm3
Sptan1
Mrpl16
Inpp5a
Ckap5
Agpat5
Cfap36
Ist1
Plxna2
Ugp2
Coa5
Gnb4
Washc2
Crbn
Hpcal4
Ppa2
Shfl
Ctss
Clip1
Kxd1
Abhd17b
Metrn
Kctd3
Hrh3
Impa1
Dgkh
Gng5
Ewsr1
Rfng
Smim19
Naa30
Ric3
Colgalt1
Pak6
Zfyve27
Hspa12a
Cops4
Car12
Psmd14
Hdac6
Ttc1
Ppp1r12a
Rbfox3
Aph1a
Ly6e
Tubg2
Rsrc2

Tax1bp1
Nprl2
Tmem183a
Osgep
Trap1
Cog7
Chd8
Etv5
Uba5
Tmem219
Ablim2
Mmd2
Dhx36
Bend6
Pbrm1
Mob4
Cst6
Smpd4
Adh5
Tesc
Rpl10
Cfap298
H13
Crat
Lst1
Nap1l5
Mrpl44
Mink1
Mrps9
Cacfd1
Wnk1
Tbc1d14
Ano8
Bptf
Mrpl43
Btbd1
Ndrg2
Lemd2
Cyld
Arpc5l
Asic4
Slc25a19
Pip5k1a
Stk39
Tmem60
Fam162a
Ddhd1
Nop56
Mrpl14
Mpp2
Med9
Vldlr

Rabac1
Exoc4
Polg
Mrpl18
Rnf34
Dag1
Rims2
Ndufaf7
Spock3
Prickle2
Ccnl2
Prelid3b
Mgrn1
Klhl22
Pnma2
Nfu1
Camk2d
Rbck1
Acd
Mrps30
Epb41l3
Atp13a2

Striatum cTBS

| DEGs Young sham vs Aged sham | cTBS DEGs unique to young adults | cTBS DEGs unique to young adults that are also ageing DEGs |
|---|---|---|
| Bc1 | Lrrc17 | Acyp2 |
| Cox8a | Rps28 | Ahcyl2 |
| Actb | Ndufa1 | Arpp21 |
| Rps24 | Pde10a | Bloc1s1 |
| Malat1 | Tomm7 | Chst2 |
| Rpl4 | Ost4 | Churc1 |
| Sst | Snhg9 | Comt |
| Fau | Tspyl4 | Cox8a |
| Dpysl2 | Araf | Cpe |
| Pcp4 | Basp1 | Cyp46a1 |
| Mt1 | Snap25 | Ddn |
| Rpl3 | Snrpf | Dlg4 |
| Rpl5 | Cox7c | Dpysl2 |
| Bloc1s1 | Rps27 | Elmo2 |
| Rasgrf2 | Bloc1s1 | Ewsr1 |
| Snca | Cox17 | Fam120a |
| Rpl37 | Atp5md | Fau |
| Rpl13 | Sem1 | Fgfr1op2 |
| Eef1a1 | Isca1 | Hspa5 |
| Ly6c1 | Dmtn | Isca1 |
| Usp50 | Snca | Itm2c |
| Nts | Ddn | Lrrc17 |
| Gamt | Smim26 | Mbp |
| Otof | Scn4b | Mt1 |
| Rpl14 | Ndrg4 | Mt2 |
| Rps10 | Snhg6 | Ndfip2 |
| Ntsr2 | Rps29 | Ndrg4 |
| Hspa5 | Phpt1 | Ndufa1 |
| Snhg6 | Slc6a1 | Ost4 |
| Ddn | Actr2 | Pdia3 |
| Lingo3 | Tatdn1 | Pet100 |
| Cpe | Arpp21 | Pfdn5 |
| Itm2c | Naa20 | Phyhipl |
| Rps8 | Ndufa3 | Plppr4 |
| Ptgds | Wsb2 | Ppp1r2 |
| Uba52 | Atp5mpl | Psd |
| Cacna1h | Car2 | R3hdm4 |
| Calm1 | Sdhd | Rap1a |
| Tsc22d1 | Ewsr1 | Rasd2 |
| Them6 | Snhg8 | Rasgrp2 |
| Atp1b1 | Pacsin1 | Rbis |
| Polr2l | Atp1a3 | Rpl10 |
| Ndrg4 | Plppr4 | Rpl13 |
| Cox7b | Sst | Rpl13a |
| Rpl13a | Fus | Rpl14 |
| Rps2 | Srsf5 | Rpl24 |
| Psd | App | Rpl27a |
| Rpl9 | Cdk5r2 | Rpl32 |
| Polb | Cops7a | Rpl4 |

Kcnip2
Strip2
Snhg11
Rps7
Josd2
Ankrd13b
Ndufa4
Nefm
Unc13c
Tac1
Pfdn5
Polr2j
Mbp
Snhg20
Necab1
Cck
Pacsin2
Arhgap33
Maneal
Mal
Higd2a
Coro7
Ccm2
Zdhhc14
Trak2
Dalrd3
Rps3a1
Jsrp1
Arpp21
Lrpap1
Camk1g
Snhg1
Srsf11
Mrpl41
Pitpnm2
Eef1a2
Gucy1a1
Sdhaf2
Chordc1
Rac3
Acvr2a
Rpl9-ps6
Rnf20
Srp72
Smpd3
Rps14
Yipf1
Morf4l1
Acsbg1
Rps27a
Mapk1ip1
Ppid

Tmem258
Rpl39
Huwe1
Txn1
Spag9
Acyp2
1810037I17Rik
Cntnap1
Rnf5
Sparcl1
Mbp
Rps27rt
Rap1gds1
Chmp5
Mt2
Slc25a22
Kctd13
Ppp1r16b
R3hdm4
Rab3c
Mt1
Ppp1r9b
Agap2
Gucy1b1
Gng13
Rgs7bp
Selenof
Kif5a
Cox8a
Rasd2
Meg3
Rpl9-ps6
Jph4
Atp1b2
Sec61g
Atp6v1a
Cs
Snrpg
Camk2a
Rheb
Lzts3
Ndufb3
Tsc22d4
Comt
Cadm4
Atxn2l
Pet100
Zwint
Arpc4
Rcn2
Sipa1l1
Atp5k

Rpl5
Rpl8
Rpl9-ps6
Rps10
Rps14
Rps19
Rps2
Rps20
Rps24
Rps27a
Rps27rt
Rps3a1
Rps4x
Rps5
Rps7
Rps8
Rpsa
Rraga
Scn4b
Smim26
Snca
Snhg11
Snhg3
Snhg6
Sst
Tmem258
Tomm7
Txn1
Usp50

| | |
|---|---|
| Oga | Usp50 |
| Sirt2 | Stxbp1 |
| Mat2a | Cpe |
| Pfn1 | Snhg11 |
| Btbd3 | Cox6c |
| Agrn | Churc1 |
| Elp2 | Senp3 |
| Ralgds | Glul |
| Sirt3 | Abat |
| Scn4b | Chst2 |
| Limd2 | Meis2 |
| Mrpl21 | Paip2 |
| Slc2a1 | Hypk |
| Emc2 | Ndfip2 |
| Lyst | Rapgef4 |
| Ly6a | Mapre2 |
| Rps4x | Fbxl16 |
| Epas1 | Ppp1r2 |
| Taf13 | Dlg4 |
| Tmem230 | Cfap20 |
| Ost4 | Prrc2a |
| Kansl2 | Eif4a2 |
| Amz2 | Snhg3 |
| Filip1 | Mtmr9 |
| Arel1 | Sdcbp |
| Slc35d3 | Cacna2d3 |
| Rgl2 | Fam120a |
| Acyp2 | Rgs14 |
| Tmub2 | Rab28 |
| Rpl30 | Ttc7b |
| Nedd4l | Rasgrp2 |
| Tubb5 | Ikbkb |
| Tbc1d9b | Ahcyl2 |
| Pttg1ip | Add2 |
| Prxl2a | Hectd4 |
| Brinp1 | B2m |
| Nlgn1 | Dbndd2 |
| Trmt1 | Hprt |
| Cbx4 | Gpm6b |
| Inf2 | Mrps28 |
| Rasd2 | Erh |
| Pvalb | Atl1 |
| Ccdc28b | Elmo2 |
| Timm9 | Phyhip |
| Grm4 | Trrap |
| Ubap2l | Rbis |
| Adcy5 | Arhgap39 |
| Stmn1 | Phyhipl |
| Trmt2a | Mrpl27 |
| Snhg3 | Pdia3 |
| Myo5b | Dnajb4 |
| Mydgf | Hspa5 |

| | |
|---|---|
| Phldb1 | Cd302 |
| Ahcyl2 | Atn1 |
| Arhgef2 | Fgfr1op2 |
| Tmem126a | Psd |
| Sptbn4 | Acsl3 |
| Kif21b | Rraga |
| Cdo1 | Wdr89 |
| Adam15 | Cyp46a1 |
| Ctxn1 | Polr2k |
| Ndufa7 | Atp5g3 |
| Hip1r | Miga1 |
| Mrps12 | Rap1a |
| Panx2 | Gas5 |
| Zmat5 | Srrm2 |
| Ptpro | Slc4a4 |
| Cinp | Ank |
| Rps6ka3 | Cstb |
| Ypel4 | Rpl18 |
| Mirg | Rps3 |
| Txndc16 | Rps4x |
| Mir124a-1hg | Rps3a1 |
| Rps20 | Eef1b2 |
| Adora2a | Syt11 |
| Cspg5 | Rpl21 |
| Fam210b | Dpysl2 |
| Ywhah | Rpsa |
| Rps5 | Rpl23a |
| Ephx1 | Rpl24 |
| Cyp51 | Rpl27a |
| Mcee | Rps7 |
| Pds5b | Rpl11 |
| Ppp5c | Rps15 |
| Mog | Rps27a |
| Cdk4 | Ttr |
| Chn2 | Rpl8 |
| Tmem208 | Nenf |
| Carm1 | Rps19 |
| Degs1 | Rplp1 |
| Fxyd2 | Rpl32 |
| Creg1 | Itm2c |
| Fyco1 | Fau |
| Large1 | Rpl5 |
| Cox19 | Rpl18a |
| Slc17a7 | Rps20 |
| Ddit3 | Eno1 |
| Hspe1 | Rps5 |
| Pop7 | Rpl17 |
| Kcnk1 | Rps2 |
| Luc7l | Rps12 |
| Dlx6os1 | Pfdn5 |
| Tmem167 | Rps10 |
| Fhl1 | Rps11 |

Rasgrp2
Mcf2l
Adarb1
Senp2
Aldh7a1
R3hdm4
Rpl24
Slco1c1
Dpf2
Tmod1
Ythdf1
Pdia6
Eml5
Ldah
Scg2
Zmym3
Polr3e
Tspan2
Kcnd3
Mobp
Dmac1
Sirt7
Plekhj1
Coq2
Cldn5
Ilkap
Nt5c3b
Mdk
Gadd45g
Chst2
Wfs1
Mmp17
Il18
Ube2f
Rell2
Dnajb11
Gap43
Lamp5
Rbm10
Trib2
Rpl27a
Zfp580
Rnf166
Ppp1r2
Mcts1
Lrrc10b
Ina
Dtx3
Rpp21
Egfl7
Otud4
Rhobtb2

Rps14
Rpl13
Rpl4
Atp9a
Rpl26
Rps8
Rpl13a
Rps24
Rpl14
Hbb-bs
Rpl10

Impdh1
Npy
Cyp46a1
Myo6
Shroom2
Anapc16
Rpl35a
Rpl8
Psmg4
Mrpl22
Rps19
Pgs1
Atp6v0a2
Pofut2
Dop1b
Cope
Ajap1
Coro2b
Ddx50
Ak3
Plcg1
Fgfr1op2
Pip4p1
Lage3
Cdk2ap2
Ndufa1
Cpsf1
Rprm
Med25
Gga3
Ppp6r3
3110039M20Rik
Elmo2
Rgs20
Ryr1
Eif4a3
Inha
Cdk16
Eloc
Olig1
Ptcd3
Mef2c
Snrpd3
Dgkd
Abhd14a
Fcor
Lrrc17
B9d2
Tram1l1
Adra2c
Rpl32
Gpr83

Tbc1d20
Med15
Ubp1
Pex6
Ahsa1
Mt2
Ttc4
Mrps18a
Pon2
Rps27rt
Ubr2
Rpsa
Ets2
Dclk2
Srm
Mrpl57
Rbis
Wipf3
Shisa4
Enah
Ccdc106
Rgs8
St3gal3
Capn2
Parp6
Cacna1c
Hs6st1
Ipo8
Adk
Akap9
Rpl36a
Mrtfa
Cops5
Reep2
Ssbp4
Plpbp
Rilpl1
Isyna1
Rida
Fndc5
Cfap410
Ino80e
Gm3764
Stk16
Clvs1
Aldh5a1
Rnf40
Ksr1
Dlg4
Traip
Xpo7
Tmem33

Tprkb
Vps4a
Smpdl3a
Tiam1
Mpi
Tmem126b
Cldn11
Dvl1
Clec2l
Ttc19
Klhl23
Igfbp7
Dpm2
Acy1
Mbnl2
Srrt
Lgi3
Psenen
Zcchc7
Psmg2
Nudt19
Nt5dc3
Pdia3
Abcc5
Eif2s3y
Dab1
Dnal4
Krt10
Rhoq
Stip1
Gpsm1
Adcy3
Cxxc1
Tmem185a
Mier2
Lrpprc
Map4k5
Smad3
Ankrd45
Surf4
Shisa7
Pdap1
Pcp4l1
Max
Bcas1
Fam120a
Tyro3
Ormdl3
Dnm2
Tars2
Dot1l
Orc4

Flna
Disp2
Sel1l
Zfr2
Rexo2
Gramd1b
Diras1
Foxo3
Prmt5
Cygb
Smox
Arih1
Tln1
Rnft2
Myh10
Tceal6
Smim27
Neat1
Ppp1r1b
Map1a
Speg
Prr7
Comt
Tmem70
Pcmtd1
Aup1
Ppp2r5a
Vrk1
Nicn1
Chid1
Tmem175
Chka
Chmp1a
Hivep2
Mpp3
Gng7
1500004A13Rik
Kcnab1
C1qb
Lratd1
Rapgefl1
Tmem234
Fbxw2
Rpl19
Cnih1
1110038F14Rik
Ppp1r13b
Il6st
Pde6d
Acap2
Ankrd46
Nubp2

Tenm1
Mrpl33
Fam193b
Gdpd1
Klhl13
Prpf8
Pip5k1c
Ramp2
Atad3a
Creld1
Rps19bp1
Magi2
Zfp511
Dagla
Mcm3ap
Med21
U2af2
Gfra4
Asic1
Mark1
Pigs
Cnot2
Usp32
Gnb2
Cmtm5
Sfxn3
Fnbp1l
Smg5
Cstf2
Ptcd2
Stub1
Lxn
Shank2
Slc25a23
Nae1
Isca2
Neu1
Limk2
Cxcl14
Nup50
Zfp871
Cbx7
Gramd1a
Syt3
Dlst
Nat8f1
Elovl5
Dnajc2
Dcaf7
Map7
Fam8a1
Vps13b

Kif17
Plpp1
Plppr3
Ints10
Rdh14
Slirp
Yipf3
Ppp6r1
Man1a2
Wasl
Rtf2
Gpr88
Jkamp
Rasgef1c
Selenos
Ndufaf5
Tango2
Stim2
Dlk2
Smim20
Baz1b
Gjb6
Czib
Ing4
Abhd17c
Comtd1
Chchd7
Opa1
Snrpa1
Anp32b
Rtn2
Trim46
Snx15
Hsph1
Mtx1
Begain
Sdr39u1
Ankrd10
Fam20c
Nr1h2
Dpm1
Krt12
Tsen15
Vcl
Abcb9
Cat
Cyb5r4
Timp3
Mocs2
Neto1
Immp1l
Car11

Sh3gl3
Psme2
Tmem223
Kansl3
Mlycd
Trak1
Ppp1cc
Mrpl19
Snapc5
Cdr1os
Rab4b
Stx8
Mrps36
Rubcn
Tomm5
Phf12
Hpcal1
Arhgap21
Cdc123
AU040320
Fads2
Ppp2r5b
Inpp1
Utrn
Polr2i
Htra1
Miga2
Inpp5j
Plxnb1
Ptma
Tsr3
Bcat1
Lrrc4b
Gtf3c2
Borcs6
Ydjc
Snx19
Timm10
Dhrs1
Naxd
Smim12
Wscd1
Scrt2
Sugt1
Ech1
Gba2
Cops2
Htr1b
Gm33651
Rap1a
Ssbp1
Tmem184b

Creld2
Ndufaf3
Kctd12
Ylpm1
Grin2a
Spsb3
Acap3
Rpl6
Pdzd2
Ppp1r37
Pik3c3
Actr1b
Ccndbp1
Syncrip
Ankrd13a
Smim26
Rtkn
Rraga
Nrbp2
Ablim1
Kifc2
Ccdc88c
Grik2
Ndufs1
Sec31a
Osbpl9
Trp53bp1
Gmpr2
Scrg1
Gnptg
Pbx2
Ss18l2
Erp29
Mrps11
Mir124-2hg
Srrd
Tmem240
Emc6
Pla2g7
Usp20
Mpv17l
Epb41l2
Nrbp1
Dctn3
Lgi2
Rrbp1
Letmd1
Fam110b
Mxd4
Dcun1d4
Med23
Tomm7

Dmtf1
Preb
Rala
Phf5a
Txnl1
Snx2
Ezr
Sucla2
Armc1
Tsc22d3
Uqcc3
Pdp1
Otud5
Zfp598
Hnrnpa2b1
Brwd1
Cuta
Phyhipl
Magoh
Lamtor3
Gapvd1
Cstf3
Dph3
Tomm34
Map4k2
Ranbp9
Rem2
Arf2
Leprot
Uchl3
Tpgs2
Pccb
Tmem258
Eif2b5
Pcdh9
Spire1
Tmem91
Smim18
Rab36
Hnrnpm
Dock10
Fn3k
Slc12a6
Eif4g3
Epb41l4aos
Pet100
Tra2a
Ncor2
Lmo7
Drd2
Zfp664
Lrrc45

Endog
Noc2l
Lars2
Rmdn3
Wdr83os
Pdzd11
Fcho1
Pno1
Kirrel3
Flot2
Zfp445
Zfp207
Csf1r
Cnot7
Btg3
Apoo
Med8
Exosc10
Asb13
Secisbp2l
Gm27032
Cnp
Pclo
Amph
Sec61b
Fut8
Cenpx
Ubac1
Lrrc42
Gt(ROSA)26Sor
Itgb1bp1
Jagn1
Eef1akmt1
Fam3c
Rin1
Stxbp6
Tssc4
Arhgef18
Slc6a11
Tspan9
Lrrtm3
Ttll7
Sacm1l
Emc8
Slain1
Rtl8a
Bap1
Tiam2
Mrpl46
Ryr3
Spata2l
Ndufaf2

Trim41
Kpna6
Slc50a1
Mrps14
Rprml
Cop1
Gtpbp6
Mctp1
Tbce
Spns1
Scn3b
Rpl10a
Trappc5
Tmem68
Gsk3a
Slc2a3
Entpd6
Gpr162
Smap2
Ccdc167
Cept1
Fbxo7
2310061I04Rik
Lysmd2
Rbm28
Bloc1s6
Mdp1
Rnf25
Ccser2
Tufm
B9d1
Tm2d2
Impdh2
Rcan1
Lrrfip1
Lyrm2
Suclg1
Vps52
Rbfa
Rimbp2
Osbpl8
Josd1
Ift22
Crabp1
Snn
Paip2b
Scamp3
Surf2
Hnrnph3
Atrx
Ubr4
Coa6

Abcd3
Ccdc97
Mb21d2
Zfp532
Sh3bgrl
Fam91a1
Rnmt
Prkab2
Srsf1
Cend1
Pnisr
Fem1b
L1cam
Serp1
Rab5c
Mark2
Ccng1
Tpp1
Kpnb1
Slc38a3
Lrrk2
Cog4
Sdhaf4
C1qtnf12
Emc4
Erbin
Nme7
Cntnap2
Armcx1
Cnot8
Ephx4
Nacc2
Stx16
Unc5a
Wac
Adam11
Csnk2a2
Ppil2
Ttll1
Dhrs7b
Cacna1b
Wtap
Leng8
Zfp385a
Dtnbp1
Mrps25
Timm29
Gga1
Ccs
Tyrobp
Zranb2
Iffo1

Kat7
Dcun1d5
Pop4
Gnb1
Chchd5
Col6a1
Rgs9
Gm10419
Homer1
Aldh1l1
Ptp4a3
Atat1
Gdpd5
Mapk6
Gmpr
Rapgef6
Herc2
Nrsn1
Cdk19
Ap3m1
Asrgl1
Mlec
Rab5if
Mrps6
Zcchc17
Ppp1r14a
Dhrs7
Rer1
Izumo4
Aip
C1qtnf4
Chrac1
Dnaja4
Usp33
Gsg1l
Dld
Gsto1
Mfap3l
Rnf181
Tmed3
Gbf1
Plppr4
Borcs8
Tmem14a
Jtb
Znhit3
Rasgef1b
Mthfsl
Wdr45
Sgcb
Hsd17b12
Kcnip4

Ramp1
Yeats4
Phyh
Dynll2
Slc25a12
Mrpl36
Ciao2b
Brd9
Fuom
2410002F23Rik
Tnks2
Wdr45b
Bax
Fam163b
Tmem38a
Cisd2
Nfyc
Tmem214
Mrpl15
Iah1
Acat1
Dnajc18
2010204K13Rik
Pnn
Mdm2
Hspbp1
Mrps17
Dtna
Trim3
Dnajb2
Dmac2
Pnpo
Isca1
Snd1
Gpr158
Chmp2b
Sp3os
Dpy30
Selenop
Nfasc
Ttbk1
Cox20
Snw1
Cacnb1
Vrk3
Foxk2
Churc1
Pim2
Pllp
Dpm3
Slc27a4
Robo2

Nr2c2
Cadm2
Ulk2
Parp1
Gng11
Vps29
Mrpl40
Med10
Copz1
Tafa5
Itch
Lamtor5
Akap5
Zpr1
Calcoco1
Bpgm
Alcam
Tmem201
Lman2
Txlna
Wdr13
Unc80
Kmt2e
Snx32
Apex1
Manf
Ube2i
Pstk
Fam107a
Ppm1b
Mtus1
Tmem127
Cnot6
BC004004
Sprn
Vps9d1
Trappc1
Tlk2
Tmbim4
Ube2h
Ccdc107
Pum1
Atp6v0e
Pin4
Paqr9
Men1
Pnrc1
Arfgef1
Dzip3
Rab24
Pkia
Clasrp

Mms19
Rgs17
Dhdds
Cyth2
Slc1a1
Hdgfl2
Dpysl4
Abca3
Txn1
Pdhb
Bcr
Atg3
Paqr4
Akap8l
Ndfip2
Rrn3
Ncbp2
Cacna1e
Mrps18b
Hectd1
Gtf2h5
Higd1a
Tpst2
Map9
C1qc
Ehd3
Capzb
Ptrhd1
Ccnh
Zmynd8
Sap30bp
Matk
Cep19
Cog1
Src
Cox14
Ppm1f
Itm2a
Api5
Zhx1
Myl12a
Myo9b
Ppip5k1
Tia1
Acadvl
Slc35e1
Ola1
Cebpzos
Med14
Emg1
Acp2
Lrtm2

1110059G10Rik
Ypel5
Agpat3
Fem1a
Prpf31
Srrm3
Sptan1
Mrpl16
Inpp5a
Ckap5
Agpat5
Cfap36
Ist1
Plxna2
Ugp2
Coa5
Gnb4
Washc2
Crbn
Hpcal4
Ppa2
Shfl
Ctss
Clip1
Kxd1
Abhd17b
Metrn
Kctd3
Hrh3
Impa1
Dgkh
Gng5
Ewsr1
Rfng
Smim19
Naa30
Ric3
Colgalt1
Pak6
Zfyve27
Hspa12a
Cops4
Car12
Psmd14
Hdac6
Ttc1
Ppp1r12a
Rbfox3
Aph1a
Ly6e
Tubg2
Rsrc2

Tax1bp1
Nprl2
Tmem183a
Osgep
Trap1
Cog7
Chd8
Etv5
Uba5
Tmem219
Ablim2
Mmd2
Dhx36
Bend6
Pbrm1
Mob4
Cst6
Smpd4
Adh5
Tesc
Rpl10
Cfap298
H13
Crat
Lst1
Nap1l5
Mrpl44
Mink1
Mrps9
Cacfd1
Wnk1
Tbc1d14
Ano8
Bptf
Mrpl43
Btbd1
Ndrg2
Lemd2
Cyld
Arpc5l
Asic4
Slc25a19
Pip5k1a
Stk39
Tmem60
Fam162a
Ddhd1
Nop56
Mrpl14
Mpp2
Med9
Vldlr

Rabac1
Exoc4
Polg
Mrpl18
Rnf34
Dag1
Rims2
Ndufaf7
Spock3
Prickle2
Ccnl2
Prelid3b
Mgrn1
Klhl22
Pnma2
Nfu1
Camk2d
Rbck1
Acd
Mrps30
Epb41l3
Atp13a2

WMT iTBS

| DEGs Young sham vs Aged sham | iTBS DEGs unique to young adults | iTBS DEGs unique to young adults that are also ageing DEGs |
|---|---|---|
| Mobp | Penk | AI593442 |
| Mbp | Rgs9 | Apod |
| Uba52 | Pde1b | Calb1 |
| Cabp1 | Calb1 | Cd82 |
| Plekhb1 | Gnal | Cldn11 |
| Bcas1 | Pcp4 | Csrp1 |
| Chn1 | Ppp1r1b | Desi1 |
| Hpca | AI593442 | Fbxl16 |
| Scn1b | Ptpn5 | Gnal |
| Bsn | Zwint | Gng11 |
| Slc17a7 | Fbxl16 | Kcnab1 |
| Tesc | Tac1 | Lrrc17 |
| Lingo1 | Prrc2b | Mag |
| Ppp3r1 | Kcnab1 | Metrn |
| Nrgn | Rap1gap | Nkx6-2 |
| Nptxr | Nedd4 | Opalin |
| Mef2c | Gng11 | Ppp1r14a |
| Rpl9-ps6 | Aplp1 | Prrc2b |
| Phf24 | Cdc42ep1 | Ptgds |
| Eef1a2 | Opalin | Ptpn5 |
| Sv2b | Abca2 | Rgs9 |
| Gria3 | Cldn11 | Rhog |
| Cacnb3 | Rhog | Tac1 |
| Lamp5 | Metrn | Tatdn1 |
| Pvalb | Csrp1 | Trf |
| Cadm3 | Cd82 | |
| Mal | Apod | |
| Mt1 | Desi1 | |
| Dnm1 | Gjb1 | |
| Rprml | Nkx6-2 | |
| Camkk2 | Ppp1r14a | |
| Plk2 | Lgi3 | |
| Pi4ka | Cnot3 | |
| Trp53inp2 | Trf | |
| Atp1a1 | Mag | |
| Fabp3 | Gm2000 | |
| Itpka | Tubb4a | |
| Tceal5 | Tatdn1 | |
| Nat8l | Ptgds | |
| Prr18 | Lrrc17 | |
| Tmeff2 | | |
| Tmsb4x | | |
| Tceal6 | | |
| Bloc1s1 | | |
| Kcnab2 | | |
| Ptk2b | | |
| Pdlim2 | | |
| Pdp1 | | |
| Matk | | |

Lmo4
Clic4
Camk2b
Ppp3ca
Sri
Cnp
Rps29
Stx1a
Lin7b
Svop
Brinp1
Mmp17
Grin1
Kalrn
Snhg6
Cplx1
Rnf112
Pgm2l1
Calm1
Tspan2
Thy1
Spock1
Nell2
Gpr162
Cx3cl1
Pea15a
Fam131a
Cdk5r1
Ptprz1
Egr1
Golga7
Polr3e
Rell2
Cap2
Fxyd1
Snap25
Atp2b2
Pkm
Syn1
Kcnip2
Trak2
Habp4
Arf3
Snph
Myh10
Celf2
Marcksl1
Mrtfb
Slc6a17
Evi2a
Rbfox3
Itpr1

Tppp3
Syp
Tmem132a
Scrg1
Camk2a
Cyfip2
R3hdm1
Ier5
Prkcg
Cplx2
Cck
St3gal5
Rasgef1a
Gnb5
Calb1
Carhsp1
Snhg9
Zbtb18
Nrxn3
Fa2h
Gls
Sccpdh
Lynx1
Gabrg2
Car11
Ncs1
Myo6
Gamt
Cacng3
Bcat1
Phgdh
Baiap2
Rab31
Plcb1
Ak3
Pianp
Jph3
Reep2
AI593442
Ugt8a
Tns3
Adcy1
Tac1
Serpini1
Nsf
Tmem59l
Mog
Josd2
Myt1l
Grina
Madd
Timm22

Gapdh
Pcsk2
Snx10
Napb
Mapk10
Tpi1
Sncb
Litaf
Ywhah
Dgkz
Dbi
Grik5
Tbc1d9b
Tatdn1
Tmeff1
Cpe
Icam5
Gng12
Snca
BC004004
Ddah1
Cmtm5
Rgs9
Rasgrp1
Syngr3
Nfe2l1
Rps28
Dcaf17
Setd7
Ptprn
Igfbp5
Peg13
St8sia3
Grb14
Cdk2ap1
Cldn11
Tusc3
Gabrb3
Ckmt1
Gnal
Paqr8
Tspan13
Slc2a1
Syt4
Qdpr
Dmtn
Sh3gl3
Ugp2
Ppp1r9a
Dbn1
Cat
Atp6v0e

Phyhip
Cdc37l1
Rnf157
Aak1
Mgst3
Lamp1
Aktip
Tsnax
Arl6ip5
Dock10
Gabbr2
Kcnj4
Rps27
Arhgef9
Ptpn11
Vbp1
Rab15
Ncor2
Gnai1
Tmem47
Ewsr1
Enc1
Ptpn5
Unc50
Eno2
Nkiras1
Id3
Pip5k1c
Tra2a
Fhl1
Myo5a
Aqp4
Cd9
Neurl1a
Reps2
Rasl10b
Stmn3
Anp32b
Efhd2
Bex2
Ppp2r5a
Rusc1
Cadps
Cfap36
2310022B05Rik
Scg2
Rab3a
Ivns1abp
Dlgap1
Syt13
Rpl4
Mt2

Tnks2
Oip5os1
Rasd2
Homer1
Strn4
Ywhaz
Dock3
Tmco1
Nefm
Tubb5
Tmem179
Evl
Fyn
Kcnq2
Rab6b
Dbp
Chga
Tuba4a
Plekhj1
Zfas1
Clstn3
Ednrb
Syn2
Add2
Dlgap4
Gsn
Kndc1
Pja2
Got2
Celf5
Cryab
Serinc5
Ddit3
Daam2
Rhog
Cadm2
Shtn1
Dgkb
Vamp2
Churc1
Rps12
Lrrn1
Mapk9
Sar1b
Adgrb2
Prkcb
Map4k4
Unc13a
Adgrl1
Kcnk1
Atp6v1b2
Zfand5

Pip4k2a
Otub1
Scd1
Asphd2
Sptbn2
Egr3
Fgf1
Idh3a
Fads2
Enah
Cnn3
Atxn7l3
B3gat3
Usp14
Arrdc3
Stk25
Scg5
Aamdc
Camta2
Scn2a
Cntn2
Sorl1
Anapc2
Nptn
Rdx
Nsg2
Wnk1
Stxbp1
Dkk3
Sc5d
C1qtnf4
Gal3st1
Ap1b1
Cyb5b
Fam234b
Zfp664
Ptprd
Kctd17
Got1
Dclk1
Prkce
Tcf4
Actr1b
Rnf130
Meg3
Atp6v0c
Picalm
Clstn1
Gpr88
Rad21
Elovl5
Adgrb1

Rian
Chst1
Nat14
Ralgds
Rps27l
Astn1
B230219D22Rik
Wasf1
Ogdh
Epb41l2
Pltp
Syt5
Fxyd7
Abhd4
Txndc9
Necap1
Ankrd13a
Plekhh1
Ndrg1
Ado
Snhg11
Plp1
Aspa
Med21
Smim13
Cdc42se2
Arhgap23
Malat1
Chn2
Ddr1
Taf13
Eif6
Mllt11
Nrbp2
Ctnna1
Tprkb
Rnf227
Olig2
Pdxdc1
Ap2a2
Strn
Nrcam
S1pr5
Vgf
Mmd
Cd81
Pou3f3
Sirt2
Rph3a
Nefl
Olig1
Cd164

Plekha1
Cdk17
Gstm7
Myl12b
Scarb2
Nhp2
Srpk2
Srsf11
Ppp2r3a
Efhd1
Gjc2
Snn
Ptk2
Pdhb
Mras
Jam3
Hepacam
Hadha
Atp6v1e1
Nacc2
Gltp
Sh2d5
Micall1
Snap91
Arpc1b
Khdrbs3
Pop5
Slc48a1
Mrpl17
Arhgdig
Fdx2
Snapin
Gabra1
Pam16
Eno1
Ufm1
Rnps1
Rtkn
Haghl
Timm23
Arpp21
Tmem134
Igfbp6
Smap2
Mrpl4
Zcchc24
Mrps26
Eci1
Adcyap1r1
Cers2
Strbp
Arhgap5

Tmem229a
Phf5a
Asns
Erbin
Ddx24
Pop4
Phlda3
Lrp1
Trim9
Pim3
Pik3r1
Pfkl
1700047M11Rik
Cxcl14
Gnptg
Slc44a1
Rpl37a
Ccdc124
Ina
Gjc3
Pdcd10
2900052N01Rik
Tusc2
Mat2b
Stard7
Dync1i1
Wscd1
Lrrc58
Kifc2
Rpl35
Sdcbp
Atp1b1
Usp5
Aldh6a1
Sptbn4
Anxa5
Adgrg1
Schip1
Ybx3
Fez1
Ifi27
Tmbim4
Ssbp4
Selenop
Hras
Trim37
Fbxl16
Rogdi
Syt7
Csnk1d
Ten1
Adk

Nfu1
Lpar1
Prkar1b
Hnrnpdl
Lamp2
Slc25a11
Camkv
Rnf181
Ppa1
Sult4a1
Zer1
Gas5
B4gat1
Hcn2
Gpbp1
Atp6v1g2
Mtus1
Snx5
Tmem208
Arhgef4
Sh3gl2
Calm2
Phldb1
Sptssa
Emc3
Wipf3
Hypk
Ccl27a
Spock2
Ei24
Prxl2b
Egln1
Commd4
Ttc9b
Znhit1
Paqr4
Apc
Cdipt
Arhgap33
Gria1
Oaz2
Sox2ot
Mapk8ip2
Pttg1ip
Cdk5r2
Pld3
Ssr3
Rbm3
Svip
Mrpl12
Pacsin1
Trim32

Map4
S1pr1
Ube2ql1
Rgs4
Prrt1
Clcn3
Map7
Ctxn1
Smim11
Nrsn2
Sugt1
Gda
Pde6d
Kcnab1
Rap1b
Sec13
Kdelr1
Usp54
Ndrg4
Rbbp4
Slain1
Dync1i2
Grhpr
Gm2a
Kif1c
Atp2b1
Ngef
Gstp1
Atg3
Echs1
Secisbp2l
Ppp1r2
Necab2
Ctdsp2
Ppt1
Cox5a
Plcl1
Gsk3a
Dnm1l
Tmem88b
Ppp2r2a
Dnajb2
Arhgef17
Commd6
Zfp207
Prmt2
Dctn4
Sulf2
Cavin3
Ociad2
Ncoa4
Car14

Rnf208
Mboat7
Chchd6
Gna12
Chgb
Cbx6
Lrrc17
Tfg
Dixdc1
Vkorc1
Scn2b
Tomm70a
Dnajc15
Kif5a
Tmem33
Ddx1
Tspan5
Ermn
Kcnj10
Il18
Nipa1
Sbf1
Sbds
Polr3h
Plin3
Gng11
Klhdc3
Bcap31
Hnrnph2
Pogk
Epas1
Iqsec1
Klf13
Camta1
Btbd1
Add3
M6pr
Gng3
Sumo3
Sema4d
Nrxn2
Slc12a2
Atp6v1a
Clk1
Gcsh
Sucla2
Psmc1
Stx1b
Pak2
Herc1
Atp6v1d
Ppp1r14b

Eif3e
Snhg12
Ncdn
Mtss2
Grn
Efnb3
Oxsr1
Spock3
Snrnp27
Ppp2r1a
Eml1
Kif3c
Syf2
Tmx4
Dlg4
Clip2
Disp2
Ypel5
Rufy3
Rit2
Bnip3
Sox10
Ppp1r11
Fnta
Scrn1
Map2k1
Papss1
Hectd4
Sh3glb1
Psph
Vstm2l
Adh5
Prpf19
Dip2a
Sfpq
Ly6e
Nptx1
2210016L21Rik
Serpinb6a
Slc25a12
Prrc2b
5031439G07Rik
Polr2l
Rnf5
Tmem98
Hk1
Pgam1
Zmiz2
Rbbp6
Tmem160
Scamp1
Slc24a2

Pcna
Eif1ax
Cyp46a1
Gja1
Dtx3
Slc25a23
Mbnl1
Agpat4
Rida
Pkig
Clmn
Mycbp2
Dcaf8
Cpd
Rasgrf1
Mink1
Camk2g
Tmcc3
Gabrd
Spin1
Fgfr2
Gclm
Dlgap3
Syngr1
Ptgds
Nsfl1c
Fnbp1
Hnrnpr
Retreg2
Lxn
Rap1a
Tmed4
Spryd3
Tmcc2
Pcif1
Arrb1
Slc35b1
Arl1
Magee1
Ubald1
Sh3glb2
Trappc2l
Paip2b
Akr7a5
Nckap1
Maf1
Ppme1
Hnrnph3
Amer2
Tmem158
Fam168b
Epdr1

Abhd17b
Fbxo9
Podxl2
Trpc4ap
Glrx5
Arpc2
Kctd3
Trim3
Etv1
Hexa
Dnm2
Vamp3
Baalc
Cldnd1
Tpd52
Jup
Dnajc5
Actr10
Ppm1a
Ctnnd2
Commd1
Dtna
Acot13
Napg
Mcts1
Spag9
Ddit4
Stard3nl
Rexo2
Rnf141
Mtmr6
Gab1
Metap2
Mtfr1l
Snap47
S100a6
Mrpl53
Ly6a
Alcam
Anln
Taok1
Polr2c
Psd3
Stk11
Dlg2
Kmt5a
Celsr2
Tmod1
Psma1
Pfkp
Ppp2r5c
Adipor2

Cxxc5
Gps1
Zeb2
Gprasp1
Epn2
Ndufa4
Cyb5r3
Slu7
Cdk19
Cdk2ap2
Ufc1
Eif3a
Tulp4
Ap1s1
Cenpb
Abi2
Asrgl1
Znrf1
Hnrnph1
Bag6
Cs
Ptprs
Fkbp8
Tspan15
Smox
Txnl1
Ccp110
Ensa
Pdk2
Ift20
Nsa2
Gprc5b
Slc32a1
Dner
Dnlz
Tmbim1
Srsf1
Hopx
Ppp1r14a
Abhd8
Txnl4a
Rnh1
Enoph1
Ttc3
Lzts2
Tpm1
Csnk2b
Hdhd2
Gad1
Cops6
Mapre3
Tnfaip6

Anp32e
Psat1
Tsg101
Tpd52l2
Epn1
Rad23b
Mrpl28
Gfap
Mrps14
Rgs7bp
Gap43
Itpk1
Gpr62
Il33
Mrpl24
Nkx6-2
Parp6
Pin1
Mrfap1
Itm2c
Cbarp
Pdcd5
Tcf25
Vdac1
Clasp2
Pllp
Ndufb5
Mapk8ip3
Eif4a3
Ppp1r16b
Uqcc2
Rplp0
Lrpap1
Adcy5
Lrrc8b
Gm10076
Mrpl42
Fbxl3
Adi1
Slc6a9
Rangap1
Frmd8
Sh3bgrl
Agap1
Hsd17b4
Trf
Smarcd1
Ola1
Abr
Dynlt3
Reep3
Mcee

Ntrk2
Arpc3
Ufd1
Ttyh2
Hnrnpf
Nsg1
Tubb3
Enpp2
Cuedc2
Capzb
Gprasp2
Tceal9
Snrpc
Eif3b
Derl1
Uqcrc1
Celf4
Dync1h1
Olfm1
Mfge8
Higd1a
Ndufa5
Gng2
Hp1bp3
Enpp5
Limch1
Pcmt1
Slc1a2
Opa1
Rnf187
Gng7
Adipor1
Gpr37
Atrn
Cmas
Trim59
Opalin
Metrn
Oxct1
Psmd8
Mapre1
Scnm1
Lap3
Arl3
BC005537
Nbea
Snx1
Itgav
Cdc42ep2
Adar
Rheb
Prmt1

Prkcsh
Epb41l1
Nkain4
Ktn1
Pdcd6
Smarca2
Bckdha
Fbxo44
Selenoh
Clcn4
Tmed9
Gnl1
Pja1
Kctd2
Mrpl54
Mag
Ube2e1
Glrx3
Kif21a
Gsk3b
Aig1
Diras2
Limd2
Snapc5
Sgta
Cfl2
Sirpa
Tra2b
Mrps34
Dnaja2
Nmral1
Pigq
Ap2m1
Nfix
Psmd2
Nucks1
Cltc
Flywch1
Pip4p1
Fcer1g
Phlpp1
Ddt
Copg1
Cd82
Psma4
Lpgat1
Plxdc2
Kazn
Tmem50b
Peg3
Ldha
Nrxn1

Slc25a5
Tex264
Tcaf1
Slc38a2
Atp6v0b
Smarcd3
Rad23a
Pdha1
Ech1
Vxn
Usp22
Ociad1
Pfn1
Gabbr1
Mrpl41
Cdc37
Rpn1
Nudt9
Chchd3
Ndufb8
Synj1
Emc8
Rhou
Dph3
Ankrd40
Hapln2
Coa3
Sema6a
Gabarapl1
Mrpl11
Pkp4
Chd3
Msrb1
P4hb
Trnp1
Cycs
St13
Cdk16
Plaat3
Ermp1
Tm7sf3
Dlg1
Tprn
Mrpl23
Efcab14
Ccdc12
Bmyc
Npc1
Cbr1
Wipi2
Mrps36
Ube2d1

Plxnb1
1500004A13Rik
Agpat3
Foxn3
Rab1b
Tsn
Apod
Pcyt2
C1qc
Myrf
Srp19
Psmd14
Trappc6b
Kidins220
Mydgf
Ehmt2
Rgs10
1190005I06Rik
Glul
Dctn6
Ssu72
Tsc22d4
Pfdn6
Nudt19
Ccdc47
Pdxp
Banf1
Sf1
Trim44
Emc1
Gpi1
Hdac5
Sap18
Agap3
Dzank1
Mrps24
Bmerb1
Rcbtb1
Polr1d
Ap3d1
Desi1
Nop53
Dlg3
Emc4
Mgrn1
1110004F10Rik
Cct8
Abhd17a
Ndufv1
Smarca4
Atp6v1c1
Mprip

Thrsp
Ndufb6
Mtpn
Ube2v1
Rnf220
Hspa9
Tubb4b
Grk2
Mtch2
Csrp1
Tafa5
Chchd1
Actr3
Ube2n
Ost4
Mrpl30
Jph4
Tmem63a
Smpd1
Pnck
Ablim2
Slc39a10
Mrps18c
Mast3
Ppid
Sesn3
Rnf13
Dpy19l1
Tomm22
Pebp1
Idh3g
Pttg1
Pon2
Lancl1
Rapgef4
Myl12a
Gnb1
Arsg
Ank3
Slc1a3
Ube2s
Nenf
Cdkn2d
Vps4a
Gng13
Arhgef10
Ank
Tmx2
Tmed7
Arf5
Azin1
C2cd2l

Csnk1g2
Pum2
Rbbp7
Tkt
Tspyl1
Ubqln2
Nono
Bod1
Hnrnpa0
Grb2
Atf4
Pfdn1
Arl8a
Dhrs1
Fbxl5
Mrps18a
Vps29
Bpgm
Acsbg1
Cyth1
Mapk1
S100b
Psmc5
Crmp1
Smim10l1
Kcna6
Stard10
Kifap3
Mrps12
Ptms
Mkrn1
Rbm39
Rpn2
Pdxk
Csnk1a1
Bcas2
Nbr1
Endod1
Ogt
Sort1
Prkaca
Ube2v2
Cdk5
Hagh
Mrpl48
Eif4a1
Acox1
Ubr4
Cntnap1
Nfasc
Ppp1r1a
Mrpl51

Prkacb
Zfp771
Vegfb
Polr2g
Ddx17
Srsf6
Gtf2i
Paics
Elovl1
Vps41

WMT cTBS

| DEGs Young sham vs Aged sham | cTBS DEGs unique to young adults | cTBS DEGs unique to young adults that are also ageing DEGs |
|---|---|---|
| Mobp | Lrrc17 | AI593442 |
| Mbp | Tatdn1 | Adgrl1 |
| Uba52 | Tspyl4 | Apod |
| Cabp1 | Ost4 | Arpc2 |
| Plekhb1 | Snrpf | Atp6v1a |
| Bcas1 | Phpt1 | Bloc1s1 |
| Chn1 | Snhg9 | Cadps |
| Hpca | Araf | Calb1 |
| Scn1b | Bloc1s1 | Cd81 |
| Bsn | Rps28 | Cdk5r2 |
| Slc17a7 | Snap25 | Celf5 |
| Tesc | Dcaf17 | Chd3 |
| Lingo1 | Ndufa1 | Clasp2 |
| Ppp3r1 | Car2 | Cnp |
| Nrgn | Rps27 | Cpe |
| Nptxr | Rps29 | Cryab |
| Mef2c | Rpl9-ps6 | Dcaf17 |
| Rpl9-ps6 | Zwint | Disp2 |
| Phf24 | Sparc | Dlg2 |
| Eef1a2 | Ppp1r16b | Dmtn |
| Sv2b | Scg2 | Dynlt3 |
| Gria3 | Gja1 | Dzank1 |
| Cacnb3 | Hectd4 | Eno1 |
| Lamp5 | Srsf5 | Epdr1 |
| Pvalb | Ptprz1 | Evl |
| Cadm3 | Actr2 | Ewsr1 |
| Mal | Snhg6 | Fbxl16 |
| Mt1 | Atp1a3 | Gad1 |
| Dnm1 | Kctd13 | Gapdh |
| Rprml | Slc6a1 | Gja1 |
| Camkk2 | Ubqln2 | Glul |
| Plk2 | Isca1 | Grk2 |
| Pi4ka | Rnf5 | Gsk3b |
| Trp53inp2 | Pacsin1 | Gsn |
| Atp1a1 | Tomm7 | Hectd4 |
| Fabp3 | Impact | Hnrnpa0 |
| Itpka | Smim26 | Homer1 |
| Tceal5 | Eif4a2 | Ina |
| Nat8l | Ndrg4 | Jph4 |
| Prr18 | Tsc22d4 | Khdrbs3 |
| Tmeff2 | Spag9 | Lmo4 |
| Tmsb4x | Atp6v1a | Lrrc17 |
| Tceal6 | Txn1 | Lynx1 |
| Bloc1s1 | Glul | Mag |
| Kcnab2 | Wdr6 | Magee1 |
| Ptk2b | Cox17 | Malat1 |
| Pdlim2 | Jph4 | Mapk9 |
| Pdp1 | Hmgb1 | Meg3 |
| Matk | Basp1 | Mog |

Lmo4
Clic4
Camk2b
Ppp3ca
Sri
Cnp
Rps29
Stx1a
Lin7b
Svop
Brinp1
Mmp17
Grin1
Kalrn
Snhg6
Cplx1
Rnf112
Pgm2l1
Calm1
Tspan2
Thy1
Spock1
Nell2
Gpr162
Cx3cl1
Pea15a
Fam131a
Cdk5r1
Ptprz1
Egr1
Golga7
Polr3e
Rell2
Cap2
Fxyd1
Snap25
Atp2b2
Pkm
Syn1
Kcnip2
Trak2
Habp4
Arf3
Snph
Myh10
Celf2
Marcksl1
Mrtfb
Slc6a17
Evi2a
Rbfox3
Itpr1
Chmp5
Sdhd
Snhg8
Sdcbp
Tmeff1
Atp5md
Zcchc12
Nedd4
Clasp2
Scn4b
Huwe1
Agap2
Fus
Selenof
Cdk5r2
Ddn
Cox7c
Dmtn
Camk2n1
Stxbp1
Slc4a4
Ank2
Psma5
Nrxn1
Tuba1b
Ndufa3
Sem1
Hprt
Ina
Gas7
Gsk3b
Polr2g
Peg3
Atp5g3
Id2
Snca
Usp50
Smarca2
Hnrnpa0
Pip5k1c
Sparcl1
Epdr1
Usp9x
Hpcal4
Hnrnpa2b1
Meg3
Pde10a
Paip2
App
Cadm4
Erh
Map2
Mycbp2
Myt1l
Ndrg4
Ndufb8
Necap1
Nrxn1
Opalin
Ost4
Pacsin1
Pdxk
Peg3
Phyhip
Pip5k1c
Polr2g
Ppp1r16b
Prkaca
Ptgds
Ptprz1
Rheb
Rian
Rit2
Rnf5
Rpl4
Rpl9-ps6
Rps12
Rps27
Rps28
Rps29
Scg2
Sdcbp
Slc44a1
Smarca2
Smim10l1
Snap25
Snca
Snhg11
Snhg6
Snhg9
Spag9
Spryd3
Stxbp1
Sucla2
Tatdn1
Tcaf1
Tmeff1
Tmx4
Tomm70a
Trf
Tsc22d4
Tsnax
Ubqln2

| | |
|---|---|
| Tppp3 | Phyhip |
| Syp | Id4 |
| Tmem132a | Rps27rt |
| Scrg1 | Cpox |
| Camk2a | Kif5c |
| Cyfip2 | Oxr1 |
| R3hdm1 | Rraga |
| Ier5 | Phyhipl |
| Prkcg | Magee1 |
| Cplx2 | Bzw1 |
| Cck | Prpf8 |
| St3gal5 | Slc25a22 |
| Rasgef1a | Cystm1 |
| Gnb5 | Tmx4 |
| Calb1 | Rheb |
| Carhsp1 | Dzank1 |
| Snhg9 | Rian |
| Zbtb18 | Calb1 |
| Nrxn3 | Cops7a |
| Fa2h | Snhg11 |
| Gls | Mcam |
| Sccpdh | Grk2 |
| Lynx1 | Ndfip2 |
| Gabrg2 | Sec61g |
| Car11 | Psip1 |
| Ncs1 | Prkaca |
| Myo6 | Necap1 |
| Gamt | B230312C02Rik |
| Cacng3 | Emc7 |
| Bcat1 | Fbxl16 |
| Phgdh | Cacybp |
| Baiap2 | Rit2 |
| Rab31 | Disp2 |
| Plcb1 | Unc80 |
| Ak3 | Arpc2 |
| Pianp | Idi1 |
| Jph3 | F3 |
| Reep2 | Mgat3 |
| AI593442 | Acyp2 |
| Ugt8a | Bpnt1 |
| Tns3 | Gad1 |
| Adcy1 | Evl |
| Tac1 | Med30 |
| Serpini1 | Rtl8a |
| Nsf | Mapre2 |
| Tmem59l | Rab28 |
| Mog | Tsnax |
| Josd2 | Prkar1a |
| Myt1l | Naa20 |
| Grina | Bcl2l2 |
| Madd | Gm20594 |
| Timm22 | Adgrl1 |

| | |
|---|---|
| Gapdh | Prrc2a |
| Pcsk2 | Gtf2a1 |
| Snx10 | Sucla2 |
| Napb | Dynlt3 |
| Mapk10 | Smim10l1 |
| Tpi1 | Frrs1l |
| Sncb | Atp6v1h |
| Litaf | Mapk9 |
| Ywhah | Rab3c |
| Dgkz | Pdxk |
| Dbi | Homer1 |
| Grik5 | AI593442 |
| Tbc1d9b | Lynx1 |
| Tatdn1 | Morf4l1 |
| Tmeff1 | Khdrbs3 |
| Cpe | Cadps |
| Icam5 | Tomm70a |
| Gng12 | Chd3 |
| Snca | Fmc1 |
| BC004004 | Lmo4 |
| Ddah1 | Tcaf1 |
| Cmtm5 | Abhd3 |
| Rgs9 | Mycbp2 |
| Rasgrp1 | Myt1l |
| Syngr3 | Pkia |
| Nfe2l1 | Ip6k1 |
| Rps28 | Dlg2 |
| Dcaf17 | Arpc4 |
| Setd7 | Cpe |
| Ptprn | Inpp4a |
| Igfbp5 | Wdr89 |
| Peg13 | Celf5 |
| St8sia3 | Spryd3 |
| Grb14 | Ewsr1 |
| Cdk2ap1 | Snrpn |
| Cldn11 | Atn1 |
| Tusc3 | Rps19 |
| Gabrb3 | Eif1 |
| Ckmt1 | Ndufb9 |
| Gnal | Ndufb8 |
| Paqr8 | Smdt1 |
| Tspan13 | Rpl23a |
| Slc2a1 | Rpl27a |
| Syt4 | Cd81 |
| Qdpr | Rpl5 |
| Dmtn | Rpl7 |
| Sh3gl3 | Rps6 |
| Ugp2 | Rpl17 |
| Ppp1r9a | Aplp1 |
| Dbn1 | Cnp |
| Cat | Rps15a |
| Atp6v0e | Rplp1 |

| | |
|---|---|
| Phyhip | Mog |
| Cdc37l1 | Eno1 |
| Rnf157 | Rps2 |
| Aak1 | Slc44a1 |
| Mgst3 | Gapdh |
| Lamp1 | Rpl4 |
| Aktip | Pacs2 |
| Tsnax | Rpl9 |
| Arl6ip5 | Rps3a1 |
| Dock10 | Fkbp2 |
| Gabbr2 | Tubb4a |
| Kcnj4 | Rpl8 |
| Rps27 | Rps15 |
| Arhgef9 | Ptma |
| Ptpn11 | Rpl32 |
| Vbp1 | Opalin |
| Rab15 | Pfdn5 |
| Ncor2 | Rpl35a |
| Gnai1 | Eef1b2 |
| Tmem47 | Rps11 |
| Ewsr1 | Ptgds |
| Enc1 | Rpl18 |
| Ptpn5 | Tpt1 |
| Unc50 | Rps14 |
| Eno2 | Rpl6 |
| Nkiras1 | Rpl26 |
| Id3 | Rpl23 |
| Pip5k1c | Rps16 |
| Tra2a | Rpl28 |
| Fhl1 | Rps4x |
| Myo5a | Trf |
| Aqp4 | Rps8 |
| Cd9 | Gjb1 |
| Neurl1a | Rpl24 |
| Reps2 | Rpl13 |
| Rasl10b | Rpl11 |
| Stmn3 | Rpl18a |
| Anp32b | Rpl30 |
| Efhd2 | Cryab |
| Bex2 | Rps12 |
| Ppp2r5a | Rps5 |
| Rusc1 | Rps27a |
| Cadps | Rps10 |
| Cfap36 | Rps7 |
| 2310022B05Rik | Rps20 |
| Scg2 | Rpl14 |
| Rab3a | Mag |
| Ivns1abp | Rps3 |
| Dlgap1 | Gsn |
| Syt13 | Rpl13a |
| Rpl4 | Fau |
| Mt2 | Rpl21 |

Tnks2
Oip5os1
Rasd2
Homer1
Strn4
Ywhaz
Dock3
Tmco1
Nefm
Tubb5
Tmem179
Evl
Fyn
Kcnq2
Rab6b
Dbp
Chga
Tuba4a
Plekhj1
Zfas1
Clstn3
Ednrb
Syn2
Add2
Dlgap4
Gsn
Kndc1
Pja2
Got2
Celf5
Cryab
Serinc5
Ddit3
Daam2
Rhog
Cadm2
Shtn1
Dgkb
Vamp2
Churc1
Rps12
Lrrn1
Mapk9
Sar1b
Adgrb2
Prkcb
Map4k4
Unc13a
Adgrl1
Kcnk1
Atp6v1b2
Zfand5
Rps24
Apod
Rpl10
Malat1
Hba-a2
Hba-a1
Hbb-bs
Ttr

Pip4k2a
Otub1
Scd1
Asphd2
Sptbn2
Egr3
Fgf1
Idh3a
Fads2
Enah
Cnn3
Atxn7l3
B3gat3
Usp14
Arrdc3
Stk25
Scg5
Aamdc
Camta2
Scn2a
Cntn2
Sorl1
Anapc2
Nptn
Rdx
Nsg2
Wnk1
Stxbp1
Dkk3
Sc5d
C1qtnf4
Gal3st1
Ap1b1
Cyb5b
Fam234b
Zfp664
Ptprd
Kctd17
Got1
Dclk1
Prkce
Tcf4
Actr1b
Rnf130
Meg3
Atp6v0c
Picalm
Clstn1
Gpr88
Rad21
Elovl5
Adgrb1

Rian
Chst1
Nat14
Ralgds
Rps27l
Astn1
B230219D22Rik
Wasf1
Ogdh
Epb41l2
Pltp
Syt5
Fxyd7
Abhd4
Txndc9
Necap1
Ankrd13a
Plekhh1
Ndrg1
Ado
Snhg11
Plp1
Aspa
Med21
Smim13
Cdc42se2
Arhgap23
Malat1
Chn2
Ddr1
Taf13
Eif6
Mllt11
Nrbp2
Ctnna1
Tprkb
Rnf227
Olig2
Pdxdc1
Ap2a2
Strn
Nrcam
S1pr5
Vgf
Mmd
Cd81
Pou3f3
Sirt2
Rph3a
Nefl
Olig1
Cd164

Plekha1
Cdk17
Gstm7
Myl12b
Scarb2
Nhp2
Srpk2
Srsf11
Ppp2r3a
Efhd1
Gjc2
Snn
Ptk2
Pdhb
Mras
Jam3
Hepacam
Hadha
Atp6v1e1
Nacc2
Gltp
Sh2d5
Micall1
Snap91
Arpc1b
Khdrbs3
Pop5
Slc48a1
Mrpl17
Arhgdig
Fdx2
Snapin
Gabra1
Pam16
Eno1
Ufm1
Rnps1
Rtkn
Haghl
Timm23
Arpp21
Tmem134
Igfbp6
Smap2
Mrpl4
Zcchc24
Mrps26
Eci1
Adcyap1r1
Cers2
Strbp
Arhgap5

Tmem229a
Phf5a
Asns
Erbin
Ddx24
Pop4
Phlda3
Lrp1
Trim9
Pim3
Pik3r1
Pfkl
1700047M11Rik
Cxcl14
Gnptg
Slc44a1
Rpl37a
Ccdc124
Ina
Gjc3
Pdcd10
2900052N01Rik
Tusc2
Mat2b
Stard7
Dync1i1
Wscd1
Lrrc58
Kifc2
Rpl35
Sdcbp
Atp1b1
Usp5
Aldh6a1
Sptbn4
Anxa5
Adgrg1
Schip1
Ybx3
Fez1
Ifi27
Tmbim4
Ssbp4
Selenop
Hras
Trim37
Fbxl16
Rogdi
Syt7
Csnk1d
Ten1
Adk

Nfu1
Lpar1
Prkar1b
Hnrnpdl
Lamp2
Slc25a11
Camkv
Rnf181
Ppa1
Sult4a1
Zer1
Gas5
B4gat1
Hcn2
Gpbp1
Atp6v1g2
Mtus1
Snx5
Tmem208
Arhgef4
Sh3gl2
Calm2
Phldb1
Sptssa
Emc3
Wipf3
Hypk
Ccl27a
Spock2
Ei24
Prxl2b
Egln1
Commd4
Ttc9b
Znhit1
Paqr4
Apc
Cdipt
Arhgap33
Gria1
Oaz2
Sox2ot
Mapk8ip2
Pttg1ip
Cdk5r2
Pld3
Ssr3
Rbm3
Svip
Mrpl12
Pacsin1
Trim32

Map4
S1pr1
Ube2ql1
Rgs4
Prrt1
Clcn3
Map7
Ctxn1
Smim11
Nrsn2
Sugt1
Gda
Pde6d
Kcnab1
Rap1b
Sec13
Kdelr1
Usp54
Ndrg4
Rbbp4
Slain1
Dync1i2
Grhpr
Gm2a
Kif1c
Atp2b1
Ngef
Gstp1
Atg3
Echs1
Secisbp2l
Ppp1r2
Necab2
Ctdsp2
Ppt1
Cox5a
Plcl1
Gsk3a
Dnm1l
Tmem88b
Ppp2r2a
Dnajb2
Arhgef17
Commd6
Zfp207
Prmt2
Dctn4
Sulf2
Cavin3
Ociad2
Ncoa4
Car14

Rnf208
Mboat7
Chchd6
Gna12
Chgb
Cbx6
Lrrc17
Tfg
Dixdc1
Vkorc1
Scn2b
Tomm70a
Dnajc15
Kif5a
Tmem33
Ddx1
Tspan5
Ermn
Kcnj10
Il18
Nipa1
Sbf1
Sbds
Polr3h
Plin3
Gng11
Klhdc3
Bcap31
Hnrnph2
Pogk
Epas1
Iqsec1
Klf13
Camta1
Btbd1
Add3
M6pr
Gng3
Sumo3
Sema4d
Nrxn2
Slc12a2
Atp6v1a
Clk1
Gcsh
Sucla2
Psmc1
Stx1b
Pak2
Herc1
Atp6v1d
Ppp1r14b

Eif3e
Snhg12
Ncdn
Mtss2
Grn
Efnb3
Oxsr1
Spock3
Snrnp27
Ppp2r1a
Eml1
Kif3c
Syf2
Tmx4
Dlg4
Clip2
Disp2
Ypel5
Rufy3
Rit2
Bnip3
Sox10
Ppp1r11
Fnta
Scrn1
Map2k1
Papss1
Hectd4
Sh3glb1
Psph
Vstm2l
Adh5
Prpf19
Dip2a
Sfpq
Ly6e
Nptx1
2210016L21Rik
Serpinb6a
Slc25a12
Prrc2b
5031439G07Rik
Polr2l
Rnf5
Tmem98
Hk1
Pgam1
Zmiz2
Rbbp6
Tmem160
Scamp1
Slc24a2

Pcna
Eif1ax
Cyp46a1
Gja1
Dtx3
Slc25a23
Mbnl1
Agpat4
Rida
Pkig
Clmn
Mycbp2
Dcaf8
Cpd
Rasgrf1
Mink1
Camk2g
Tmcc3
Gabrd
Spin1
Fgfr2
Gclm
Dlgap3
Syngr1
Ptgds
Nsfl1c
Fnbp1
Hnrnpr
Retreg2
Lxn
Rap1a
Tmed4
Spryd3
Tmcc2
Pcif1
Arrb1
Slc35b1
Arl1
Magee1
Ubald1
Sh3glb2
Trappc2l
Paip2b
Akr7a5
Nckap1
Maf1
Ppme1
Hnrnph3
Amer2
Tmem158
Fam168b
Epdr1

Abhd17b
Fbxo9
Podxl2
Trpc4ap
Glrx5
Arpc2
Kctd3
Trim3
Etv1
Hexa
Dnm2
Vamp3
Baalc
Cldnd1
Tpd52
Jup
Dnajc5
Actr10
Ppm1a
Ctnnd2
Commd1
Dtna
Acot13
Napg
Mcts1
Spag9
Ddit4
Stard3nl
Rexo2
Rnf141
Mtmr6
Gab1
Metap2
Mtfr1l
Snap47
S100a6
Mrpl53
Ly6a
Alcam
Anln
Taok1
Polr2c
Psd3
Stk11
Dlg2
Kmt5a
Celsr2
Tmod1
Psma1
Pfkp
Ppp2r5c
Adipor2

Cxxc5
Gps1
Zeb2
Gprasp1
Epn2
Ndufa4
Cyb5r3
Slu7
Cdk19
Cdk2ap2
Ufc1
Eif3a
Tulp4
Ap1s1
Cenpb
Abi2
Asrgl1
Znrf1
Hnrnph1
Bag6
Cs
Ptprs
Fkbp8
Tspan15
Smox
Txnl1
Ccp110
Ensa
Pdk2
Ift20
Nsa2
Gprc5b
Slc32a1
Dner
Dnlz
Tmbim1
Srsf1
Hopx
Ppp1r14a
Abhd8
Txnl4a
Rnh1
Enoph1
Ttc3
Lzts2
Tpm1
Csnk2b
Hdhd2
Gad1
Cops6
Mapre3
Tnfaip6

Anp32e
Psat1
Tsg101
Tpd52l2
Epn1
Rad23b
Mrpl28
Gfap
Mrps14
Rgs7bp
Gap43
Itpk1
Gpr62
Il33
Mrpl24
Nkx6-2
Parp6
Pin1
Mrfap1
Itm2c
Cbarp
Pdcd5
Tcf25
Vdac1
Clasp2
Pllp
Ndufb5
Mapk8ip3
Eif4a3
Ppp1r16b
Uqcc2
Rplp0
Lrpap1
Adcy5
Lrrc8b
Gm10076
Mrpl42
Fbxl3
Adi1
Slc6a9
Rangap1
Frmd8
Sh3bgrl
Agap1
Hsd17b4
Trf
Smarcd1
Ola1
Abr
Dynlt3
Reep3
Mcee

Ntrk2
Arpc3
Ufd1
Ttyh2
Hnrnpf
Nsg1
Tubb3
Enpp2
Cuedc2
Capzb
Gprasp2
Tceal9
Snrpc
Eif3b
Derl1
Uqcrc1
Celf4
Dync1h1
Olfm1
Mfge8
Higd1a
Ndufa5
Gng2
Hp1bp3
Enpp5
Limch1
Pcmt1
Slc1a2
Opa1
Rnf187
Gng7
Adipor1
Gpr37
Atrn
Cmas
Trim59
Opalin
Metrn
Oxct1
Psmd8
Mapre1
Scnm1
Lap3
Arl3
BC005537
Nbea
Snx1
Itgav
Cdc42ep2
Adar
Rheb
Prmt1

Prkcsh
Epb41l1
Nkain4
Ktn1
Pdcd6
Smarca2
Bckdha
Fbxo44
Selenoh
Clcn4
Tmed9
Gnl1
Pja1
Kctd2
Mrpl54
Mag
Ube2e1
Glrx3
Kif21a
Gsk3b
Aig1
Diras2
Limd2
Snapc5
Sgta
Cfl2
Sirpa
Tra2b
Mrps34
Dnaja2
Nmral1
Pigq
Ap2m1
Nfix
Psmd2
Nucks1
Cltc
Flywch1
Pip4p1
Fcer1g
Phlpp1
Ddt
Copg1
Cd82
Psma4
Lpgat1
Plxdc2
Kazn
Tmem50b
Peg3
Ldha
Nrxn1

Slc25a5
Tex264
Tcaf1
Slc38a2
Atp6v0b
Smarcd3
Rad23a
Pdha1
Ech1
Vxn
Usp22
Ociad1
Pfn1
Gabbr1
Mrpl41
Cdc37
Rpn1
Nudt9
Chchd3
Ndufb8
Synj1
Emc8
Rhou
Dph3
Ankrd40
Hapln2
Coa3
Sema6a
Gabarapl1
Mrpl11
Pkp4
Chd3
Msrb1
P4hb
Trnp1
Cycs
St13
Cdk16
Plaat3
Ermp1
Tm7sf3
Dlg1
Tprn
Mrpl23
Efcab14
Ccdc12
Bmyc
Npc1
Cbr1
Wipi2
Mrps36
Ube2d1

Plxnb1
1500004A13Rik
Agpat3
Foxn3
Rab1b
Tsn
Apod
Pcyt2
C1qc
Myrf
Srp19
Psmd14
Trappc6b
Kidins220
Mydgf
Ehmt2
Rgs10
1190005I06Rik
Glul
Dctn6
Ssu72
Tsc22d4
Pfdn6
Nudt19
Ccdc47
Pdxp
Banf1
Sf1
Trim44
Emc1
Gpi1
Hdac5
Sap18
Agap3
Dzank1
Mrps24
Bmerb1
Rcbtb1
Polr1d
Ap3d1
Desi1
Nop53
Dlg3
Emc4
Mgrn1
1110004F10Rik
Cct8
Abhd17a
Ndufv1
Smarca4
Atp6v1c1
Mprip

Thrsp
Ndufb6
Mtpn
Ube2v1
Rnf220
Hspa9
Tubb4b
Grk2
Mtch2
Csrp1
Tafa5
Chchd1
Actr3
Ube2n
Ost4
Mrpl30
Jph4
Tmem63a
Smpd1
Pnck
Ablim2
Slc39a10
Mrps18c
Mast3
Ppid
Sesn3
Rnf13
Dpy19l1
Tomm22
Pebp1
Idh3g
Pttg1
Pon2
Lancl1
Rapgef4
Myl12a
Gnb1
Arsg
Ank3
Slc1a3
Ube2s
Nenf
Cdkn2d
Vps4a
Gng13
Arhgef10
Ank
Tmx2
Tmed7
Arf5
Azin1
C2cd2l

Csnk1g2
Pum2
Rbbp7
Tkt
Tspyl1
Ubqln2
Nono
Bod1
Hnrnpa0
Grb2
Atf4
Pfdn1
Arl8a
Dhrs1
Fbxl5
Mrps18a
Vps29
Bpgm
Acsbg1
Cyth1
Mapk1
S100b
Psmc5
Crmp1
Smim10l1
Kcna6
Stard10
Kifap3
Mrps12
Ptms
Mkrn1
Rbm39
Rpn2
Pdxk
Csnk1a1
Bcas2
Nbr1
Endod1
Ogt
Sort1
Prkaca
Ube2v2
Cdk5
Hagh
Mrpl48
Eif4a1
Acox1
Ubr4
Cntnap1
Nfasc
Ppp1r1a
Mrpl51

Prkacb
Zfp771
Vegfb
Polr2g
Ddx17
Srsf6
Gtf2i
Paics
Elovl1
Vps41
